# Effects of phosphorylation on protein backbone dynamics and conformational preferences

**DOI:** 10.1101/2024.02.15.580491

**Authors:** David Bickel, Wim Vranken

## Abstract

Phosphorylations are the most common and extensively studied post-translational modification (PTM) of proteins in eukaryotes. They constitute a major regulatory mechanism, modulating protein function, protein-protein interactions, as well as subcellular localization. Phosphorylation sites are preferably located in intrinsically disordered regions and have been shown to trigger structural rearrangements and order-to-disorder transitions. They can therefore have a significant effect on protein backbone dynamics or conformation, but only sparse experimental data are available. To obtain a more general description of how and when phosphorylations have a significant effect on protein behavior, molecular dynamics (MD) currently provides the only suitable framework to study these effects at a large scale in atomistic detail.

This study develops a systematic MD simulation framework to explore the influence of phosphorylations on the local backbone dynamics and conformational propensities of proteins. Through a series of glycine-backbone peptides, we studied the effects of amino acid residues including the three most common phosphorylations (Ser, Thr, and Tyr), on local backbone dynamics and conformational propensities. We further extended our study to investigate the interactions of all such residues between position *i* to positions *i*+1, *i*+2, *i*+3, and *i*+4 in such peptides. The final dataset is comprised of structural ensembles for 3,393 sequences with more than 1 µs of sampling for each ensemble. To validate the relevance of the results, the structural and conformational properties extracted from the MD simulations are compared to NMR data from the Biological Magnetic Resonance Data Bank.

The systematic nature of this study enables the projection of the gained knowledge onto any phosphorylation-site in the proteome and provides a general framework for the study of further PTMs. The full dataset is publicly available, as a training and reference set.

## Introduction

Phosphorylations are the most common and well-studied post-translational modification (PTM) in proteins. They play a key role in regulating a variety of cellular processes, like cellular signaling,^1^ cell-cycle regulation,^2^ phase separation,^3^ and subcellular localization.^4^ Dysregulation of phosphorylation is associated with diseases such as cancer and neurodegeneration. The phosphorylation of the amino acid serine (Ser) is the most common, followed by threonine (Thr) and tyrosine (Tyr).^5^ Phosphorylation of histidine, arginine, lysine, aspartate, glutamate, and cysteine also occurs but only infrequently.^6,7^

Protein phosphorylation is a reversible process during which a phosphoryl group is attached to the side chain of an amino acid. This phosphorylation is mediated by protein kinases while the dephosphorylation is mediated by phosphatases. The attachment of the phosphoryl group significantly changes the local amino acid environment, most notably by introducing a negative charge and by increasing the volume of the amino acid side chain. The latter effect is most notable for phosphorylation sites with small side chains, like Ser and Thr, where the amino acids’ volume is changed by a factor of ∼1.5.^8^ In addition, the phosphate group introduces constitutive negative charges to the primarily neutral amino acids. With p*K*_a_ values of 2 and 5.8-6.3 the phosphoryl group’s di-anionic form is the most common at physiological pH, being about 10-times more prevalent than the mono-anionic form.^8,9^ Thus, the equilibrium between the mono- and di-anionic forms of the phosphoryl group are strongly subject to the cellular microenvironment and the subcellular localization.

Phosphorylation sites are typically found in intrinsically disordered regions (IDRs) of proteins and flexible residues are substantially enriched around phosphorylation sites, with an estimated 65% of all phosphorylations occurring in such lowly structured regions (Ser: ∼70%, Thr: ∼60%, Tyr: ∼40%).^10,11^ It is hypothesized that such flexible regions facilitate access for kinases, which transfer phosphate groups to the amino acid side chain.^12^ In addition, IDRs commonly contain sequence motifs involved in protein recognition, so called short linear motifs (SLiMs). Phosphorylations in such motifs can be mediated by kinase recognition motifs, so further modulating a protein’s interactions with partners.^8,12^ Further complexity is introduced by the ability of phosphorylation to induce significant changes in a protein’s conformational preferences and dynamics, even resulting in folding of IDRs.^8,13^ As a means to bring about functional changes in a protein, phosphorylations are able to modulate the protein’s conformational landscape. The protein structures that have been resolved by X-ray crystallography in the phosphorylated and the native state do capture conformational changes,^14^ but are limited to static structures and fail to capture more gradual changes in conformational preferences in IDRs and IDPs. In contrast, Nuclear Magnetic Resonance (NMR) spectroscopy is well suited to observe significant changes in the overall behavior of IDRs upon phosphorylation, often with detailed experiments dedicated to capture dynamics.^15^ Such changes are also encompassed by NMR chemical shifts, which can be interpreted to infer information on a protein’s conformation and dynamics at residue resolution.^16–18^ Chemical shifts are especially useful as they are readily publicly available from the Biological Magnetic Resonance Data Bank (BMRB)^19^ and therefore amenable for large scale analysis. Circular dichroism (CD) spectroscopy can also be used to capture phosphorylation-induced changes by measuring changes in secondary structure content, including order-disorder transitions.^20^ Finally, small-angle X-ray or neutron scattering (SAXS/SANS) provides low-resolution information on the shape and compactness of proteins, again capturing large conformational changes.^21^

Despite the various techniques to study the impact of phosphorylation on protein behavior, experimental data remains sparse. The gained insights pertain to single proteins and while highly interesting, it is unclear to which extent and under which conditions the observed effects induced by phosphorylation can be generalized. In this work we use extensive molecular dynamics (MD) simulations to generate a conformational ensemble dataset that systematically explores the effects of phosphorylations of Ser, Thr and Tyr on local backbone dynamics. Similar yet more limited studies on regular peptides have been carried out successfully before, providing valuable insights into their dynamics and intrinsic conformational preferences.^22–24^ This study aims to expand on these studies, with the underlying reasoning that small changes in conformational preferences in peptides indicate differences in side-chain interactions and/or dynamics that remain relevant for proteins, especially when intrinsically disordered. A detailed analysis of the generated MD dataset indeed reveals consistent conformational effects of phosphorylations, which are validated against publicly available NMR data from the BMRB and compared to prior studies. The full conformational data set is made publicly available as described under **Data Availability**.

## Materials and Methods

The code used to run the simulations and perform the analyses described below has been made available on GitHub (see **Data availability**).

### Preparation of molecular systems

To prepare the 3-dimensional structures from the peptide sequences, we used the PeptideBuilder library,^25^ which we extended by the additional residues in **Table 2** as well as acetyl (ACE) and *N*-methylamide (NME) capping groups. For the scaffold, glycine residues were chosen since glycine, due to its lack of side chains, features the least restricted conformational free energy landscape of all amino acids and thus represents a neutral environment to study the effect of other residues on each other. For each sequence we built four conformations with differing backbone dihedrals (*φ*, *ψ*) as starting points for independent simulations (**Suppl. Table 1**). The dihedrals were chosen to minimize intramolecular hydrogen bonds, allowing the peptides to rapidly explore conformational states unrelating to their starting conformation.

### Molecular dynamics simulations

#### Force field selection

A common problem for MD simulations on phosphorylated amino acids is that the phosphate parameters of many force fields are not well validated, due to the very limited experimental data that is available. Moreover, phosphorylations show different effects in different force fields.^26^ Recent studies comparing multiple force fields found the Amber-family ff99SB-ILDN force field^27–29^ in combination with TIP4P-D water^30^ to best reproduce the limited experimental data, which we therefore chose for this study.^31–33^

#### Simulation setup

For each peptide sequence four independent replica simulations were run from different starting conformations (see above). The peptides were solvated with TIP4P-D water^30^ in a rhombic-dodecahedral periodic boundary box with box vector lengths dependent on the peptide size: 4.2 nm for pentapeptides, 4.4 nm for hexapeptides, 4.7 nm for heptapeptides, and 5.0 nm for octa- and nonapeptides. Sodium and chloride ions were added up to a concentration of 0.15 mol l^-1^ neutralizing potential net charges on the peptides. The ff99SB-ILDN force field was used for the peptides with parameters for phosphorylated residues from Homeyer *et al.* and Steinbrecher *et al.*.^27–29^ The simulations were carried out using GROMACS v. 2021.3. The leap-frog algorithm was used with an integration time step of 2 fs. Nonbonded interactions were treated with a Verlet list cutoff scheme with a 1.0 nm cutoff. Long-range dispersion correction was applied to energy and pressure. Particle mesh Ewald method was used to treat long range electrostatic interactions with a gridspacing of 0.12 nm. The LINCS algorithm was used to constrain bonds with hydrogen atoms.

#### System equilibration

Each system was first subjected to three iterations of energy minimization for 10,000 steps of steepest descend. First only solvent atoms were minimized applying restraints of 1000 kJ mol^-1^ nm^-2^ on all peptide atoms. Then, only the solutes were minimized applying the same restraints to the water atoms. Finally, a minimization without restraints was done. Over 50 ps of NVT simulation, the temperature was stabilized around 298 K using velocity-rescaling with separate heat bath couplings for the peptide and the solvent.^34^ Thereafter, the pressure was equilibrated to 1 bar over 250 ps NPT simulation with the Berendsen barostat,^35^ using a time constant of 0.5 ps and 4.5·10^−5^ bar^−1^ isothermal compressibility.

#### Production simulations

Finally, the production simulations were performed without any restraints. For temperature control we used the Nosé-Hoover thermostat at 298 K with separate heat bath couplings for the peptide and the solvent.^36,37^ For the pressure the Parrinello-Rahman barostat was used with a relaxation time of 2 ps, and 4.5·10^−5^ bar^−1^ isothermal compressibility.^38,39^ Initially, we simulated a few pentapeptides for more than 1 µs per replica to analyze how quickly the conformational sampling convergences (see **Analysis of convergence**). We found that most simulations converged after 300 ns of sampling per replica. Thus, the remaining systems were run for 300 ns pre replica (1.2 µs cumulative sampling) by default. Simulations that showed poor convergence after that time were further extended to up to 900 ns (3.6 µs cumulative sampling).

### Analysis of the molecular dynamics simulations

For all analyses except the hydrogen bond analysis water molecules and ions were stripped from the trajectories. Replica simulations of the same peptides were concatenated and analyzed as a single trajectory (though *post hoc* separation of the data points into the individual replicas remained possible). The peptides were imaged and centered to avoid breaks across the periodic boundary box using the *gmx trjconv* module.

#### Backbone dihedrals

The (*φ*, *ψ*) backbone dihedrals were extracted as a time series for all residues over the course of the simulation using the *MDAnalysis* python package.^40^ To compare the dihedral profiles between different residues and peptides, the (*φ*, *ψ*) angle pairs were binned into 9° × 9°-bins (resulting in 1,600 bins). The dihedral pair counts in the bins were then normalized by the total number of dihedral pairs, such that *f*_*X*_(*i*, *j*) denotes the fraction of angle pairs for residue *X* that fall into the bin *ij*. Consequently, differences between the dihedral profiles of two residues *X*_1_ and *X*_2_can be calculated as root mean square deviations (RMSD) of the relative bin populations, as described in **equation 1**.

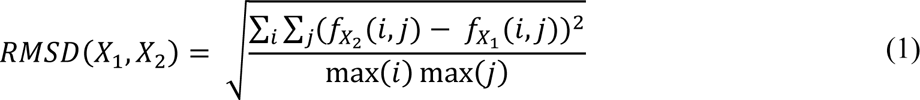

Further the circular variance can be calculated as a measure of diversity of the (*φ*, *ψ*) angle pairs as described in **equation 2**.

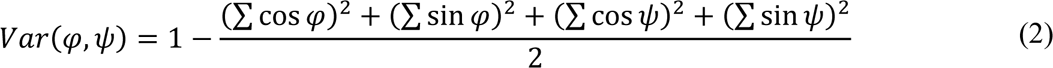

#### Analysis of convergence

The backbone dihedral angles (*φ*, *ψ*) were calculated for the stripped and imaged trajectories using the *gmx angle* module. To perform the principal component analysis (PCA) the angles were transformed into continuous cartesian space by storing their sines and cosines, so that an individual residue’s conformation is represented as a point in a four-dimensional space (cos(*φ*), sin(*φ*), cos(*ψ*), sin(*ψ*)), and the backbone conformation can be represented in 4 * *n*_residues_ dimensions. The eigenvectors and eigenvalues were calculated across all replica simulations using *gmx covar*. Finally, projections for each replica individually along the eigenvectors were calculated using *gmx anaeig.* To compare the projections along the eigenvectors of the individual replica simulations, only the first five eigenvectors were chosen and binned along their variance ranges into five equally sized bins resulting in a total of 3,125 combinations of bins per peptide. This permitted for some variation in the bins, so allowing conformationally related structures to end up in corresponding bins. Finally, the replica correlation was calculated for each pair of replicas (*a*, *b*) by calculating the correlation coefficient between the populations of the individual bins (*a_k_*, *b_k_*) using **equation 3**. Bins that were not populated in either of the replicas *a* and *b* were not included in the calculation of the correlation coefficient. Including such unpopulated bins leads to an overestimation of correlation coefficients in short trajectories when only a small portion of the phase space has been sampled yet.

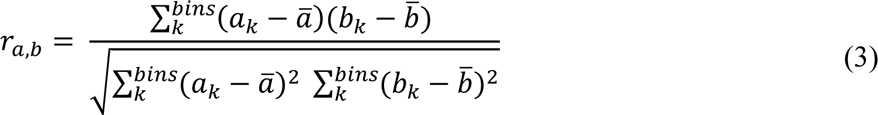

#### Conformational state propensities

The conformational state propensities of the conformational ensembles were calculated using the Constava software package,^41^ v. 0.1.0, using the default conformational state models and sliding-window subsampling with a window size of 3. For the comparison to chemical shift-derived secondary structure propensities, the helix properties (*core helix*, *surrounding helix*) as well as the sheet propensities (*core sheet*, *surrounding sheet*) were added up. Moreover, the values were smoothed using a weighted moving average (width: 3, weights: [0.25, 0.50, 0.25]).

#### Hydrogen bond network analysis

From the trajectories, hydrogen bonds were recorded using the *MDAnalysis* python package.^40^ Hydrogen bonds were defined as having a hydrogen bond donor-acceptor distance of less than 3 Å and a donor-H→acceptor angle of greater 150°.

### Validation against NMR data

#### Data collection

The chemical shift data were obtained from the BMRB.^19^ We queried the database for entries with phosphorylated residues identified by containing residues named ’SEP’, ’TPO’, or ’PTR’ resulting in 120 entries. We then used BLAST,^42^ v. 2.13.0, to identify BMRB entries with high sequence identity that were non-phosphorylated. The BLAST hits were further filtered to have 100% sequence identity around the phosphorylation site (±8 residues). This resulted in 144 paired entries. For each entry the S^2^_RCI_ (random coil index-derived S^2^ order parameter) was calculated using **equations 4** and **5**.^43,44^ The secondary structure propensities determined using δ2D.^17^

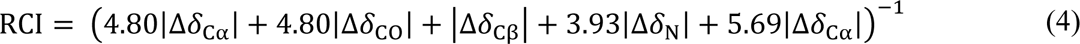

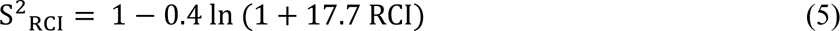

#### Comparison to MD simulations

The paired BMRB entries were aligned along their sequences. Entries with S^2^_RCI_ > 0.8 around the phosphorylation site were excluded. For each matched residue pair the differences in S^2^_RCI_ and secondary structure propensities between the phosphorylated and unphosphorylated state were calculated.

## Results

We conceived a systematic MD framework to study the effects of individual residues and their local interactions on protein dynamics and conformational propensities. In this setup, a series of glycine peptide scaffolds are used for all simulations and only one to two amino acid residues are substituted to investigate their respective and joint effects on the conformational sampling (**Table 1**). The simulations included all canonical amino acids as well as physiologically relevant alternative protonation states and the three most commonly occurring phosphorylations, leading to nine additional non-standard amino acids (**Table 2**). In total we generated 3,393 peptides in which we studied the individual residues’ effects as well as binary interactions in positions *i*+1, *i*+2, *i*+3, and *i*+4.

**Table 1.** Overview of simulated peptides. Configurations of the peptides simulated in this study. X_1_ and X_2_ are wildcard residues that were independently exchanged by any of 29 amino acids investigated in this study. These include the 20 canonical amino acids, alternate protonation states for Asp, Glu, and His, as well as phosphorylated Ser, Thr, and Tyr in mono- and diionic form. Ace refers to a N-terminal N-Acetyl cap. Nme refers to a C-terminal N-Methyl amide cap. Each peptide was simulated in four independent replicas.

| Length | Peptide scaffolds | Combinations | Total |
| --- | --- | --- | --- |
| 5 | Ace-GGX<sub>1</sub>GG-Nme | 29 | 29 |
| 6 | Ace-GGX<sub>1</sub>X<sub>2</sub>GG-Nme | 29 * 29 | 841 |
| 7 | Ace-GGX<sub>1</sub>GX<sub>2</sub>GG-Nme | 29 * 29 | 841 |
| 8 | Ace-GGX<sub>1</sub>GGX<sub>2</sub>GG-Nme | 29 * 29 | 841 |
| 9 | Ace-GGX<sub>1</sub>GGGX<sub>2</sub>GG-Nme | 29 * 29 | 841 |
|  |  |  | **3,393** |

**Table 2:**
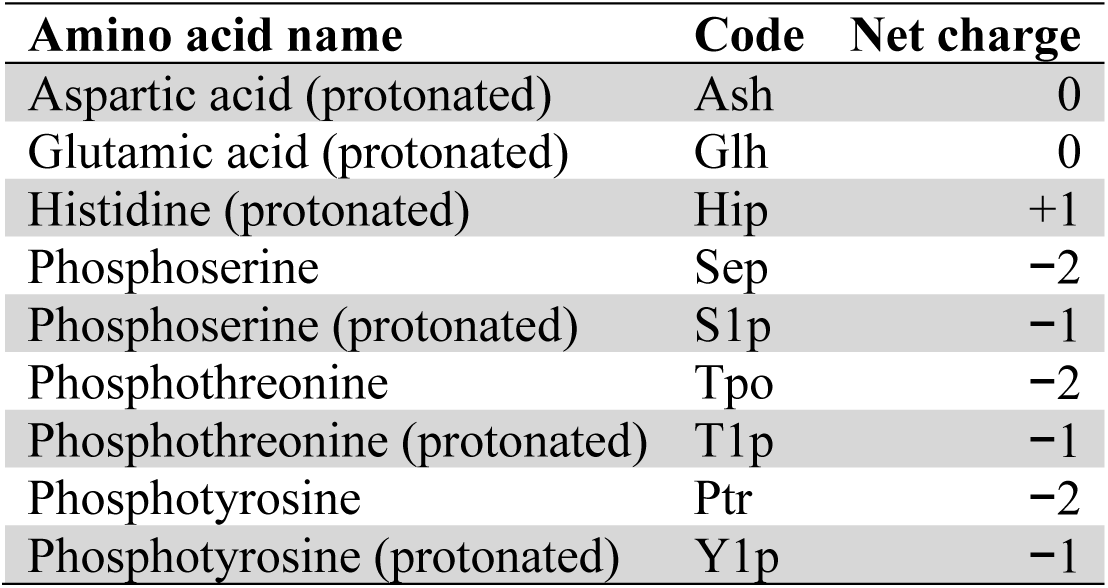
Non-standard amino acids that were investigated in this study. The codes used for the amino acids correspond to their ff99SB-ILDN residue codes.

| Amino acid name | Code | Net charge |
| --- | --- | --- |
| Aspartic acid (protonated) | Ash | 0 |
| Glutamic acid (protonated) | Glh | 0 |
| Histidine (protonated) | Hip | +1 |
| Phosphoserine | Sep | -2 |
| Phosphoserine (protonated) | S1p | -1 |
| Phosphothreonine | Tpo | -2 |
| Phosphothreonine (protonated) | T1p | -1 |
| Phosphotyrosine | Ptr | -2 |
| Phosphotyrosine (protonated) | Y1p | -1 |

### Threonine phosphorylation decelerates fast interchange of conformational states

To ensure that the MD results are statistically viable and reproducible we compared how well the conformational sampling of the different replica simulations correlated with one another. Under the assumption that all replica simulations sample from the same conformational free energy landscape, over the course of the sampling all replicas should converge to increasingly accurate representations of that same free energy landscape. The correlation coefficient of populations of individual conformational states between the replicas (*r*_replica_) provides an intuitive and unbiased measure of this convergence.

To compare the conformational ensembles of individual replica simulations, we adopted the method by Campos and Baptista,^45^ where conformational ensembles are represented as energy landscapes in a multidimensional space based on backbone coordinates. By employing principal component analysis (PCA), the number of dimensions can be further reduced allowing for the efficient comparison of conformational ensembles in the same transformed space. We reduced the space to the first five principal components that still cover more than 35% of the conformational variability. While this limits the accuracy of the representation of the conformational space, it allows for conformationally related structures to be clustered together and facilitates the comparison between the replicas. Unlike the original publication and given the conformational variability of these small peptides where there is no overall context from a protein fold, we performed the PCA on the backbone dihedrals (*φ*, *ψ*) directly, eliminating the need for a centroid structure that is expensive to calculate for large ensembles.

Initial analyses showed that most simulations converged reasonably well after 300 ns of sampling per replica (1.2 µs of cumulative sampling). Therefore, we simulated all systems for 300 ns by default. Subsequently, we extended simulations with poor correlation between replicas (minimum pairwise *r*_replica_ < 0.3) progressively up to 900 ns pre replica (3.6 µs of cumulative sampling). In the end, more than 81% of the simulations show high *r*_replica_ of > 0.7 (**Figure 1**), with all *r*_replica_ values provided as a measure of confidence for any results derived from these simulations by us or third parties (**Suppl. Table 2**). Only a small fraction of the simulations did not converge. The latter is a strong indication that in these simulations either (meta-)stable conformations are adopted, or high energy barriers persist, both of which restrict backbone dynamics and thus the conformational sampling. Notably, a high number of the simulations with poor convergence feature phosphothreonine side chains (Tpo, **Suppl. Fig. 1**). This indicates that Tpo restricts backbone dynamics, so potentially stabilizing otherwise disordered regions.

**Figure 1.**
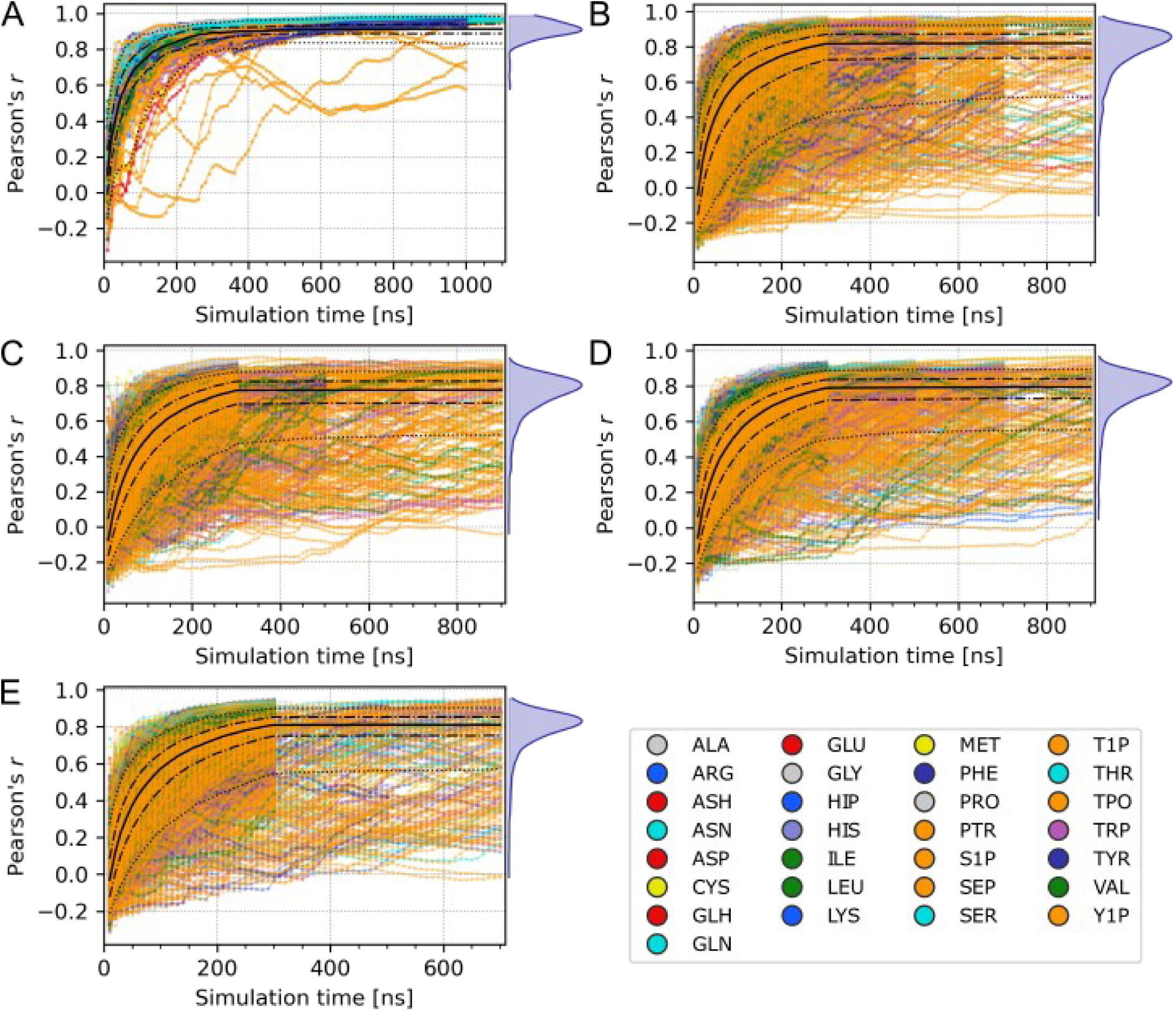
Correlation coefficients between replicas (r_replica_) as a function over the simulation time. The color of the lines indicate the amino acid type (X_1_) investigated in the corresponding simulation. The dotted, dashed, and solid black lines represent the 5%, 25%, 50%, 75%, and 95% percentiles. The total distribution of the correlation coefficients over all simulations is visualized as a blue density curve on the right. **A.** Simulations of pentapeptides (GGX_1_GG). **B.** Simulations of hexapeptides (GGX_1_X_2_GG). **C.** Simulations of heptapeptides (GGX_1_GX_2_GG). **D.** Simulations of octapeptides (GGX_1_GGX_2_GG). **E.** Simulations of nonapeptides (GGX_1_GGGX_2_GG).

### Individual amino acids show only minor differences in their backbone dihedral energy landscapes

To investigate the conformational preferences of individual amino acids, we generated Ramachandran plots for the central residue in the GGXGG peptide series, plotting all (*φ*, *ψ*) along the simulations. With exception to glycine and proline, which have unique Ramachandran profiles, the profiles of all remaining residues are, as expected, characterized by five energy minima (**Figure 2H**): α_R_-region, α’_R_-region, β-region, ppII-region, and α_L_-region. Differences in the relative populations of these minima show the individual conformational preferences of the corresponding residues. These differences are, as expected, not very pronounced since the glycine backbone does not stabilize any emerging secondary structure. Still, the interactions from the single amino acid side chain itself already generates differences in overall peptide behavior.

**Figure 2.**
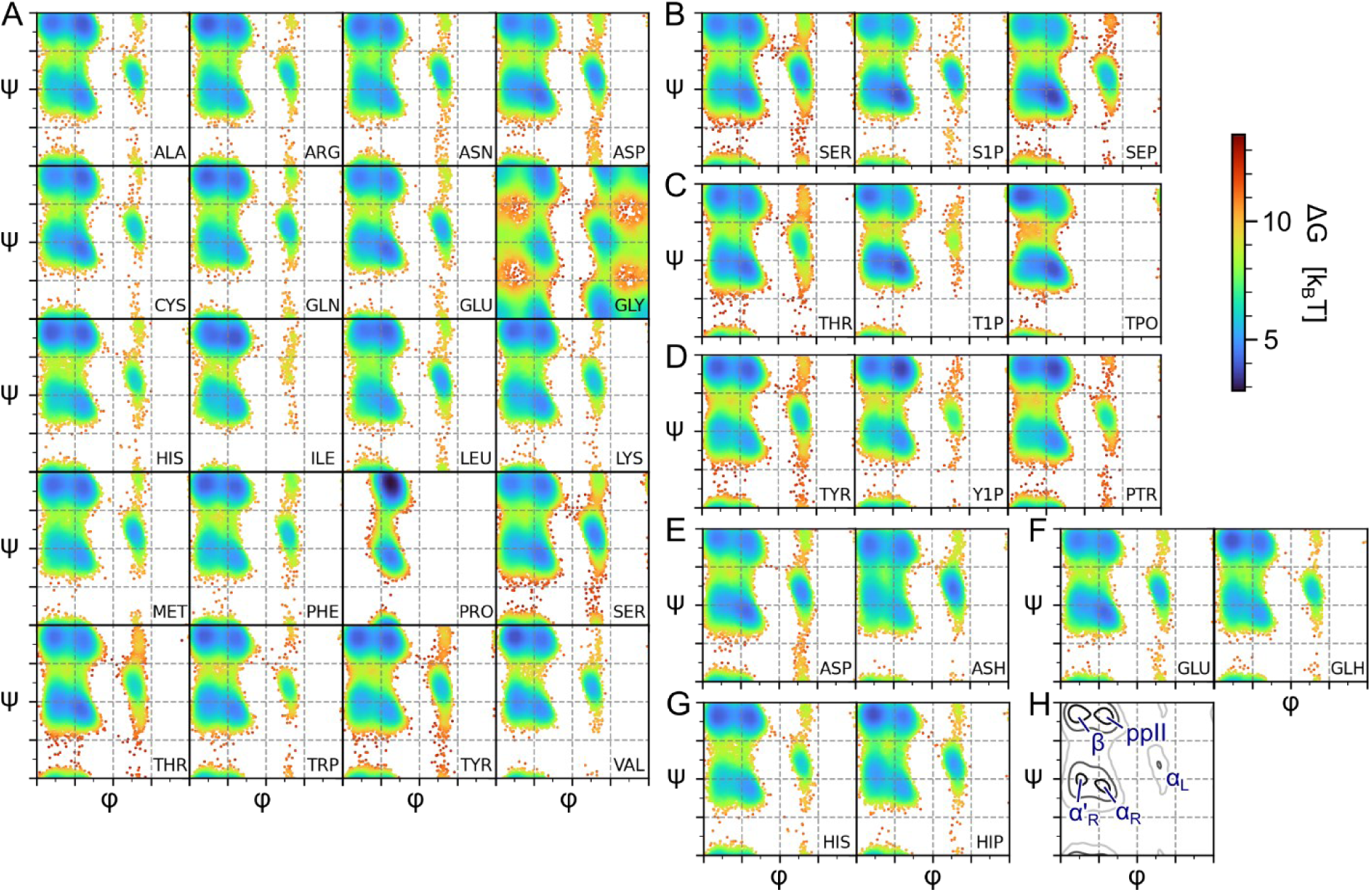
Ramachandran plots obtained from the sampling of GGXGG peptides. The backbone dihedrals (φ, ψ) of the central residue (shown in the lower right of each plot) are plotted in the range [-180°, 180]. From the sampling populations relative free energies for all areas of the plot were inferred indicated by the coloring of the data points. **A.** Standard amino acids in alphabetical order. **B.** Serine and its phosphorylated forms (S1P, SEP). **C.** Threonine and its phosphorylated forms (T1P, TPO). **D.** Tyrosine and its phosphorylated forms (Y1P, PTR). **E.** Aspartate and its neutral form aspartic acid (ASH). **F.** Glutamate and its neutral form glutamic acid (GLH). **G.** Histidine and its protonated form (HIP). **H.** Schematic overview of the energy minima exhibited in most of the Ramachandran plots.

One general pattern that emerges is that negative charges on the amino acid side chain lead to an increase in the population of the α_R_-region. This can be observed when comparing Asp to Ash/Asn, or Glu to Glh/Gln. When looking at the phosphorylation of Ser (S1p, Sep), the same increase in the α_R_-region is observed, accompanied by a slight decline in the β- and ppII-region. In contrast, positive charges show a slight tendency to promote the β-region. This can be observed in Arg and the protonation of His (Hip). In Lys, however, the effect is barely visible. For Thr (T1p, Tpo), the strongest changes upon phosphorylation are observed. There is again an increase in the α_R_-region as well as in the β-region, while the ppII-region decreases. Interestingly, the connection between the α_R_-region and β-region is also less populated than in other amino acids, showing that Tpo would stabilize either of these conformations with reduced exchange between them, which is likely related to the lack of convergence for the Tpo simulations. The population of the α_L_-region decreases too upon phosphorylation, to the point that in the di-anionic form (Tpo) this state was not sampled at all. Finally, for the Tyr-phosphorylation (Y1p, Ptr) only slight increases in the α_R_- region and the ppII-region are observed. The effects are less pronounced though than in the other phosphorylations, probably due to the distal position of the phosphoryl group on an already bulky side chain, so limiting its additional effect on the peptides’ backbone conformation and dynamics.

### Phosphorylations of Thr and Ser promote ordered conformations

Following the investigation into the influence of individual amino acid types on peptide conformation within GGXGG peptides, we further studied how the amino acid type changes conformational propensities in a more complex local sequence context that attempts to capture interactions between amino acid side chains; are such conformational tendencies consistent across different peptide sequences? To address this, we analyzed our conformational ensembles using the *Constava* software package.^41^ This software employs a probabilistic model to extract propensities for distinct conformational states that capture typical combinations of conformation and dynamics as observed by NMR in solution. The conformational states are *core helix* (exclusively α-helical with minimal backbone dynamics), *surrounding helix* (mostly α-helical with significant backbone dynamics), c*ore sheet* (exclusively β-sheet with minimal backbone dynamics), *surrounding sheet* (β-sheet and ppII with significant backbone dynamics), *turn* (mostly turn conformation and highly dynamic), and *other* (mostly coil conformation and highly dynamic).

Our analysis revealed that *core helix* propensities that are very low in the isolated amino acid generally remained quite low in the more complex peptides (*e.g.*, Val), with Pro, Sep, and Tpo showing the most substantial propensities for this conformational state (**Figure 3A**). Conversely, *surrounding helix* displayed a similar amino acid type pattern, yet exhibited higher overall propensities, reflecting the dynamic nature of the short peptides in our study (**Figure 3B**). Phosphorylation led to increased helix propensities for both Ser and Tyr, with this effect being more pronounced for Ser. Although Thr exhibited a trend of elevated core helix propensities following phosphorylation, surrounding helix propensities displayed bidirectional trends. Generally, negatively charged residues seem to promote *core*/*surrounding helix* propensities. This is observed in the phosphorylated residues as well as for the protonation states of Glu and Asp. In terms of *core sheet* and *surrounding sheet* propensities, dissimilar patterns emerged (**Figure 3D, 3E**). Here, phosphorylation of Ser phosphorylation led to decreased propensities in both *core sheet* and *surrounding sheet* propensities. In contrast, Tyr phosphorylation resulted in reduced *core sheet* propensity but increased *surrounding sheet* propensity. Thr phosphorylation increased *core sheet* propensities, while *surrounding sheet* propensities displayed a divergent trend similar to the helical conformational states. The *core*/*surrounding sheet* propensities for Glu and Asp increase when protonated. Finally, the *turn* and *other* conformational states are linked to high backbone dynamics. Upon phosphorylation, both Ser and Thr show decreases in *other* propensity when phosphorylated (**Figure 3F**), while Thr also showed decreased *turn* propensities (**Figure 3C**). Phosphorylation had no effect on *turn* propensities for Ser or Tyr. Mono-anionic Y1p exhibited a slight increase in other propensities, while Ptr showed no changes.

**Figure 3.**
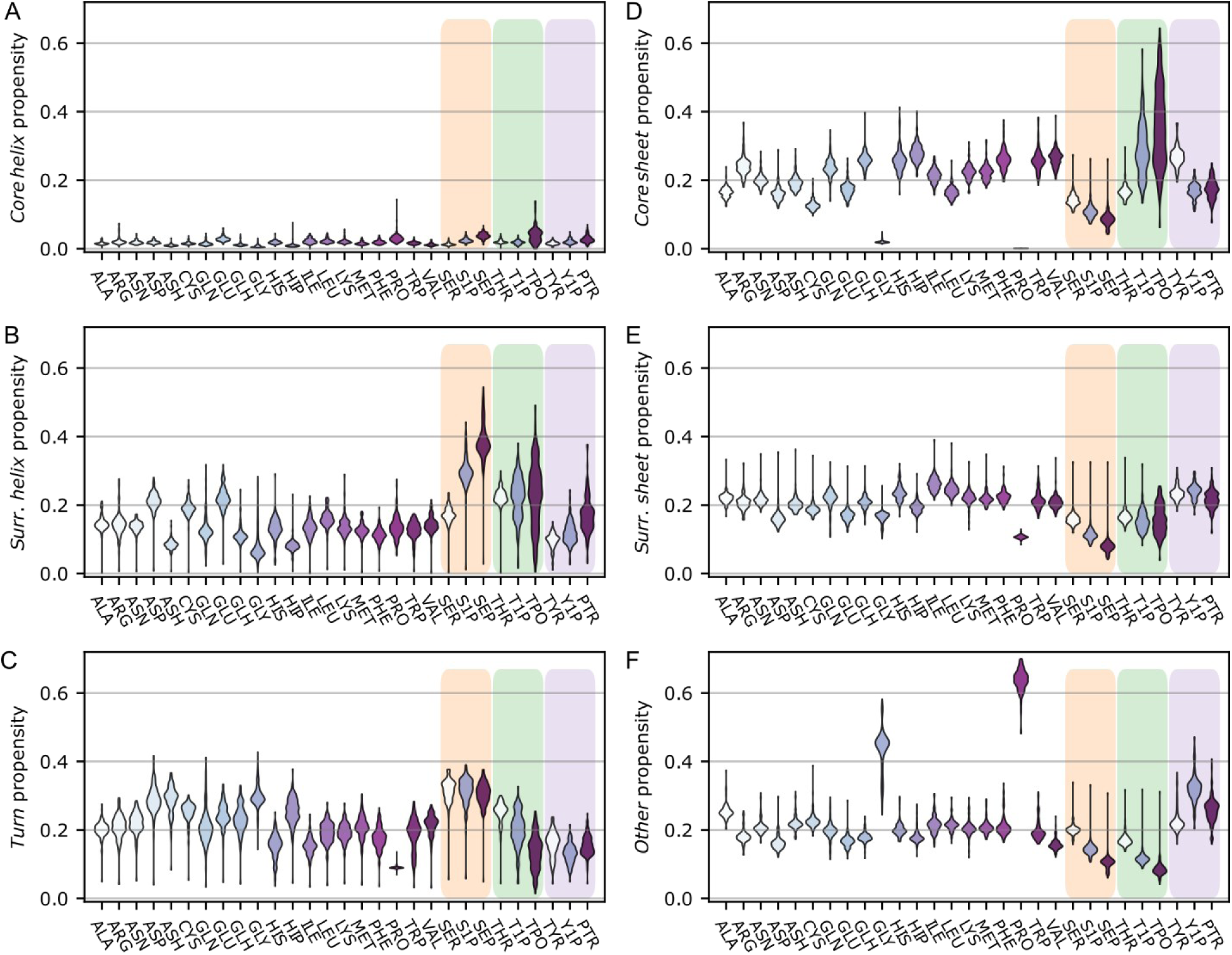
Conformational state propensities of amino acid types. The conformational state propensities of individual residues are shown as violin plots over all simulated peptides. For glycine only residues in the positions X_1_ and X_2_ are analyzed, ignoring the “scaffold glycines”. Serine, threonine, and tyrosine (and their phosphorylated forms) are highlighted in orange, green, and purple, respectively. **A.** Core helix propensities per residue type. **B.** Core sheet propensities per residue type. **C.** Surrounding helix propensities per residue type. **D.** Surrounding sheet propensities per residue type. **E.** Turn propensities per residue type. **F.** Other propensities per residue type.

Intriguingly, Tpo consistently exhibited greater energy barriers between conformational states compared to other residues. So, it might serve as a conformational lock, locking the backbone into specific conformations that depend on the local sequence context. The poor convergence of the Tpo simulations severely limits such an interpretation. The observed decreases in the *other* and turn propensities indicate that both Sep and Tpo exhibit higher propensities for structured backbones compared to their non-phosphorylated counterparts.

### Residues restrict the conformational space of N- and C-terminal neighbors

We further wanted to investigate how residues affect neighboring residues in C-terminal and N-terminal direction. To that end, we compared the Ramachandran plots for all residues in peptides of type GGXGGGGGG and GGGGGGXGG, with corresponding residues in the all-glycine nonapeptide (X = Gly). In this way, any effect the residue X has on its neighboring residues in N-or C-terminal direction would become apparent as a change of the dihedral profile in comparison to the reference peptide.

The results show that the effects of non-glycine residues in position *i* are most apparent in the adjacent position *i* ± 1 and can stretch up to position *i* ± 2 (**Figure 4**). Beyond that position, any residual effects cannot be discerned from the normal sampling variability. Further, residues that themselves show pronounced differences in their conformational preferences to glycine also have stronger influences on their neighboring residues. Most residues preferentially exhibit effects in C-terminal direction, with exception to Pro and Tyr that most strongly act in N-terminal direction. For Pro the N-terminal effect can be partly explained by its unique structure, that prevents backbone hydrogen bonds to the proline’s nitrogen form being formed. This disrupts any type of right-handed helix in N-terminal direction of the Pro residue. For Tyr, the interpretation of the N-terminal effect is less clear. We observe hydrogen bond formation between the phenolic OH-group and the carbonyl oxygen in position *i*-4, that could serve as a n explanation. These hydrogen bonds are, however, not particularly persistent. For phosphorylations it becomes apparent that phosphorylated residues generally show stronger effects than their respective unphosphorylated counterparts and that the effects act in C-terminal direction. Interestingly, for the phosphorylation of Tyr this results in an inversion in the main direction of effect, where Tyr shows stronger effects in N-terminal direction while Ptr acts C-terminally, though the effect is not very strong. Tpo shows the strongest effect in C-terminal direction overall, followed by Sep. This further shows that phosphorylated residues tend to stabilize backbone conformations.

**Figure 4.**
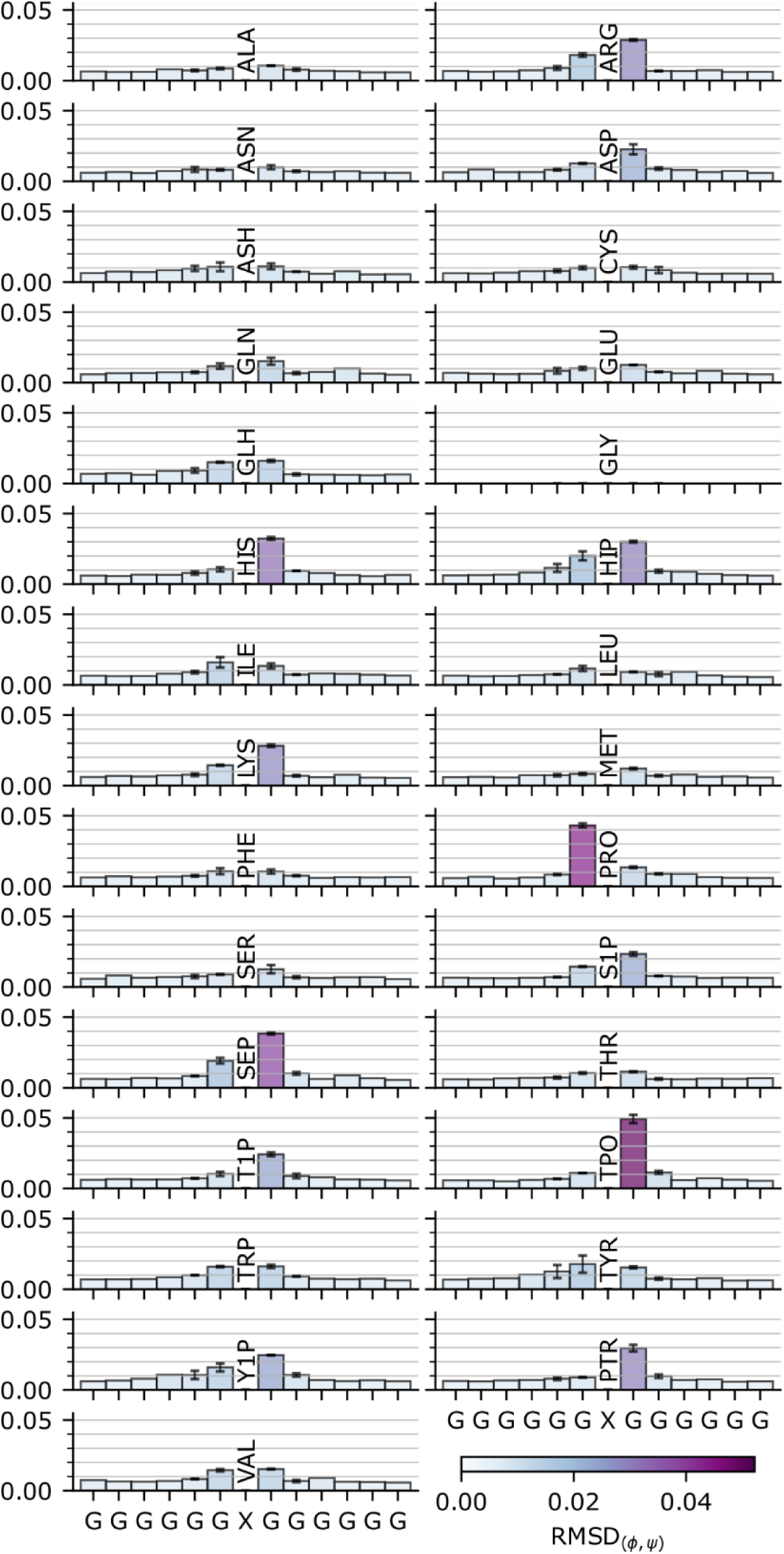
Effects of non-glycine residues on neighboring residues. The magnitude of the effect is shown as the RMSD of the respective Ramachandran plots in relation to a glycine in an all-glycine peptide. For the analysis peptides of types GGXGGGGGG and GGGGGGXGG were investigated (with X=G being the reference peptides). For the overlapping residues the errorbars indicate the upper and lower bounds of the RMSD. A representation of the individual peptides is given in **Suppl. Fig. 2**.

### Local sequence context influences conformational preferences

Since the previous results showed that residues have notable effects on the backbone dynamics, even in the absence of other nearby amino acid side chains, we analyzed if there are particular types of effect exerted by different types of amino acids. In peptides of type GGX_1_X_2_GG, we investigated how the residue X_1_ changes the conformational preferences of the residue X_2_ in comparison to glycine and vice versa (**Figure 5**). We did the same analysis for residues in positions *i*±2, *i*±3, and *i*±4 using peptides of type GGX_1_GX_2_GG, GGX_1_GGX_2_GG, and GGX_1_GGGX_2_GG. Though the average magnitude of the effects diminishes, individual residues combinations lead still to notable effects even when spaced out by three glycine residues (**Suppl. Fig. 3-5)**.

**Figure 5.**
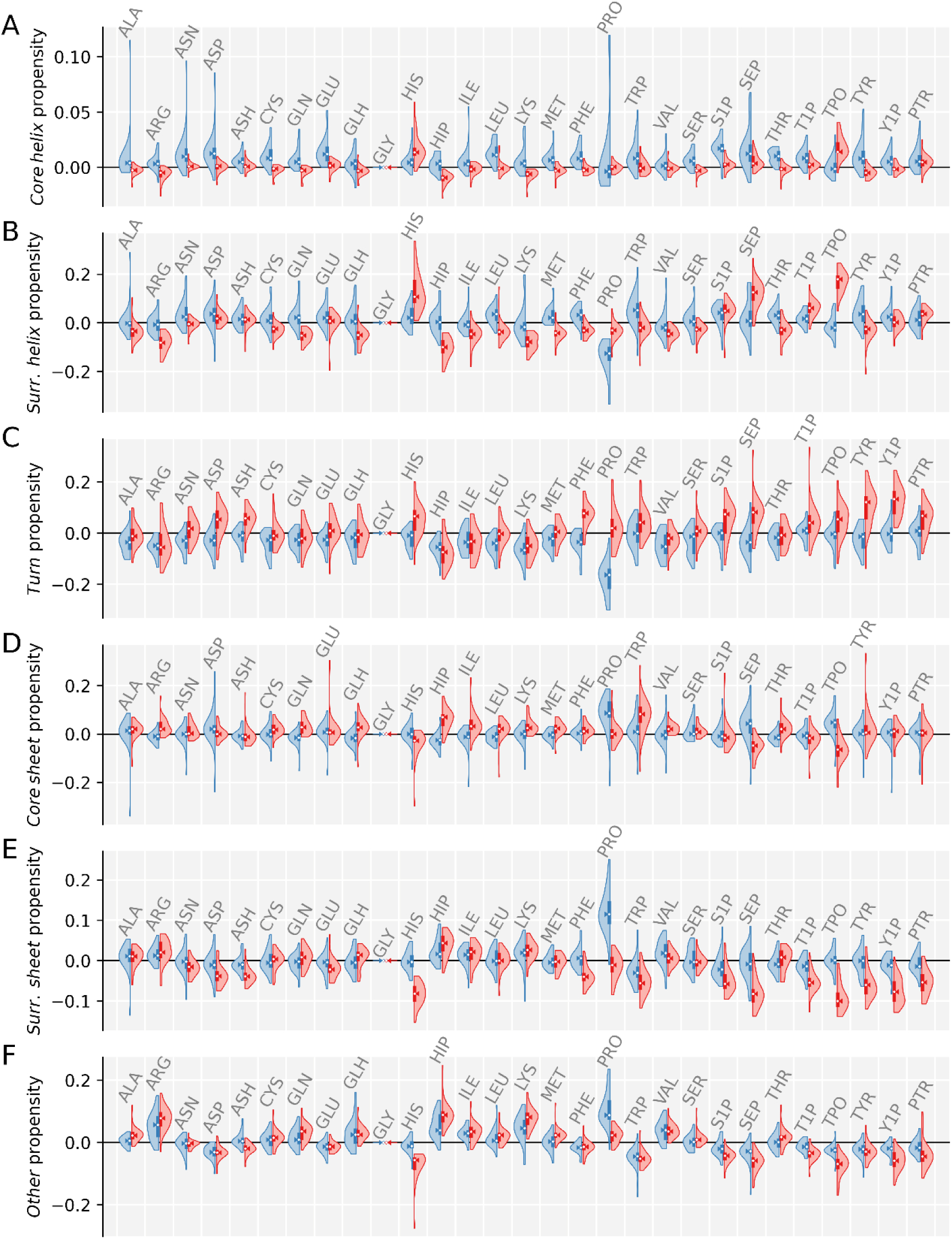
Changes of conformational propensities induced on residues in position i±1. For each residue type, two distributions are shown: the effects on residues in position i-1 (blue), and the effect on residues in position i+1 (red). All effects are shown relative to the effects of a Gly residue. **A.** Changes in core helix propensities per residue type. **B.** Changes in surrounding helix propensities per residue type. **C.** Changes in turn propensities per residue type. **D.** Changes in core sheet propensities per residue type. **E.** Changes in surrounding sheet propensities per residue type. **F.** Changes in other propensities per residue type.

Concordant to the individual residue propensities, the analysis of the *core helix* conformational state only shows small effects. His, Sep, and Tpo lead to increased *surrounding helix* conformational state propensities in C-terminal direction, suggesting that they promote the formation of α-helical structures. Intriguingly, no such effect is observed for unphosphorylated Ser and Thr residues. This highlights the capacity of Ser and Thr phosphorylation to function as switches capable of inducing secondary structure.^46–48^ In the context of the *core sheet* conformational state, both Ser and Thr phosphorylation lead to marginal reductions in propensities, albeit with minor impact. The *surrounding sheet* conformational state encompasses next to β-sheet like extended structures also the polyproline II state. Pro residues show the strongest effect here, strongly promoting this conformational state in N-terminal direction, whilst strongly reducing the *surrounding helix* propensity especially in that direction. In contrast, phosphorylated residues generally lead to decreased *surrounding sheet* propensities in C-terminal direction. In the same line *other* propensities which correspond to random coil motions are generally disfavored after phosphorylated residues. In contrast, both Sep and Tpo show increased propensities for turn-like conformations. Many more subtle effects are present in the data, for example Phe increases *surrounding helix* propensity towards the N-terminus but decreases it towards the C-terminus.

A general pattern for the phosphorylated residues is that they tend to promote bent and more compact conformations in our simulation. They disfavor extended conformational states like the *surrounding sheet* or *other* conformational state.

### Changes in the hydrogen bond network upon phosphorylation

The phosphorylation of the amino acid side chains also introduces strong hydrogen bond acceptors (and a donor in S1p, T1p, and Y1p). This allows for new hydrogen bonds to be formed between the phosphate group and adjacent residues, contributing to the changes in the conformational preferences described above. To analyze these effects, we visualized the relative occurrence of hydrogen bonds across our peptide series.

For Ser peptides we find the hydrogen bond N*_i_*_+2_ →O*_i_*_-1_ to be the most prevalent hydrogen bond (**Figure 6A**). This hydrogen bond corresponds to a single turn in a 3_10_ helix showing a tendency to form transient helical structures. Upon phosphorylation, these 3_10_ helix hydrogen bonds become more frequent indicating that the phosphorylation stabilizes helical conformations (**Figure 6B**). Furthermore, we see more interactions between the phosphoryl group and other side chains than in Ser. Finally, the phosphoryl group occasionally forms hydrogen bonds with the amide nitrogen of the Sep, that way rigidifying the backbone.

**Figure 6.**
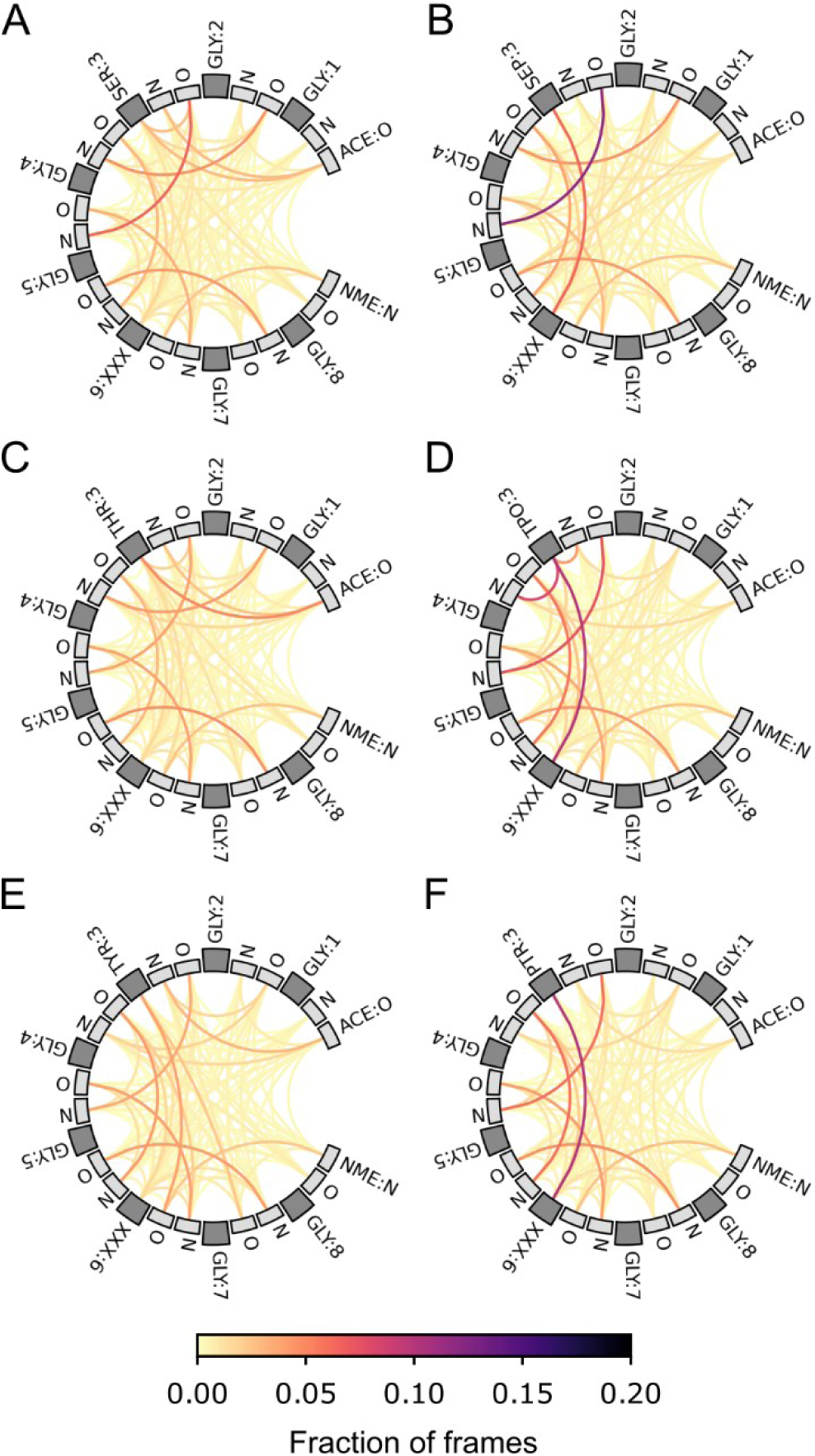
Changes in the hydrogen bond network in octapeptides upon phosphorylation. The peptides are drawn depicted on a circle from the N-terminal acetyl cap (ACE) to the C-terminal N-methyl cap (NME). Dark boxes indicate the side chain of the corresponding residue, while the grey boxes left and right of the side chains are the backbone N and O. Hydrogen bonds are drawn as lines between a donor and an acceptor. The color of the line indicates the fraction of the MD simulations in which the particular hydrogen bond was observed. The hydrogen bond networks are shown for **A.** serine peptides (Ser), **B.** phophoserine peptides (Sep), **C.** threonine peptides (Thr), **D.** phosphothreonine peptides (Tpo), **E.** tyrosine peptides (Tyr), and **F.** phosphothreonine peptides (Ptr). The respective residue is always in position three while position six (XXX) is any residue.

A similar pattern is observed for Thr peptides, where the phosphorylation increases 3_10_ helix hydrogen bonds, and leads to more sidechain-sidechain hydrogen bonds (**Figure 6C, 6D**). For Tpo, however, hydrogen bonds to the backbone are more prevalent than for Sep. Tpo forms strong hydrogen bonds between the phosphoryl group and N*_i_* / N*_i_*_+1_. These hydrogen bonds are likely the reason for the strong rigidification of the backbone, which makes the Tpo simulations particularly slow to converge.

Finally, for Tyr peptides changes in the hydrogen network upon phosphorylation are less prominent than in the other two peptide series (**Figure 6E, 6F**). Ptr is unable to for those hydrogen bonds to directly adjacent backbone nitrogens. So, its phosphoryl group primarily forms hydrogen bonds to hydrogen bond acceptors on other side chains.

Generally, hydrogen bonds are more prevalent in the phosphorylated peptides than they are in their unphosphorylated counter parts. While Sep and Tpo for hydrogen bonds with the local backbone, for Ptr the most notable hydrogen bonds are formed to other amino acid side chains. The hydrogen bonds of Sep and Tpo with the local backbone in particular, likely contribute to their more restricted backbone dynamics.

### Comparison of MD ensemble-derived properties to NMR chemical shift-derived descriptors

Given the computational provenance of the dataset, a validation of the observations against experimental data is vital to assess its accuracy. NMR chemical shifts are a good source for such validation, as they are readily available from the BMRB^19^ and contain information on peptides’ and proteins’ backbone conformation and dynamics. Even though few chemicals shifts are available for phosphorylated residues, several samples for each type of phosphorylation could be obtained from the BMRB. Comparing such chemical shift information, which encompasses the effect of phosphorylation on a specific system with a sequence containing diverse amino acids, to our conformational ensembles is non-trivial. Chemical shifts capture the time-averaged backbone conformation in solution, while in the MD simulation individual conformations are resolved at atomistic detail. Thus, we compared the higher-level conformational state propensities from the MD conformational ensembles with corresponding secondary structure propensities derived from the chemical shifts.

To that end, we searched the BMRB for matching pairs of proteins in a non-phosphorylated and phosphorylated state and analyzed the changes in the chemical shift-derived conformational descriptors. To avoid biases in the data from highly ordered proteins, where the overall fold is very stable and unlikely to be affected by local changes, we focused on highly dynamic proteins by excluding proteins with random coil index-derived S^2^ parameters (S^2^_RCI_) > 0.8. We further excluded experiments that were performed under denaturing conditions (*i.e*, high concentration of urea or methanol). These chemical shift-derived conformational descriptors were then compared to properties of the conformational ensembles. S^2^_RCI_ as a measure of backbone rigidity is compared to the circular variance (variance of the backbone dihedrals), which should be inversely correlated. The chemical shift-derived secondary structure populations are compared to the conformational state propensities obtained from Constava.^41^ Note that these measures inherently follow different scales and thus the following comparison is limited to the relative trends of the changes rather than comparing absolute values.

While in the experimental measurements more fluctuations are observed than in the MD dataset, the observations from the simulations overall match the observations from NMR experiments. For both Ser and Thr, phosphorylation leads to an increase of the S^2^_RCI_ which shows a reduction of backbone dynamics (**Figure 7A, 7B**). This is matched in the MD simulations by reduction of the circular variance. For Tyr phosphorylation, though, only minor effects are observed, with both the MD data and the NMR data fluctuating around zero (**Figure 7C**).

**Figure 7.**
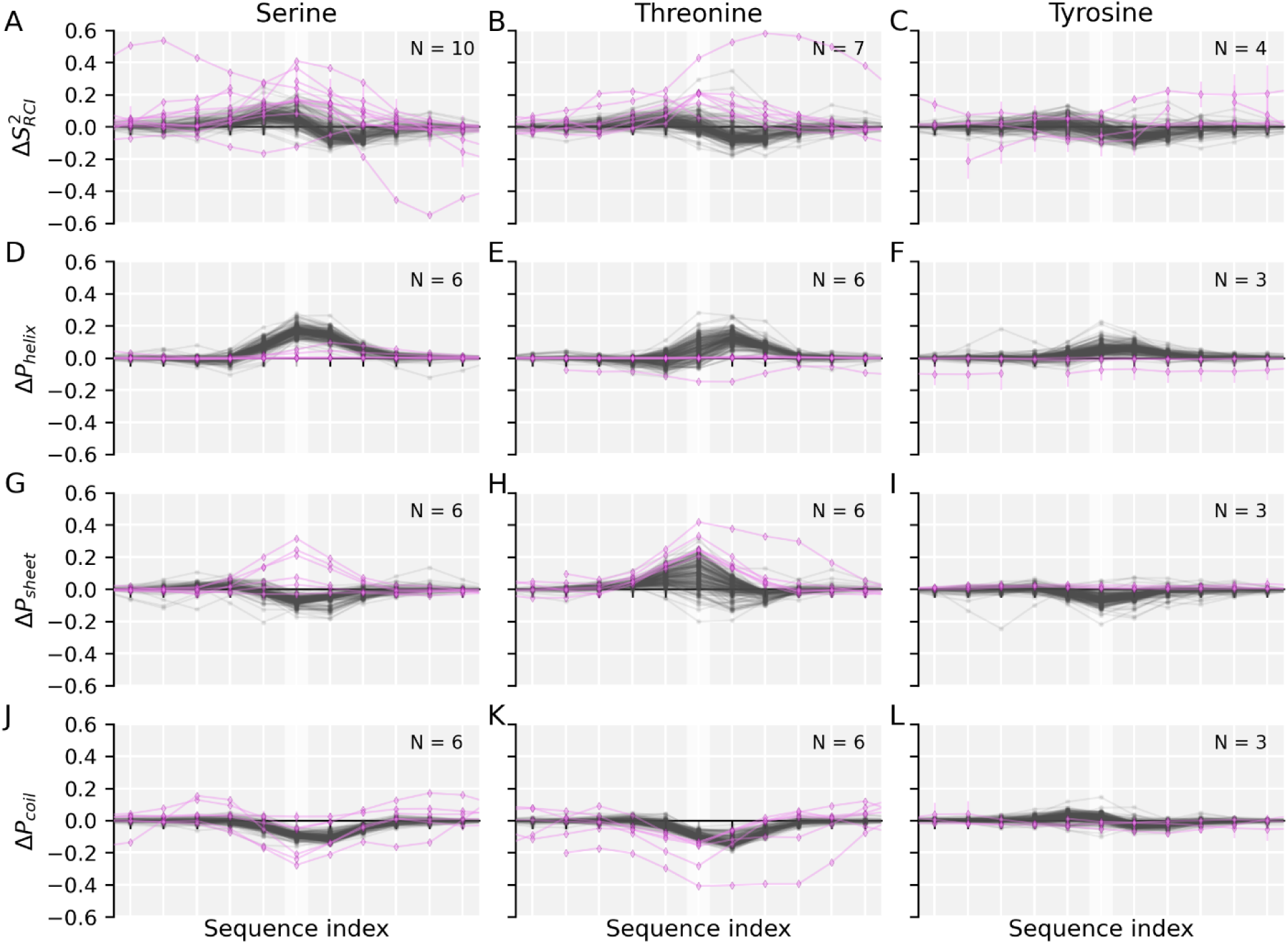
Trends of changes in backbone conformations upon phosphorylation compared between MD results and chemical shift-derived descriptors. The conformational changes induced by phosphorylations are compared between the MD simulations (grey) and NMR data from the BMRB (purple). The phosphorylation site is highlighted by a light ribbon. The changes are given for Ser, Thr, and Tyr phosphorylations in the left (**A**,**D**,**G**,**J**), center (**B**,**E**,**H**,**K**), and right (**C**, **F**, **I**, **L**) columns respectively. **A-C.** Change in S^2^ is compared to the negative change in circular variance of the backbone dihedrals (phi, psi) in the simulation. **D-F.** Change in helix propensities upon phosphorylation. **G-I.** Change in sheet propensities upon phosphorylation. **J-L.** Change in coil propensities upon phosphorylation is compared.

For Ser phosphorylation, the MD simulations show an increase of helix propensity around the phosphorylation site (**Figure 7D**). Such an increase is also observed in the NMR data, though the maximum helix propensity is reached at position *i*+1 rather than directly at the phosphorylation site. For Tpo and Ptr no increase in helix propensity is observed in the available NMR data, though in the MD data slight increases in helix propensity around the phosphorylation site are observed (**Figure 7E, 7F**). When comparing sheet propensities between the simulations and the chemical shift data, we found that upon Ser phosphorylation half the chemical shift sets indicated an increase in sheet propensity while the others only showed very subtle effects (**Figure 7G**). In contrast, the MD simulations show a slight decrease in sheet propensity, which was only observed in one of the NMR experiments. A reason here could be the missing sequence context and tertiary structure in our simulations that stabilizes sheet conformations in these particular proteins. However, it can also not be entirely excluded, that this is a bias of the force field. For Thr phosphorylation both MD simulations and NMR data indicate a strong increase in sheet propensity (**Figure 7H**). Again, for Ptr only small changes in the sheet propensity are observed (**Figure 7I**). Finally, coil propensities obtained from chemical shifts were compared against *other* conformational state propensities. For both Tpo and Sep coil propensities decrease around the phosphorylation site both in the simulations as well as in the NMR data (**Figure 7J, 7K**). This is again less prominent in Ptr, where only small fluctuations around zero occur (**Figure 7L**).

In summary, the MD simulations seem to capture the structural changes in the phosphorylated residues reasonably well. There are however notable limitations to this comparison. The simulated peptides are rich in glycine and thus more dynamic than any of the peptides from the BMRB, where ample sequence context and side-chain interactions will be present. These will promote secondary structure or long-range interactions that can bias the data towards certain conformations that cannot be observed in the short peptides we simulated. Given the lack of available experimental data and few case studies to compare the MD simulation results to, our comparison is therefore qualitative and lacks statistical power. Furthermore, especially for the Tpo simulations, poor convergence limits the confidence of the obtained values. Despite these limitations, the comparison does support some key findings of our analysis, *e.g.*, that phosphorylations generally induce changes to more ordered conformations.

## Discussion

A major concern when running large-scale MD simulations is to ensure that the results are statistically viable and reproducible. This is ideally achieved by reaching ergodic sampling, where the entire accessible conformational space has been explored, and the relative occupations of individual conformations relate to their respective free energies. However, such a sampling is rarely possible for any but the simplest systems. In a rigorous attempt to assess the ergodicity of our sampling, the convergence of replica simulations of the same peptides on the same conformational energy landscape was assessed. We could show that more than 81% of the simulations converged well. Other simulations were extended to improve overall convergence, but some simulations did not converge. Intriguingly, many of these simulations feature Tpo residues. This led to the hypothesis that the conformational space of Tpo is characterized by higher energy barriers than other residues. Analysis of the dihedral profile of Tpo in a GGXGG peptide, where Tpo emerged as the conformationally most restricted residue second only to Pro, supported this hypothesis further (**Figure 2**). NMR studies on phosphothreonine peptides showed that they form strong intrapeptide hydrogen bonds to the amide nitrogens in positions *i* and *i-*1 which would explain the restricted conformational space.^49^

We further showed that side chain phosphorylation generally leads to changes in the conformational preferences of amino acids, effects that extend to neighboring residues. These observed effects were strongest in Tpo, and Sep, while for Ptr only subtle changes were observed (**Figure 2**). The same observations were made in secondary chemical shift (SCS) analyses where both Tpo and Sep induced significant changes in the C_α_ chemical shifts, while for Ptr no significant differences from Tyr were observed.^18^ Similarly, the protonation state of side chains can influence conformational preferences, with negative charges in general promoting helical conformation. This can be observed in the phosphorylated residues (e.g., Sep or Tpo), as well as for Asp and Glu. Conversely, the positive charge induced by His protonation shifts the conformational preference towards extended sheet conformations. Such protonation effects, which can be induced by changes in environmental pH, could play an important role in for example the adaptation of proteins to different subcellular environments.^50,51^

The most notable concrete effect of all phosphorylations observed in the conformational ensembles is a general promotion of ordered conformational states over disordered conformational states. This is in line with NMR observations that show both Sep as well as Tpo form strong hydrogen bonds to their own amide nitrogen.^49,52^ Their phosphoryl groups elicit their conformational effects by directly interacting with the local backbone. In Ptr such interactions with the local backbone are prevented by the long and rigid sidechain. Thus, interactions of Ptr are more focused on electorstatic interactions with sequence-wise distant residues.^53^ In fact, in our study we observed only low conformational effects of Ptr on backbone dynamics in comparison to Sep or Tpr, since the peptides in our study were chosen to study local backbone effects which are limited for Ptr. To study long-range interaction larger proteins that exhibit tertiary structure would be required. This would however limit the transferability of our results to proteins with similar folds, thus, we did not include such cases.

Phosphorylated residues are indeed frequently discussed to induce structure in disordered peptides.^54^ This is confirmed by the general effects of the phosphorylations on adjacent residues in N- and C-terminal direction. These effects were generally stronger in the C-terminal direction, with especially phosphorylation of Ser increasing *surrounding helix* propensities in C-terminal direction (**Figure 5**). This effect is stronger for the di-anionic species (Sep) than the mono-anionic species (S1p). The same tendency was found in CD spectroscopy, where Ser phosphorylation was shown to increase helix propensity, if the phosphorylation site was N-terminal to an existing helix and destabilize helices if Ser phosphorylation was C-terminal to the helix.^55^ In the simulations we found that Sep induces higher helix propensity in C-terminal direction, which is in line with those findings. We do not observe though, a decrease in N-terminal direction. Here, no pronounced effect is observed, which could be because in the glycine-based peptides the helix propensity was low to begin with.

This also shows one of the limitations of this study. To study the residues individual effects, we embedded all residues in glycine backbones. This allowed us to extract generalizable conclusions but makes direct comparison with specific peptides in specific folds of proteins less straightforward. Furthermore, we limited ourselves to the local effects of phosphorylations on the local amino acid backbone, since such interactions would be generalizable across the whole proteome. However, this excluded long-range ionic interactions from our analysis. While these interactions arguably play an important role in the effects of phosphorylations, they are highly dependent on tertiary structure and therefore highly specific to certain systems.

Overall, though, many observations from the conformational ensembles have been observed in experimental case studies, highlighting that our study reveals general effects that can be transferred to proteins even on the sequence level.

## Conclusion

In this work we employed large-scale MD simulations to generate a conformational dataset containing a total of 3,393 different peptides. The peptides were selected to study the effects of protein phosphorylations on the backbone dynamics and conformational preferences, but also encompass general interaction effects between natural amino acids, as well as changes in protonation state. Analyses of the dataset revealed general effects of phosphorylations in IDPs and IDRs on their backbone dynamics and conformational propensities purely based on the local sequence context, which could be partially validated with NMR chemical shift data. This makes the information derived from the dataset highly transferrable. The conformational ensembles generated over the course of this study are publicly available and free to use for any further applications. Moreover, we provide the *r*_replica_ for all simulations together with the conformational data as a measure of confidence for all simulations. Furthermore, the methodology can be further extended to study the conformational effects of other post-translational modifications (*e.g.*, methylations, acetylations) or non-standard amino acids, for many of which very little is known.

## Supporting information

replica correlation coefficients

## Data availability

We bundled all scripts and templates for preparing, running and analyzing our simulations in a GitHub repository: https://github.com/Bio2Byte/bio2byte-mdutils. The fork of PeptideBuilder^25^ that was used to generate the phosphorylated and capped peptides is available in a separate GitHub repository: https://github.com/Bio2Byte/bio2byte-peptidebuilder.

The conformational dataset as well as related analyses presented in this work have been deposited on Zenodo (∼85 GiB):

- Pentapeptide simulations: <u>10.5281/zenodo.10517328</u>
- Hexapeptides simulations: <u>10.5281/zenodo.10518872</u>
- Heptapeptides simulations: <u>10.5281/zenodo.10518971</u>
- Octapeptides simulations: <u>10.5281/zenodo.10518993</u>
- Nonapeptides simulations: <u>10.5281/zenodo.10519033</u>

To limit the file size, solvent molecules have been removed from the conformational ensembles prior to upload. For the unprocessed trajectories (∼7.3 TiB), please, contact the corresponding author.

## Acknowledgements

The work in this study was founded by Research Foundation Flanders (FWO) by grant G028821N to D.B. The resources and services used in this work were provided by the VSC (Flemish Supercomputer Center), funded by the Research Foundation – Flanders (FWO) and the Flemish Government.

## Author contributions

W.V. conceptualized the study. D.B. carried out the simulations and analyses. All authors contributed to discussing and interpreting the results and the writing of the manuscript.

## Notes

https://zenodo.org/doi/10.5281/zenodo.10517328

https://zenodo.org/doi/10.5281/zenodo.10518872

https://zenodo.org/doi/10.5281/zenodo.10518971

https://zenodo.org/doi/10.5281/zenodo.10518993

https://zenodo.org/doi/10.5281/zenodo.10519033

https://github.com/Bio2Byte/bio2byte-mdutils

https://github.com/Bio2Byte/bio2byte-peptidebuilder

### Competing Interest Statement

The authors have declared no competing interest.

### Summary of Updates

* Code and templates that were used throughout this study for performing and analyzing the simulations have been made publicly available * Various clarifications throughout the Materials and Methods part, including adding the equations for calculating the RMSD and S2RCI * Streamlining of the first paragraph of the Results section (with parts being moved to Materials and Methods) * Various clarifications throughout the Results section * Adding labels in Figure 7 to improve readability

