## Supplementary material for "Effects of phosphorylation on protein backbone dynamics and conformational preferences"

#### Supplementary figures

Correlation coefficients between replicate simulations ($r_{replica}$).

This work is shared under a CC BY 4.0 license.

### Supplementary figures

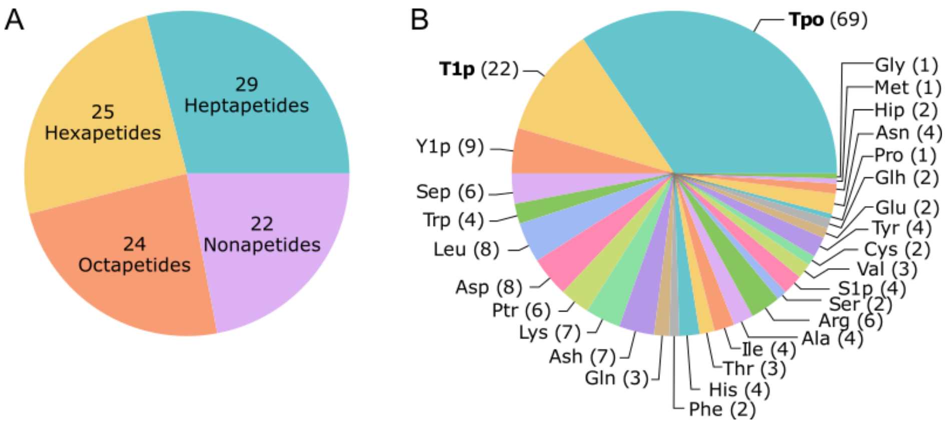

***Suppl. Fig. 1. Peptide lengths and residues of unconverged peptide simulations.*** *This figure summarizes properties of the 100 peptide simulations that showed the poorest convergence after extending the simulation time to up to 900 ns per replica ($r_{replica} \leq 0.330$).* ***A****. Frequency of the various peptide lengths observed in those simulations. Notably, the pentapeptides that only had one non-glycine residue instead of two do not show up in this figure. Therefore, we hypothesize that non-glycine sidechain interactions are a main factor that prevents simulations from converging.* ***B.*** *Residue counts in the peptides that showed the slowest convergence. Note that there are two residues $X_1$ and $X_2$ that are counted here without respect of their combination or respective position. Out of the 100 peptides 86 had at least one phosphothreonine residue (Tpo or T1p). This showcases that threonine phosphorylation exerts a strong effect on the free conformational sampling.*

This work is shared under a CC BY 4.0 license.

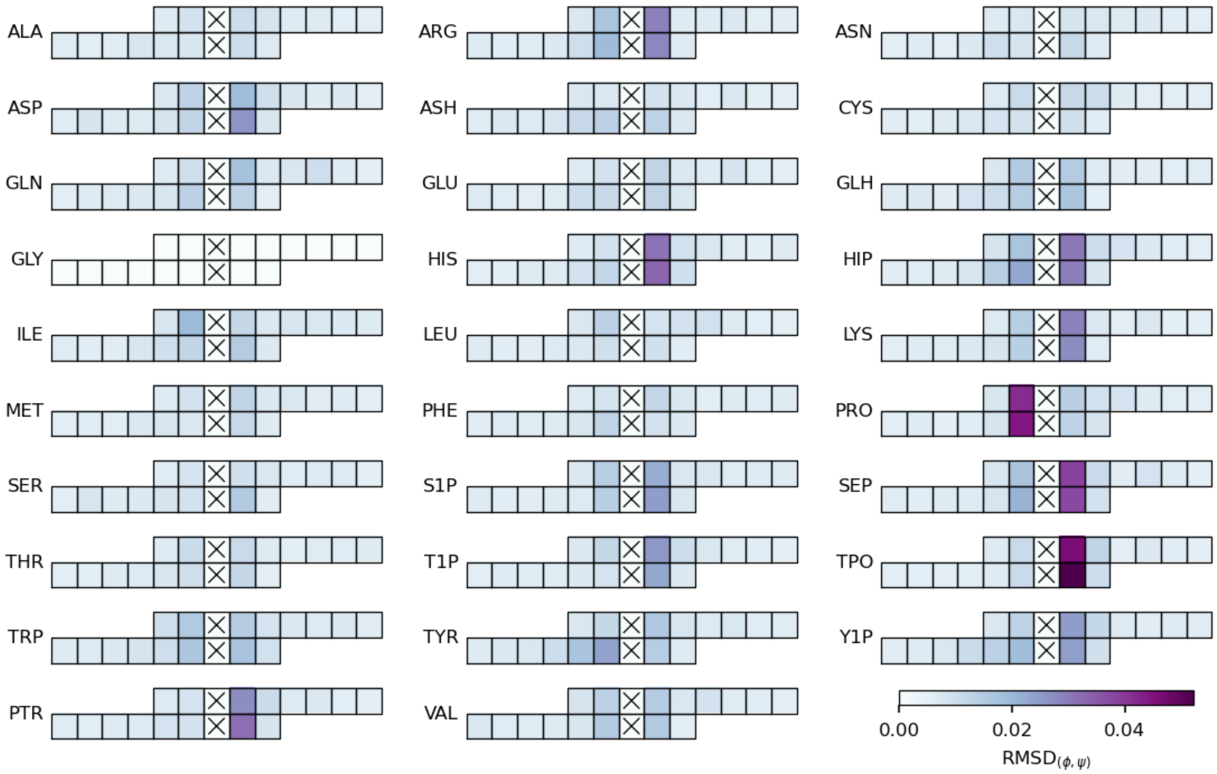

***Suppl. Fig. 2. Effects of non-glycine residues on neighboring residues.*** *The magnitude of the effect is shown as the RMSD of the respective Ramachandran plots in relation to a glycine in an all-glycine peptide. For the analysis peptides of types GGXGGGGGG and GGGGGGXGG were investigated (with X=G being the reference peptides).*

This work is shared under a CC BY 4.0 license.

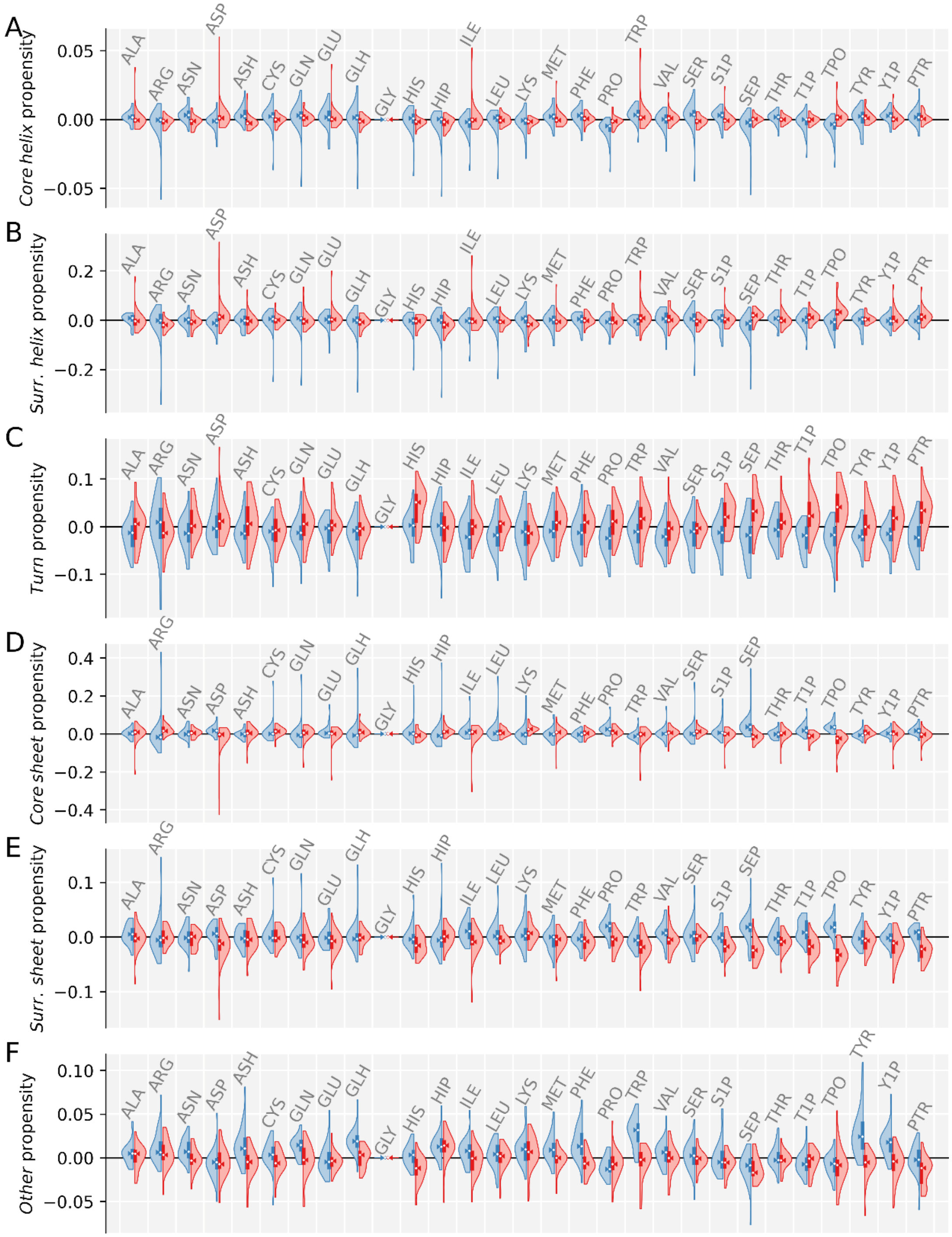

***Suppl. Fig. 3. Changes of conformational propensities on residues in positions i±2.*** *For each residue type, two distributions are shown: the effects on residues in position i-2 (blue), and the effect on residues in position i+2 (red). All effects are shown relative to the effects of a Gly residue.* ***A.*** *Changes in* core helix *propensities per residue type.* ***B.*** *Changes in* surrounding helix *propensities per residue type.* ***C.*** *Changes in* core sheet *propensities per residue type.* ***D.*** *Changes in* surrounding sheet *propensities per residue type.* ***E.*** *Changes in* turn *propensities per residue type.* ***F.*** *Changes in* other *propensities per residue type.*

This work is shared under a CC BY 4.0 license.

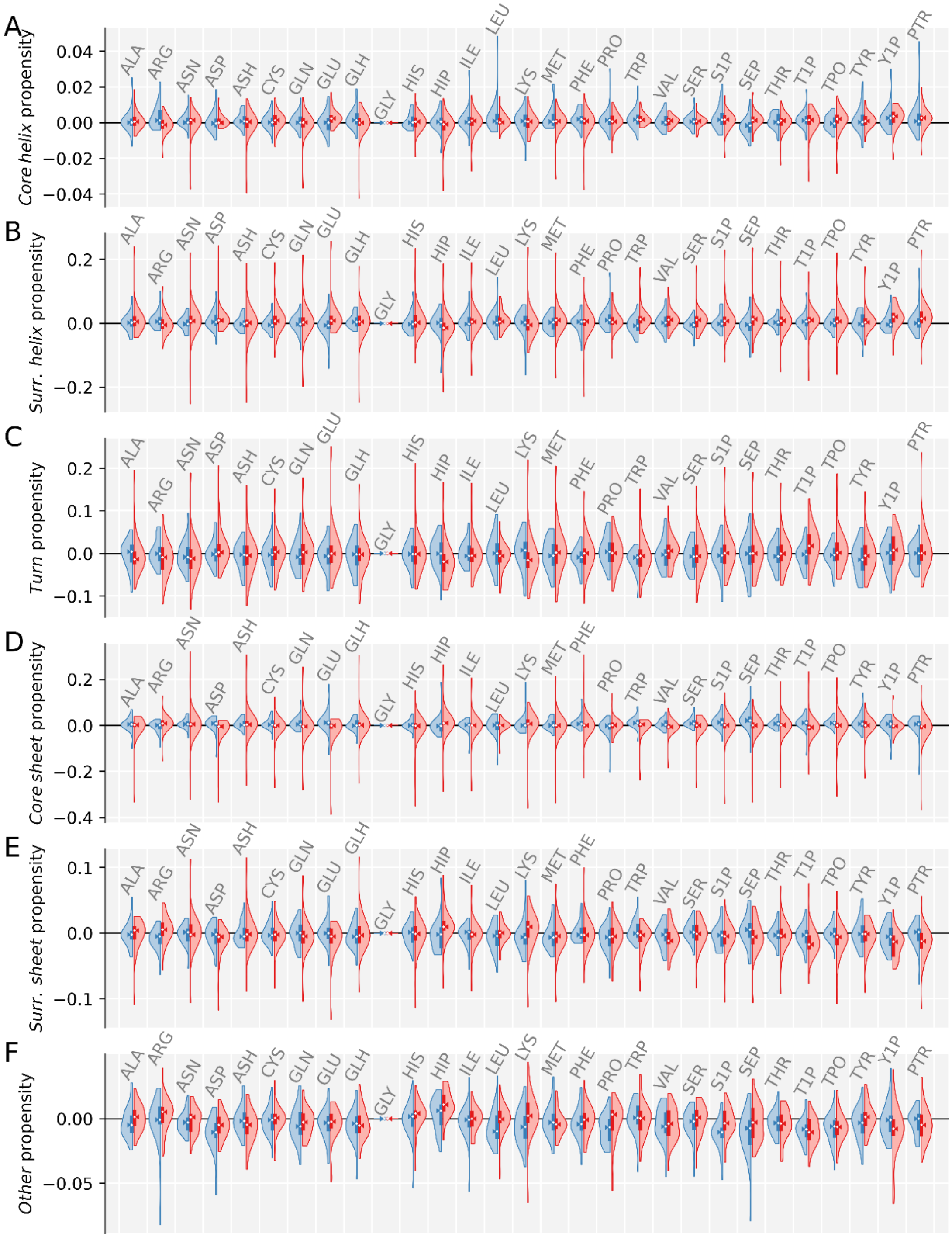

***Suppl. Fig. 4. Changes of conformational propensities on residues in positions i±3.*** *For each residue type, two distributions are shown: the effects on residues in position i-3 (blue), and the effect on residues in position i+3 (red). All effects are shown relative to the effects of a Gly residue.* ***A.*** *Changes in* core helix *propensities per residue type.* ***B.*** *Changes in* surrounding helix *propensities per residue type.* ***C.*** *Changes in* core sheet *propensities per residue type.* ***D.*** *Changes in* surrounding sheet *propensities per residue type.* ***E.*** *Changes in* turn *propensities per residue type.* ***F.*** *Changes in* other *propensities per residue type.*

This work is shared under a CC BY 4.0 license.

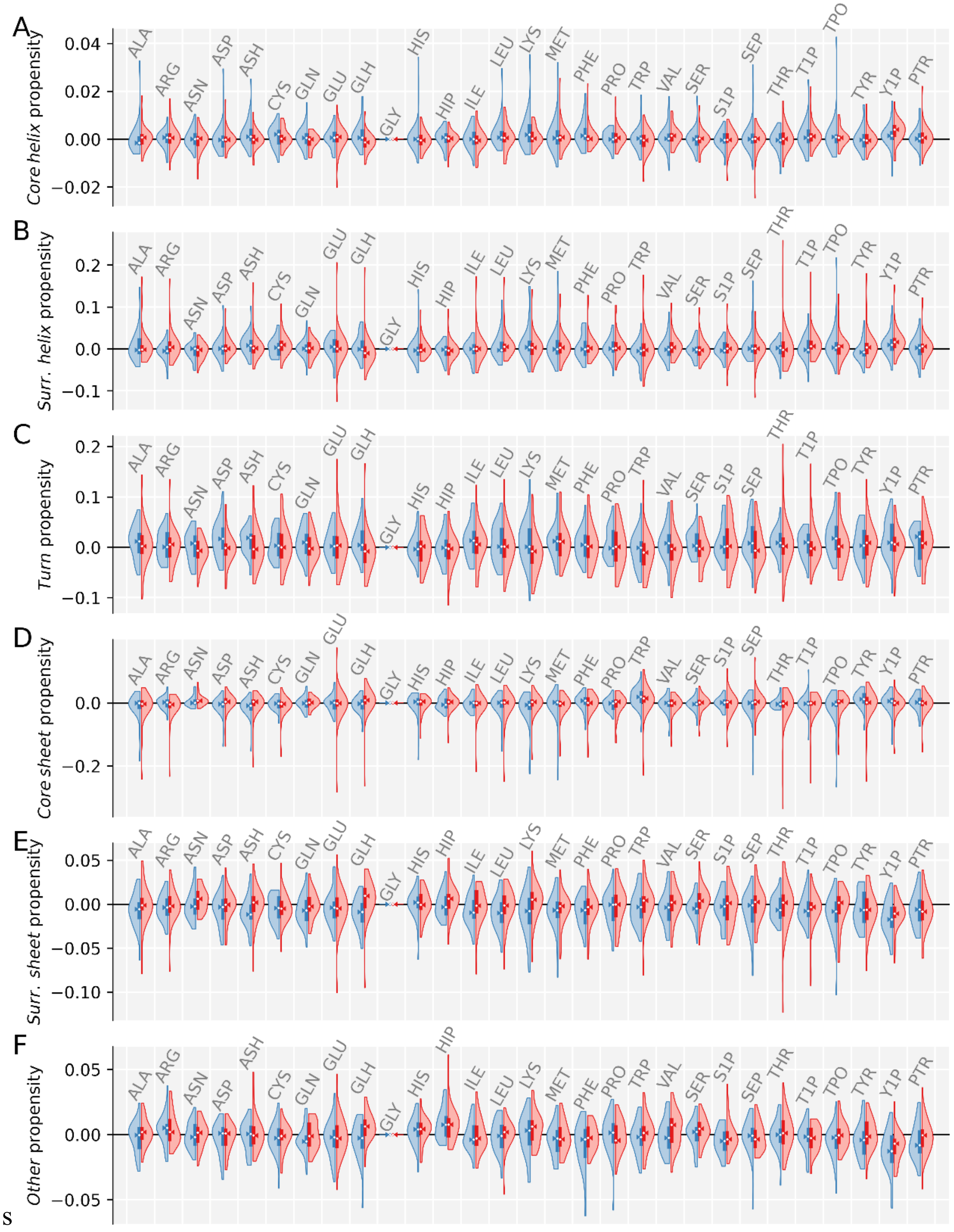

***Suppl. Fig. 5. Changes of conformational propensities on residues in positions i±4.*** *For each residue type, two distributions are shown: the effects on residues in position i-4 (blue), and the effect on residues in position i+4 (red). All effects are shown relative to the effects of a Gly residue.* ***A.*** *Changes in* core helix *propensities per residue type.* ***B.*** *Changes in* surrounding helix *propensities per residue type.* ***C.*** *Changes in* core sheet *propensities per residue type.* ***D.*** *Changes in* surrounding sheet *propensities per residue type.* ***E.*** *Changes in* turn *propensities per residue type.* ***F.*** *Changes in* other *propensities per residue type.*

This work is shared under a CC BY 4.0 license.

### Supplementary tables

***Suppl. Table 1. Starting conformations of the peptides.*** *The peptides were generated with the PeptideBuilder library from sets of backbone dihedrals. These dihedrals were chosen to minimize intramolecular hydrogen bonds.*

| | $(\varphi,\psi)_{res1}$ | $(\varphi,\psi)_{res2}$ | $(\varphi,\psi)_{res3}$ | $(\varphi,\psi)_{res4}$ | $(\varphi,\psi)_{res5}$ | $(\varphi,\psi)_{res6}$ | $(\varphi,\psi)_{res7}$ | $(\varphi,\psi)_{res8}$ | $(\varphi,\psi)_{res9}$ |
|---|---|---|---|---|---|---|---|---|---|
| ***Pentapeptides of type:*** **Gly-Gly-Xxx-Gly-Gly** | | | | | | | | | |
| **Replica 1** | −135°<br>135° | −135°<br>135° | −60°<br>−40° | −135°<br>135° | −135°<br>135° | | | | |
| **Replica 2** | −135°<br>135° | −60°<br>−40° | −135°<br>135° | −135°<br>135° | −60°<br>−40° | | | | |
| **Replica 3** | −60°<br>−40° | −60°<br>−40° | −135°<br>135° | −60°<br>−40° | −60°<br>−40° | | | | |
| **Replica 4** | −60°<br>−40° | −135°<br>135° | −60°<br>−40° | −60°<br>−40° | −135°<br>135° | | | | |
| ***Hexapeptides of type:*** **Gly-Gly-Xxx$_1$-Xxx$_2$-Gly-Gly** | | | | | | | | | |
| **Replica 1** | −135°<br>135° | −135°<br>135° | −60°<br>−40° | −135°<br>135° | −135°<br>135° | −60°<br>−40° | | | |
| **Replica 2** | −135°<br>135° | −60°<br>−40° | −135°<br>135° | −135°<br>135° | −60°<br>−40° | −135°<br>135° | | | |
| **Replica 3** | −60°<br>−40° | −60°<br>−40° | −135°<br>135° | −60°<br>−40° | −60°<br>−40° | −135°<br>135° | | | |
| **Replica 4** | −60°<br>−40° | −135°<br>135° | −60°<br>−40° | −60°<br>−40° | −135°<br>135° | −60°<br>−40° | | | |
| ***Heptapeptides of type:*** **Gly-Gly-Xxx$_1$-Gly-Xxx$_2$-Gly-Gly** | | | | | | | | | |
| **Replica 1** | −135°<br>135° | −135°<br>135° | −60°<br>−40° | −135°<br>135° | −135°<br>135° | −60°<br>−40° | −135°<br>135° | | |
| **Replica 2** | −135°<br>135° | −60°<br>−40° | −135°<br>135° | −135°<br>135° | −60°<br>−40° | −135°<br>135° | −135°<br>135° | | |
| **Replica 3** | −60°<br>−40° | −60°<br>−40° | −135°<br>135° | −60°<br>−40° | −60°<br>−40° | −135°<br>135° | −60°<br>−40° | | |
| **Replica 4** | −60°<br>−40° | −135°<br>135° | −60°<br>−40° | −60°<br>−40° | −135°<br>135° | −60°<br>−40° | −60°<br>−40° | | |
| ***Heptapeptides of type:*** **Gly-Gly-Xxx$_1$-Gly-Gly-Xxx$_2$-Gly-Gly** | | | | | | | | | |
| **Replica 1** | −135°<br>135° | −135°<br>135° | −60°<br>−40° | −135°<br>135° | −135°<br>135° | −60°<br>−40° | −135°<br>135° | −135°<br>135° | |
| **Replica 2** | −135°<br>135° | −60°<br>−40° | −135°<br>135° | −135°<br>135° | −60°<br>−40° | −135°<br>135° | −135°<br>135° | −60°<br>−40° | |
| **Replica 3** | −60°<br>−40° | −60°<br>−40° | −135°<br>135° | −60°<br>−40° | −60°<br>−40° | −135°<br>135° | −60°<br>−40° | −60°<br>−40° | |
| **Replica 4** | −60°<br>−40° | −135°<br>135° | −60°<br>−40° | −60°<br>−40° | −135°<br>135° | −60°<br>−40° | −60°<br>−40° | −135°<br>135° | |
| ***Nonapeptides of type:*** **Gly-Gly-Xxx$_1$-Gly-Gly-Gly-Xxx$_2$-Gly-Gly** | | | | | | | | | |
| **Replica 1** | −135°<br>135° | −135°<br>135° | −60°<br>−40° | −135°<br>135° | −135°<br>135° | −60°<br>−40° | −135°<br>135° | −135°<br>135° | −60°<br>−40° |
| **Replica 2** | −135°<br>135° | −60°<br>−40° | −135°<br>135° | −135°<br>135° | −60°<br>−40° | −135°<br>135° | −135°<br>135° | −60°<br>−40° | −135°<br>135° |
| **Replica 3** | −60°<br>−40° | −60°<br>−40° | −135°<br>135° | −60°<br>−40° | −60°<br>−40° | −135°<br>135° | −60°<br>−40° | −60°<br>−40° | −135°<br>135° |
| **Replica 4** | −60°<br>−40° | −135°<br>135° | −60°<br>−40° | −60°<br>−40° | −135°<br>135° | −60°<br>−40° | −60°<br>−40° | −135°<br>135° | −60°<br>−40° |

This work is shared under a CC BY 4.0 license.

***Suppl. Table 2. Correlation coefficients between replicate simulations ($r_{replica}$).*** *The pairwise correlation coefficients are given between all independent simulations of a molecular system. If simulations were extended all replicates were extended by the same amount of time. For a machine-readable version of this table consult the supplemental file:* rreplica_data.tsv.

| | **Peptide** | $r_{replica}$**(1,2)** | $r_{replica}$**(1,3)** | $r_{replica}$**(1,4)** | $r_{replica}$**(2,3)** | $r_{replica}$**(2,4)** | $r_{replica}$**(3,4)** | **Median** |
|---|---|---|---|---|---|---|---|---|
| 0 | GG**Ala**GG | 0.922 | 0.932 | 0.939 | 0.936 | 0.936 | 0.939 | 0.936 |
| 1 | GG**Arg**GG | 0.911 | 0.896 | 0.890 | 0.894 | 0.893 | 0.856 | 0.894 |
| 2 | GG**Ash**GG | 0.904 | 0.889 | 0.900 | 0.887 | 0.927 | 0.879 | 0.894 |
| 3 | GG**Asn**GG | 0.928 | 0.924 | 0.921 | 0.950 | 0.917 | 0.914 | 0.923 |
| 4 | GG**Asp**GG | 0.903 | 0.905 | 0.887 | 0.898 | 0.874 | 0.857 | 0.893 |
| 5 | GG**Cys**GG | 0.916 | 0.934 | 0.923 | 0.908 | 0.909 | 0.936 | 0.920 |
| 6 | GG**Glh**GG | 0.899 | 0.923 | 0.920 | 0.928 | 0.884 | 0.913 | 0.917 |
| 7 | GG**Gln**GG | 0.919 | 0.940 | 0.925 | 0.931 | 0.942 | 0.944 | 0.935 |
| 8 | GG**Glu**GG | 0.855 | 0.920 | 0.862 | 0.831 | 0.871 | 0.875 | 0.866 |
| 9 | GG**Gly**GG | 0.985 | 0.982 | 0.990 | 0.983 | 0.985 | 0.986 | 0.985 |
| 10 | GG**Hip**GG | 0.899 | 0.897 | 0.904 | 0.888 | 0.887 | 0.887 | 0.893 |
| 11 | GG**His**GG | 0.891 | 0.893 | 0.883 | 0.846 | 0.877 | 0.954 | 0.887 |
| 12 | GG**Ile**GG | 0.891 | 0.904 | 0.914 | 0.928 | 0.898 | 0.942 | 0.909 |
| 13 | GG**Leu**GG | 0.901 | 0.890 | 0.860 | 0.906 | 0.878 | 0.848 | 0.884 |
| 14 | GG**Lys**GG | 0.883 | 0.901 | 0.888 | 0.908 | 0.907 | 0.918 | 0.904 |
| 15 | GG**Met**GG | 0.826 | 0.870 | 0.833 | 0.882 | 0.868 | 0.919 | 0.869 |
| 16 | GG**Phe**GG | 0.933 | 0.940 | 0.911 | 0.959 | 0.928 | 0.927 | 0.930 |
| 17 | GG**Pro**GG | 0.989 | 0.988 | 0.984 | 0.988 | 0.985 | 0.989 | 0.988 |
| 18 | GG**Ptr**GG | 0.959 | 0.965 | 0.970 | 0.950 | 0.940 | 0.963 | 0.961 |
| 19 | GG**S1p**GG | 0.840 | 0.875 | 0.884 | 0.885 | 0.813 | 0.851 | 0.863 |
| 20 | GG**Sep**GG | 0.960 | 0.951 | 0.961 | 0.950 | 0.956 | 0.971 | 0.958 |
| 21 | GG**Ser**GG | 0.971 | 0.969 | 0.970 | 0.961 | 0.959 | 0.971 | 0.969 |
| 22 | GG**T1p**GG | 0.933 | 0.844 | 0.839 | 0.850 | 0.833 | 0.615 | 0.841 |
| 23 | GG**Thr**GG | 0.959 | 0.974 | 0.969 | 0.983 | 0.983 | 0.985 | 0.978 |
| 24 | GG**Tpo**GG | 0.932 | 0.579 | 0.689 | 0.733 | 0.825 | 0.952 | 0.779 |
| 25 | GG**Trp**GG | 0.874 | 0.902 | 0.860 | 0.918 | 0.928 | 0.904 | 0.903 |
| 26 | GG**Tyr**GG | 0.938 | 0.965 | 0.968 | 0.924 | 0.926 | 0.949 | 0.944 |
| 27 | GG**Val**GG | 0.905 | 0.897 | 0.895 | 0.838 | 0.903 | 0.893 | 0.896 |
| 28 | GG**Y1p**GG | 0.932 | 0.892 | 0.937 | 0.900 | 0.938 | 0.892 | 0.916 |
| 29 | GG**AlaAla**GG | 0.938 | 0.948 | 0.932 | 0.934 | 0.919 | 0.946 | 0.936 |
| 30 | GG**AlaArg**GG | 0.939 | 0.908 | 0.775 | 0.906 | 0.787 | 0.803 | 0.854 |
| 31 | GG**AlaAsh**GG | 0.764 | 0.800 | 0.763 | 0.897 | 0.936 | 0.924 | 0.849 |
| 32 | GG**AlaAsn**GG | 0.839 | 0.804 | 0.831 | 0.887 | 0.917 | 0.911 | 0.863 |
| 33 | GG**AlaAsp**GG | 0.840 | 0.893 | 0.900 | 0.861 | 0.822 | 0.876 | 0.868 |
| 34 | GG**AlaCys**GG | 0.871 | 0.869 | 0.807 | 0.887 | 0.824 | 0.928 | 0.870 |
| 35 | GG**AlaGlh**GG | 0.924 | 0.857 | 0.916 | 0.898 | 0.898 | 0.850 | 0.898 |
| 36 | GG**AlaGln**GG | 0.909 | 0.922 | 0.847 | 0.944 | 0.841 | 0.817 | 0.878 |
| 37 | GG**AlaGlu**GG | 0.729 | 0.728 | 0.715 | 0.889 | 0.867 | 0.854 | 0.792 |
| 38 | GG**AlaGly**GG | 0.901 | 0.896 | 0.911 | 0.910 | 0.899 | 0.884 | 0.900 |
| 39 | GG**AlaHip**GG | 0.891 | 0.925 | 0.935 | 0.913 | 0.917 | 0.933 | 0.921 |
| 40 | GG**AlaHis**GG | 0.821 | 0.814 | 0.800 | 0.898 | 0.888 | 0.918 | 0.855 |
| 41 | GG**AlaIle**GG | 0.791 | 0.851 | 0.790 | 0.805 | 0.802 | 0.797 | 0.800 |
| 42 | GG**AlaLeu**GG | 0.855 | 0.837 | 0.870 | 0.853 | 0.819 | 0.869 | 0.854 |
| 43 | GG**AlaLys**GG | 0.864 | 0.823 | 0.903 | 0.795 | 0.910 | 0.816 | 0.843 |
| 44 | GG**AlaMet**GG | 0.906 | 0.870 | 0.814 | 0.905 | 0.867 | 0.884 | 0.877 |
| 45 | GG**AlaPhe**GG | 0.910 | 0.899 | 0.896 | 0.884 | 0.890 | 0.921 | 0.897 |
| 46 | GG**AlaPro**GG | 0.953 | 0.963 | 0.953 | 0.960 | 0.959 | 0.945 | 0.956 |
| 47 | GG**AlaPtr**GG | 0.719 | 0.765 | 0.681 | 0.827 | 0.806 | 0.827 | 0.786 |
| 48 | GG**AlaS1p**GG | 0.803 | 0.787 | 0.623 | 0.776 | 0.659 | 0.628 | 0.717 |
| 49 | GG**AlaSep**GG | 0.726 | 0.833 | 0.692 | 0.822 | 0.691 | 0.764 | 0.745 |

This work is shared under a CC BY 4.0 license.

| | Peptide | $r_{replica}$(1,2) | $r_{replica}$(1,3) | $r_{replica}$(1,4) | $r_{replica}$(2,3) | $r_{replica}$(2,4) | $r_{replica}$(3,4) | Median |
|---|---|---|---|---|---|---|---|---|
| 50 | GG**AlaSer**GG | 0.886 | 0.823 | 0.902 | 0.851 | 0.893 | 0.829 | 0.869 |
| 51 | GG**AlaT1p**GG | 0.705 | 0.377 | 0.465 | 0.737 | 0.761 | 0.888 | 0.721 |
| 52 | GG**AlaThr**GG | 0.842 | 0.859 | 0.844 | 0.866 | 0.877 | 0.884 | 0.862 |
| 53 | GG**AlaTpo**GG | 0.713 | 0.741 | 0.667 | 0.531 | 0.832 | 0.363 | 0.690 |
| 54 | GG**AlaTrp**GG | 0.838 | 0.774 | 0.840 | 0.774 | 0.756 | 0.844 | 0.806 |
| 55 | GG**AlaTyr**GG | 0.906 | 0.892 | 0.726 | 0.873 | 0.774 | 0.702 | 0.823 |
| 56 | GG**AlaVal**GG | 0.927 | 0.910 | 0.856 | 0.894 | 0.850 | 0.833 | 0.875 |
| 57 | GG**AlaY1p**GG | 0.843 | 0.786 | 0.830 | 0.839 | 0.879 | 0.850 | 0.841 |
| 58 | GG**ArgAla**GG | 0.888 | 0.914 | 0.878 | 0.901 | 0.851 | 0.899 | 0.894 |
| 59 | GG**ArgArg**GG | 0.899 | 0.889 | 0.863 | 0.894 | 0.860 | 0.866 | 0.878 |
| 60 | GG**ArgAsh**GG | 0.879 | 0.709 | 0.828 | 0.680 | 0.796 | 0.739 | 0.767 |
| 61 | GG**ArgAsn**GG | 0.889 | 0.860 | 0.808 | 0.877 | 0.821 | 0.854 | 0.857 |
| 62 | GG**ArgAsp**GG | 0.894 | 0.834 | 0.868 | 0.807 | 0.891 | 0.826 | 0.851 |
| 63 | GG**ArgCys**GG | 0.776 | 0.854 | 0.800 | 0.842 | 0.938 | 0.832 | 0.837 |
| 64 | GG**ArgGlh**GG | 0.895 | 0.925 | 0.901 | 0.882 | 0.859 | 0.912 | 0.898 |
| 65 | GG**ArgGln**GG | 0.915 | 0.782 | 0.902 | 0.777 | 0.896 | 0.834 | 0.865 |
| 66 | GG**ArgGlu**GG | 0.885 | 0.807 | 0.855 | 0.818 | 0.867 | 0.858 | 0.856 |
| 67 | GG**ArgGly**GG | 0.851 | 0.774 | 0.772 | 0.712 | 0.727 | 0.827 | 0.773 |
| 68 | GG**ArgHip**GG | 0.945 | 0.852 | 0.939 | 0.865 | 0.914 | 0.836 | 0.890 |
| 69 | GG**ArgHis**GG | 0.784 | 0.870 | 0.817 | 0.826 | 0.834 | 0.839 | 0.830 |
| 70 | GG**ArgIle**GG | 0.901 | 0.900 | 0.893 | 0.907 | 0.909 | 0.889 | 0.900 |
| 71 | GG**ArgLeu**GG | 0.803 | 0.669 | 0.870 | 0.797 | 0.810 | 0.680 | 0.800 |
| 72 | GG**ArgLys**GG | 0.828 | 0.906 | 0.885 | 0.854 | 0.790 | 0.853 | 0.853 |
| 73 | GG**ArgMet**GG | 0.876 | 0.889 | 0.916 | 0.862 | 0.880 | 0.902 | 0.885 |
| 74 | GG**ArgPhe**GG | 0.833 | 0.833 | 0.801 | 0.870 | 0.795 | 0.853 | 0.833 |
| 75 | GG**ArgPro**GG | 0.921 | 0.894 | 0.929 | 0.915 | 0.925 | 0.910 | 0.918 |
| 76 | GG**ArgPtr**GG | 0.593 | 0.377 | 0.666 | 0.814 | 0.884 | 0.766 | 0.716 |
| 77 | GG**ArgS1p**GG | 0.790 | 0.706 | 0.729 | 0.669 | 0.775 | 0.524 | 0.717 |
| 78 | GG**ArgSep**GG | 0.747 | 0.767 | 0.804 | 0.341 | 0.926 | 0.395 | 0.757 |
| 79 | GG**ArgSer**GG | 0.894 | 0.899 | 0.829 | 0.900 | 0.830 | 0.834 | 0.864 |
| 80 | GG**ArgT1p**GG | 0.743 | 0.757 | 0.765 | 0.683 | 0.768 | 0.683 | 0.750 |
| 81 | GG**ArgThr**GG | 0.582 | 0.559 | 0.697 | 0.834 | 0.881 | 0.828 | 0.762 |
| 82 | GG**ArgTpo**GG | 0.695 | 0.626 | 0.718 | 0.797 | 0.825 | 0.802 | 0.758 |
| 83 | GG**ArgTrp**GG | 0.765 | 0.827 | 0.802 | 0.837 | 0.834 | 0.861 | 0.830 |
| 84 | GG**ArgTyr**GG | 0.800 | 0.813 | 0.797 | 0.787 | 0.753 | 0.845 | 0.798 |
| 85 | GG**ArgVal**GG | 0.883 | 0.849 | 0.863 | 0.937 | 0.883 | 0.852 | 0.873 |
| 86 | GG**ArgY1p**GG | 0.723 | 0.840 | 0.614 | 0.725 | 0.716 | 0.576 | 0.719 |
| 87 | GG**AshAla**GG | 0.886 | 0.898 | 0.905 | 0.870 | 0.887 | 0.901 | 0.892 |
| 88 | GG**AshArg**GG | 0.770 | 0.782 | 0.806 | 0.865 | 0.856 | 0.843 | 0.824 |
| 89 | GG**AshAsh**GG | 0.450 | 0.432 | 0.572 | 0.902 | 0.825 | 0.840 | 0.699 |
| 90 | GG**AshAsn**GG | 0.916 | 0.732 | 0.926 | 0.788 | 0.902 | 0.732 | 0.845 |
| 91 | GG**AshAsp**GG | 0.680 | 0.851 | 0.347 | 0.637 | 0.697 | 0.446 | 0.659 |
| 92 | GG**AshCys**GG | 0.893 | 0.898 | 0.922 | 0.919 | 0.940 | 0.907 | 0.913 |
| 93 | GG**AshGlh**GG | 0.817 | 0.911 | 0.868 | 0.820 | 0.768 | 0.842 | 0.831 |
| 94 | GG**AshGln**GG | 0.791 | 0.868 | 0.881 | 0.854 | 0.854 | 0.894 | 0.861 |
| 95 | GG**AshGlu**GG | 0.538 | 0.839 | 0.824 | 0.506 | 0.575 | 0.834 | 0.700 |
| 96 | GG**AshGly**GG | 0.904 | 0.885 | 0.916 | 0.872 | 0.905 | 0.892 | 0.898 |
| 97 | GG**AshHip**GG | 0.880 | 0.881 | 0.885 | 0.814 | 0.858 | 0.867 | 0.873 |
| 98 | GG**AshHis**GG | 0.659 | 0.869 | 0.700 | 0.774 | 0.778 | 0.831 | 0.776 |
| 99 | GG**AshIle**GG | 0.889 | 0.745 | 0.726 | 0.677 | 0.735 | 0.695 | 0.731 |

This work is shared under a CC BY 4.0 license.

| | Peptide | $r_{replica}$(1,2) | $r_{replica}$(1,3) | $r_{replica}$(1,4) | $r_{replica}$(2,3) | $r_{replica}$(2,4) | $r_{replica}$(3,4) | Median |
|---|---|---|---|---|---|---|---|---|
| 100 | GG**AshLeu**GG | 0.846 | 0.835 | 0.789 | 0.841 | 0.876 | 0.844 | 0.843 |
| 101 | GG**AshLys**GG | 0.933 | 0.929 | 0.698 | 0.907 | 0.749 | 0.686 | 0.828 |
| 102 | GG**AshMet**GG | 0.830 | 0.845 | 0.813 | 0.816 | 0.787 | 0.825 | 0.820 |
| 103 | GG**AshPhe**GG | 0.879 | 0.868 | 0.831 | 0.858 | 0.838 | 0.777 | 0.848 |
| 104 | GG**AshPro**GG | 0.896 | 0.936 | 0.881 | 0.933 | 0.880 | 0.904 | 0.900 |
| 105 | GG**AshPtr**GG | 0.670 | 0.624 | 0.724 | 0.760 | 0.673 | 0.813 | 0.699 |
| 106 | GG**AshS1p**GG | 0.836 | 0.859 | 0.861 | 0.875 | 0.860 | 0.836 | 0.859 |
| 107 | GG**AshSep**GG | 0.782 | 0.814 | 0.877 | 0.621 | 0.773 | 0.782 | 0.782 |
| 108 | GG**AshSer**GG | 0.856 | 0.817 | 0.840 | 0.879 | 0.869 | 0.845 | 0.851 |
| 109 | GG**AshT1p**GG | 0.807 | 0.454 | 0.931 | 0.669 | 0.822 | 0.564 | 0.738 |
| 110 | GG**AshThr**GG | 0.882 | 0.858 | 0.912 | 0.869 | 0.873 | 0.866 | 0.871 |
| 111 | GG**AshTpo**GG | 0.891 | 0.828 | 0.834 | 0.847 | 0.888 | 0.748 | 0.841 |
| 112 | GG**AshTrp**GG | 0.803 | 0.897 | 0.781 | 0.848 | 0.822 | 0.865 | 0.835 |
| 113 | GG**AshTyr**GG | 0.878 | 0.820 | 0.782 | 0.876 | 0.823 | 0.850 | 0.836 |
| 114 | GG**AshVal**GG | 0.874 | 0.881 | 0.858 | 0.832 | 0.883 | 0.838 | 0.866 |
| 115 | GG**AshY1p**GG | 0.655 | 0.814 | 0.828 | 0.602 | 0.669 | 0.786 | 0.727 |
| 116 | GG**AsnAla**GG | 0.840 | 0.827 | 0.866 | 0.834 | 0.836 | 0.869 | 0.838 |
| 117 | GG**AsnArg**GG | 0.892 | 0.885 | 0.874 | 0.909 | 0.914 | 0.852 | 0.889 |
| 118 | GG**AsnAsh**GG | 0.799 | 0.665 | 0.702 | 0.775 | 0.813 | 0.843 | 0.787 |
| 119 | GG**AsnAsn**GG | 0.880 | 0.727 | 0.876 | 0.741 | 0.874 | 0.667 | 0.807 |
| 120 | GG**AsnAsp**GG | 0.931 | 0.439 | 0.670 | 0.413 | 0.659 | 0.738 | 0.664 |
| 121 | GG**AsnCys**GG | 0.845 | 0.863 | 0.854 | 0.827 | 0.814 | 0.804 | 0.836 |
| 122 | GG**AsnGlh**GG | 0.889 | 0.912 | 0.926 | 0.860 | 0.897 | 0.912 | 0.905 |
| 123 | GG**AsnGln**GG | 0.892 | 0.889 | 0.917 | 0.885 | 0.914 | 0.908 | 0.900 |
| 124 | GG**AsnGlu**GG | 0.879 | 0.825 | 0.891 | 0.858 | 0.860 | 0.853 | 0.859 |
| 125 | GG**AsnGly**GG | 0.894 | 0.910 | 0.899 | 0.907 | 0.901 | 0.901 | 0.901 |
| 126 | GG**AsnHip**GG | 0.849 | 0.835 | 0.878 | 0.773 | 0.844 | 0.869 | 0.847 |
| 127 | GG**AsnHis**GG | 0.815 | 0.830 | 0.758 | 0.799 | 0.753 | 0.791 | 0.795 |
| 128 | GG**AsnIle**GG | 0.809 | 0.804 | 0.864 | 0.903 | 0.721 | 0.803 | 0.807 |
| 129 | GG**AsnLeu**GG | 0.823 | 0.839 | 0.856 | 0.733 | 0.751 | 0.905 | 0.831 |
| 130 | GG**AsnLys**GG | 0.859 | 0.790 | 0.837 | 0.785 | 0.803 | 0.822 | 0.812 |
| 131 | GG**AsnMet**GG | 0.898 | 0.846 | 0.799 | 0.863 | 0.856 | 0.817 | 0.851 |
| 132 | GG**AsnPhe**GG | 0.871 | 0.838 | 0.854 | 0.881 | 0.893 | 0.859 | 0.865 |
| 133 | GG**AsnPro**GG | 0.910 | 0.906 | 0.887 | 0.905 | 0.877 | 0.903 | 0.904 |
| 134 | GG**AsnPtr**GG | 0.806 | 0.821 | 0.736 | 0.878 | 0.745 | 0.748 | 0.777 |
| 135 | GG**AsnS1p**GG | 0.841 | 0.848 | 0.722 | 0.858 | 0.758 | 0.727 | 0.800 |
| 136 | GG**AsnSep**GG | 0.778 | 0.876 | 0.693 | 0.785 | 0.818 | 0.716 | 0.781 |
| 137 | GG**AsnSer**GG | 0.841 | 0.802 | 0.714 | 0.880 | 0.660 | 0.667 | 0.758 |
| 138 | GG**AsnT1p**GG | 0.829 | 0.887 | 0.920 | 0.916 | 0.886 | 0.904 | 0.896 |
| 139 | GG**AsnThr**GG | 0.888 | 0.869 | 0.881 | 0.854 | 0.859 | 0.885 | 0.875 |
| 140 | GG**AsnTpo**GG | 0.818 | 0.843 | 0.877 | 0.794 | 0.776 | 0.790 | 0.806 |
| 141 | GG**AsnTrp**GG | 0.792 | 0.806 | 0.783 | 0.804 | 0.860 | 0.822 | 0.805 |
| 142 | GG**AsnTyr**GG | 0.851 | 0.869 | 0.787 | 0.798 | 0.807 | 0.822 | 0.814 |
| 143 | GG**AsnVal**GG | 0.833 | 0.895 | 0.826 | 0.890 | 0.799 | 0.868 | 0.851 |
| 144 | GG**AsnY1p**GG | 0.787 | 0.804 | 0.797 | 0.767 | 0.809 | 0.777 | 0.792 |
| 145 | GG**AspAla**GG | 0.849 | 0.887 | 0.821 | 0.832 | 0.761 | 0.794 | 0.826 |
| 146 | GG**AspArg**GG | 0.746 | 0.720 | 0.711 | 0.731 | 0.714 | 0.736 | 0.725 |
| 147 | GG**AspAsh**GG | 0.536 | 0.458 | 0.396 | 0.606 | 0.805 | 0.676 | 0.571 |
| 148 | GG**AspAsn**GG | 0.740 | 0.819 | 0.665 | 0.738 | 0.684 | 0.634 | 0.711 |
| 149 | GG**AspAsp**GG | 0.818 | 0.789 | 0.794 | 0.810 | 0.831 | 0.835 | 0.814 |

This work is shared under a CC BY 4.0 license.

| | Peptide | $r_{replica}(1,2)$ | $r_{replica}(1,3)$ | $r_{replica}(1,4)$ | $r_{replica}(2,3)$ | $r_{replica}(2,4)$ | $r_{replica}(3,4)$ | Median |
|---|---|---|---|---|---|---|---|---|
| 150 | GG**AspCys**GG | 0.859 | 0.732 | 0.798 | 0.731 | 0.743 | 0.804 | 0.771 |
| 151 | GG**AspGlh**GG | 0.465 | 0.426 | 0.434 | 0.664 | 0.643 | 0.893 | 0.554 |
| 152 | GG**AspGln**GG | 0.517 | 0.525 | 0.577 | 0.788 | 0.726 | 0.790 | 0.652 |
| 153 | GG**AspGlu**GG | 0.745 | 0.810 | 0.831 | 0.774 | 0.832 | 0.830 | 0.820 |
| 154 | GG**AspGly**GG | 0.805 | 0.821 | 0.739 | 0.779 | 0.724 | 0.780 | 0.779 |
| 155 | GG**AspHip**GG | 0.707 | 0.713 | 0.572 | 0.752 | 0.722 | 0.714 | 0.714 |
| 156 | GG**AspHis**GG | 0.806 | 0.828 | 0.861 | 0.790 | 0.839 | 0.779 | 0.817 |
| 157 | GG**AspIle**GG | 0.901 | 0.716 | 0.878 | 0.802 | 0.882 | 0.829 | 0.853 |
| 158 | GG**AspLeu**GG | 0.665 | 0.846 | 0.642 | 0.722 | 0.508 | 0.630 | 0.654 |
| 159 | GG**AspLys**GG | 0.736 | 0.745 | 0.687 | 0.726 | 0.421 | 0.630 | 0.706 |
| 160 | GG**AspMet**GG | 0.748 | 0.703 | 0.783 | 0.738 | 0.738 | 0.811 | 0.743 |
| 161 | GG**AspPhe**GG | 0.835 | 0.838 | 0.526 | 0.855 | 0.543 | 0.596 | 0.715 |
| 162 | GG**AspPro**GG | 0.946 | 0.942 | 0.895 | 0.921 | 0.910 | 0.888 | 0.915 |
| 163 | GG**AspPtr**GG | 0.807 | 0.685 | 0.816 | 0.623 | 0.716 | 0.630 | 0.701 |
| 164 | GG**AspS1p**GG | 0.808 | 0.728 | 0.813 | 0.701 | 0.816 | 0.712 | 0.768 |
| 165 | GG**AspSep**GG | 0.847 | 0.601 | 0.525 | 0.546 | 0.719 | 0.535 | 0.574 |
| 166 | GG**AspSer**GG | 0.791 | 0.836 | 0.866 | 0.844 | 0.782 | 0.771 | 0.813 |
| 167 | GG**AspT1p**GG | 0.567 | 0.301 | 0.525 | 0.829 | 0.886 | 0.884 | 0.698 |
| 168 | GG**AspThr**GG | 0.830 | 0.842 | 0.881 | 0.703 | 0.799 | 0.876 | 0.836 |
| 169 | GG**AspTpo**GG | 0.778 | 0.832 | 0.792 | 0.852 | 0.721 | 0.684 | 0.785 |
| 170 | GG**AspTrp**GG | 0.691 | 0.747 | 0.737 | 0.757 | 0.555 | 0.621 | 0.714 |
| 171 | GG**AspTyr**GG | 0.701 | 0.711 | 0.789 | 0.869 | 0.766 | 0.776 | 0.771 |
| 172 | GG**AspVal**GG | 0.854 | 0.873 | 0.827 | 0.880 | 0.821 | 0.825 | 0.840 |
| 173 | GG**AspY1p**GG | 0.752 | 0.748 | 0.608 | 0.701 | 0.541 | 0.529 | 0.655 |
| 174 | GG**CysAla**GG | 0.900 | 0.890 | 0.890 | 0.914 | 0.921 | 0.894 | 0.897 |
| 175 | GG**CysArg**GG | 0.899 | 0.825 | 0.810 | 0.853 | 0.836 | 0.832 | 0.834 |
| 176 | GG**CysAsh**GG | 0.821 | 0.836 | 0.901 | 0.815 | 0.830 | 0.847 | 0.833 |
| 177 | GG**CysAsn**GG | 0.886 | 0.880 | 0.774 | 0.887 | 0.808 | 0.837 | 0.858 |
| 178 | GG**CysAsp**GG | 0.762 | 0.753 | 0.598 | 0.814 | 0.821 | 0.704 | 0.758 |
| 179 | GG**CysCys**GG | 0.851 | 0.760 | 0.793 | 0.883 | 0.898 | 0.892 | 0.867 |
| 180 | GG**CysGlh**GG | 0.751 | 0.807 | 0.816 | 0.843 | 0.755 | 0.808 | 0.807 |
| 181 | GG**CysGln**GG | 0.725 | 0.827 | 0.880 | 0.743 | 0.771 | 0.909 | 0.799 |
| 182 | GG**CysGlu**GG | 0.832 | 0.767 | 0.722 | 0.836 | 0.782 | 0.872 | 0.807 |
| 183 | GG**CysGly**GG | 0.791 | 0.856 | 0.866 | 0.794 | 0.839 | 0.849 | 0.844 |
| 184 | GG**CysHip**GG | 0.677 | 0.819 | 0.598 | 0.809 | 0.378 | 0.536 | 0.638 |
| 185 | GG**CysHis**GG | 0.786 | 0.752 | 0.754 | 0.809 | 0.818 | 0.834 | 0.797 |
| 186 | GG**CysIle**GG | 0.896 | 0.682 | 0.715 | 0.810 | 0.786 | 0.849 | 0.798 |
| 187 | GG**CysLeu**GG | 0.830 | 0.910 | 0.880 | 0.837 | 0.792 | 0.860 | 0.848 |
| 188 | GG**CysLys**GG | 0.777 | 0.894 | 0.907 | 0.750 | 0.785 | 0.861 | 0.823 |
| 189 | GG**CysMet**GG | 0.807 | 0.733 | 0.767 | 0.822 | 0.838 | 0.815 | 0.811 |
| 190 | GG**CysPhe**GG | 0.780 | 0.842 | 0.747 | 0.810 | 0.814 | 0.812 | 0.811 |
| 191 | GG**CysPro**GG | 0.925 | 0.930 | 0.939 | 0.938 | 0.931 | 0.909 | 0.931 |
| 192 | GG**CysPtr**GG | 0.716 | 0.738 | 0.713 | 0.782 | 0.789 | 0.754 | 0.746 |
| 193 | GG**CysS1p**GG | 0.790 | 0.835 | 0.795 | 0.757 | 0.813 | 0.800 | 0.798 |
| 194 | GG**CysSep**GG | 0.846 | 0.796 | 0.836 | 0.746 | 0.764 | 0.800 | 0.798 |
| 195 | GG**CysSer**GG | 0.763 | 0.790 | 0.768 | 0.855 | 0.804 | 0.815 | 0.797 |
| 196 | GG**CysT1p**GG | 0.344 | 0.665 | 0.855 | 0.785 | 0.603 | 0.812 | 0.725 |
| 197 | GG**CysThr**GG | 0.806 | 0.775 | 0.887 | 0.794 | 0.806 | 0.828 | 0.806 |
| 198 | GG**CysTpo**GG | 0.789 | 0.821 | 0.782 | 0.738 | 0.769 | 0.789 | 0.785 |
| 199 | GG**CysTrp**GG | 0.708 | 0.749 | 0.785 | 0.814 | 0.861 | 0.852 | 0.799 |

This work is shared under a CC BY 4.0 license.

| | Peptide | $r_{replica}$(1,2) | $r_{replica}$(1,3) | $r_{replica}$(1,4) | $r_{replica}$(2,3) | $r_{replica}$(2,4) | $r_{replica}$(3,4) | Median |
|---|---|---|---|---|---|---|---|---|
| 200 | GG**CysTyr**GG | 0.791 | 0.867 | 0.888 | 0.875 | 0.821 | 0.873 | 0.870 |
| 201 | GG**CysVal**GG | 0.808 | 0.812 | 0.788 | 0.832 | 0.838 | 0.824 | 0.818 |
| 202 | GG**CysY1p**GG | 0.757 | 0.739 | 0.817 | 0.717 | 0.807 | 0.806 | 0.782 |
| 203 | GG**GlhAla**GG | 0.901 | 0.928 | 0.875 | 0.915 | 0.910 | 0.899 | 0.906 |
| 204 | GG**GlhArg**GG | 0.707 | 0.849 | 0.645 | 0.743 | 0.718 | 0.747 | 0.730 |
| 205 | GG**GlhAsh**GG | 0.857 | 0.888 | 0.921 | 0.890 | 0.857 | 0.926 | 0.889 |
| 206 | GG**GlhAsn**GG | 0.837 | 0.915 | 0.439 | 0.807 | 0.574 | 0.400 | 0.690 |
| 207 | GG**GlhAsp**GG | 0.851 | 0.436 | 0.880 | 0.390 | 0.913 | 0.447 | 0.649 |
| 208 | GG**GlhCys**GG | 0.849 | 0.843 | 0.860 | 0.849 | 0.857 | 0.847 | 0.849 |
| 209 | GG**GlhGlh**GG | 0.906 | 0.921 | 0.852 | 0.928 | 0.885 | 0.871 | 0.896 |
| 210 | GG**GlhGln**GG | 0.847 | 0.845 | 0.864 | 0.824 | 0.799 | 0.834 | 0.839 |
| 211 | GG**GlhGlu**GG | 0.743 | 0.660 | 0.672 | 0.775 | 0.646 | 0.554 | 0.666 |
| 212 | GG**GlhGly**GG | 0.721 | 0.805 | 0.666 | 0.847 | 0.823 | 0.788 | 0.797 |
| 213 | GG**GlhHip**GG | 0.886 | 0.865 | 0.877 | 0.890 | 0.905 | 0.899 | 0.888 |
| 214 | GG**GlhHis**GG | 0.770 | 0.724 | 0.828 | 0.867 | 0.851 | 0.807 | 0.817 |
| 215 | GG**GlhIle**GG | 0.803 | 0.853 | 0.881 | 0.778 | 0.747 | 0.875 | 0.828 |
| 216 | GG**GlhLeu**GG | 0.792 | 0.888 | 0.881 | 0.794 | 0.787 | 0.897 | 0.838 |
| 217 | GG**GlhLys**GG | 0.884 | 0.852 | 0.900 | 0.918 | 0.935 | 0.923 | 0.909 |
| 218 | GG**GlhMet**GG | 0.913 | 0.494 | 0.919 | 0.364 | 0.946 | 0.436 | 0.704 |
| 219 | GG**GlhPhe**GG | 0.776 | 0.764 | 0.770 | 0.850 | 0.734 | 0.759 | 0.767 |
| 220 | GG**GlhPro**GG | 0.904 | 0.898 | 0.882 | 0.916 | 0.902 | 0.899 | 0.901 |
| 221 | GG**GlhPtr**GG | 0.601 | 0.604 | 0.718 | 0.871 | 0.536 | 0.476 | 0.602 |
| 222 | GG**GlhS1p**GG | 0.755 | 0.716 | 0.743 | 0.820 | 0.702 | 0.671 | 0.730 |
| 223 | GG**GlhSep**GG | 0.713 | 0.692 | 0.704 | 0.838 | 0.787 | 0.771 | 0.742 |
| 224 | GG**GlhSer**GG | 0.870 | 0.844 | 0.881 | 0.868 | 0.878 | 0.876 | 0.873 |
| 225 | GG**GlhT1p**GG | 0.388 | 0.725 | 0.585 | 0.394 | 0.730 | 0.592 | 0.588 |
| 226 | GG**GlhThr**GG | 0.867 | 0.858 | 0.865 | 0.846 | 0.841 | 0.868 | 0.862 |
| 227 | GG**GlhTpo**GG | 0.261 | 0.215 | 0.931 | 0.948 | 0.141 | 0.092 | 0.238 |
| 228 | GG**GlhTrp**GG | 0.737 | 0.686 | 0.822 | 0.624 | 0.748 | 0.797 | 0.743 |
| 229 | GG**GlhTyr**GG | 0.800 | 0.735 | 0.821 | 0.847 | 0.857 | 0.863 | 0.834 |
| 230 | GG**GlhVal**GG | 0.900 | 0.892 | 0.900 | 0.852 | 0.916 | 0.884 | 0.896 |
| 231 | GG**GlhY1p**GG | 0.744 | 0.862 | 0.857 | 0.780 | 0.781 | 0.848 | 0.814 |
| 232 | GG**GlnAla**GG | 0.874 | 0.915 | 0.906 | 0.880 | 0.870 | 0.930 | 0.893 |
| 233 | GG**GlnArg**GG | 0.923 | 0.919 | 0.828 | 0.936 | 0.834 | 0.847 | 0.883 |
| 234 | GG**GlnAsh**GG | 0.848 | 0.866 | 0.863 | 0.833 | 0.868 | 0.868 | 0.864 |
| 235 | GG**GlnAsn**GG | 0.736 | 0.824 | 0.838 | 0.764 | 0.721 | 0.808 | 0.786 |
| 236 | GG**GlnAsp**GG | 0.837 | 0.841 | 0.830 | 0.855 | 0.824 | 0.829 | 0.833 |
| 237 | GG**GlnCys**GG | 0.813 | 0.719 | 0.885 | 0.802 | 0.833 | 0.734 | 0.807 |
| 238 | GG**GlnGlh**GG | 0.893 | 0.889 | 0.630 | 0.868 | 0.588 | 0.716 | 0.792 |
| 239 | GG**GlnGln**GG | 0.705 | 0.683 | 0.854 | 0.753 | 0.818 | 0.843 | 0.786 |
| 240 | GG**GlnGlu**GG | 0.884 | 0.812 | 0.835 | 0.816 | 0.845 | 0.825 | 0.830 |
| 241 | GG**GlnGly**GG | 0.887 | 0.935 | 0.911 | 0.860 | 0.890 | 0.891 | 0.891 |
| 242 | GG**GlnHip**GG | 0.700 | 0.645 | 0.723 | 0.917 | 0.890 | 0.837 | 0.780 |
| 243 | GG**GlnHis**GG | 0.745 | 0.751 | 0.733 | 0.794 | 0.657 | 0.728 | 0.739 |
| 244 | GG**GlnIle**GG | 0.873 | 0.858 | 0.824 | 0.843 | 0.833 | 0.841 | 0.842 |
| 245 | GG**GlnLeu**GG | 0.753 | 0.757 | 0.753 | 0.878 | 0.864 | 0.866 | 0.811 |
| 246 | GG**GlnLys**GG | 0.754 | 0.887 | 0.922 | 0.807 | 0.701 | 0.864 | 0.835 |
| 247 | GG**GlnMet**GG | 0.842 | 0.879 | 0.883 | 0.881 | 0.838 | 0.877 | 0.878 |
| 248 | GG**GlnPhe**GG | 0.861 | 0.739 | 0.906 | 0.743 | 0.848 | 0.709 | 0.795 |
| 249 | GG**GlnPro**GG | 0.934 | 0.900 | 0.923 | 0.881 | 0.912 | 0.903 | 0.908 |

This work is shared under a CC BY 4.0 license.

| | Peptide | $r_{replica}$(1,2) | $r_{replica}$(1,3) | $r_{replica}$(1,4) | $r_{replica}$(2,3) | $r_{replica}$(2,4) | $r_{replica}$(3,4) | Median |
|---|---|---|---|---|---|---|---|---|
| 250 | GG**GlnPtr**GG | 0.827 | 0.784 | 0.615 | 0.837 | 0.675 | 0.810 | 0.797 |
| 251 | GG**GlnS1p**GG | 0.744 | 0.810 | 0.761 | 0.801 | 0.742 | 0.813 | 0.781 |
| 252 | GG**GlnSep**GG | 0.677 | 0.601 | 0.643 | 0.926 | 0.953 | 0.928 | 0.801 |
| 253 | GG**GlnSer**GG | 0.881 | 0.908 | 0.846 | 0.876 | 0.865 | 0.841 | 0.871 |
| 254 | GG**GlnT1p**GG | 0.687 | 0.787 | 0.518 | 0.663 | 0.792 | 0.400 | 0.675 |
| 255 | GG**GlnThr**GG | 0.875 | 0.869 | 0.838 | 0.841 | 0.823 | 0.820 | 0.839 |
| 256 | GG**GlnTpo**GG | 0.799 | 0.853 | 0.812 | 0.841 | 0.869 | 0.860 | 0.847 |
| 257 | GG**GlnTrp**GG | 0.828 | 0.716 | 0.820 | 0.721 | 0.761 | 0.776 | 0.769 |
| 258 | GG**GlnTyr**GG | 0.827 | 0.880 | 0.828 | 0.856 | 0.860 | 0.853 | 0.854 |
| 259 | GG**GlnVal**GG | 0.889 | 0.920 | 0.895 | 0.885 | 0.909 | 0.883 | 0.892 |
| 260 | GG**GlnY1p**GG | 0.885 | 0.875 | 0.879 | 0.862 | 0.907 | 0.870 | 0.877 |
| 261 | GG**GluAla**GG | 0.782 | 0.796 | 0.725 | 0.844 | 0.858 | 0.761 | 0.789 |
| 262 | GG**GluArg**GG | 0.699 | 0.730 | 0.761 | 0.772 | 0.692 | 0.752 | 0.741 |
| 263 | GG**GluAsh**GG | 0.753 | 0.756 | 0.635 | 0.874 | 0.445 | 0.451 | 0.694 |
| 264 | GG**GluAsn**GG | 0.838 | 0.853 | 0.761 | 0.835 | 0.811 | 0.801 | 0.823 |
| 265 | GG**GluAsp**GG | 0.846 | 0.669 | 0.835 | 0.790 | 0.863 | 0.748 | 0.812 |
| 266 | GG**GluCys**GG | 0.864 | 0.841 | 0.824 | 0.837 | 0.793 | 0.851 | 0.839 |
| 267 | GG**GluGlh**GG | 0.467 | 0.334 | 0.497 | 0.841 | 0.749 | 0.765 | 0.623 |
| 268 | GG**GluGln**GG | 0.732 | 0.692 | 0.661 | 0.843 | 0.752 | 0.851 | 0.742 |
| 269 | GG**GluGlu**GG | 0.784 | 0.796 | 0.794 | 0.836 | 0.799 | 0.782 | 0.795 |
| 270 | GG**GluGly**GG | 0.787 | 0.859 | 0.807 | 0.816 | 0.832 | 0.790 | 0.812 |
| 271 | GG**GluHip**GG | 0.893 | 0.553 | 0.893 | 0.514 | 0.895 | 0.540 | 0.723 |
| 272 | GG**GluHis**GG | 0.823 | 0.768 | 0.732 | 0.775 | 0.769 | 0.820 | 0.772 |
| 273 | GG**GluIle**GG | 0.818 | 0.896 | 0.860 | 0.803 | 0.815 | 0.834 | 0.826 |
| 274 | GG**GluLeu**GG | 0.821 | 0.826 | 0.474 | 0.880 | 0.568 | 0.540 | 0.694 |
| 275 | GG**GluLys**GG | 0.503 | 0.531 | 0.528 | 0.839 | 0.860 | 0.797 | 0.664 |
| 276 | GG**GluMet**GG | 0.808 | 0.800 | 0.842 | 0.822 | 0.879 | 0.793 | 0.815 |
| 277 | GG**GluPhe**GG | 0.833 | 0.715 | 0.712 | 0.716 | 0.751 | 0.690 | 0.716 |
| 278 | GG**GluPro**GG | 0.909 | 0.923 | 0.805 | 0.926 | 0.818 | 0.806 | 0.863 |
| 279 | GG**GluPtr**GG | 0.601 | 0.651 | 0.563 | 0.599 | 0.628 | 0.565 | 0.600 |
| 280 | GG**GluS1p**GG | 0.876 | 0.638 | 0.745 | 0.757 | 0.739 | 0.732 | 0.742 |
| 281 | GG**GluSep**GG | 0.881 | 0.668 | 0.907 | 0.472 | 0.897 | 0.620 | 0.775 |
| 282 | GG**GluSer**GG | 0.772 | 0.807 | 0.791 | 0.830 | 0.822 | 0.831 | 0.815 |
| 283 | GG**GluT1p**GG | 0.828 | 0.853 | 0.871 | 0.876 | 0.876 | 0.850 | 0.862 |
| 284 | GG**GluThr**GG | 0.784 | 0.838 | 0.783 | 0.771 | 0.818 | 0.789 | 0.786 |
| 285 | GG**GluTpo**GG | 0.791 | 0.564 | 0.559 | 0.670 | 0.664 | 0.777 | 0.667 |
| 286 | GG**GluTrp**GG | 0.711 | 0.593 | 0.697 | 0.576 | 0.788 | 0.727 | 0.704 |
| 287 | GG**GluTyr**GG | 0.898 | 0.900 | 0.892 | 0.877 | 0.912 | 0.881 | 0.895 |
| 288 | GG**GluVal**GG | 0.865 | 0.861 | 0.868 | 0.817 | 0.870 | 0.850 | 0.863 |
| 289 | GG**GluY1p**GG | 0.752 | 0.820 | 0.698 | 0.759 | 0.764 | 0.661 | 0.755 |
| 290 | GG**GlyAla**GG | 0.942 | 0.930 | 0.933 | 0.927 | 0.939 | 0.936 | 0.934 |
| 291 | GG**GlyArg**GG | 0.819 | 0.858 | 0.872 | 0.852 | 0.841 | 0.876 | 0.855 |
| 292 | GG**GlyAsh**GG | 0.902 | 0.799 | 0.890 | 0.850 | 0.912 | 0.862 | 0.876 |
| 293 | GG**GlyAsn**GG | 0.820 | 0.891 | 0.869 | 0.844 | 0.847 | 0.844 | 0.845 |
| 294 | GG**GlyAsp**GG | 0.763 | 0.856 | 0.776 | 0.771 | 0.531 | 0.794 | 0.773 |
| 295 | GG**GlyCys**GG | 0.895 | 0.901 | 0.886 | 0.898 | 0.891 | 0.903 | 0.896 |
| 296 | GG**GlyGlh**GG | 0.876 | 0.903 | 0.775 | 0.872 | 0.798 | 0.815 | 0.843 |
| 297 | GG**GlyGln**GG | 0.921 | 0.886 | 0.889 | 0.876 | 0.892 | 0.870 | 0.887 |
| 298 | GG**GlyGlu**GG | 0.880 | 0.840 | 0.853 | 0.863 | 0.868 | 0.863 | 0.863 |
| 299 | GG**GlyGly**GG | 0.911 | 0.922 | 0.898 | 0.930 | 0.911 | 0.937 | 0.916 |

This work is shared under a CC BY 4.0 license.

| | Peptide | $r_{replica}$(1,2) | $r_{replica}$(1,3) | $r_{replica}$(1,4) | $r_{replica}$(2,3) | $r_{replica}$(2,4) | $r_{replica}$(3,4) | Median |
|---|---|---|---|---|---|---|---|---|
| 300 | GG**GlyHip**GG | 0.849 | 0.823 | 0.829 | 0.874 | 0.926 | 0.876 | 0.861 |
| 301 | GG**GlyHis**GG | 0.840 | 0.861 | 0.848 | 0.879 | 0.874 | 0.902 | 0.867 |
| 302 | GG**GlyIle**GG | 0.858 | 0.843 | 0.900 | 0.841 | 0.884 | 0.853 | 0.855 |
| 303 | GG**GlyLeu**GG | 0.868 | 0.888 | 0.899 | 0.908 | 0.891 | 0.901 | 0.895 |
| 304 | GG**GlyLys**GG | 0.867 | 0.834 | 0.834 | 0.882 | 0.872 | 0.859 | 0.863 |
| 305 | GG**GlyMet**GG | 0.867 | 0.909 | 0.875 | 0.865 | 0.866 | 0.913 | 0.871 |
| 306 | GG**GlyPhe**GG | 0.893 | 0.910 | 0.924 | 0.890 | 0.885 | 0.919 | 0.902 |
| 307 | GG**GlyPro**GG | 0.862 | 0.883 | 0.887 | 0.919 | 0.895 | 0.933 | 0.891 |
| 308 | GG**GlyPtr**GG | 0.776 | 0.716 | 0.746 | 0.890 | 0.829 | 0.791 | 0.783 |
| 309 | GG**GlyS1p**GG | 0.772 | 0.836 | 0.790 | 0.817 | 0.718 | 0.753 | 0.781 |
| 310 | GG**GlySep**GG | 0.868 | 0.886 | 0.872 | 0.900 | 0.884 | 0.906 | 0.885 |
| 311 | GG**GlySer**GG | 0.896 | 0.867 | 0.881 | 0.893 | 0.901 | 0.876 | 0.887 |
| 312 | GG**GlyT1p**GG | 0.654 | 0.840 | 0.781 | 0.809 | 0.840 | 0.841 | 0.825 |
| 313 | GG**GlyThr**GG | 0.924 | 0.912 | 0.871 | 0.897 | 0.882 | 0.882 | 0.890 |
| 314 | GG**GlyTpo**GG | 0.846 | 0.930 | 0.581 | 0.885 | 0.791 | 0.618 | 0.819 |
| 315 | GG**GlyTrp**GG | 0.758 | 0.823 | 0.829 | 0.728 | 0.706 | 0.843 | 0.790 |
| 316 | GG**GlyTyr**GG | 0.890 | 0.897 | 0.908 | 0.901 | 0.902 | 0.893 | 0.899 |
| 317 | GG**GlyVal**GG | 0.885 | 0.889 | 0.894 | 0.853 | 0.860 | 0.865 | 0.875 |
| 318 | GG**GlyY1p**GG | 0.864 | 0.855 | 0.857 | 0.818 | 0.874 | 0.847 | 0.856 |
| 319 | GG**HipAla**GG | 0.883 | 0.900 | 0.878 | 0.902 | 0.902 | 0.920 | 0.901 |
| 320 | GG**HipArg**GG | 0.941 | 0.920 | 0.918 | 0.924 | 0.944 | 0.942 | 0.933 |
| 321 | GG**HipAsh**GG | 0.849 | 0.884 | 0.807 | 0.882 | 0.889 | 0.880 | 0.881 |
| 322 | GG**HipAsn**GG | 0.949 | 0.920 | 0.912 | 0.893 | 0.909 | 0.910 | 0.911 |
| 323 | GG**HipAsp**GG | 0.820 | 0.876 | 0.831 | 0.818 | 0.765 | 0.829 | 0.824 |
| 324 | GG**HipCys**GG | 0.904 | 0.922 | 0.917 | 0.916 | 0.878 | 0.932 | 0.917 |
| 325 | GG**HipGlh**GG | 0.939 | 0.942 | 0.791 | 0.899 | 0.835 | 0.726 | 0.867 |
| 326 | GG**HipGln**GG | 0.853 | 0.899 | 0.919 | 0.839 | 0.805 | 0.911 | 0.876 |
| 327 | GG**HipGlu**GG | 0.860 | 0.922 | 0.819 | 0.889 | 0.854 | 0.825 | 0.857 |
| 328 | GG**HipGly**GG | 0.898 | 0.873 | 0.904 | 0.874 | 0.873 | 0.861 | 0.874 |
| 329 | GG**HipHip**GG | 0.908 | 0.925 | 0.911 | 0.902 | 0.872 | 0.925 | 0.910 |
| 330 | GG**HipHis**GG | 0.847 | 0.803 | 0.868 | 0.861 | 0.826 | 0.870 | 0.854 |
| 331 | GG**HipIle**GG | 0.792 | 0.867 | 0.913 | 0.842 | 0.799 | 0.855 | 0.848 |
| 332 | GG**HipLeu**GG | 0.924 | 0.920 | 0.934 | 0.937 | 0.931 | 0.924 | 0.927 |
| 333 | GG**HipLys**GG | 0.958 | 0.951 | 0.902 | 0.924 | 0.894 | 0.889 | 0.913 |
| 334 | GG**HipMet**GG | 0.843 | 0.831 | 0.791 | 0.864 | 0.853 | 0.887 | 0.848 |
| 335 | GG**HipPhe**GG | 0.911 | 0.918 | 0.872 | 0.881 | 0.820 | 0.926 | 0.896 |
| 336 | GG**HipPro**GG | 0.888 | 0.926 | 0.894 | 0.915 | 0.912 | 0.929 | 0.914 |
| 337 | GG**HipPtr**GG | 0.765 | 0.700 | 0.727 | 0.906 | 0.916 | 0.903 | 0.834 |
| 338 | GG**HipS1p**GG | 0.608 | 0.610 | 0.840 | 0.623 | 0.605 | 0.766 | 0.617 |
| 339 | GG**HipSep**GG | 0.657 | 0.576 | 0.572 | 0.856 | 0.863 | 0.895 | 0.757 |
| 340 | GG**HipSer**GG | 0.918 | 0.905 | 0.908 | 0.914 | 0.911 | 0.903 | 0.910 |
| 341 | GG**HipT1p**GG | 0.836 | 0.741 | 0.896 | 0.691 | 0.807 | 0.802 | 0.804 |
| 342 | GG**HipThr**GG | 0.869 | 0.890 | 0.892 | 0.876 | 0.874 | 0.924 | 0.883 |
| 343 | GG**HipTpo**GG | 0.641 | 0.901 | 0.532 | 0.665 | 0.623 | 0.548 | 0.632 |
| 344 | GG**HipTrp**GG | 0.682 | 0.901 | 0.833 | 0.735 | 0.724 | 0.780 | 0.758 |
| 345 | GG**HipTyr**GG | 0.821 | 0.865 | 0.803 | 0.803 | 0.767 | 0.757 | 0.803 |
| 346 | GG**HipVal**GG | 0.921 | 0.814 | 0.929 | 0.833 | 0.912 | 0.863 | 0.888 |
| 347 | GG**HipY1p**GG | 0.735 | 0.632 | 0.771 | 0.910 | 0.872 | 0.840 | 0.806 |
| 348 | GG**HisAla**GG | 0.744 | 0.706 | 0.742 | 0.880 | 0.912 | 0.910 | 0.812 |
| 349 | GG**HisArg**GG | 0.853 | 0.836 | 0.782 | 0.790 | 0.763 | 0.872 | 0.813 |

This work is shared under a CC BY 4.0 license.

| | Peptide | $r_{replica}(1,2)$ | $r_{replica}(1,3)$ | $r_{replica}(1,4)$ | $r_{replica}(2,3)$ | $r_{replica}(2,4)$ | $r_{replica}(3,4)$ | Median |
|---|---|---|---|---|---|---|---|---|
| 350 | GG**HisAsh**GG | 0.735 | 0.748 | 0.777 | 0.666 | 0.667 | 0.909 | 0.742 |
| 351 | GG**HisAsn**GG | 0.768 | 0.771 | 0.603 | 0.847 | 0.888 | 0.809 | 0.790 |
| 352 | GG**HisAsp**GG | 0.727 | 0.606 | 0.758 | 0.780 | 0.872 | 0.755 | 0.756 |
| 353 | GG**HisCys**GG | 0.890 | 0.938 | 0.888 | 0.880 | 0.889 | 0.886 | 0.889 |
| 354 | GG**HisGlh**GG | 0.622 | 0.651 | 0.590 | 0.917 | 0.913 | 0.878 | 0.764 |
| 355 | GG**HisGln**GG | 0.863 | 0.762 | 0.804 | 0.582 | 0.850 | 0.648 | 0.783 |
| 356 | GG**HisGlu**GG | 0.785 | 0.756 | 0.827 | 0.863 | 0.825 | 0.898 | 0.826 |
| 357 | GG**HisGly**GG | 0.793 | 0.772 | 0.799 | 0.795 | 0.827 | 0.768 | 0.794 |
| 358 | GG**HisHip**GG | 0.844 | 0.794 | 0.788 | 0.842 | 0.877 | 0.829 | 0.836 |
| 359 | GG**HisHis**GG | 0.767 | 0.748 | 0.832 | 0.779 | 0.780 | 0.876 | 0.780 |
| 360 | GG**HisIle**GG | 0.816 | 0.861 | 0.777 | 0.749 | 0.810 | 0.772 | 0.794 |
| 361 | GG**HisLeu**GG | 0.809 | 0.757 | 0.758 | 0.791 | 0.691 | 0.610 | 0.758 |
| 362 | GG**HisLys**GG | 0.688 | 0.886 | 0.854 | 0.664 | 0.668 | 0.896 | 0.771 |
| 363 | GG**HisMet**GG | 0.872 | 0.680 | 0.898 | 0.473 | 0.851 | 0.644 | 0.765 |
| 364 | GG**HisPhe**GG | 0.782 | 0.898 | 0.802 | 0.892 | 0.858 | 0.864 | 0.861 |
| 365 | GG**HisPro**GG | 0.882 | 0.883 | 0.878 | 0.869 | 0.904 | 0.902 | 0.882 |
| 366 | GG**HisPtr**GG | 0.599 | 0.956 | 0.624 | 0.618 | 0.596 | 0.601 | 0.609 |
| 367 | GG**HisS1p**GG | 0.917 | 0.905 | 0.875 | 0.906 | 0.936 | 0.868 | 0.906 |
| 368 | GG**HisSep**GG | 0.833 | 0.913 | 0.763 | 0.796 | 0.868 | 0.745 | 0.814 |
| 369 | GG**HisSer**GG | 0.880 | 0.881 | 0.845 | 0.912 | 0.799 | 0.819 | 0.863 |
| 370 | GG**HisT1p**GG | 0.786 | 0.772 | 0.832 | 0.908 | 0.923 | 0.909 | 0.870 |
| 371 | GG**HisThr**GG | 0.818 | 0.867 | 0.723 | 0.794 | 0.843 | 0.701 | 0.806 |
| 372 | GG**HisTpo**GG | 0.624 | 0.843 | 0.735 | 0.810 | 0.469 | 0.687 | 0.711 |
| 373 | GG**HisTrp**GG | 0.892 | 0.879 | 0.907 | 0.914 | 0.900 | 0.889 | 0.896 |
| 374 | GG**HisTyr**GG | 0.729 | 0.577 | 0.679 | 0.681 | 0.885 | 0.716 | 0.699 |
| 375 | GG**HisVal**GG | 0.899 | 0.902 | 0.910 | 0.925 | 0.944 | 0.930 | 0.918 |
| 376 | GG**HisY1p**GG | 0.538 | 0.466 | 0.586 | 0.752 | 0.868 | 0.712 | 0.649 |
| 377 | GG**IleAla**GG | 0.873 | 0.873 | 0.874 | 0.842 | 0.876 | 0.858 | 0.873 |
| 378 | GG**IleArg**GG | 0.896 | 0.866 | 0.902 | 0.852 | 0.927 | 0.866 | 0.881 |
| 379 | GG**IleAsh**GG | 0.707 | 0.800 | 0.798 | 0.837 | 0.703 | 0.748 | 0.773 |
| 380 | GG**IleAsn**GG | 0.776 | 0.845 | 0.798 | 0.836 | 0.851 | 0.854 | 0.840 |
| 381 | GG**IleAsp**GG | 0.625 | 0.667 | 0.549 | 0.850 | 0.619 | 0.612 | 0.622 |
| 382 | GG**IleCys**GG | 0.802 | 0.805 | 0.894 | 0.908 | 0.829 | 0.847 | 0.838 |
| 383 | GG**IleGlh**GG | 0.880 | 0.745 | 0.829 | 0.661 | 0.773 | 0.764 | 0.769 |
| 384 | GG**IleGln**GG | 0.803 | 0.886 | 0.858 | 0.861 | 0.918 | 0.889 | 0.873 |
| 385 | GG**IleGlu**GG | 0.748 | 0.693 | 0.709 | 0.841 | 0.845 | 0.841 | 0.794 |
| 386 | GG**IleGly**GG | 0.882 | 0.849 | 0.878 | 0.883 | 0.854 | 0.853 | 0.866 |
| 387 | GG**IleHip**GG | 0.846 | 0.857 | 0.891 | 0.808 | 0.868 | 0.879 | 0.862 |
| 388 | GG**IleHis**GG | 0.786 | 0.823 | 0.803 | 0.785 | 0.726 | 0.837 | 0.795 |
| 389 | GG**IleIle**GG | 0.850 | 0.849 | 0.867 | 0.841 | 0.818 | 0.820 | 0.845 |
| 390 | GG**IleLeu**GG | 0.635 | 0.800 | 0.825 | 0.673 | 0.505 | 0.822 | 0.737 |
| 391 | GG**IleLys**GG | 0.825 | 0.873 | 0.934 | 0.843 | 0.812 | 0.858 | 0.850 |
| 392 | GG**IleMet**GG | 0.807 | 0.890 | 0.644 | 0.780 | 0.688 | 0.628 | 0.734 |
| 393 | GG**IlePhe**GG | 0.865 | 0.851 | 0.858 | 0.847 | 0.869 | 0.824 | 0.854 |
| 394 | GG**IlePro**GG | 0.843 | 0.891 | 0.896 | 0.876 | 0.843 | 0.892 | 0.884 |
| 395 | GG**IlePtr**GG | 0.842 | 0.769 | 0.756 | 0.693 | 0.666 | 0.744 | 0.750 |
| 396 | GG**IleS1p**GG | 0.834 | 0.815 | 0.843 | 0.755 | 0.810 | 0.818 | 0.817 |
| 397 | GG**IleSep**GG | 0.587 | 0.877 | 0.889 | 0.562 | 0.527 | 0.863 | 0.725 |
| 398 | GG**IleSer**GG | 0.791 | 0.795 | 0.668 | 0.807 | 0.720 | 0.790 | 0.791 |
| 399 | GG**IleT1p**GG | 0.805 | 0.742 | 0.733 | 0.616 | 0.763 | 0.553 | 0.738 |

This work is shared under a CC BY 4.0 license.

| | Peptide | $r_{replica}$(1,2) | $r_{replica}$(1,3) | $r_{replica}$(1,4) | $r_{replica}$(2,3) | $r_{replica}$(2,4) | $r_{replica}$(3,4) | Median |
|---|---|---|---|---|---|---|---|---|
| 400 | GG**IleThr**GG | 0.856 | 0.824 | 0.847 | 0.871 | 0.824 | 0.828 | 0.837 |
| 401 | GG**IleTpo**GG | 0.934 | 0.917 | 0.141 | 0.949 | 0.395 | 0.374 | 0.656 |
| 402 | GG**IleTrp**GG | 0.688 | 0.597 | 0.766 | 0.747 | 0.867 | 0.798 | 0.756 |
| 403 | GG**IleTyr**GG | 0.808 | 0.745 | 0.554 | 0.827 | 0.727 | 0.779 | 0.762 |
| 404 | GG**IleVal**GG | 0.898 | 0.857 | 0.873 | 0.882 | 0.874 | 0.882 | 0.878 |
| 405 | GG**IleY1p**GG | 0.844 | 0.786 | 0.803 | 0.833 | 0.814 | 0.798 | 0.808 |
| 406 | GG**LeuAla**GG | 0.878 | 0.832 | 0.900 | 0.832 | 0.860 | 0.785 | 0.846 |
| 407 | GG**LeuArg**GG | 0.918 | 0.759 | 0.929 | 0.705 | 0.911 | 0.741 | 0.835 |
| 408 | GG**LeuAsh**GG | 0.786 | 0.864 | 0.895 | 0.843 | 0.792 | 0.885 | 0.854 |
| 409 | GG**LeuAsn**GG | 0.783 | 0.673 | 0.742 | 0.550 | 0.861 | 0.431 | 0.707 |
| 410 | GG**LeuAsp**GG | 0.810 | 0.819 | 0.845 | 0.851 | 0.806 | 0.789 | 0.815 |
| 411 | GG**LeuCys**GG | 0.837 | 0.822 | 0.795 | 0.803 | 0.819 | 0.801 | 0.811 |
| 412 | GG**LeuGlh**GG | 0.941 | 0.463 | 0.954 | 0.573 | 0.923 | 0.530 | 0.748 |
| 413 | GG**LeuGln**GG | 0.734 | 0.830 | 0.818 | 0.740 | 0.801 | 0.783 | 0.792 |
| 414 | GG**LeuGlu**GG | 0.572 | 0.543 | 0.616 | 0.850 | 0.820 | 0.803 | 0.709 |
| 415 | GG**LeuGly**GG | 0.910 | 0.919 | 0.929 | 0.928 | 0.918 | 0.920 | 0.920 |
| 416 | GG**LeuHip**GG | 0.880 | 0.742 | 0.851 | 0.805 | 0.880 | 0.832 | 0.842 |
| 417 | GG**LeuHis**GG | 0.797 | 0.770 | 0.865 | 0.734 | 0.791 | 0.700 | 0.781 |
| 418 | GG**LeuIle**GG | 0.843 | 0.848 | 0.859 | 0.880 | 0.893 | 0.904 | 0.870 |
| 419 | GG**LeuLeu**GG | 0.684 | 0.898 | 0.882 | 0.635 | 0.676 | 0.853 | 0.768 |
| 420 | GG**LeuLys**GG | 0.806 | 0.827 | 0.805 | 0.845 | 0.862 | 0.843 | 0.835 |
| 421 | GG**LeuMet**GG | 0.903 | 0.850 | 0.852 | 0.829 | 0.848 | 0.852 | 0.851 |
| 422 | GG**LeuPhe**GG | 0.796 | 0.774 | 0.874 | 0.811 | 0.827 | 0.871 | 0.819 |
| 423 | GG**LeuPro**GG | 0.877 | 0.888 | 0.847 | 0.908 | 0.906 | 0.919 | 0.897 |
| 424 | GG**LeuPtr**GG | 0.850 | 0.744 | 0.859 | 0.739 | 0.875 | 0.766 | 0.808 |
| 425 | GG**LeuS1p**GG | 0.775 | 0.450 | 0.705 | 0.413 | 0.822 | 0.414 | 0.577 |
| 426 | GG**LeuSep**GG | 0.802 | 0.881 | 0.823 | 0.819 | 0.825 | 0.841 | 0.824 |
| 427 | GG**LeuSer**GG | 0.841 | 0.836 | 0.833 | 0.814 | 0.795 | 0.840 | 0.835 |
| 428 | GG**LeuT1p**GG | 0.608 | 0.383 | 0.341 | 0.640 | 0.594 | 0.706 | 0.601 |
| 429 | GG**LeuThr**GG | 0.785 | 0.789 | 0.788 | 0.853 | 0.806 | 0.832 | 0.798 |
| 430 | GG**LeuTpo**GG | 0.415 | 0.844 | 0.405 | 0.730 | 0.902 | 0.704 | 0.717 |
| 431 | GG**LeuTrp**GG | 0.722 | 0.679 | 0.817 | 0.822 | 0.709 | 0.696 | 0.716 |
| 432 | GG**LeuTyr**GG | 0.867 | 0.845 | 0.858 | 0.761 | 0.839 | 0.864 | 0.852 |
| 433 | GG**LeuVal**GG | 0.895 | 0.876 | 0.757 | 0.838 | 0.775 | 0.730 | 0.807 |
| 434 | GG**LeuY1p**GG | 0.662 | 0.833 | 0.860 | 0.691 | 0.651 | 0.792 | 0.742 |
| 435 | GG**LysAla**GG | 0.930 | 0.900 | 0.918 | 0.923 | 0.916 | 0.923 | 0.920 |
| 436 | GG**LysArg**GG | 0.882 | 0.870 | 0.895 | 0.885 | 0.945 | 0.898 | 0.890 |
| 437 | GG**LysAsh**GG | 0.936 | 0.911 | 0.925 | 0.877 | 0.939 | 0.901 | 0.918 |
| 438 | GG**LysAsn**GG | 0.869 | 0.885 | 0.832 | 0.844 | 0.846 | 0.864 | 0.855 |
| 439 | GG**LysAsp**GG | 0.807 | 0.906 | 0.868 | 0.826 | 0.868 | 0.882 | 0.868 |
| 440 | GG**LysCys**GG | 0.866 | 0.906 | 0.771 | 0.866 | 0.773 | 0.785 | 0.825 |
| 441 | GG**LysGlh**GG | 0.935 | 0.882 | 0.881 | 0.916 | 0.893 | 0.909 | 0.901 |
| 442 | GG**LysGln**GG | 0.869 | 0.726 | 0.878 | 0.793 | 0.917 | 0.822 | 0.845 |
| 443 | GG**LysGlu**GG | 0.915 | 0.750 | 0.706 | 0.789 | 0.759 | 0.833 | 0.774 |
| 444 | GG**LysGly**GG | 0.854 | 0.849 | 0.861 | 0.849 | 0.901 | 0.873 | 0.857 |
| 445 | GG**LysHip**GG | 0.886 | 0.824 | 0.881 | 0.941 | 0.930 | 0.895 | 0.890 |
| 446 | GG**LysHis**GG | 0.858 | 0.848 | 0.882 | 0.935 | 0.903 | 0.909 | 0.892 |
| 447 | GG**LysIle**GG | 0.863 | 0.900 | 0.921 | 0.902 | 0.877 | 0.919 | 0.901 |
| 448 | GG**LysLeu**GG | 0.841 | 0.927 | 0.842 | 0.838 | 0.818 | 0.859 | 0.841 |
| 449 | GG**LysLys**GG | 0.917 | 0.938 | 0.929 | 0.922 | 0.931 | 0.945 | 0.930 |

This work is shared under a CC BY 4.0 license.

| | **Peptide** | $r_{replica}$**(1,2)** | $r_{replica}$**(1,3)** | $r_{replica}$**(1,4)** | $r_{replica}$**(2,3)** | $r_{replica}$**(2,4)** | $r_{replica}$**(3,4)** | **Median** |
|---|---|---|---|---|---|---|---|---|
| 450 | GG**LysMet**GG | 0.935 | 0.951 | 0.920 | 0.941 | 0.944 | 0.941 | 0.941 |
| 451 | GG**LysPhe**GG | 0.831 | 0.916 | 0.838 | 0.773 | 0.866 | 0.751 | 0.834 |
| 452 | GG**LysPro**GG | 0.936 | 0.925 | 0.933 | 0.935 | 0.927 | 0.912 | 0.930 |
| 453 | GG**LysPtr**GG | 0.857 | 0.854 | 0.661 | 0.888 | 0.657 | 0.705 | 0.779 |
| 454 | GG**LysS1p**GG | 0.740 | 0.561 | 0.766 | 0.539 | 0.645 | 0.747 | 0.693 |
| 455 | GG**LysSep**GG | 0.829 | 0.868 | 0.872 | 0.816 | 0.853 | 0.853 | 0.853 |
| 456 | GG**LysSer**GG | 0.889 | 0.909 | 0.789 | 0.890 | 0.849 | 0.807 | 0.869 |
| 457 | GG**LysT1p**GG | 0.779 | 0.692 | 0.761 | 0.705 | 0.850 | 0.669 | 0.733 |
| 458 | GG**LysThr**GG | 0.847 | 0.868 | 0.849 | 0.900 | 0.894 | 0.873 | 0.870 |
| 459 | GG**LysTpo**GG | 0.753 | 0.881 | 0.898 | 0.825 | 0.741 | 0.867 | 0.846 |
| 460 | GG**LysTrp**GG | 0.593 | 0.820 | 0.748 | 0.740 | 0.711 | 0.784 | 0.744 |
| 461 | GG**LysTyr**GG | 0.854 | 0.868 | 0.836 | 0.819 | 0.822 | 0.835 | 0.835 |
| 462 | GG**LysVal**GG | 0.903 | 0.931 | 0.932 | 0.912 | 0.922 | 0.925 | 0.923 |
| 463 | GG**LysY1p**GG | 0.724 | 0.849 | 0.843 | 0.800 | 0.776 | 0.860 | 0.822 |
| 464 | GG**MetAla**GG | 0.891 | 0.731 | 0.909 | 0.806 | 0.916 | 0.789 | 0.848 |
| 465 | GG**MetArg**GG | 0.651 | 0.714 | 0.712 | 0.797 | 0.863 | 0.787 | 0.751 |
| 466 | GG**MetAsh**GG | 0.822 | 0.770 | 0.777 | 0.799 | 0.814 | 0.865 | 0.806 |
| 467 | GG**MetAsn**GG | 0.840 | 0.873 | 0.610 | 0.916 | 0.571 | 0.559 | 0.725 |
| 468 | GG**MetAsp**GG | 0.918 | 0.876 | 0.859 | 0.915 | 0.864 | 0.870 | 0.873 |
| 469 | GG**MetCys**GG | 0.799 | 0.832 | 0.776 | 0.846 | 0.826 | 0.814 | 0.820 |
| 470 | GG**MetGlh**GG | 0.835 | 0.854 | 0.738 | 0.838 | 0.745 | 0.746 | 0.790 |
| 471 | GG**MetGln**GG | 0.898 | 0.889 | 0.909 | 0.877 | 0.909 | 0.851 | 0.894 |
| 472 | GG**MetGlu**GG | 0.873 | 0.883 | 0.786 | 0.866 | 0.747 | 0.676 | 0.826 |
| 473 | GG**MetGly**GG | 0.854 | 0.838 | 0.830 | 0.893 | 0.806 | 0.821 | 0.834 |
| 474 | GG**MetHip**GG | 0.868 | 0.933 | 0.906 | 0.912 | 0.927 | 0.940 | 0.920 |
| 475 | GG**MetHis**GG | 0.857 | 0.782 | 0.724 | 0.807 | 0.783 | 0.760 | 0.783 |
| 476 | GG**MetIle**GG | 0.797 | 0.910 | 0.914 | 0.770 | 0.779 | 0.908 | 0.853 |
| 477 | GG**MetLeu**GG | 0.832 | 0.603 | 0.527 | 0.621 | 0.541 | 0.893 | 0.612 |
| 478 | GG**MetLys**GG | 0.917 | 0.757 | 0.917 | 0.724 | 0.875 | 0.794 | 0.835 |
| 479 | GG**MetMet**GG | 0.865 | 0.819 | 0.834 | 0.840 | 0.826 | 0.782 | 0.830 |
| 480 | GG**MetPhe**GG | 0.765 | 0.723 | 0.725 | 0.855 | 0.882 | 0.814 | 0.790 |
| 481 | GG**MetPro**GG | 0.909 | 0.949 | 0.916 | 0.947 | 0.915 | 0.916 | 0.916 |
| 482 | GG**MetPtr**GG | 0.472 | 0.440 | 0.420 | 0.831 | 0.821 | 0.783 | 0.627 |
| 483 | GG**MetS1p**GG | 0.838 | 0.659 | 0.788 | 0.708 | 0.862 | 0.725 | 0.756 |
| 484 | GG**MetSep**GG | 0.861 | 0.806 | 0.844 | 0.873 | 0.911 | 0.868 | 0.865 |
| 485 | GG**MetSer**GG | 0.692 | 0.831 | 0.840 | 0.791 | 0.818 | 0.880 | 0.824 |
| 486 | GG**MetT1p**GG | 0.716 | 0.874 | 0.827 | 0.745 | 0.828 | 0.836 | 0.828 |
| 487 | GG**MetThr**GG | 0.840 | 0.852 | 0.883 | 0.822 | 0.805 | 0.877 | 0.846 |
| 488 | GG**MetTpo**GG | 0.644 | 0.609 | 0.798 | 0.794 | 0.469 | 0.509 | 0.627 |
| 489 | GG**MetTrp**GG | 0.895 | 0.931 | 0.592 | 0.906 | 0.568 | 0.629 | 0.762 |
| 490 | GG**MetTyr**GG | 0.858 | 0.816 | 0.834 | 0.855 | 0.898 | 0.812 | 0.845 |
| 491 | GG**MetVal**GG | 0.899 | 0.863 | 0.885 | 0.857 | 0.904 | 0.878 | 0.881 |
| 492 | GG**MetY1p**GG | 0.729 | 0.880 | 0.551 | 0.797 | 0.724 | 0.618 | 0.726 |
| 493 | GG**PheAla**GG | 0.848 | 0.850 | 0.914 | 0.883 | 0.902 | 0.898 | 0.891 |
| 494 | GG**PheArg**GG | 0.804 | 0.889 | 0.844 | 0.864 | 0.878 | 0.881 | 0.871 |
| 495 | GG**PheAsh**GG | 0.867 | 0.757 | 0.657 | 0.602 | 0.449 | 0.756 | 0.707 |
| 496 | GG**PheAsn**GG | 0.766 | 0.859 | 0.904 | 0.795 | 0.803 | 0.878 | 0.831 |
| 497 | GG**PheAsp**GG | 0.785 | 0.768 | 0.764 | 0.830 | 0.768 | 0.741 | 0.768 |
| 498 | GG**PheCys**GG | 0.806 | 0.864 | 0.877 | 0.758 | 0.896 | 0.832 | 0.848 |
| 499 | GG**PheGlh**GG | 0.339 | 0.340 | 0.578 | 0.905 | 0.806 | 0.847 | 0.692 |

This work is shared under a CC BY 4.0 license.

| | Peptide | $r_{replica}(1,2)$ | $r_{replica}(1,3)$ | $r_{replica}(1,4)$ | $r_{replica}(2,3)$ | $r_{replica}(2,4)$ | $r_{replica}(3,4)$ | Median |
|---|---|---|---|---|---|---|---|---|
| 500 | GG**PheGln**GG | 0.626 | 0.641 | 0.630 | 0.861 | 0.930 | 0.870 | 0.751 |
| 501 | GG**PheGlu**GG | 0.728 | 0.729 | 0.776 | 0.622 | 0.555 | 0.816 | 0.729 |
| 502 | GG**PheGly**GG | 0.881 | 0.897 | 0.895 | 0.895 | 0.916 | 0.894 | 0.895 |
| 503 | GG**PheHip**GG | 0.870 | 0.900 | 0.890 | 0.749 | 0.905 | 0.799 | 0.880 |
| 504 | GG**PheHis**GG | 0.798 | 0.729 | 0.880 | 0.721 | 0.815 | 0.713 | 0.764 |
| 505 | GG**PheIle**GG | 0.563 | 0.805 | 0.741 | 0.621 | 0.668 | 0.747 | 0.705 |
| 506 | GG**PheLeu**GG | 0.717 | 0.914 | 0.889 | 0.708 | 0.769 | 0.918 | 0.829 |
| 507 | GG**PheLys**GG | 0.733 | 0.796 | 0.844 | 0.815 | 0.822 | 0.854 | 0.818 |
| 508 | GG**PheMet**GG | 0.800 | 0.797 | 0.696 | 0.782 | 0.709 | 0.603 | 0.745 |
| 509 | GG**PhePhe**GG | 0.812 | 0.807 | 0.754 | 0.775 | 0.689 | 0.834 | 0.791 |
| 510 | GG**PhePro**GG | 0.921 | 0.928 | 0.924 | 0.901 | 0.909 | 0.929 | 0.922 |
| 511 | GG**PhePtr**GG | 0.786 | 0.747 | 0.751 | 0.774 | 0.808 | 0.799 | 0.780 |
| 512 | GG**PheS1p**GG | 0.603 | 0.715 | 0.810 | 0.690 | 0.659 | 0.870 | 0.702 |
| 513 | GG**PheSep**GG | 0.331 | 0.312 | 0.324 | 0.883 | 0.886 | 0.897 | 0.607 |
| 514 | GG**PheSer**GG | 0.777 | 0.707 | 0.708 | 0.783 | 0.705 | 0.780 | 0.743 |
| 515 | GG**PheT1p**GG | 0.622 | 0.843 | 0.709 | 0.849 | 0.819 | 0.880 | 0.831 |
| 516 | GG**PheThr**GG | 0.870 | 0.830 | 0.843 | 0.853 | 0.865 | 0.846 | 0.849 |
| 517 | GG**PheTpo**GG | 0.741 | 0.724 | 0.727 | 0.555 | 0.673 | 0.731 | 0.726 |
| 518 | GG**PheTrp**GG | 0.858 | 0.863 | 0.815 | 0.775 | 0.744 | 0.838 | 0.827 |
| 519 | GG**PheTyr**GG | 0.878 | 0.844 | 0.785 | 0.920 | 0.782 | 0.753 | 0.815 |
| 520 | GG**PheVal**GG | 0.853 | 0.857 | 0.835 | 0.895 | 0.897 | 0.921 | 0.876 |
| 521 | GG**PheY1p**GG | 0.756 | 0.784 | 0.832 | 0.625 | 0.734 | 0.739 | 0.748 |
| 522 | GG**ProAla**GG | 0.927 | 0.874 | 0.903 | 0.876 | 0.913 | 0.943 | 0.908 |
| 523 | GG**ProArg**GG | 0.815 | 0.700 | 0.802 | 0.849 | 0.924 | 0.894 | 0.832 |
| 524 | GG**ProAsh**GG | 0.642 | 0.911 | 0.936 | 0.708 | 0.621 | 0.867 | 0.787 |
| 525 | GG**ProAsn**GG | 0.761 | 0.817 | 0.840 | 0.780 | 0.804 | 0.942 | 0.811 |
| 526 | GG**ProAsp**GG | 0.650 | 0.679 | 0.737 | 0.778 | 0.831 | 0.723 | 0.730 |
| 527 | GG**ProCys**GG | 0.935 | 0.854 | 0.911 | 0.869 | 0.887 | 0.902 | 0.895 |
| 528 | GG**ProGlh**GG | 0.956 | 0.912 | 0.934 | 0.922 | 0.950 | 0.922 | 0.928 |
| 529 | GG**ProGln**GG | 0.388 | 0.918 | 0.927 | 0.558 | 0.387 | 0.913 | 0.735 |
| 530 | GG**ProGlu**GG | 0.839 | 0.851 | 0.820 | 0.909 | 0.677 | 0.744 | 0.830 |
| 531 | GG**ProGly**GG | 0.865 | 0.898 | 0.910 | 0.919 | 0.926 | 0.933 | 0.915 |
| 532 | GG**ProHip**GG | 0.746 | 0.791 | 0.895 | 0.918 | 0.831 | 0.843 | 0.837 |
| 533 | GG**ProHis**GG | 0.806 | 0.871 | 0.849 | 0.734 | 0.786 | 0.796 | 0.801 |
| 534 | GG**ProIle**GG | 0.865 | 0.681 | 0.926 | 0.811 | 0.853 | 0.664 | 0.832 |
| 535 | GG**ProLeu**GG | 0.931 | 0.333 | 0.668 | 0.316 | 0.662 | 0.773 | 0.665 |
| 536 | GG**ProLys**GG | 0.954 | 0.720 | 0.707 | 0.738 | 0.731 | 0.886 | 0.734 |
| 537 | GG**ProMet**GG | 0.588 | 0.808 | 0.810 | 0.596 | 0.782 | 0.784 | 0.783 |
| 538 | GG**ProPhe**GG | 0.939 | 0.506 | 0.618 | 0.422 | 0.524 | 0.922 | 0.571 |
| 539 | GG**ProPro**GG | 0.917 | 0.523 | 0.781 | 0.511 | 0.825 | 0.761 | 0.771 |
| 540 | GG**ProPtr**GG | 0.736 | 0.782 | 0.852 | 0.789 | 0.824 | 0.785 | 0.787 |
| 541 | GG**ProS1p**GG | 0.760 | 0.712 | 0.817 | 0.790 | 0.789 | 0.830 | 0.790 |
| 542 | GG**ProSep**GG | 0.619 | 0.759 | 0.549 | 0.773 | 0.733 | 0.772 | 0.746 |
| 543 | GG**ProSer**GG | 0.788 | 0.909 | 0.927 | 0.669 | 0.746 | 0.901 | 0.845 |
| 544 | GG**ProT1p**GG | 0.859 | 0.844 | 0.744 | 0.764 | 0.623 | 0.887 | 0.804 |
| 545 | GG**ProThr**GG | 0.813 | 0.926 | 0.849 | 0.826 | 0.792 | 0.876 | 0.837 |
| 546 | GG**ProTpo**GG | 0.838 | 0.478 | 0.671 | 0.410 | 0.621 | 0.878 | 0.646 |
| 547 | GG**ProTrp**GG | 0.869 | 0.839 | 0.686 | 0.747 | 0.605 | 0.629 | 0.716 |
| 548 | GG**ProTyr**GG | 0.857 | 0.806 | 0.687 | 0.811 | 0.693 | 0.685 | 0.750 |
| 549 | GG**ProVal**GG | 0.955 | 0.952 | 0.955 | 0.924 | 0.938 | 0.973 | 0.953 |

This work is shared under a CC BY 4.0 license.

| | Peptide | $r_{replica}(1,2)$ | $r_{replica}(1,3)$ | $r_{replica}(1,4)$ | $r_{replica}(2,3)$ | $r_{replica}(2,4)$ | $r_{replica}(3,4)$ | Median |
|---|---|---|---|---|---|---|---|---|
| 550 | GG**ProY1p**GG | 0.892 | 0.894 | 0.715 | 0.880 | 0.750 | 0.703 | 0.815 |
| 551 | GG**PtrAla**GG | 0.452 | 0.547 | 0.404 | 0.852 | 0.791 | 0.838 | 0.669 |
| 552 | GG**PtrArg**GG | 0.362 | 0.928 | 0.706 | 0.477 | 0.337 | 0.632 | 0.555 |
| 553 | GG**PtrAsh**GG | 0.440 | 0.768 | 0.855 | 0.485 | 0.540 | 0.691 | 0.615 |
| 554 | GG**PtrAsn**GG | 0.821 | 0.643 | 0.893 | 0.661 | 0.826 | 0.716 | 0.769 |
| 555 | GG**PtrAsp**GG | 0.525 | 0.646 | 0.438 | 0.691 | 0.739 | 0.649 | 0.648 |
| 556 | GG**PtrCys**GG | 0.763 | 0.706 | 0.692 | 0.559 | 0.789 | 0.373 | 0.699 |
| 557 | GG**PtrGlh**GG | 0.840 | 0.662 | 0.840 | 0.313 | 0.859 | 0.568 | 0.751 |
| 558 | GG**PtrGln**GG | 0.717 | 0.662 | 0.772 | 0.805 | 0.602 | 0.645 | 0.689 |
| 559 | GG**PtrGlu**GG | 0.676 | 0.815 | 0.822 | 0.679 | 0.780 | 0.735 | 0.758 |
| 560 | GG**PtrGly**GG | 0.839 | 0.763 | 0.797 | 0.820 | 0.841 | 0.843 | 0.830 |
| 561 | GG**PtrHip**GG | 0.744 | 0.778 | 0.835 | 0.808 | 0.441 | 0.478 | 0.761 |
| 562 | GG**PtrHis**GG | 0.847 | 0.737 | 0.806 | 0.676 | 0.876 | 0.707 | 0.772 |
| 563 | GG**PtrIle**GG | 0.746 | 0.338 | 0.727 | 0.359 | 0.815 | 0.376 | 0.552 |
| 564 | GG**PtrLeu**GG | 0.774 | 0.765 | 0.672 | 0.887 | 0.640 | 0.609 | 0.719 |
| 565 | GG**PtrLys**GG | 0.780 | 0.495 | 0.390 | 0.599 | 0.522 | 0.875 | 0.560 |
| 566 | GG**PtrMet**GG | 0.823 | 0.368 | 0.455 | 0.443 | 0.558 | 0.826 | 0.506 |
| 567 | GG**PtrPhe**GG | 0.726 | 0.831 | 0.441 | 0.777 | 0.656 | 0.507 | 0.691 |
| 568 | GG**PtrPro**GG | 0.786 | 0.884 | 0.868 | 0.785 | 0.911 | 0.854 | 0.861 |
| 569 | GG**PtrPtr**GG | 0.451 | 0.667 | 0.533 | 0.770 | 0.833 | 0.873 | 0.718 |
| 570 | GG**PtrS1p**GG | 0.506 | 0.487 | 0.724 | 0.610 | 0.556 | 0.730 | 0.583 |
| 571 | GG**PtrSep**GG | 0.909 | 0.870 | 0.888 | 0.928 | 0.879 | 0.897 | 0.892 |
| 572 | GG**PtrSer**GG | 0.712 | 0.832 | 0.823 | 0.702 | 0.667 | 0.784 | 0.748 |
| 573 | GG**PtrT1p**GG | 0.846 | 0.515 | 0.734 | 0.583 | 0.795 | 0.639 | 0.687 |
| 574 | GG**PtrThr**GG | 0.943 | 0.372 | 0.872 | 0.494 | 0.909 | 0.664 | 0.768 |
| 575 | GG**PtrTpo**GG | 0.800 | 0.728 | 0.633 | 0.673 | 0.721 | 0.713 | 0.717 |
| 576 | GG**PtrTrp**GG | 0.507 | 0.780 | 0.662 | 0.610 | 0.434 | 0.589 | 0.599 |
| 577 | GG**PtrTyr**GG | 0.649 | 0.824 | 0.735 | 0.748 | 0.785 | 0.793 | 0.767 |
| 578 | GG**PtrVal**GG | 0.547 | 0.502 | 0.495 | 0.688 | 0.832 | 0.685 | 0.616 |
| 579 | GG**PtrY1p**GG | 0.751 | 0.647 | 0.515 | 0.717 | 0.456 | 0.607 | 0.627 |
| 580 | GG**S1pAla**GG | 0.834 | 0.815 | 0.839 | 0.842 | 0.839 | 0.844 | 0.839 |
| 581 | GG**S1pArg**GG | 0.837 | 0.572 | 0.789 | 0.447 | 0.725 | 0.612 | 0.669 |
| 582 | GG**S1pAsh**GG | 0.864 | 0.922 | 0.604 | 0.836 | 0.637 | 0.510 | 0.737 |
| 583 | GG**S1pAsn**GG | 0.767 | 0.811 | 0.794 | 0.817 | 0.768 | 0.805 | 0.800 |
| 584 | GG**S1pAsp**GG | 0.737 | 0.805 | 0.857 | 0.827 | 0.824 | 0.815 | 0.819 |
| 585 | GG**S1pCys**GG | 0.740 | 0.898 | 0.775 | 0.740 | 0.781 | 0.772 | 0.773 |
| 586 | GG**S1pGlh**GG | 0.764 | 0.790 | 0.799 | 0.839 | 0.849 | 0.864 | 0.819 |
| 587 | GG**S1pGln**GG | 0.870 | 0.885 | 0.831 | 0.848 | 0.844 | 0.866 | 0.857 |
| 588 | GG**S1pGlu**GG | 0.806 | 0.829 | 0.680 | 0.806 | 0.655 | 0.624 | 0.743 |
| 589 | GG**S1pGly**GG | 0.816 | 0.873 | 0.866 | 0.851 | 0.830 | 0.898 | 0.859 |
| 590 | GG**S1pHip**GG | 0.861 | 0.770 | 0.877 | 0.846 | 0.811 | 0.751 | 0.828 |
| 591 | GG**S1pHis**GG | 0.747 | 0.768 | 0.819 | 0.792 | 0.775 | 0.847 | 0.784 |
| 592 | GG**S1pIle**GG | 0.857 | 0.499 | 0.729 | 0.515 | 0.786 | 0.470 | 0.622 |
| 593 | GG**S1pLeu**GG | 0.669 | 0.659 | 0.614 | 0.683 | 0.686 | 0.632 | 0.664 |
| 594 | GG**S1pLys**GG | 0.724 | 0.926 | 0.621 | 0.757 | 0.573 | 0.588 | 0.673 |
| 595 | GG**S1pMet**GG | 0.750 | 0.769 | 0.777 | 0.768 | 0.762 | 0.793 | 0.769 |
| 596 | GG**S1pPhe**GG | 0.859 | 0.754 | 0.802 | 0.776 | 0.806 | 0.823 | 0.804 |
| 597 | GG**S1pPro**GG | 0.890 | 0.910 | 0.879 | 0.905 | 0.883 | 0.908 | 0.898 |
| 598 | GG**S1pPtr**GG | 0.790 | 0.682 | 0.784 | 0.587 | 0.743 | 0.665 | 0.712 |
| 599 | GG**S1pS1p**GG | 0.820 | 0.697 | 0.738 | 0.785 | 0.738 | 0.870 | 0.762 |

This work is shared under a CC BY 4.0 license.

| | Peptide | $r_{replica}$(1,2) | $r_{replica}$(1,3) | $r_{replica}$(1,4) | $r_{replica}$(2,3) | $r_{replica}$(2,4) | $r_{replica}$(3,4) | Median |
|---|---|---|---|---|---|---|---|---|
| 600 | GG**S1pSep**GG | 0.670 | 0.882 | 0.783 | 0.707 | 0.779 | 0.809 | 0.781 |
| 601 | GG**S1pSer**GG | 0.810 | 0.886 | 0.896 | 0.711 | 0.765 | 0.888 | 0.848 |
| 602 | GG**S1pT1p**GG | 0.862 | 0.843 | 0.555 | 0.899 | 0.456 | 0.430 | 0.699 |
| 603 | GG**S1pThr**GG | 0.895 | 0.784 | 0.859 | 0.730 | 0.789 | 0.740 | 0.787 |
| 604 | GG**S1pTpo**GG | 0.651 | 0.468 | 0.671 | 0.840 | 0.811 | 0.750 | 0.711 |
| 605 | GG**S1pTrp**GG | 0.857 | 0.784 | 0.849 | 0.813 | 0.826 | 0.694 | 0.819 |
| 606 | GG**S1pTyr**GG | 0.764 | 0.554 | 0.666 | 0.694 | 0.802 | 0.682 | 0.688 |
| 607 | GG**S1pVal**GG | 0.716 | 0.794 | 0.744 | 0.849 | 0.747 | 0.821 | 0.771 |
| 608 | GG**S1pY1p**GG | 0.696 | 0.674 | 0.742 | 0.695 | 0.710 | 0.682 | 0.696 |
| 609 | GG**SepAla**GG | 0.920 | 0.880 | 0.888 | 0.870 | 0.884 | 0.858 | 0.882 |
| 610 | GG**SepArg**GG | 0.907 | 0.816 | 0.895 | 0.831 | 0.915 | 0.827 | 0.863 |
| 611 | GG**SepAsh**GG | 0.772 | 0.760 | 0.814 | 0.793 | 0.786 | 0.866 | 0.790 |
| 612 | GG**SepAsn**GG | 0.872 | 0.876 | 0.912 | 0.883 | 0.873 | 0.876 | 0.876 |
| 613 | GG**SepAsp**GG | 0.784 | 0.886 | 0.879 | 0.793 | 0.773 | 0.898 | 0.836 |
| 614 | GG**SepCys**GG | 0.891 | 0.817 | 0.859 | 0.799 | 0.849 | 0.796 | 0.833 |
| 615 | GG**SepGlh**GG | 0.909 | 0.885 | 0.870 | 0.886 | 0.845 | 0.863 | 0.877 |
| 616 | GG**SepGln**GG | 0.898 | 0.869 | 0.885 | 0.845 | 0.858 | 0.899 | 0.877 |
| 617 | GG**SepGlu**GG | 0.846 | 0.802 | 0.838 | 0.832 | 0.750 | 0.826 | 0.829 |
| 618 | GG**SepGly**GG | 0.863 | 0.832 | 0.747 | 0.841 | 0.771 | 0.767 | 0.801 |
| 619 | GG**SepHip**GG | 0.593 | 0.662 | 0.758 | 0.501 | 0.548 | 0.811 | 0.627 |
| 620 | GG**SepHis**GG | 0.545 | 0.680 | 0.618 | 0.712 | 0.817 | 0.796 | 0.696 |
| 621 | GG**SepIle**GG | 0.753 | 0.759 | 0.835 | 0.843 | 0.840 | 0.848 | 0.837 |
| 622 | GG**SepLeu**GG | 0.783 | 0.807 | 0.786 | 0.778 | 0.868 | 0.764 | 0.785 |
| 623 | GG**SepLys**GG | 0.901 | 0.897 | 0.921 | 0.911 | 0.925 | 0.922 | 0.916 |
| 624 | GG**SepMet**GG | 0.757 | 0.830 | 0.855 | 0.732 | 0.764 | 0.875 | 0.797 |
| 625 | GG**SepPhe**GG | 0.875 | 0.844 | 0.824 | 0.860 | 0.796 | 0.774 | 0.834 |
| 626 | GG**SepPro**GG | 0.960 | 0.949 | 0.956 | 0.946 | 0.968 | 0.951 | 0.953 |
| 627 | GG**SepPtr**GG | 0.843 | 0.377 | 0.751 | 0.357 | 0.742 | 0.615 | 0.678 |
| 628 | GG**SepS1p**GG | 0.868 | 0.864 | 0.820 | 0.832 | 0.842 | 0.818 | 0.837 |
| 629 | GG**SepSep**GG | 0.908 | 0.842 | 0.899 | 0.877 | 0.884 | 0.880 | 0.882 |
| 630 | GG**SepSer**GG | 0.842 | 0.896 | 0.916 | 0.847 | 0.838 | 0.897 | 0.872 |
| 631 | GG**SepT1p**GG | 0.370 | 0.924 | 0.661 | 0.413 | 0.772 | 0.684 | 0.673 |
| 632 | GG**SepThr**GG | 0.831 | 0.726 | 0.746 | 0.803 | 0.779 | 0.823 | 0.791 |
| 633 | GG**SepTpo**GG | 0.864 | 0.428 | 0.306 | 0.580 | 0.426 | 0.819 | 0.504 |
| 634 | GG**SepTrp**GG | 0.890 | 0.525 | 0.651 | 0.572 | 0.703 | 0.845 | 0.677 |
| 635 | GG**SepTyr**GG | 0.598 | 0.654 | 0.664 | 0.820 | 0.885 | 0.834 | 0.742 |
| 636 | GG**SepVal**GG | 0.774 | 0.791 | 0.790 | 0.752 | 0.869 | 0.812 | 0.790 |
| 637 | GG**SepY1p**GG | 0.818 | 0.663 | 0.776 | 0.646 | 0.698 | 0.683 | 0.690 |
| 638 | GG**SerAla**GG | 0.842 | 0.867 | 0.880 | 0.906 | 0.882 | 0.894 | 0.881 |
| 639 | GG**SerArg**GG | 0.802 | 0.771 | 0.496 | 0.845 | 0.636 | 0.702 | 0.736 |
| 640 | GG**SerAsh**GG | 0.810 | 0.749 | 0.798 | 0.765 | 0.878 | 0.763 | 0.781 |
| 641 | GG**SerAsn**GG | 0.877 | 0.865 | 0.861 | 0.848 | 0.848 | 0.789 | 0.854 |
| 642 | GG**SerAsp**GG | 0.800 | 0.870 | 0.840 | 0.767 | 0.686 | 0.853 | 0.820 |
| 643 | GG**SerCys**GG | 0.892 | 0.872 | 0.875 | 0.868 | 0.868 | 0.894 | 0.874 |
| 644 | GG**SerGlh**GG | 0.665 | 0.741 | 0.622 | 0.722 | 0.807 | 0.627 | 0.693 |
| 645 | GG**SerGln**GG | 0.803 | 0.743 | 0.770 | 0.836 | 0.822 | 0.798 | 0.800 |
| 646 | GG**SerGlu**GG | 0.662 | 0.845 | 0.867 | 0.612 | 0.626 | 0.886 | 0.753 |
| 647 | GG**SerGly**GG | 0.765 | 0.850 | 0.882 | 0.765 | 0.745 | 0.868 | 0.807 |
| 648 | GG**SerHip**GG | 0.817 | 0.836 | 0.858 | 0.804 | 0.837 | 0.920 | 0.836 |
| 649 | GG**SerHis**GG | 0.749 | 0.770 | 0.790 | 0.800 | 0.797 | 0.850 | 0.794 |

This work is shared under a CC BY 4.0 license.

| | Peptide | $r_{replica}$(1,2) | $r_{replica}$(1,3) | $r_{replica}$(1,4) | $r_{replica}$(2,3) | $r_{replica}$(2,4) | $r_{replica}$(3,4) | Median |
|---|---|---|---|---|---|---|---|---|
| 650 | GG**SerIle**GG | 0.862 | 0.888 | 0.820 | 0.862 | 0.733 | 0.793 | 0.841 |
| 651 | GG**SerLeu**GG | 0.837 | 0.807 | 0.802 | 0.823 | 0.890 | 0.799 | 0.815 |
| 652 | GG**SerLys**GG | 0.823 | 0.836 | 0.850 | 0.922 | 0.914 | 0.939 | 0.882 |
| 653 | GG**SerMet**GG | 0.855 | 0.857 | 0.922 | 0.857 | 0.885 | 0.878 | 0.868 |
| 654 | GG**SerPhe**GG | 0.898 | 0.877 | 0.870 | 0.879 | 0.877 | 0.897 | 0.878 |
| 655 | GG**SerPro**GG | 0.922 | 0.914 | 0.931 | 0.926 | 0.903 | 0.903 | 0.918 |
| 656 | GG**SerPtr**GG | 0.855 | 0.752 | 0.752 | 0.743 | 0.682 | 0.740 | 0.748 |
| 657 | GG**SerS1p**GG | 0.794 | 0.790 | 0.795 | 0.777 | 0.845 | 0.797 | 0.794 |
| 658 | GG**SerSep**GG | 0.723 | 0.556 | 0.546 | 0.363 | 0.355 | 0.914 | 0.551 |
| 659 | GG**SerSer**GG | 0.784 | 0.835 | 0.795 | 0.843 | 0.856 | 0.847 | 0.839 |
| 660 | GG**SerT1p**GG | 0.445 | 0.872 | 0.633 | 0.599 | 0.826 | 0.764 | 0.699 |
| 661 | GG**SerThr**GG | 0.813 | 0.803 | 0.777 | 0.854 | 0.799 | 0.832 | 0.808 |
| 662 | GG**SerTpo**GG | 0.749 | 0.883 | 0.863 | 0.840 | 0.573 | 0.732 | 0.794 |
| 663 | GG**SerTrp**GG | 0.849 | 0.804 | 0.706 | 0.834 | 0.768 | 0.875 | 0.819 |
| 664 | GG**SerTyr**GG | 0.807 | 0.788 | 0.861 | 0.831 | 0.843 | 0.809 | 0.820 |
| 665 | GG**SerVal**GG | 0.719 | 0.816 | 0.775 | 0.809 | 0.728 | 0.818 | 0.792 |
| 666 | GG**SerY1p**GG | 0.759 | 0.814 | 0.839 | 0.760 | 0.795 | 0.796 | 0.795 |
| 667 | GG**T1pAla**GG | 0.865 | 0.939 | 0.807 | 0.906 | 0.913 | 0.853 | 0.885 |
| 668 | GG**T1pArg**GG | 0.273 | 0.306 | 0.751 | 0.961 | 0.771 | 0.764 | 0.758 |
| 669 | GG**T1pAsh**GG | 0.559 | 0.411 | 0.550 | 0.817 | 0.797 | 0.824 | 0.678 |
| 670 | GG**T1pAsn**GG | 0.467 | 0.824 | 0.605 | 0.546 | 0.896 | 0.522 | 0.575 |
| 671 | GG**T1pAsp**GG | 0.898 | 0.894 | 0.879 | 0.889 | 0.907 | 0.887 | 0.892 |
| 672 | GG**T1pCys**GG | 0.909 | 0.942 | 0.482 | 0.939 | 0.423 | 0.369 | 0.696 |
| 673 | GG**T1pGlh**GG | 0.646 | 0.550 | 0.604 | 0.864 | 0.819 | 0.789 | 0.718 |
| 674 | GG**T1pGln**GG | 0.810 | 0.848 | 0.872 | 0.876 | 0.795 | 0.824 | 0.836 |
| 675 | GG**T1pGlu**GG | 0.590 | 0.759 | 0.441 | 0.860 | 0.887 | 0.790 | 0.775 |
| 676 | GG**T1pGly**GG | 0.898 | 0.573 | 0.898 | 0.635 | 0.916 | 0.543 | 0.766 |
| 677 | GG**T1pHip**GG | 0.869 | 0.224 | 0.664 | 0.128 | 0.927 | 0.006 | 0.444 |
| 678 | GG**T1pHis**GG | 0.848 | 0.575 | 0.776 | 0.430 | 0.801 | 0.742 | 0.759 |
| 679 | GG**T1pIle**GG | 0.529 | 0.841 | 0.581 | 0.435 | 0.770 | 0.515 | 0.555 |
| 680 | GG**T1pLeu**GG | 0.605 | 0.767 | 0.878 | 0.777 | 0.325 | 0.538 | 0.686 |
| 681 | GG**T1pLys**GG | 0.641 | 0.693 | 0.887 | 0.827 | 0.751 | 0.806 | 0.778 |
| 682 | GG**T1pMet**GG | 0.519 | 0.713 | 0.776 | 0.832 | 0.886 | 0.890 | 0.804 |
| 683 | GG**T1pPhe**GG | 0.909 | 0.885 | 0.649 | 0.867 | 0.761 | 0.742 | 0.814 |
| 684 | GG**T1pPro**GG | 0.866 | 0.903 | 0.864 | 0.868 | 0.861 | 0.911 | 0.867 |
| 685 | GG**T1pPtr**GG | 0.716 | 0.722 | 0.603 | 0.683 | 0.561 | 0.684 | 0.683 |
| 686 | GG**T1pS1p**GG | 0.592 | 0.578 | 0.308 | 0.737 | 0.702 | 0.671 | 0.631 |
| 687 | GG**T1pSep**GG | 0.866 | 0.463 | 0.476 | 0.659 | 0.645 | 0.799 | 0.652 |
| 688 | GG**T1pSer**GG | 0.726 | 0.921 | 0.395 | 0.777 | 0.350 | 0.527 | 0.627 |
| 689 | GG**T1pT1p**GG | 0.709 | 0.876 | 0.766 | 0.630 | 0.291 | 0.784 | 0.737 |
| 690 | GG**T1pThr**GG | 0.922 | 0.698 | 0.759 | 0.559 | 0.630 | 0.912 | 0.728 |
| 691 | GG**T1pTpo**GG | 0.916 | 0.334 | 0.753 | 0.313 | 0.727 | 0.778 | 0.740 |
| 692 | GG**T1pTrp**GG | 0.842 | 0.696 | 0.951 | 0.662 | 0.844 | 0.768 | 0.805 |
| 693 | GG**T1pTyr**GG | 0.620 | 0.614 | 0.815 | 0.431 | 0.758 | 0.538 | 0.617 |
| 694 | GG**T1pVal**GG | 0.845 | 0.853 | 0.825 | 0.830 | 0.733 | 0.843 | 0.837 |
| 695 | GG**T1pY1p**GG | 0.452 | 0.547 | 0.614 | 0.458 | 0.428 | 0.857 | 0.503 |
| 696 | GG**ThrAla**GG | 0.876 | 0.887 | 0.839 | 0.905 | 0.865 | 0.871 | 0.874 |
| 697 | GG**ThrArg**GG | 0.784 | 0.799 | 0.766 | 0.840 | 0.931 | 0.846 | 0.819 |
| 698 | GG**ThrAsh**GG | 0.600 | 0.789 | 0.783 | 0.733 | 0.753 | 0.837 | 0.768 |
| 699 | GG**ThrAsn**GG | 0.820 | 0.839 | 0.847 | 0.847 | 0.848 | 0.838 | 0.843 |

This work is shared under a CC BY 4.0 license.

| | Peptide | $r_{replica}(1,2)$ | $r_{replica}(1,3)$ | $r_{replica}(1,4)$ | $r_{replica}(2,3)$ | $r_{replica}(2,4)$ | $r_{replica}(3,4)$ | Median |
|---|---|---|---|---|---|---|---|---|
| 700 | GG**ThrAsp**GG | 0.747 | 0.667 | 0.762 | 0.720 | 0.865 | 0.740 | 0.743 |
| 701 | GG**ThrCys**GG | 0.785 | 0.768 | 0.715 | 0.796 | 0.806 | 0.832 | 0.791 |
| 702 | GG**ThrGlh**GG | 0.917 | 0.905 | 0.874 | 0.880 | 0.901 | 0.851 | 0.891 |
| 703 | GG**ThrGln**GG | 0.840 | 0.844 | 0.765 | 0.780 | 0.766 | 0.696 | 0.773 |
| 704 | GG**ThrGlu**GG | 0.811 | 0.864 | 0.886 | 0.859 | 0.871 | 0.881 | 0.867 |
| 705 | GG**ThrGly**GG | 0.884 | 0.869 | 0.868 | 0.847 | 0.876 | 0.879 | 0.873 |
| 706 | GG**ThrHip**GG | 0.770 | 0.756 | 0.890 | 0.665 | 0.818 | 0.799 | 0.785 |
| 707 | GG**ThrHis**GG | 0.837 | 0.884 | 0.861 | 0.831 | 0.748 | 0.837 | 0.837 |
| 708 | GG**ThrIle**GG | 0.825 | 0.859 | 0.853 | 0.839 | 0.823 | 0.866 | 0.846 |
| 709 | GG**ThrLeu**GG | 0.863 | 0.615 | 0.826 | 0.716 | 0.821 | 0.641 | 0.769 |
| 710 | GG**ThrLys**GG | 0.878 | 0.775 | 0.855 | 0.649 | 0.833 | 0.754 | 0.804 |
| 711 | GG**ThrMet**GG | 0.788 | 0.820 | 0.830 | 0.799 | 0.877 | 0.830 | 0.825 |
| 712 | GG**ThrPhe**GG | 0.730 | 0.658 | 0.592 | 0.850 | 0.798 | 0.805 | 0.764 |
| 713 | GG**ThrPro**GG | 0.947 | 0.946 | 0.951 | 0.936 | 0.944 | 0.934 | 0.945 |
| 714 | GG**ThrPtr**GG | 0.707 | 0.710 | 0.764 | 0.583 | 0.756 | 0.646 | 0.708 |
| 715 | GG**ThrS1p**GG | 0.602 | 0.626 | 0.604 | 0.538 | 0.499 | 0.388 | 0.570 |
| 716 | GG**ThrSep**GG | 0.815 | 0.873 | 0.823 | 0.813 | 0.811 | 0.811 | 0.814 |
| 717 | GG**ThrSer**GG | 0.844 | 0.808 | 0.844 | 0.847 | 0.825 | 0.774 | 0.835 |
| 718 | GG**ThrT1p**GG | 0.724 | 0.745 | 0.675 | 0.880 | 0.448 | 0.538 | 0.699 |
| 719 | GG**ThrThr**GG | 0.850 | 0.815 | 0.836 | 0.801 | 0.893 | 0.819 | 0.827 |
| 720 | GG**ThrTpo**GG | 0.281 | 0.179 | 0.322 | 0.902 | 0.918 | 0.921 | 0.612 |
| 721 | GG**ThrTrp**GG | 0.650 | 0.619 | 0.693 | 0.703 | 0.861 | 0.757 | 0.698 |
| 722 | GG**ThrTyr**GG | 0.836 | 0.816 | 0.839 | 0.870 | 0.846 | 0.822 | 0.837 |
| 723 | GG**ThrVal**GG | 0.784 | 0.788 | 0.844 | 0.705 | 0.807 | 0.709 | 0.786 |
| 724 | GG**ThrY1p**GG | 0.432 | 0.711 | 0.741 | 0.727 | 0.579 | 0.732 | 0.719 |
| 725 | GG**TpoAla**GG | 0.905 | 0.872 | 0.884 | 0.888 | 0.889 | 0.852 | 0.886 |
| 726 | GG**TpoArg**GG | 0.710 | 0.724 | 0.645 | 0.587 | 0.924 | 0.671 | 0.691 |
| 727 | GG**TpoAsh**GG | 0.730 | 0.291 | 0.628 | 0.587 | 0.024 | -0.159 | 0.439 |
| 728 | GG**TpoAsn**GG | 0.384 | 0.876 | 0.937 | 0.715 | 0.118 | 0.717 | 0.716 |
| 729 | GG**TpoAsp**GG | 0.264 | 0.958 | 0.950 | 0.112 | 0.136 | 0.963 | 0.607 |
| 730 | GG**TpoCys**GG | 0.567 | 0.598 | 0.940 | 0.693 | 0.554 | 0.560 | 0.583 |
| 731 | GG**TpoGlh**GG | 0.740 | 0.890 | 0.381 | 0.741 | 0.577 | 0.345 | 0.658 |
| 732 | GG**TpoGln**GG | 0.499 | 0.600 | 0.513 | 0.879 | 0.744 | 0.837 | 0.672 |
| 733 | GG**TpoGlu**GG | 0.790 | 0.740 | 0.440 | 0.819 | 0.654 | 0.572 | 0.697 |
| 734 | GG**TpoGly**GG | 0.433 | 0.635 | 0.777 | 0.847 | 0.697 | 0.795 | 0.737 |
| 735 | GG**TpoHip**GG | 0.950 | 0.692 | 0.521 | 0.662 | 0.577 | 0.366 | 0.619 |
| 736 | GG**TpoHis**GG | 0.573 | 0.836 | 0.659 | 0.451 | 0.887 | 0.609 | 0.634 |
| 737 | GG**TpoIle**GG | 0.636 | 0.578 | 0.486 | 0.681 | 0.534 | 0.823 | 0.607 |
| 738 | GG**TpoLeu**GG | 0.968 | 0.545 | 0.303 | 0.599 | 0.384 | 0.815 | 0.572 |
| 739 | GG**TpoLys**GG | 0.208 | 0.625 | 0.570 | 0.552 | 0.728 | 0.952 | 0.598 |
| 740 | GG**TpoMet**GG | 0.817 | 0.696 | 0.696 | 0.433 | 0.450 | 0.715 | 0.696 |
| 741 | GG**TpoPhe**GG | 0.826 | 0.810 | 0.438 | 0.735 | 0.487 | 0.626 | 0.680 |
| 742 | GG**TpoPro**GG | 0.927 | 0.446 | 0.741 | 0.474 | 0.785 | 0.799 | 0.763 |
| 743 | GG**TpoPtr**GG | 0.621 | 0.425 | 0.404 | 0.655 | 0.550 | 0.583 | 0.566 |
| 744 | GG**TpoS1p**GG | 0.961 | -0.011 | 0.741 | -0.038 | 0.739 | 0.526 | 0.633 |
| 745 | GG**TpoSep**GG | 0.443 | 0.675 | 0.409 | 0.874 | 0.942 | 0.900 | 0.774 |
| 746 | GG**TpoSer**GG | 0.695 | 0.897 | 0.845 | 0.684 | 0.606 | 0.916 | 0.770 |
| 747 | GG**TpoT1p**GG | 0.796 | 0.788 | 0.437 | 0.839 | 0.693 | 0.664 | 0.740 |
| 748 | GG**TpoThr**GG | 0.435 | 0.444 | 0.536 | 0.851 | 0.854 | 0.864 | 0.694 |
| 749 | GG**TpoTpo**GG | 0.817 | 0.904 | 0.457 | 0.750 | 0.761 | 0.393 | 0.756 |

This work is shared under a CC BY 4.0 license.

| | Peptide | $r_{replica}$(1,2) | $r_{replica}$(1,3) | $r_{replica}$(1,4) | $r_{replica}$(2,3) | $r_{replica}$(2,4) | $r_{replica}$(3,4) | Median |
|---|---|---|---|---|---|---|---|---|
| 750 | GG**TpoTrp**GG | 0.809 | 0.603 | 0.735 | 0.416 | 0.598 | 0.699 | 0.651 |
| 751 | GG**TpoTyr**GG | 0.188 | 0.809 | 0.869 | 0.622 | 0.373 | 0.843 | 0.716 |
| 752 | GG**TpoVal**GG | 0.697 | 0.782 | 0.753 | 0.727 | 0.743 | 0.788 | 0.748 |
| 753 | GG**TpoY1p**GG | 0.299 | 0.648 | 0.323 | 0.744 | 0.923 | 0.745 | 0.696 |
| 754 | GG**TrpAla**GG | 0.849 | 0.857 | 0.865 | 0.829 | 0.877 | 0.863 | 0.860 |
| 755 | GG**TrpArg**GG | 0.817 | 0.763 | 0.889 | 0.945 | 0.725 | 0.635 | 0.790 |
| 756 | GG**TrpAsh**GG | 0.915 | 0.932 | 0.513 | 0.946 | 0.460 | 0.461 | 0.714 |
| 757 | GG**TrpAsn**GG | 0.882 | 0.828 | 0.900 | 0.837 | 0.917 | 0.836 | 0.860 |
| 758 | GG**TrpAsp**GG | 0.818 | 0.823 | 0.816 | 0.825 | 0.819 | 0.804 | 0.818 |
| 759 | GG**TrpCys**GG | 0.910 | 0.743 | 0.919 | 0.662 | 0.901 | 0.633 | 0.822 |
| 760 | GG**TrpGlh**GG | 0.621 | 0.770 | 0.592 | 0.868 | 0.697 | 0.772 | 0.734 |
| 761 | GG**TrpGln**GG | 0.897 | 0.884 | 0.866 | 0.925 | 0.873 | 0.868 | 0.879 |
| 762 | GG**TrpGlu**GG | 0.564 | 0.777 | 0.577 | 0.609 | 0.451 | 0.571 | 0.574 |
| 763 | GG**TrpGly**GG | 0.887 | 0.839 | 0.809 | 0.819 | 0.807 | 0.861 | 0.829 |
| 764 | GG**TrpHip**GG | 0.597 | 0.893 | 0.822 | 0.583 | 0.580 | 0.884 | 0.710 |
| 765 | GG**TrpHis**GG | 0.763 | 0.524 | 0.827 | 0.654 | 0.657 | 0.323 | 0.655 |
| 766 | GG**TrpIle**GG | 0.768 | 0.749 | 0.911 | 0.780 | 0.788 | 0.789 | 0.784 |
| 767 | GG**TrpLeu**GG | 0.893 | 0.648 | 0.784 | 0.699 | 0.703 | 0.696 | 0.701 |
| 768 | GG**TrpLys**GG | 0.799 | 0.913 | 0.830 | 0.860 | 0.872 | 0.882 | 0.866 |
| 769 | GG**TrpMet**GG | 0.661 | 0.591 | 0.684 | 0.894 | 0.868 | 0.854 | 0.769 |
| 770 | GG**TrpPhe**GG | 0.563 | 0.520 | 0.724 | 0.447 | 0.798 | 0.579 | 0.571 |
| 771 | GG**TrpPro**GG | 0.813 | 0.840 | 0.851 | 0.868 | 0.905 | 0.925 | 0.859 |
| 772 | GG**TrpPtr**GG | 0.748 | 0.630 | 0.804 | 0.597 | 0.715 | 0.688 | 0.701 |
| 773 | GG**TrpS1p**GG | 0.782 | 0.832 | 0.816 | 0.634 | 0.678 | 0.712 | 0.747 |
| 774 | GG**TrpSep**GG | 0.851 | 0.931 | 0.926 | 0.805 | 0.853 | 0.882 | 0.867 |
| 775 | GG**TrpSer**GG | 0.841 | 0.694 | 0.783 | 0.760 | 0.864 | 0.763 | 0.773 |
| 776 | GG**TrpT1p**GG | 0.792 | 0.720 | 0.456 | 0.653 | 0.452 | 0.567 | 0.610 |
| 777 | GG**TrpThr**GG | 0.743 | 0.632 | 0.759 | 0.716 | 0.868 | 0.712 | 0.730 |
| 778 | GG**TrpTpo**GG | 0.519 | 0.506 | 0.294 | 0.680 | 0.780 | 0.788 | 0.600 |
| 779 | GG**TrpTrp**GG | 0.741 | 0.826 | 0.708 | 0.674 | 0.518 | 0.710 | 0.709 |
| 780 | GG**TrpTyr**GG | 0.490 | 0.380 | 0.362 | 0.840 | 0.858 | 0.913 | 0.665 |
| 781 | GG**TrpVal**GG | 0.900 | 0.645 | 0.677 | 0.770 | 0.779 | 0.834 | 0.774 |
| 782 | GG**TrpY1p**GG | 0.799 | 0.870 | 0.737 | 0.793 | 0.794 | 0.698 | 0.794 |
| 783 | GG**TyrAla**GG | 0.822 | 0.855 | 0.909 | 0.921 | 0.870 | 0.889 | 0.879 |
| 784 | GG**TyrArg**GG | 0.859 | 0.691 | 0.790 | 0.702 | 0.796 | 0.823 | 0.793 |
| 785 | GG**TyrAsh**GG | 0.902 | 0.875 | 0.739 | 0.863 | 0.683 | 0.784 | 0.824 |
| 786 | GG**TyrAsn**GG | 0.817 | 0.841 | 0.844 | 0.821 | 0.702 | 0.732 | 0.819 |
| 787 | GG**TyrAsp**GG | 0.645 | 0.582 | 0.851 | 0.731 | 0.700 | 0.800 | 0.715 |
| 788 | GG**TyrCys**GG | 0.874 | 0.625 | 0.814 | 0.767 | 0.895 | 0.806 | 0.810 |
| 789 | GG**TyrGlh**GG | 0.409 | 0.716 | 0.706 | 0.419 | 0.345 | 0.680 | 0.550 |
| 790 | GG**TyrGln**GG | 0.831 | 0.712 | 0.855 | 0.508 | 0.893 | 0.558 | 0.772 |
| 791 | GG**TyrGlu**GG | 0.850 | 0.739 | 0.575 | 0.743 | 0.524 | 0.356 | 0.657 |
| 792 | GG**TyrGly**GG | 0.807 | 0.861 | 0.828 | 0.877 | 0.861 | 0.863 | 0.861 |
| 793 | GG**TyrHip**GG | 0.781 | 0.778 | 0.916 | 0.839 | 0.845 | 0.826 | 0.833 |
| 794 | GG**TyrHis**GG | 0.798 | 0.774 | 0.835 | 0.648 | 0.650 | 0.893 | 0.786 |
| 795 | GG**TyrIle**GG | 0.735 | 0.777 | 0.536 | 0.929 | 0.726 | 0.740 | 0.738 |
| 796 | GG**TyrLeu**GG | 0.674 | 0.823 | 0.888 | 0.741 | 0.737 | 0.861 | 0.782 |
| 797 | GG**TyrLys**GG | 0.768 | 0.835 | 0.857 | 0.599 | 0.803 | 0.820 | 0.811 |
| 798 | GG**TyrMet**GG | 0.574 | 0.833 | 0.759 | 0.726 | 0.860 | 0.804 | 0.782 |
| 799 | GG**TyrPhe**GG | 0.888 | 0.793 | 0.768 | 0.818 | 0.764 | 0.747 | 0.780 |

This work is shared under a CC BY 4.0 license.

| | Peptide | $r_{replica}$(1,2) | $r_{replica}$(1,3) | $r_{replica}$(1,4) | $r_{replica}$(2,3) | $r_{replica}$(2,4) | $r_{replica}$(3,4) | Median |
|---|---|---|---|---|---|---|---|---|
| 800 | GG**TyrPro**GG | 0.899 | 0.822 | 0.838 | 0.883 | 0.895 | 0.926 | 0.889 |
| 801 | GG**TyrPtr**GG | 0.848 | 0.522 | 0.628 | 0.594 | 0.820 | 0.548 | 0.611 |
| 802 | GG**TyrS1p**GG | 0.766 | 0.680 | 0.836 | 0.764 | 0.899 | 0.752 | 0.765 |
| 803 | GG**TyrSep**GG | 0.546 | 0.610 | 0.400 | 0.893 | 0.824 | 0.740 | 0.675 |
| 804 | GG**TyrSer**GG | 0.896 | 0.882 | 0.919 | 0.897 | 0.899 | 0.904 | 0.898 |
| 805 | GG**TyrT1p**GG | 0.847 | 0.844 | 0.792 | 0.881 | 0.693 | 0.634 | 0.818 |
| 806 | GG**TyrThr**GG | 0.840 | 0.674 | 0.737 | 0.892 | 0.886 | 0.890 | 0.863 |
| 807 | GG**TyrTpo**GG | 0.652 | 0.666 | 0.724 | 0.724 | 0.838 | 0.689 | 0.707 |
| 808 | GG**TyrTrp**GG | 0.857 | 0.848 | 0.718 | 0.768 | 0.755 | 0.591 | 0.762 |
| 809 | GG**TyrTyr**GG | 0.809 | 0.791 | 0.825 | 0.766 | 0.789 | 0.775 | 0.790 |
| 810 | GG**TyrVal**GG | 0.890 | 0.825 | 0.847 | 0.811 | 0.823 | 0.882 | 0.836 |
| 811 | GG**TyrY1p**GG | 0.755 | 0.760 | 0.692 | 0.747 | 0.763 | 0.641 | 0.751 |
| 812 | GG**ValAla**GG | 0.931 | 0.919 | 0.887 | 0.883 | 0.883 | 0.890 | 0.888 |
| 813 | GG**ValArg**GG | 0.865 | 0.845 | 0.882 | 0.760 | 0.802 | 0.862 | 0.854 |
| 814 | GG**ValAsh**GG | 0.765 | 0.794 | 0.868 | 0.639 | 0.722 | 0.754 | 0.759 |
| 815 | GG**ValAsn**GG | 0.625 | 0.782 | 0.625 | 0.776 | 0.739 | 0.689 | 0.714 |
| 816 | GG**ValAsp**GG | 0.816 | 0.857 | 0.865 | 0.836 | 0.801 | 0.823 | 0.829 |
| 817 | GG**ValCys**GG | 0.901 | 0.898 | 0.926 | 0.901 | 0.876 | 0.874 | 0.900 |
| 818 | GG**ValGlh**GG | 0.734 | 0.780 | 0.863 | 0.586 | 0.793 | 0.698 | 0.757 |
| 819 | GG**ValGln**GG | 0.601 | 0.594 | 0.617 | 0.902 | 0.690 | 0.677 | 0.647 |
| 820 | GG**ValGlu**GG | 0.893 | 0.692 | 0.725 | 0.638 | 0.694 | 0.672 | 0.693 |
| 821 | GG**ValGly**GG | 0.897 | 0.895 | 0.828 | 0.881 | 0.854 | 0.864 | 0.873 |
| 822 | GG**ValHip**GG | 0.710 | 0.844 | 0.804 | 0.843 | 0.913 | 0.893 | 0.843 |
| 823 | GG**ValHis**GG | 0.885 | 0.862 | 0.808 | 0.870 | 0.852 | 0.839 | 0.857 |
| 824 | GG**ValIle**GG | 0.892 | 0.912 | 0.676 | 0.894 | 0.683 | 0.674 | 0.787 |
| 825 | GG**ValLeu**GG | 0.698 | 0.585 | 0.809 | 0.864 | 0.667 | 0.550 | 0.683 |
| 826 | GG**ValLys**GG | 0.853 | 0.794 | 0.823 | 0.700 | 0.934 | 0.663 | 0.809 |
| 827 | GG**ValMet**GG | 0.833 | 0.806 | 0.853 | 0.886 | 0.886 | 0.823 | 0.843 |
| 828 | GG**ValPhe**GG | 0.808 | 0.805 | 0.816 | 0.749 | 0.812 | 0.795 | 0.806 |
| 829 | GG**ValPro**GG | 0.896 | 0.877 | 0.909 | 0.911 | 0.912 | 0.915 | 0.910 |
| 830 | GG**ValPtr**GG | 0.712 | 0.708 | 0.562 | 0.707 | 0.838 | 0.638 | 0.707 |
| 831 | GG**ValS1p**GG | 0.764 | 0.759 | 0.807 | 0.773 | 0.811 | 0.763 | 0.768 |
| 832 | GG**ValSep**GG | 0.878 | 0.779 | 0.831 | 0.814 | 0.889 | 0.795 | 0.823 |
| 833 | GG**ValSer**GG | 0.919 | 0.797 | 0.880 | 0.783 | 0.865 | 0.847 | 0.856 |
| 834 | GG**ValT1p**GG | 0.856 | 0.758 | 0.726 | 0.563 | 0.521 | 0.835 | 0.742 |
| 835 | GG**ValThr**GG | 0.741 | 0.810 | 0.887 | 0.782 | 0.727 | 0.771 | 0.777 |
| 836 | GG**ValTpo**GG | 0.713 | 0.718 | 0.737 | 0.461 | 0.779 | 0.636 | 0.715 |
| 837 | GG**ValTrp**GG | 0.847 | 0.832 | 0.831 | 0.827 | 0.845 | 0.799 | 0.831 |
| 838 | GG**ValTyr**GG | 0.907 | 0.875 | 0.885 | 0.893 | 0.919 | 0.913 | 0.900 |
| 839 | GG**ValVal**GG | 0.663 | 0.949 | 0.913 | 0.657 | 0.700 | 0.906 | 0.803 |
| 840 | GG**ValY1p**GG | 0.833 | 0.745 | 0.806 | 0.739 | 0.785 | 0.776 | 0.780 |
| 841 | GG**Y1pAla**GG | 0.767 | 0.836 | 0.762 | 0.828 | 0.655 | 0.770 | 0.769 |
| 842 | GG**Y1pArg**GG | 0.448 | 0.673 | 0.914 | 0.800 | 0.456 | 0.728 | 0.700 |
| 843 | GG**Y1pAsh**GG | 0.734 | 0.738 | 0.824 | 0.908 | 0.542 | 0.537 | 0.736 |
| 844 | GG**Y1pAsn**GG | 0.895 | 0.497 | 0.856 | 0.411 | 0.870 | 0.553 | 0.704 |
| 845 | GG**Y1pAsp**GG | 0.839 | 0.805 | 0.855 | 0.859 | 0.825 | 0.803 | 0.832 |
| 846 | GG**Y1pCys**GG | 0.414 | 0.631 | 0.729 | 0.647 | 0.732 | 0.665 | 0.656 |
| 847 | GG**Y1pGlh**GG | 0.833 | 0.748 | 0.511 | 0.718 | 0.678 | 0.521 | 0.698 |
| 848 | GG**Y1pGln**GG | 0.771 | 0.756 | 0.777 | 0.908 | 0.905 | 0.921 | 0.841 |
| 849 | GG**Y1pGlu**GG | 0.630 | 0.804 | 0.720 | 0.684 | 0.930 | 0.774 | 0.747 |

This work is shared under a CC BY 4.0 license.

| | Peptide | $r_{replica}(1,2)$ | $r_{replica}(1,3)$ | $r_{replica}(1,4)$ | $r_{replica}(2,3)$ | $r_{replica}(2,4)$ | $r_{replica}(3,4)$ | Median |
|---|---|---|---|---|---|---|---|---|
| 850 | GG**Y1pGly**GG | 0.785 | 0.873 | 0.812 | 0.870 | 0.823 | 0.876 | 0.846 |
| 851 | GG**Y1pHip**GG | 0.857 | 0.531 | 0.928 | 0.591 | 0.867 | 0.572 | 0.724 |
| 852 | GG**Y1pHis**GG | 0.663 | 0.630 | 0.824 | 0.817 | 0.819 | 0.772 | 0.794 |
| 853 | GG**Y1pIle**GG | 0.876 | 0.593 | 0.835 | 0.550 | 0.840 | 0.664 | 0.749 |
| 854 | GG**Y1pLeu**GG | 0.768 | 0.698 | 0.713 | 0.859 | 0.779 | 0.828 | 0.774 |
| 855 | GG**Y1pLys**GG | 0.651 | 0.871 | 0.715 | 0.709 | 0.881 | 0.772 | 0.744 |
| 856 | GG**Y1pMet**GG | 0.744 | 0.746 | 0.606 | 0.921 | 0.669 | 0.629 | 0.706 |
| 857 | GG**Y1pPhe**GG | 0.891 | 0.667 | 0.771 | 0.756 | 0.744 | 0.767 | 0.761 |
| 858 | GG**Y1pPro**GG | 0.827 | 0.847 | 0.839 | 0.873 | 0.761 | 0.850 | 0.843 |
| 859 | GG**Y1pPtr**GG | 0.687 | 0.437 | 0.402 | 0.329 | 0.373 | 0.665 | 0.419 |
| 860 | GG**Y1pS1p**GG | 0.478 | 0.835 | 0.946 | 0.781 | 0.612 | 0.908 | 0.808 |
| 861 | GG**Y1pSep**GG | 0.788 | 0.811 | 0.526 | 0.663 | 0.593 | 0.648 | 0.655 |
| 862 | GG**Y1pSer**GG | 0.902 | 0.857 | 0.694 | 0.880 | 0.648 | 0.703 | 0.780 |
| 863 | GG**Y1pT1p**GG | 0.519 | 0.815 | 0.879 | 0.429 | 0.675 | 0.881 | 0.745 |
| 864 | GG**Y1pThr**GG | 0.866 | 0.754 | 0.715 | 0.664 | 0.639 | 0.819 | 0.734 |
| 865 | GG**Y1pTpo**GG | 0.598 | 0.613 | 0.686 | 0.554 | 0.782 | 0.560 | 0.605 |
| 866 | GG**Y1pTrp**GG | 0.455 | 0.466 | 0.916 | 0.405 | 0.447 | 0.499 | 0.460 |
| 867 | GG**Y1pTyr**GG | 0.591 | 0.795 | 0.835 | 0.471 | 0.465 | 0.849 | 0.693 |
| 868 | GG**Y1pVal**GG | 0.887 | 0.571 | 0.377 | 0.779 | 0.679 | 0.807 | 0.729 |
| 869 | GG**Y1pY1p**GG | 0.724 | 0.710 | 0.723 | 0.746 | 0.732 | 0.765 | 0.728 |
| 870 | GG**Ala**G**Ala**GG | 0.845 | 0.870 | 0.875 | 0.894 | 0.879 | 0.898 | 0.877 |
| 871 | GG**Ala**G**Arg**GG | 0.737 | 0.784 | 0.574 | 0.740 | 0.558 | 0.571 | 0.656 |
| 872 | GG**Ala**G**Ash**GG | 0.886 | 0.864 | 0.901 | 0.895 | 0.909 | 0.894 | 0.895 |
| 873 | GG**Ala**G**Asn**GG | 0.890 | 0.914 | 0.929 | 0.881 | 0.871 | 0.891 | 0.891 |
| 874 | GG**Ala**G**Asp**GG | 0.803 | 0.820 | 0.874 | 0.776 | 0.794 | 0.864 | 0.812 |
| 875 | GG**Ala**G**Cys**GG | 0.826 | 0.808 | 0.836 | 0.790 | 0.773 | 0.791 | 0.799 |
| 876 | GG**Ala**G**Glh**GG | 0.845 | 0.847 | 0.843 | 0.874 | 0.869 | 0.861 | 0.854 |
| 877 | GG**Ala**G**Gln**GG | 0.797 | 0.824 | 0.842 | 0.781 | 0.837 | 0.794 | 0.810 |
| 878 | GG**Ala**G**Glu**GG | 0.877 | 0.807 | 0.851 | 0.789 | 0.882 | 0.725 | 0.829 |
| 879 | GG**Ala**G**Gly**GG | 0.938 | 0.904 | 0.904 | 0.912 | 0.881 | 0.895 | 0.904 |
| 880 | GG**Ala**G**Hip**GG | 0.828 | 0.837 | 0.865 | 0.843 | 0.806 | 0.872 | 0.840 |
| 881 | GG**Ala**G**His**GG | 0.848 | 0.819 | 0.822 | 0.855 | 0.800 | 0.863 | 0.835 |
| 882 | GG**Ala**G**Ile**GG | 0.849 | 0.845 | 0.864 | 0.864 | 0.877 | 0.876 | 0.864 |
| 883 | GG**Ala**G**Leu**GG | 0.729 | 0.775 | 0.760 | 0.783 | 0.815 | 0.825 | 0.779 |
| 884 | GG**Ala**G**Lys**GG | 0.841 | 0.866 | 0.857 | 0.843 | 0.877 | 0.865 | 0.861 |
| 885 | GG**Ala**G**Met**GG | 0.797 | 0.887 | 0.879 | 0.839 | 0.868 | 0.873 | 0.871 |
| 886 | GG**Ala**G**Phe**GG | 0.857 | 0.850 | 0.881 | 0.818 | 0.840 | 0.809 | 0.845 |
| 887 | GG**Ala**G**Pro**GG | 0.902 | 0.886 | 0.882 | 0.894 | 0.889 | 0.886 | 0.887 |
| 888 | GG**Ala**G**Ptr**GG | 0.849 | 0.784 | 0.846 | 0.779 | 0.831 | 0.887 | 0.839 |
| 889 | GG**Ala**G**S1p**GG | 0.848 | 0.792 | 0.840 | 0.817 | 0.868 | 0.841 | 0.841 |
| 890 | GG**Ala**G**Sep**GG | 0.879 | 0.897 | 0.856 | 0.896 | 0.837 | 0.852 | 0.868 |
| 891 | GG**Ala**G**Ser**GG | 0.899 | 0.872 | 0.784 | 0.862 | 0.819 | 0.804 | 0.840 |
| 892 | GG**Ala**G**T1p**GG | 0.507 | 0.771 | 0.673 | 0.616 | 0.715 | 0.745 | 0.694 |
| 893 | GG**Ala**G**Thr**GG | 0.724 | 0.755 | 0.696 | 0.783 | 0.744 | 0.784 | 0.749 |
| 894 | GG**Ala**G**Tpo**GG | 0.809 | 0.883 | 0.718 | 0.731 | 0.353 | 0.809 | 0.770 |
| 895 | GG**Ala**G**Trp**GG | 0.776 | 0.766 | 0.758 | 0.747 | 0.712 | 0.794 | 0.762 |
| 896 | GG**Ala**G**Tyr**GG | 0.697 | 0.859 | 0.841 | 0.666 | 0.723 | 0.883 | 0.782 |
| 897 | GG**Ala**G**Val**GG | 0.809 | 0.740 | 0.763 | 0.818 | 0.781 | 0.680 | 0.772 |
| 898 | GG**Ala**G**Y1p**GG | 0.765 | 0.837 | 0.816 | 0.811 | 0.787 | 0.789 | 0.800 |
| 899 | GG**Arg**G**Ala**GG | 0.858 | 0.844 | 0.829 | 0.839 | 0.823 | 0.754 | 0.834 |

This work is shared under a CC BY 4.0 license.

| | Peptide | $r_{replica}$(1,2) | $r_{replica}$(1,3) | $r_{replica}$(1,4) | $r_{replica}$(2,3) | $r_{replica}$(2,4) | $r_{replica}$(3,4) | Median |
|---|---|---|---|---|---|---|---|---|
| 900 | GG**Arg**G**Arg**GG | 0.626 | 0.644 | 0.698 | 0.726 | 0.729 | 0.701 | 0.699 |
| 901 | GG**Arg**G**Ash**GG | 0.630 | 0.574 | 0.738 | 0.802 | 0.734 | 0.658 | 0.696 |
| 902 | GG**Arg**G**Asn**GG | 0.695 | 0.764 | 0.671 | 0.833 | 0.840 | 0.799 | 0.782 |
| 903 | GG**Arg**G**Asp**GG | 0.742 | 0.750 | 0.705 | 0.839 | 0.763 | 0.699 | 0.746 |
| 904 | GG**Arg**G**Cys**GG | 0.783 | 0.813 | 0.798 | 0.823 | 0.731 | 0.789 | 0.794 |
| 905 | GG**Arg**G**Glh**GG | 0.567 | 0.665 | 0.514 | 0.776 | 0.810 | 0.725 | 0.695 |
| 906 | GG**Arg**G**Gln**GG | 0.826 | 0.820 | 0.810 | 0.867 | 0.731 | 0.747 | 0.815 |
| 907 | GG**Arg**G**Glu**GG | 0.654 | 0.760 | 0.695 | 0.782 | 0.687 | 0.790 | 0.728 |
| 908 | GG**Arg**G**Gly**GG | 0.796 | 0.791 | 0.755 | 0.767 | 0.797 | 0.766 | 0.779 |
| 909 | GG**Arg**G**Hip**GG | 0.744 | 0.826 | 0.799 | 0.750 | 0.851 | 0.837 | 0.813 |
| 910 | GG**Arg**G**His**GG | 0.803 | 0.805 | 0.814 | 0.783 | 0.775 | 0.831 | 0.804 |
| 911 | GG**Arg**G**Ile**GG | 0.738 | 0.606 | 0.789 | 0.813 | 0.787 | 0.679 | 0.763 |
| 912 | GG**Arg**G**Leu**GG | 0.820 | 0.811 | 0.828 | 0.797 | 0.811 | 0.823 | 0.816 |
| 913 | GG**Arg**G**Lys**GG | 0.746 | 0.802 | 0.771 | 0.772 | 0.792 | 0.850 | 0.782 |
| 914 | GG**Arg**G**Met**GG | 0.743 | 0.813 | 0.814 | 0.677 | 0.806 | 0.770 | 0.788 |
| 915 | GG**Arg**G**Phe**GG | 0.697 | 0.817 | 0.664 | 0.806 | 0.729 | 0.724 | 0.727 |
| 916 | GG**Arg**G**Pro**GG | 0.809 | 0.842 | 0.711 | 0.739 | 0.715 | 0.759 | 0.749 |
| 917 | GG**Arg**G**Ptr**GG | 0.633 | 0.613 | 0.729 | 0.672 | 0.743 | 0.702 | 0.687 |
| 918 | GG**Arg**G**S1p**GG | 0.809 | 0.716 | 0.785 | 0.705 | 0.781 | 0.725 | 0.753 |
| 919 | GG**Arg**G**Sep**GG | 0.767 | 0.835 | 0.718 | 0.760 | 0.629 | 0.604 | 0.739 |
| 920 | GG**Arg**G**Ser**GG | 0.817 | 0.814 | 0.768 | 0.784 | 0.750 | 0.739 | 0.776 |
| 921 | GG**Arg**G**T1p**GG | 0.646 | 0.642 | 0.739 | 0.870 | 0.582 | 0.551 | 0.644 |
| 922 | GG**Arg**G**Thr**GG | 0.732 | 0.672 | 0.712 | 0.797 | 0.705 | 0.730 | 0.721 |
| 923 | GG**Arg**G**Tpo**GG | 0.737 | 0.686 | 0.655 | 0.818 | 0.740 | 0.692 | 0.714 |
| 924 | GG**Arg**G**Trp**GG | 0.777 | 0.551 | 0.763 | 0.609 | 0.773 | 0.549 | 0.686 |
| 925 | GG**Arg**G**Tyr**GG | 0.798 | 0.770 | 0.745 | 0.767 | 0.697 | 0.676 | 0.756 |
| 926 | GG**Arg**G**Val**GG | 0.715 | 0.731 | 0.678 | 0.773 | 0.712 | 0.726 | 0.720 |
| 927 | GG**Arg**G**Y1p**GG | 0.629 | 0.589 | 0.571 | 0.600 | 0.569 | 0.530 | 0.580 |
| 928 | GG**Ash**G**Ala**GG | 0.861 | 0.894 | 0.864 | 0.849 | 0.889 | 0.846 | 0.862 |
| 929 | GG**Ash**G**Arg**GG | 0.677 | 0.770 | 0.730 | 0.718 | 0.721 | 0.780 | 0.726 |
| 930 | GG**Ash**G**Ash**GG | 0.801 | 0.693 | 0.730 | 0.792 | 0.650 | 0.624 | 0.711 |
| 931 | GG**Ash**G**Asn**GG | 0.839 | 0.762 | 0.728 | 0.744 | 0.726 | 0.744 | 0.744 |
| 932 | GG**Ash**G**Asp**GG | 0.795 | 0.738 | 0.821 | 0.713 | 0.741 | 0.769 | 0.755 |
| 933 | GG**Ash**G**Cys**GG | 0.863 | 0.864 | 0.865 | 0.843 | 0.836 | 0.835 | 0.853 |
| 934 | GG**Ash**G**Glh**GG | 0.775 | 0.715 | 0.833 | 0.681 | 0.822 | 0.665 | 0.745 |
| 935 | GG**Ash**G**Gln**GG | 0.802 | 0.782 | 0.845 | 0.698 | 0.745 | 0.786 | 0.784 |
| 936 | GG**Ash**G**Glu**GG | 0.731 | 0.811 | 0.728 | 0.674 | 0.797 | 0.693 | 0.729 |
| 937 | GG**Ash**G**Gly**GG | 0.865 | 0.824 | 0.859 | 0.857 | 0.895 | 0.857 | 0.858 |
| 938 | GG**Ash**G**Hip**GG | 0.841 | 0.805 | 0.713 | 0.808 | 0.759 | 0.700 | 0.782 |
| 939 | GG**Ash**G**His**GG | 0.827 | 0.745 | 0.837 | 0.768 | 0.772 | 0.770 | 0.771 |
| 940 | GG**Ash**G**Ile**GG | 0.753 | 0.804 | 0.825 | 0.752 | 0.753 | 0.791 | 0.772 |
| 941 | GG**Ash**G**Leu**GG | 0.837 | 0.829 | 0.831 | 0.786 | 0.830 | 0.852 | 0.830 |
| 942 | GG**Ash**G**Lys**GG | 0.788 | 0.749 | 0.763 | 0.721 | 0.756 | 0.804 | 0.760 |
| 943 | GG**Ash**G**Met**GG | 0.681 | 0.564 | 0.612 | 0.582 | 0.686 | 0.650 | 0.631 |
| 944 | GG**Ash**G**Phe**GG | 0.841 | 0.750 | 0.722 | 0.799 | 0.712 | 0.766 | 0.758 |
| 945 | GG**Ash**G**Pro**GG | 0.822 | 0.889 | 0.854 | 0.831 | 0.829 | 0.860 | 0.842 |
| 946 | GG**Ash**G**Ptr**GG | 0.509 | 0.860 | 0.556 | 0.702 | 0.874 | 0.731 | 0.717 |
| 947 | GG**Ash**G**S1p**GG | 0.895 | 0.752 | 0.693 | 0.788 | 0.738 | 0.773 | 0.762 |
| 948 | GG**Ash**G**Sep**GG | 0.834 | 0.792 | 0.876 | 0.827 | 0.849 | 0.811 | 0.831 |
| 949 | GG**Ash**G**Ser**GG | 0.818 | 0.828 | 0.810 | 0.784 | 0.804 | 0.755 | 0.807 |

This work is shared under a CC BY 4.0 license.

| | Peptide | $r_{replica}(1,2)$ | $r_{replica}(1,3)$ | $r_{replica}(1,4)$ | $r_{replica}(2,3)$ | $r_{replica}(2,4)$ | $r_{replica}(3,4)$ | Median |
|---|---|---|---|---|---|---|---|---|
| 950 | GG**Ash**G**T1p**GG | 0.569 | 0.725 | 0.752 | 0.745 | 0.624 | 0.778 | 0.735 |
| 951 | GG**Ash**G**Thr**GG | 0.790 | 0.798 | 0.807 | 0.710 | 0.691 | 0.798 | 0.794 |
| 952 | GG**Ash**G**Tpo**GG | 0.823 | 0.866 | 0.521 | 0.875 | 0.351 | 0.311 | 0.672 |
| 953 | GG**Ash**G**Trp**GG | 0.514 | 0.730 | 0.642 | 0.477 | 0.423 | 0.811 | 0.578 |
| 954 | GG**Ash**G**Tyr**GG | 0.912 | 0.832 | 0.858 | 0.814 | 0.860 | 0.781 | 0.845 |
| 955 | GG**Ash**G**Val**GG | 0.759 | 0.739 | 0.670 | 0.773 | 0.685 | 0.733 | 0.736 |
| 956 | GG**Ash**G**Y1p**GG | 0.750 | 0.727 | 0.761 | 0.695 | 0.681 | 0.807 | 0.739 |
| 957 | GG**Asn**G**Ala**GG | 0.841 | 0.870 | 0.887 | 0.818 | 0.857 | 0.879 | 0.863 |
| 958 | GG**Asn**G**Arg**GG | 0.832 | 0.776 | 0.804 | 0.807 | 0.822 | 0.770 | 0.805 |
| 959 | GG**Asn**G**Ash**GG | 0.872 | 0.904 | 0.888 | 0.887 | 0.839 | 0.883 | 0.885 |
| 960 | GG**Asn**G**Asn**GG | 0.864 | 0.829 | 0.858 | 0.800 | 0.826 | 0.800 | 0.827 |
| 961 | GG**Asn**G**Asp**GG | 0.749 | 0.675 | 0.741 | 0.708 | 0.773 | 0.809 | 0.745 |
| 962 | GG**Asn**G**Cys**GG | 0.881 | 0.797 | 0.858 | 0.756 | 0.844 | 0.785 | 0.820 |
| 963 | GG**Asn**G**Glh**GG | 0.815 | 0.818 | 0.838 | 0.797 | 0.825 | 0.758 | 0.817 |
| 964 | GG**Asn**G**Gln**GG | 0.645 | 0.637 | 0.631 | 0.829 | 0.777 | 0.834 | 0.711 |
| 965 | GG**Asn**G**Glu**GG | 0.589 | 0.875 | 0.874 | 0.558 | 0.446 | 0.878 | 0.732 |
| 966 | GG**Asn**G**Gly**GG | 0.890 | 0.899 | 0.897 | 0.926 | 0.887 | 0.888 | 0.894 |
| 967 | GG**Asn**G**Hip**GG | 0.718 | 0.717 | 0.709 | 0.735 | 0.674 | 0.683 | 0.713 |
| 968 | GG**Asn**G**His**GG | 0.846 | 0.882 | 0.864 | 0.859 | 0.858 | 0.896 | 0.862 |
| 969 | GG**Asn**G**Ile**GG | 0.829 | 0.780 | 0.844 | 0.852 | 0.880 | 0.863 | 0.848 |
| 970 | GG**Asn**G**Leu**GG | 0.801 | 0.800 | 0.788 | 0.796 | 0.746 | 0.847 | 0.798 |
| 971 | GG**Asn**G**Lys**GG | 0.814 | 0.843 | 0.878 | 0.751 | 0.833 | 0.801 | 0.823 |
| 972 | GG**Asn**G**Met**GG | 0.785 | 0.797 | 0.824 | 0.767 | 0.754 | 0.786 | 0.786 |
| 973 | GG**Asn**G**Phe**GG | 0.787 | 0.749 | 0.848 | 0.679 | 0.827 | 0.709 | 0.768 |
| 974 | GG**Asn**G**Pro**GG | 0.791 | 0.888 | 0.867 | 0.777 | 0.792 | 0.843 | 0.818 |
| 975 | GG**Asn**G**Ptr**GG | 0.842 | 0.841 | 0.775 | 0.843 | 0.777 | 0.860 | 0.842 |
| 976 | GG**Asn**G**S1p**GG | 0.793 | 0.846 | 0.830 | 0.811 | 0.759 | 0.881 | 0.820 |
| 977 | GG**Asn**G**Sep**GG | 0.807 | 0.587 | 0.715 | 0.740 | 0.741 | 0.716 | 0.728 |
| 978 | GG**Asn**G**Ser**GG | 0.829 | 0.836 | 0.816 | 0.878 | 0.835 | 0.816 | 0.832 |
| 979 | GG**Asn**G**T1p**GG | 0.543 | 0.886 | 0.782 | 0.394 | 0.863 | 0.701 | 0.741 |
| 980 | GG**Asn**G**Thr**GG | 0.869 | 0.813 | 0.796 | 0.834 | 0.822 | 0.835 | 0.828 |
| 981 | GG**Asn**G**Tpo**GG | 0.374 | 0.321 | 0.544 | 0.903 | 0.836 | 0.849 | 0.690 |
| 982 | GG**Asn**G**Trp**GG | 0.690 | 0.768 | 0.746 | 0.780 | 0.751 | 0.782 | 0.760 |
| 983 | GG**Asn**G**Tyr**GG | 0.796 | 0.737 | 0.784 | 0.806 | 0.817 | 0.758 | 0.790 |
| 984 | GG**Asn**G**Val**GG | 0.824 | 0.822 | 0.803 | 0.864 | 0.840 | 0.835 | 0.829 |
| 985 | GG**Asn**G**Y1p**GG | 0.778 | 0.803 | 0.831 | 0.851 | 0.862 | 0.870 | 0.841 |
| 986 | GG**Asp**G**Ala**GG | 0.875 | 0.775 | 0.831 | 0.730 | 0.823 | 0.800 | 0.812 |
| 987 | GG**Asp**G**Arg**GG | 0.383 | 0.655 | 0.581 | 0.318 | 0.446 | 0.718 | 0.514 |
| 988 | GG**Asp**G**Ash**GG | 0.667 | 0.746 | 0.628 | 0.780 | 0.701 | 0.721 | 0.711 |
| 989 | GG**Asp**G**Asn**GG | 0.716 | 0.768 | 0.772 | 0.763 | 0.746 | 0.821 | 0.766 |
| 990 | GG**Asp**G**Asp**GG | 0.682 | 0.828 | 0.675 | 0.716 | 0.727 | 0.721 | 0.718 |
| 991 | GG**Asp**G**Cys**GG | 0.532 | 0.556 | 0.765 | 0.762 | 0.566 | 0.708 | 0.637 |
| 992 | GG**Asp**G**Glh**GG | 0.662 | 0.790 | 0.673 | 0.703 | 0.797 | 0.653 | 0.688 |
| 993 | GG**Asp**G**Gln**GG | 0.771 | 0.623 | 0.802 | 0.747 | 0.780 | 0.731 | 0.759 |
| 994 | GG**Asp**G**Glu**GG | 0.702 | 0.503 | 0.670 | 0.590 | 0.690 | 0.675 | 0.673 |
| 995 | GG**Asp**G**Gly**GG | 0.660 | 0.750 | 0.664 | 0.698 | 0.786 | 0.710 | 0.704 |
| 996 | GG**Asp**G**Hip**GG | 0.689 | 0.765 | 0.827 | 0.396 | 0.546 | 0.783 | 0.727 |
| 997 | GG**Asp**G**His**GG | 0.647 | 0.674 | 0.707 | 0.770 | 0.645 | 0.667 | 0.671 |
| 998 | GG**Asp**G**Ile**GG | 0.719 | 0.664 | 0.688 | 0.783 | 0.702 | 0.661 | 0.695 |
| 999 | GG**Asp**G**Leu**GG | 0.638 | 0.490 | 0.595 | 0.723 | 0.698 | 0.662 | 0.650 |

This work is shared under a CC BY 4.0 license.

| | Peptide | $r_{replica}$(1,2) | $r_{replica}$(1,3) | $r_{replica}$(1,4) | $r_{replica}$(2,3) | $r_{replica}$(2,4) | $r_{replica}$(3,4) | Median |
|---|---|---|---|---|---|---|---|---|
| 1000 | GG**Asp**G**Lys**GG | 0.607 | 0.614 | 0.620 | 0.727 | 0.698 | 0.781 | 0.659 |
| 1001 | GG**Asp**G**Met**GG | 0.741 | 0.757 | 0.739 | 0.823 | 0.737 | 0.631 | 0.740 |
| 1002 | GG**Asp**G**Phe**GG | 0.701 | 0.730 | 0.726 | 0.710 | 0.647 | 0.810 | 0.718 |
| 1003 | GG**Asp**G**Pro**GG | 0.704 | 0.787 | 0.812 | 0.789 | 0.758 | 0.808 | 0.788 |
| 1004 | GG**Asp**G**Ptr**GG | 0.721 | 0.755 | 0.740 | 0.702 | 0.791 | 0.749 | 0.745 |
| 1005 | GG**Asp**G**S1p**GG | 0.657 | 0.550 | 0.624 | 0.741 | 0.656 | 0.659 | 0.657 |
| 1006 | GG**Asp**G**Sep**GG | 0.871 | 0.809 | 0.822 | 0.788 | 0.841 | 0.857 | 0.831 |
| 1007 | GG**Asp**G**Ser**GG | 0.842 | 0.837 | 0.830 | 0.840 | 0.833 | 0.839 | 0.838 |
| 1008 | GG**Asp**G**T1p**GG | 0.716 | 0.858 | 0.779 | 0.721 | 0.608 | 0.783 | 0.750 |
| 1009 | GG**Asp**G**Thr**GG | 0.721 | 0.339 | 0.608 | 0.464 | 0.630 | 0.635 | 0.619 |
| 1010 | GG**Asp**G**Tpo**GG | 0.858 | 0.740 | 0.864 | 0.782 | 0.825 | 0.818 | 0.822 |
| 1011 | GG**Asp**G**Trp**GG | 0.743 | 0.744 | 0.785 | 0.764 | 0.846 | 0.783 | 0.773 |
| 1012 | GG**Asp**G**Tyr**GG | 0.680 | 0.787 | 0.610 | 0.738 | 0.751 | 0.643 | 0.709 |
| 1013 | GG**Asp**G**Val**GG | 0.704 | 0.721 | 0.646 | 0.760 | 0.714 | 0.643 | 0.709 |
| 1014 | GG**Asp**G**Y1p**GG | 0.647 | 0.638 | 0.616 | 0.722 | 0.722 | 0.651 | 0.649 |
| 1015 | GG**Cys**G**Ala**GG | 0.887 | 0.863 | 0.840 | 0.880 | 0.849 | 0.883 | 0.872 |
| 1016 | GG**Cys**G**Arg**GG | 0.774 | 0.758 | 0.821 | 0.832 | 0.828 | 0.821 | 0.821 |
| 1017 | GG**Cys**G**Ash**GG | 0.837 | 0.821 | 0.796 | 0.814 | 0.841 | 0.815 | 0.818 |
| 1018 | GG**Cys**G**Asn**GG | 0.785 | 0.804 | 0.794 | 0.800 | 0.767 | 0.807 | 0.797 |
| 1019 | GG**Cys**G**Asp**GG | 0.765 | 0.708 | 0.743 | 0.815 | 0.661 | 0.713 | 0.728 |
| 1020 | GG**Cys**G**Cys**GG | 0.808 | 0.874 | 0.681 | 0.860 | 0.722 | 0.748 | 0.778 |
| 1021 | GG**Cys**G**Glh**GG | 0.740 | 0.855 | 0.839 | 0.762 | 0.771 | 0.839 | 0.805 |
| 1022 | GG**Cys**G**Gln**GG | 0.779 | 0.767 | 0.764 | 0.774 | 0.732 | 0.692 | 0.766 |
| 1023 | GG**Cys**G**Glu**GG | 0.694 | 0.747 | 0.833 | 0.769 | 0.813 | 0.767 | 0.768 |
| 1024 | GG**Cys**G**Gly**GG | 0.874 | 0.872 | 0.874 | 0.833 | 0.848 | 0.877 | 0.873 |
| 1025 | GG**Cys**G**Hip**GG | 0.754 | 0.749 | 0.685 | 0.721 | 0.751 | 0.738 | 0.744 |
| 1026 | GG**Cys**G**His**GG | 0.766 | 0.793 | 0.802 | 0.803 | 0.800 | 0.798 | 0.799 |
| 1027 | GG**Cys**G**Ile**GG | 0.558 | 0.648 | 0.664 | 0.794 | 0.766 | 0.853 | 0.715 |
| 1028 | GG**Cys**G**Leu**GG | 0.616 | 0.781 | 0.747 | 0.724 | 0.722 | 0.803 | 0.736 |
| 1029 | GG**Cys**G**Lys**GG | 0.819 | 0.680 | 0.701 | 0.747 | 0.730 | 0.749 | 0.738 |
| 1030 | GG**Cys**G**Met**GG | 0.750 | 0.783 | 0.802 | 0.824 | 0.734 | 0.787 | 0.785 |
| 1031 | GG**Cys**G**Phe**GG | 0.813 | 0.723 | 0.748 | 0.726 | 0.792 | 0.741 | 0.744 |
| 1032 | GG**Cys**G**Pro**GG | 0.849 | 0.856 | 0.825 | 0.850 | 0.838 | 0.895 | 0.849 |
| 1033 | GG**Cys**G**Ptr**GG | 0.706 | 0.667 | 0.697 | 0.766 | 0.780 | 0.791 | 0.736 |
| 1034 | GG**Cys**G**S1p**GG | 0.771 | 0.761 | 0.679 | 0.767 | 0.662 | 0.800 | 0.764 |
| 1035 | GG**Cys**G**Sep**GG | 0.809 | 0.794 | 0.695 | 0.809 | 0.820 | 0.817 | 0.809 |
| 1036 | GG**Cys**G**Ser**GG | 0.852 | 0.786 | 0.764 | 0.811 | 0.818 | 0.800 | 0.805 |
| 1037 | GG**Cys**G**T1p**GG | 0.717 | 0.744 | 0.537 | 0.628 | 0.446 | 0.712 | 0.670 |
| 1038 | GG**Cys**G**Thr**GG | 0.796 | 0.823 | 0.789 | 0.788 | 0.786 | 0.814 | 0.793 |
| 1039 | GG**Cys**G**Tpo**GG | 0.781 | 0.840 | 0.798 | 0.803 | 0.781 | 0.868 | 0.800 |
| 1040 | GG**Cys**G**Trp**GG | 0.829 | 0.705 | 0.583 | 0.676 | 0.573 | 0.680 | 0.678 |
| 1041 | GG**Cys**G**Tyr**GG | 0.782 | 0.637 | 0.857 | 0.801 | 0.786 | 0.684 | 0.784 |
| 1042 | GG**Cys**G**Val**GG | 0.784 | 0.726 | 0.834 | 0.623 | 0.770 | 0.700 | 0.748 |
| 1043 | GG**Cys**G**Y1p**GG | 0.695 | 0.805 | 0.746 | 0.747 | 0.612 | 0.704 | 0.725 |
| 1044 | GG**Glh**G**Ala**GG | 0.892 | 0.772 | 0.861 | 0.789 | 0.884 | 0.780 | 0.825 |
| 1045 | GG**Glh**G**Arg**GG | 0.769 | 0.821 | 0.797 | 0.799 | 0.809 | 0.799 | 0.799 |
| 1046 | GG**Glh**G**Ash**GG | 0.799 | 0.811 | 0.773 | 0.765 | 0.777 | 0.822 | 0.788 |
| 1047 | GG**Glh**G**Asn**GG | 0.750 | 0.800 | 0.846 | 0.569 | 0.701 | 0.830 | 0.775 |
| 1048 | GG**Glh**G**Asp**GG | 0.847 | 0.666 | 0.854 | 0.662 | 0.817 | 0.702 | 0.760 |
| 1049 | GG**Glh**G**Cys**GG | 0.718 | 0.799 | 0.832 | 0.813 | 0.772 | 0.841 | 0.806 |

This work is shared under a CC BY 4.0 license.

| | Peptide | $r_{replica}(1,2)$ | $r_{replica}(1,3)$ | $r_{replica}(1,4)$ | $r_{replica}(2,3)$ | $r_{replica}(2,4)$ | $r_{replica}(3,4)$ | Median |
|---|---|---|---|---|---|---|---|---|
| 1050 | GG**Glh**G**Glh**GG | 0.747 | 0.678 | 0.773 | 0.698 | 0.786 | 0.716 | 0.731 |
| 1051 | GG**Glh**G**Gln**GG | 0.856 | 0.798 | 0.830 | 0.827 | 0.893 | 0.821 | 0.828 |
| 1052 | GG**Glh**G**Glu**GG | 0.828 | 0.792 | 0.845 | 0.809 | 0.779 | 0.852 | 0.818 |
| 1053 | GG**Glh**G**Gly**GG | 0.919 | 0.908 | 0.589 | 0.912 | 0.579 | 0.533 | 0.748 |
| 1054 | GG**Glh**G**Hip**GG | 0.746 | 0.803 | 0.593 | 0.809 | 0.661 | 0.685 | 0.715 |
| 1055 | GG**Glh**G**His**GG | 0.838 | 0.779 | 0.844 | 0.761 | 0.809 | 0.818 | 0.814 |
| 1056 | GG**Glh**G**Ile**GG | 0.785 | 0.794 | 0.736 | 0.782 | 0.774 | 0.813 | 0.784 |
| 1057 | GG**Glh**G**Leu**GG | 0.748 | 0.829 | 0.849 | 0.740 | 0.769 | 0.865 | 0.799 |
| 1058 | GG**Glh**G**Lys**GG | 0.823 | 0.860 | 0.841 | 0.861 | 0.821 | 0.873 | 0.850 |
| 1059 | GG**Glh**G**Met**GG | 0.762 | 0.748 | 0.820 | 0.773 | 0.841 | 0.809 | 0.791 |
| 1060 | GG**Glh**G**Phe**GG | 0.750 | 0.875 | 0.818 | 0.752 | 0.716 | 0.861 | 0.785 |
| 1061 | GG**Glh**G**Pro**GG | 0.831 | 0.813 | 0.861 | 0.881 | 0.870 | 0.855 | 0.858 |
| 1062 | GG**Glh**G**Ptr**GG | 0.751 | 0.760 | 0.793 | 0.780 | 0.772 | 0.742 | 0.766 |
| 1063 | GG**Glh**G**S1p**GG | 0.821 | 0.727 | 0.807 | 0.742 | 0.776 | 0.673 | 0.759 |
| 1064 | GG**Glh**G**Sep**GG | 0.834 | 0.826 | 0.755 | 0.815 | 0.790 | 0.725 | 0.803 |
| 1065 | GG**Glh**G**Ser**GG | 0.881 | 0.872 | 0.808 | 0.879 | 0.818 | 0.792 | 0.845 |
| 1066 | GG**Glh**G**T1p**GG | 0.719 | 0.644 | 0.787 | 0.713 | 0.715 | 0.689 | 0.714 |
| 1067 | GG**Glh**G**Thr**GG | 0.773 | 0.780 | 0.762 | 0.789 | 0.683 | 0.686 | 0.768 |
| 1068 | GG**Glh**G**Tpo**GG | 0.762 | 0.616 | 0.618 | 0.437 | 0.507 | 0.723 | 0.617 |
| 1069 | GG**Glh**G**Trp**GG | 0.763 | 0.768 | 0.817 | 0.784 | 0.714 | 0.798 | 0.776 |
| 1070 | GG**Glh**G**Tyr**GG | 0.795 | 0.731 | 0.546 | 0.802 | 0.505 | 0.503 | 0.638 |
| 1071 | GG**Glh**G**Val**GG | 0.826 | 0.833 | 0.678 | 0.855 | 0.744 | 0.785 | 0.805 |
| 1072 | GG**Glh**G**Y1p**GG | 0.726 | 0.746 | 0.771 | 0.784 | 0.708 | 0.682 | 0.736 |
| 1073 | GG**Gln**G**Ala**GG | 0.879 | 0.891 | 0.787 | 0.897 | 0.829 | 0.804 | 0.854 |
| 1074 | GG**Gln**G**Arg**GG | 0.663 | 0.651 | 0.494 | 0.830 | 0.354 | 0.367 | 0.572 |
| 1075 | GG**Gln**G**Ash**GG | 0.873 | 0.838 | 0.812 | 0.867 | 0.848 | 0.882 | 0.858 |
| 1076 | GG**Gln**G**Asn**GG | 0.798 | 0.899 | 0.842 | 0.796 | 0.783 | 0.850 | 0.820 |
| 1077 | GG**Gln**G**Asp**GG | 0.756 | 0.748 | 0.802 | 0.744 | 0.796 | 0.779 | 0.767 |
| 1078 | GG**Gln**G**Cys**GG | 0.813 | 0.699 | 0.777 | 0.810 | 0.873 | 0.797 | 0.803 |
| 1079 | GG**Gln**G**Glh**GG | 0.723 | 0.824 | 0.499 | 0.807 | 0.629 | 0.502 | 0.676 |
| 1080 | GG**Gln**G**Gln**GG | 0.757 | 0.767 | 0.721 | 0.794 | 0.794 | 0.746 | 0.762 |
| 1081 | GG**Gln**G**Glu**GG | 0.725 | 0.831 | 0.828 | 0.756 | 0.779 | 0.857 | 0.804 |
| 1082 | GG**Gln**G**Gly**GG | 0.874 | 0.872 | 0.854 | 0.858 | 0.836 | 0.907 | 0.865 |
| 1083 | GG**Gln**G**Hip**GG | 0.844 | 0.833 | 0.829 | 0.804 | 0.769 | 0.806 | 0.817 |
| 1084 | GG**Gln**G**His**GG | 0.614 | 0.888 | 0.755 | 0.623 | 0.649 | 0.786 | 0.702 |
| 1085 | GG**Gln**G**Ile**GG | 0.807 | 0.836 | 0.769 | 0.830 | 0.725 | 0.812 | 0.809 |
| 1086 | GG**Gln**G**Leu**GG | 0.833 | 0.875 | 0.863 | 0.855 | 0.804 | 0.847 | 0.851 |
| 1087 | GG**Gln**G**Lys**GG | 0.697 | 0.754 | 0.795 | 0.756 | 0.645 | 0.782 | 0.755 |
| 1088 | GG**Gln**G**Met**GG | 0.845 | 0.859 | 0.779 | 0.885 | 0.838 | 0.808 | 0.841 |
| 1089 | GG**Gln**G**Phe**GG | 0.693 | 0.673 | 0.712 | 0.846 | 0.873 | 0.833 | 0.772 |
| 1090 | GG**Gln**G**Pro**GG | 0.880 | 0.861 | 0.908 | 0.885 | 0.914 | 0.894 | 0.890 |
| 1091 | GG**Gln**G**Ptr**GG | 0.794 | 0.587 | 0.687 | 0.565 | 0.548 | 0.636 | 0.612 |
| 1092 | GG**Gln**G**S1p**GG | 0.826 | 0.858 | 0.843 | 0.808 | 0.797 | 0.842 | 0.834 |
| 1093 | GG**Gln**G**Sep**GG | 0.706 | 0.725 | 0.865 | 0.912 | 0.802 | 0.832 | 0.817 |
| 1094 | GG**Gln**G**Ser**GG | 0.823 | 0.872 | 0.851 | 0.858 | 0.849 | 0.913 | 0.854 |
| 1095 | GG**Gln**G**T1p**GG | 0.716 | 0.768 | 0.709 | 0.646 | 0.502 | 0.654 | 0.681 |
| 1096 | GG**Gln**G**Thr**GG | 0.724 | 0.824 | 0.801 | 0.730 | 0.808 | 0.805 | 0.803 |
| 1097 | GG**Gln**G**Tpo**GG | 0.389 | 0.329 | 0.736 | 0.837 | 0.374 | 0.401 | 0.395 |
| 1098 | GG**Gln**G**Trp**GG | 0.820 | 0.648 | 0.783 | 0.716 | 0.863 | 0.751 | 0.767 |
| 1099 | GG**Gln**G**Tyr**GG | 0.857 | 0.790 | 0.801 | 0.813 | 0.792 | 0.751 | 0.796 |

This work is shared under a CC BY 4.0 license.

| | Peptide | $r_{replica}$(1,2) | $r_{replica}$(1,3) | $r_{replica}$(1,4) | $r_{replica}$(2,3) | $r_{replica}$(2,4) | $r_{replica}$(3,4) | Median |
|---|---|---|---|---|---|---|---|---|
| 1100 | GG**Gln**G**Val**GG | 0.761 | 0.767 | 0.816 | 0.837 | 0.828 | 0.828 | 0.822 |
| 1101 | GG**Gln**G**Y1p**GG | 0.762 | 0.713 | 0.826 | 0.766 | 0.803 | 0.675 | 0.764 |
| 1102 | GG**Glu**G**Ala**GG | 0.846 | 0.858 | 0.823 | 0.829 | 0.778 | 0.815 | 0.826 |
| 1103 | GG**Glu**G**Arg**GG | 0.548 | 0.657 | 0.691 | 0.672 | 0.629 | 0.702 | 0.664 |
| 1104 | GG**Glu**G**Ash**GG | 0.768 | 0.832 | 0.793 | 0.844 | 0.821 | 0.882 | 0.826 |
| 1105 | GG**Glu**G**Asn**GG | 0.814 | 0.807 | 0.774 | 0.778 | 0.829 | 0.778 | 0.793 |
| 1106 | GG**Glu**G**Asp**GG | 0.830 | 0.678 | 0.794 | 0.678 | 0.713 | 0.750 | 0.731 |
| 1107 | GG**Glu**G**Cys**GG | 0.750 | 0.679 | 0.673 | 0.832 | 0.807 | 0.785 | 0.767 |
| 1108 | GG**Glu**G**Glh**GG | 0.712 | 0.674 | 0.713 | 0.779 | 0.773 | 0.787 | 0.743 |
| 1109 | GG**Glu**G**Gln**GG | 0.805 | 0.806 | 0.785 | 0.829 | 0.815 | 0.804 | 0.806 |
| 1110 | GG**Glu**G**Glu**GG | 0.816 | 0.824 | 0.830 | 0.822 | 0.775 | 0.799 | 0.819 |
| 1111 | GG**Glu**G**Gly**GG | 0.859 | 0.880 | 0.881 | 0.855 | 0.867 | 0.889 | 0.874 |
| 1112 | GG**Glu**G**Hip**GG | 0.632 | 0.548 | 0.509 | 0.730 | 0.723 | 0.820 | 0.678 |
| 1113 | GG**Glu**G**His**GG | 0.772 | 0.736 | 0.760 | 0.706 | 0.782 | 0.795 | 0.766 |
| 1114 | GG**Glu**G**Ile**GG | 0.786 | 0.680 | 0.686 | 0.793 | 0.784 | 0.763 | 0.773 |
| 1115 | GG**Glu**G**Leu**GG | 0.648 | 0.498 | 0.755 | 0.464 | 0.535 | 0.537 | 0.536 |
| 1116 | GG**Glu**G**Lys**GG | 0.742 | 0.831 | 0.745 | 0.772 | 0.749 | 0.760 | 0.754 |
| 1117 | GG**Glu**G**Met**GG | 0.750 | 0.738 | 0.753 | 0.753 | 0.781 | 0.812 | 0.753 |
| 1118 | GG**Glu**G**Phe**GG | 0.768 | 0.683 | 0.503 | 0.735 | 0.544 | 0.594 | 0.638 |
| 1119 | GG**Glu**G**Pro**GG | 0.856 | 0.884 | 0.893 | 0.857 | 0.849 | 0.865 | 0.861 |
| 1120 | GG**Glu**G**Ptr**GG | 0.716 | 0.774 | 0.564 | 0.655 | 0.568 | 0.571 | 0.613 |
| 1121 | GG**Glu**G**S1p**GG | 0.749 | 0.693 | 0.731 | 0.571 | 0.696 | 0.723 | 0.710 |
| 1122 | GG**Glu**G**Sep**GG | 0.798 | 0.788 | 0.728 | 0.866 | 0.803 | 0.817 | 0.800 |
| 1123 | GG**Glu**G**Ser**GG | 0.727 | 0.713 | 0.776 | 0.762 | 0.834 | 0.770 | 0.766 |
| 1124 | GG**Glu**G**T1p**GG | 0.728 | 0.862 | 0.339 | 0.740 | 0.730 | 0.415 | 0.729 |
| 1125 | GG**Glu**G**Thr**GG | 0.781 | 0.726 | 0.774 | 0.756 | 0.803 | 0.754 | 0.765 |
| 1126 | GG**Glu**G**Tpo**GG | 0.253 | 0.646 | 0.109 | 0.824 | 0.920 | 0.743 | 0.695 |
| 1127 | GG**Glu**G**Trp**GG | 0.672 | 0.622 | 0.810 | 0.587 | 0.711 | 0.628 | 0.650 |
| 1128 | GG**Glu**G**Tyr**GG | 0.810 | 0.829 | 0.711 | 0.685 | 0.794 | 0.585 | 0.752 |
| 1129 | GG**Glu**G**Val**GG | 0.785 | 0.843 | 0.770 | 0.823 | 0.719 | 0.780 | 0.782 |
| 1130 | GG**Glu**G**Y1p**GG | 0.759 | 0.790 | 0.746 | 0.811 | 0.736 | 0.780 | 0.769 |
| 1131 | GG**Gly**G**Ala**GG | 0.883 | 0.896 | 0.907 | 0.886 | 0.890 | 0.884 | 0.888 |
| 1132 | GG**Gly**G**Arg**GG | 0.786 | 0.782 | 0.809 | 0.843 | 0.841 | 0.798 | 0.803 |
| 1133 | GG**Gly**G**Ash**GG | 0.893 | 0.886 | 0.877 | 0.905 | 0.860 | 0.857 | 0.882 |
| 1134 | GG**Gly**G**Asn**GG | 0.854 | 0.868 | 0.849 | 0.839 | 0.856 | 0.860 | 0.855 |
| 1135 | GG**Gly**G**Asp**GG | 0.874 | 0.870 | 0.851 | 0.866 | 0.834 | 0.802 | 0.859 |
| 1136 | GG**Gly**G**Cys**GG | 0.851 | 0.831 | 0.906 | 0.870 | 0.854 | 0.840 | 0.853 |
| 1137 | GG**Gly**G**Glh**GG | 0.860 | 0.880 | 0.866 | 0.894 | 0.846 | 0.857 | 0.863 |
| 1138 | GG**Gly**G**Gln**GG | 0.835 | 0.824 | 0.817 | 0.856 | 0.849 | 0.857 | 0.842 |
| 1139 | GG**Gly**G**Glu**GG | 0.930 | 0.889 | 0.931 | 0.867 | 0.915 | 0.885 | 0.902 |
| 1140 | GG**Gly**G**Gly**GG | 0.933 | 0.926 | 0.951 | 0.946 | 0.951 | 0.946 | 0.946 |
| 1141 | GG**Gly**G**Hip**GG | 0.823 | 0.849 | 0.848 | 0.835 | 0.832 | 0.828 | 0.834 |
| 1142 | GG**Gly**G**His**GG | 0.901 | 0.792 | 0.744 | 0.815 | 0.715 | 0.788 | 0.790 |
| 1143 | GG**Gly**G**Ile**GG | 0.865 | 0.872 | 0.853 | 0.899 | 0.822 | 0.856 | 0.861 |
| 1144 | GG**Gly**G**Leu**GG | 0.887 | 0.871 | 0.910 | 0.863 | 0.883 | 0.899 | 0.885 |
| 1145 | GG**Gly**G**Lys**GG | 0.846 | 0.825 | 0.833 | 0.871 | 0.823 | 0.807 | 0.829 |
| 1146 | GG**Gly**G**Met**GG | 0.904 | 0.915 | 0.919 | 0.906 | 0.903 | 0.913 | 0.909 |
| 1147 | GG**Gly**G**Phe**GG | 0.920 | 0.901 | 0.891 | 0.895 | 0.904 | 0.896 | 0.899 |
| 1148 | GG**Gly**G**Pro**GG | 0.912 | 0.899 | 0.924 | 0.921 | 0.910 | 0.911 | 0.912 |
| 1149 | GG**Gly**G**Ptr**GG | 0.871 | 0.852 | 0.874 | 0.876 | 0.868 | 0.854 | 0.870 |

This work is shared under a CC BY 4.0 license.

| | Peptide | $r_{replica}(1,2)$ | $r_{replica}(1,3)$ | $r_{replica}(1,4)$ | $r_{replica}(2,3)$ | $r_{replica}(2,4)$ | $r_{replica}(3,4)$ | Median |
|---|---|---|---|---|---|---|---|---|
| 1150 | GG**Gly**G**S1p**GG | 0.853 | 0.882 | 0.841 | 0.838 | 0.843 | 0.842 | 0.842 |
| 1151 | GG**Gly**G**Sep**GG | 0.920 | 0.895 | 0.910 | 0.915 | 0.905 | 0.881 | 0.907 |
| 1152 | GG**Gly**G**Ser**GG | 0.870 | 0.908 | 0.898 | 0.866 | 0.846 | 0.901 | 0.884 |
| 1153 | GG**Gly**G**T1p**GG | 0.891 | 0.847 | 0.704 | 0.840 | 0.693 | 0.843 | 0.841 |
| 1154 | GG**Gly**G**Thr**GG | 0.866 | 0.891 | 0.814 | 0.862 | 0.856 | 0.846 | 0.859 |
| 1155 | GG**Gly**G**Tpo**GG | 0.920 | 0.921 | 0.588 | 0.900 | 0.532 | 0.633 | 0.767 |
| 1156 | GG**Gly**G**Trp**GG | 0.821 | 0.801 | 0.838 | 0.888 | 0.809 | 0.785 | 0.815 |
| 1157 | GG**Gly**G**Tyr**GG | 0.878 | 0.902 | 0.928 | 0.932 | 0.899 | 0.913 | 0.907 |
| 1158 | GG**Gly**G**Val**GG | 0.878 | 0.866 | 0.871 | 0.864 | 0.852 | 0.821 | 0.865 |
| 1159 | GG**Gly**G**Y1p**GG | 0.750 | 0.814 | 0.824 | 0.825 | 0.841 | 0.850 | 0.824 |
| 1160 | GG**Hip**G**Ala**GG | 0.813 | 0.862 | 0.832 | 0.845 | 0.896 | 0.876 | 0.854 |
| 1161 | GG**Hip**G**Arg**GG | 0.749 | 0.788 | 0.729 | 0.831 | 0.797 | 0.826 | 0.793 |
| 1162 | GG**Hip**G**Ash**GG | 0.878 | 0.812 | 0.904 | 0.789 | 0.845 | 0.850 | 0.847 |
| 1163 | GG**Hip**G**Asn**GG | 0.660 | 0.726 | 0.673 | 0.824 | 0.900 | 0.831 | 0.775 |
| 1164 | GG**Hip**G**Asp**GG | 0.796 | 0.699 | 0.819 | 0.698 | 0.792 | 0.686 | 0.745 |
| 1165 | GG**Hip**G**Cys**GG | 0.799 | 0.813 | 0.707 | 0.819 | 0.684 | 0.723 | 0.761 |
| 1166 | GG**Hip**G**Glh**GG | 0.790 | 0.834 | 0.824 | 0.838 | 0.893 | 0.864 | 0.836 |
| 1167 | GG**Hip**G**Gln**GG | 0.803 | 0.784 | 0.828 | 0.767 | 0.811 | 0.716 | 0.793 |
| 1168 | GG**Hip**G**Glu**GG | 0.718 | 0.839 | 0.540 | 0.756 | 0.662 | 0.546 | 0.690 |
| 1169 | GG**Hip**G**Gly**GG | 0.887 | 0.860 | 0.885 | 0.893 | 0.890 | 0.901 | 0.888 |
| 1170 | GG**Hip**G**Hip**GG | 0.755 | 0.818 | 0.838 | 0.764 | 0.854 | 0.790 | 0.804 |
| 1171 | GG**Hip**G**His**GG | 0.810 | 0.766 | 0.756 | 0.816 | 0.773 | 0.797 | 0.785 |
| 1172 | GG**Hip**G**Ile**GG | 0.713 | 0.756 | 0.793 | 0.793 | 0.755 | 0.763 | 0.760 |
| 1173 | GG**Hip**G**Leu**GG | 0.739 | 0.725 | 0.716 | 0.843 | 0.804 | 0.722 | 0.732 |
| 1174 | GG**Hip**G**Lys**GG | 0.860 | 0.792 | 0.693 | 0.821 | 0.737 | 0.627 | 0.765 |
| 1175 | GG**Hip**G**Met**GG | 0.671 | 0.888 | 0.839 | 0.629 | 0.803 | 0.819 | 0.811 |
| 1176 | GG**Hip**G**Phe**GG | 0.775 | 0.677 | 0.755 | 0.802 | 0.740 | 0.802 | 0.765 |
| 1177 | GG**Hip**G**Pro**GG | 0.830 | 0.784 | 0.818 | 0.870 | 0.901 | 0.888 | 0.850 |
| 1178 | GG**Hip**G**Ptr**GG | 0.710 | 0.549 | 0.763 | 0.537 | 0.685 | 0.538 | 0.617 |
| 1179 | GG**Hip**G**S1p**GG | 0.861 | 0.695 | 0.793 | 0.700 | 0.747 | 0.544 | 0.724 |
| 1180 | GG**Hip**G**Sep**GG | 0.856 | 0.893 | 0.897 | 0.850 | 0.858 | 0.868 | 0.863 |
| 1181 | GG**Hip**G**Ser**GG | 0.859 | 0.798 | 0.805 | 0.797 | 0.854 | 0.845 | 0.825 |
| 1182 | GG**Hip**G**T1p**GG | 0.681 | 0.656 | 0.670 | 0.472 | 0.820 | 0.415 | 0.663 |
| 1183 | GG**Hip**G**Thr**GG | 0.677 | 0.841 | 0.789 | 0.754 | 0.746 | 0.816 | 0.772 |
| 1184 | GG**Hip**G**Tpo**GG | 0.565 | 0.637 | 0.806 | 0.760 | 0.745 | 0.777 | 0.753 |
| 1185 | GG**Hip**G**Trp**GG | 0.666 | 0.818 | 0.805 | 0.532 | 0.731 | 0.806 | 0.768 |
| 1186 | GG**Hip**G**Tyr**GG | 0.678 | 0.642 | 0.796 | 0.711 | 0.685 | 0.653 | 0.682 |
| 1187 | GG**Hip**G**Val**GG | 0.805 | 0.813 | 0.848 | 0.839 | 0.834 | 0.790 | 0.823 |
| 1188 | GG**Hip**G**Y1p**GG | 0.816 | 0.846 | 0.327 | 0.814 | 0.338 | 0.424 | 0.619 |
| 1189 | GG**His**G**Ala**GG | 0.846 | 0.674 | 0.846 | 0.727 | 0.791 | 0.652 | 0.759 |
| 1190 | GG**His**G**Arg**GG | 0.575 | 0.633 | 0.663 | 0.690 | 0.590 | 0.716 | 0.648 |
| 1191 | GG**His**G**Ash**GG | 0.778 | 0.718 | 0.801 | 0.758 | 0.880 | 0.782 | 0.780 |
| 1192 | GG**His**G**Asn**GG | 0.617 | 0.730 | 0.800 | 0.815 | 0.615 | 0.777 | 0.754 |
| 1193 | GG**His**G**Asp**GG | 0.816 | 0.666 | 0.514 | 0.756 | 0.363 | 0.477 | 0.590 |
| 1194 | GG**His**G**Cys**GG | 0.791 | 0.766 | 0.721 | 0.836 | 0.798 | 0.806 | 0.795 |
| 1195 | GG**His**G**Glh**GG | 0.647 | 0.809 | 0.597 | 0.627 | 0.613 | 0.732 | 0.637 |
| 1196 | GG**His**G**Gln**GG | 0.706 | 0.672 | 0.635 | 0.628 | 0.659 | 0.664 | 0.661 |
| 1197 | GG**His**G**Glu**GG | 0.831 | 0.804 | 0.735 | 0.764 | 0.722 | 0.794 | 0.779 |
| 1198 | GG**His**G**Gly**GG | 0.836 | 0.808 | 0.846 | 0.803 | 0.855 | 0.763 | 0.822 |
| 1199 | GG**His**G**Hip**GG | 0.764 | 0.599 | 0.745 | 0.501 | 0.784 | 0.526 | 0.672 |

This work is shared under a CC BY 4.0 license.

| | Peptide | $r_{replica}(1,2)$ | $r_{replica}(1,3)$ | $r_{replica}(1,4)$ | $r_{replica}(2,3)$ | $r_{replica}(2,4)$ | $r_{replica}(3,4)$ | Median |
|---|---|---|---|---|---|---|---|---|
| 1200 | GG**His**G**His**GG | 0.857 | 0.810 | 0.674 | 0.773 | 0.687 | 0.751 | 0.762 |
| 1201 | GG**His**G**Ile**GG | 0.809 | 0.687 | 0.832 | 0.792 | 0.861 | 0.775 | 0.801 |
| 1202 | GG**His**G**Leu**GG | 0.844 | 0.813 | 0.725 | 0.827 | 0.715 | 0.765 | 0.789 |
| 1203 | GG**His**G**Lys**GG | 0.747 | 0.782 | 0.727 | 0.776 | 0.793 | 0.786 | 0.779 |
| 1204 | GG**His**G**Met**GG | 0.821 | 0.753 | 0.777 | 0.787 | 0.795 | 0.800 | 0.791 |
| 1205 | GG**His**G**Phe**GG | 0.666 | 0.776 | 0.837 | 0.707 | 0.785 | 0.833 | 0.781 |
| 1206 | GG**His**G**Pro**GG | 0.831 | 0.842 | 0.790 | 0.842 | 0.818 | 0.809 | 0.825 |
| 1207 | GG**His**G**Ptr**GG | 0.747 | 0.798 | 0.806 | 0.759 | 0.682 | 0.744 | 0.753 |
| 1208 | GG**His**G**S1p**GG | 0.699 | 0.763 | 0.689 | 0.791 | 0.543 | 0.554 | 0.694 |
| 1209 | GG**His**G**Sep**GG | 0.826 | 0.730 | 0.742 | 0.644 | 0.811 | 0.718 | 0.736 |
| 1210 | GG**His**G**Ser**GG | 0.851 | 0.806 | 0.850 | 0.804 | 0.828 | 0.816 | 0.822 |
| 1211 | GG**His**G**T1p**GG | 0.676 | 0.702 | 0.808 | 0.837 | 0.430 | 0.550 | 0.689 |
| 1212 | GG**His**G**Thr**GG | 0.762 | 0.829 | 0.754 | 0.764 | 0.742 | 0.767 | 0.763 |
| 1213 | GG**His**G**Tpo**GG | 0.756 | 0.701 | 0.769 | 0.795 | 0.848 | 0.793 | 0.781 |
| 1214 | GG**His**G**Trp**GG | 0.707 | 0.826 | 0.741 | 0.715 | 0.827 | 0.721 | 0.731 |
| 1215 | GG**His**G**Tyr**GG | 0.757 | 0.831 | 0.837 | 0.742 | 0.757 | 0.840 | 0.794 |
| 1216 | GG**His**G**Val**GG | 0.859 | 0.743 | 0.813 | 0.811 | 0.822 | 0.848 | 0.818 |
| 1217 | GG**His**G**Y1p**GG | 0.631 | 0.746 | 0.791 | 0.666 | 0.668 | 0.834 | 0.707 |
| 1218 | GG**Ile**G**Ala**GG | 0.734 | 0.687 | 0.672 | 0.871 | 0.858 | 0.853 | 0.794 |
| 1219 | GG**Ile**G**Arg**GG | 0.686 | 0.685 | 0.712 | 0.803 | 0.866 | 0.771 | 0.741 |
| 1220 | GG**Ile**G**Ash**GG | 0.800 | 0.809 | 0.738 | 0.744 | 0.775 | 0.706 | 0.760 |
| 1221 | GG**Ile**G**Asn**GG | 0.813 | 0.875 | 0.812 | 0.873 | 0.785 | 0.830 | 0.821 |
| 1222 | GG**Ile**G**Asp**GG | 0.756 | 0.747 | 0.637 | 0.750 | 0.781 | 0.780 | 0.753 |
| 1223 | GG**Ile**G**Cys**GG | 0.740 | 0.838 | 0.816 | 0.701 | 0.773 | 0.774 | 0.773 |
| 1224 | GG**Ile**G**Glh**GG | 0.850 | 0.613 | 0.839 | 0.565 | 0.773 | 0.619 | 0.696 |
| 1225 | GG**Ile**G**Gln**GG | 0.741 | 0.738 | 0.810 | 0.717 | 0.739 | 0.752 | 0.740 |
| 1226 | GG**Ile**G**Glu**GG | 0.738 | 0.712 | 0.706 | 0.776 | 0.838 | 0.742 | 0.740 |
| 1227 | GG**Ile**G**Gly**GG | 0.909 | 0.846 | 0.891 | 0.857 | 0.887 | 0.846 | 0.872 |
| 1228 | GG**Ile**G**Hip**GG | 0.725 | 0.832 | 0.760 | 0.735 | 0.725 | 0.730 | 0.733 |
| 1229 | GG**Ile**G**His**GG | 0.562 | 0.555 | 0.522 | 0.583 | 0.728 | 0.618 | 0.572 |
| 1230 | GG**Ile**G**Ile**GG | 0.739 | 0.887 | 0.829 | 0.710 | 0.776 | 0.799 | 0.788 |
| 1231 | GG**Ile**G**Leu**GG | 0.792 | 0.638 | 0.756 | 0.634 | 0.722 | 0.760 | 0.739 |
| 1232 | GG**Ile**G**Lys**GG | 0.565 | 0.587 | 0.653 | 0.348 | 0.357 | 0.492 | 0.529 |
| 1233 | GG**Ile**G**Met**GG | 0.738 | 0.755 | 0.778 | 0.738 | 0.787 | 0.742 | 0.749 |
| 1234 | GG**Ile**G**Phe**GG | 0.814 | 0.847 | 0.734 | 0.832 | 0.665 | 0.697 | 0.774 |
| 1235 | GG**Ile**G**Pro**GG | 0.808 | 0.772 | 0.761 | 0.808 | 0.780 | 0.795 | 0.788 |
| 1236 | GG**Ile**G**Ptr**GG | 0.639 | 0.712 | 0.681 | 0.655 | 0.694 | 0.719 | 0.687 |
| 1237 | GG**Ile**G**S1p**GG | 0.756 | 0.809 | 0.776 | 0.671 | 0.670 | 0.785 | 0.766 |
| 1238 | GG**Ile**G**Sep**GG | 0.837 | 0.748 | 0.805 | 0.796 | 0.756 | 0.725 | 0.776 |
| 1239 | GG**Ile**G**Ser**GG | 0.848 | 0.814 | 0.723 | 0.782 | 0.729 | 0.775 | 0.779 |
| 1240 | GG**Ile**G**T1p**GG | 0.278 | 0.648 | 0.908 | 0.835 | 0.350 | 0.695 | 0.671 |
| 1241 | GG**Ile**G**Thr**GG | 0.686 | 0.779 | 0.797 | 0.722 | 0.770 | 0.793 | 0.774 |
| 1242 | GG**Ile**G**Tpo**GG | 0.916 | 0.373 | 0.747 | 0.357 | 0.772 | 0.704 | 0.726 |
| 1243 | GG**Ile**G**Trp**GG | 0.714 | 0.650 | 0.571 | 0.717 | 0.660 | 0.473 | 0.655 |
| 1244 | GG**Ile**G**Tyr**GG | 0.654 | 0.777 | 0.879 | 0.779 | 0.692 | 0.842 | 0.778 |
| 1245 | GG**Ile**G**Val**GG | 0.725 | 0.582 | 0.696 | 0.454 | 0.563 | 0.540 | 0.572 |
| 1246 | GG**Ile**G**Y1p**GG | 0.735 | 0.734 | 0.751 | 0.779 | 0.674 | 0.646 | 0.734 |
| 1247 | GG**Leu**G**Ala**GG | 0.858 | 0.856 | 0.871 | 0.810 | 0.865 | 0.852 | 0.857 |
| 1248 | GG**Leu**G**Arg**GG | 0.810 | 0.773 | 0.844 | 0.795 | 0.807 | 0.819 | 0.809 |
| 1249 | GG**Leu**G**Ash**GG | 0.835 | 0.858 | 0.869 | 0.846 | 0.816 | 0.855 | 0.850 |

This work is shared under a CC BY 4.0 license.

| | Peptide | $r_{replica}(1,2)$ | $r_{replica}(1,3)$ | $r_{replica}(1,4)$ | $r_{replica}(2,3)$ | $r_{replica}(2,4)$ | $r_{replica}(3,4)$ | Median |
|---|---|---|---|---|---|---|---|---|
| 1250 | GG**Leu**G**Asn**GG | 0.742 | 0.813 | 0.837 | 0.698 | 0.797 | 0.814 | 0.805 |
| 1251 | GG**Leu**G**Asp**GG | 0.675 | 0.719 | 0.714 | 0.678 | 0.750 | 0.767 | 0.716 |
| 1252 | GG**Leu**G**Cys**GG | 0.809 | 0.765 | 0.797 | 0.793 | 0.786 | 0.793 | 0.793 |
| 1253 | GG**Leu**G**Glh**GG | 0.749 | 0.762 | 0.827 | 0.749 | 0.794 | 0.770 | 0.766 |
| 1254 | GG**Leu**G**Gln**GG | 0.837 | 0.680 | 0.686 | 0.699 | 0.677 | 0.668 | 0.683 |
| 1255 | GG**Leu**G**Glu**GG | 0.788 | 0.790 | 0.784 | 0.784 | 0.805 | 0.729 | 0.786 |
| 1256 | GG**Leu**G**Gly**GG | 0.868 | 0.920 | 0.866 | 0.850 | 0.844 | 0.861 | 0.863 |
| 1257 | GG**Leu**G**Hip**GG | 0.771 | 0.798 | 0.755 | 0.666 | 0.767 | 0.709 | 0.761 |
| 1258 | GG**Leu**G**His**GG | 0.744 | 0.750 | 0.729 | 0.797 | 0.853 | 0.820 | 0.773 |
| 1259 | GG**Leu**G**Ile**GG | 0.303 | 0.827 | 0.685 | 0.327 | 0.633 | 0.717 | 0.659 |
| 1260 | GG**Leu**G**Leu**GG | 0.601 | 0.681 | 0.711 | 0.612 | 0.700 | 0.724 | 0.691 |
| 1261 | GG**Leu**G**Lys**GG | 0.742 | 0.737 | 0.683 | 0.846 | 0.709 | 0.723 | 0.730 |
| 1262 | GG**Leu**G**Met**GG | 0.791 | 0.787 | 0.776 | 0.790 | 0.735 | 0.647 | 0.781 |
| 1263 | GG**Leu**G**Phe**GG | 0.835 | 0.622 | 0.817 | 0.674 | 0.840 | 0.716 | 0.766 |
| 1264 | GG**Leu**G**Pro**GG | 0.824 | 0.818 | 0.811 | 0.831 | 0.803 | 0.850 | 0.821 |
| 1265 | GG**Leu**G**Ptr**GG | 0.790 | 0.672 | 0.751 | 0.770 | 0.722 | 0.644 | 0.736 |
| 1266 | GG**Leu**G**S1p**GG | 0.700 | 0.561 | 0.613 | 0.620 | 0.742 | 0.450 | 0.616 |
| 1267 | GG**Leu**G**Sep**GG | 0.598 | 0.697 | 0.781 | 0.736 | 0.668 | 0.765 | 0.717 |
| 1268 | GG**Leu**G**Ser**GG | 0.777 | 0.749 | 0.763 | 0.846 | 0.600 | 0.572 | 0.756 |
| 1269 | GG**Leu**G**T1p**GG | 0.806 | 0.719 | 0.689 | 0.821 | 0.787 | 0.899 | 0.797 |
| 1270 | GG**Leu**G**Thr**GG | 0.738 | 0.791 | 0.841 | 0.756 | 0.711 | 0.782 | 0.769 |
| 1271 | GG**Leu**G**Tpo**GG | 0.634 | 0.619 | 0.605 | 0.810 | 0.463 | 0.452 | 0.612 |
| 1272 | GG**Leu**G**Trp**GG | 0.691 | 0.822 | 0.849 | 0.758 | 0.627 | 0.812 | 0.785 |
| 1273 | GG**Leu**G**Tyr**GG | 0.826 | 0.799 | 0.822 | 0.834 | 0.820 | 0.778 | 0.821 |
| 1274 | GG**Leu**G**Val**GG | 0.788 | 0.756 | 0.850 | 0.650 | 0.813 | 0.682 | 0.772 |
| 1275 | GG**Leu**G**Y1p**GG | 0.752 | 0.791 | 0.803 | 0.792 | 0.783 | 0.754 | 0.787 |
| 1276 | GG**Lys**G**Ala**GG | 0.849 | 0.848 | 0.880 | 0.846 | 0.831 | 0.817 | 0.847 |
| 1277 | GG**Lys**G**Arg**GG | 0.841 | 0.781 | 0.826 | 0.841 | 0.853 | 0.764 | 0.834 |
| 1278 | GG**Lys**G**Ash**GG | 0.689 | 0.703 | 0.665 | 0.762 | 0.781 | 0.761 | 0.732 |
| 1279 | GG**Lys**G**Asn**GG | 0.774 | 0.850 | 0.767 | 0.811 | 0.859 | 0.801 | 0.806 |
| 1280 | GG**Lys**G**Asp**GG | 0.828 | 0.853 | 0.835 | 0.817 | 0.767 | 0.854 | 0.831 |
| 1281 | GG**Lys**G**Cys**GG | 0.727 | 0.698 | 0.780 | 0.836 | 0.851 | 0.829 | 0.804 |
| 1282 | GG**Lys**G**Glh**GG | 0.686 | 0.810 | 0.826 | 0.788 | 0.749 | 0.847 | 0.799 |
| 1283 | GG**Lys**G**Gln**GG | 0.713 | 0.792 | 0.854 | 0.819 | 0.682 | 0.731 | 0.761 |
| 1284 | GG**Lys**G**Glu**GG | 0.714 | 0.676 | 0.714 | 0.733 | 0.711 | 0.819 | 0.714 |
| 1285 | GG**Lys**G**Gly**GG | 0.870 | 0.838 | 0.883 | 0.861 | 0.869 | 0.851 | 0.865 |
| 1286 | GG**Lys**G**Hip**GG | 0.840 | 0.779 | 0.669 | 0.761 | 0.761 | 0.719 | 0.761 |
| 1287 | GG**Lys**G**His**GG | 0.841 | 0.800 | 0.756 | 0.790 | 0.754 | 0.798 | 0.794 |
| 1288 | GG**Lys**G**Ile**GG | 0.829 | 0.788 | 0.838 | 0.756 | 0.837 | 0.786 | 0.808 |
| 1289 | GG**Lys**G**Leu**GG | 0.837 | 0.794 | 0.788 | 0.776 | 0.804 | 0.831 | 0.799 |
| 1290 | GG**Lys**G**Lys**GG | 0.777 | 0.763 | 0.777 | 0.776 | 0.811 | 0.789 | 0.777 |
| 1291 | GG**Lys**G**Met**GG | 0.842 | 0.735 | 0.901 | 0.815 | 0.830 | 0.729 | 0.822 |
| 1292 | GG**Lys**G**Phe**GG | 0.854 | 0.778 | 0.826 | 0.807 | 0.807 | 0.845 | 0.816 |
| 1293 | GG**Lys**G**Pro**GG | 0.834 | 0.817 | 0.864 | 0.768 | 0.834 | 0.794 | 0.825 |
| 1294 | GG**Lys**G**Ptr**GG | 0.669 | 0.584 | 0.561 | 0.773 | 0.742 | 0.689 | 0.679 |
| 1295 | GG**Lys**G**S1p**GG | 0.741 | 0.792 | 0.776 | 0.780 | 0.821 | 0.777 | 0.778 |
| 1296 | GG**Lys**G**Sep**GG | 0.828 | 0.887 | 0.884 | 0.875 | 0.832 | 0.888 | 0.879 |
| 1297 | GG**Lys**G**Ser**GG | 0.769 | 0.806 | 0.823 | 0.752 | 0.780 | 0.847 | 0.793 |
| 1298 | GG**Lys**G**T1p**GG | 0.777 | 0.754 | 0.835 | 0.690 | 0.803 | 0.714 | 0.765 |
| 1299 | GG**Lys**G**Thr**GG | 0.741 | 0.687 | 0.683 | 0.749 | 0.698 | 0.607 | 0.693 |

This work is shared under a CC BY 4.0 license.

| | Peptide | $r_{replica}(1,2)$ | $r_{replica}(1,3)$ | $r_{replica}(1,4)$ | $r_{replica}(2,3)$ | $r_{replica}(2,4)$ | $r_{replica}(3,4)$ | Median |
|---|---|---|---|---|---|---|---|---|
| 1300 | GG**Lys**G**Tpo**GG | 0.819 | 0.770 | 0.810 | 0.627 | 0.817 | 0.730 | 0.790 |
| 1301 | GG**Lys**G**Trp**GG | 0.727 | 0.823 | 0.767 | 0.663 | 0.778 | 0.757 | 0.762 |
| 1302 | GG**Lys**G**Tyr**GG | 0.853 | 0.807 | 0.866 | 0.765 | 0.738 | 0.788 | 0.798 |
| 1303 | GG**Lys**G**Val**GG | 0.814 | 0.839 | 0.884 | 0.822 | 0.858 | 0.855 | 0.847 |
| 1304 | GG**Lys**G**Y1p**GG | 0.754 | 0.725 | 0.741 | 0.772 | 0.750 | 0.705 | 0.746 |
| 1305 | GG**Met**G**Ala**GG | 0.840 | 0.820 | 0.876 | 0.899 | 0.880 | 0.863 | 0.869 |
| 1306 | GG**Met**G**Arg**GG | 0.823 | 0.784 | 0.487 | 0.744 | 0.369 | 0.518 | 0.631 |
| 1307 | GG**Met**G**Ash**GG | 0.466 | 0.820 | 0.834 | 0.419 | 0.409 | 0.882 | 0.643 |
| 1308 | GG**Met**G**Asn**GG | 0.784 | 0.819 | 0.781 | 0.780 | 0.808 | 0.737 | 0.783 |
| 1309 | GG**Met**G**Asp**GG | 0.864 | 0.845 | 0.815 | 0.826 | 0.824 | 0.833 | 0.829 |
| 1310 | GG**Met**G**Cys**GG | 0.675 | 0.706 | 0.694 | 0.805 | 0.835 | 0.881 | 0.755 |
| 1311 | GG**Met**G**Glh**GG | 0.853 | 0.723 | 0.719 | 0.656 | 0.597 | 0.605 | 0.687 |
| 1312 | GG**Met**G**Gln**GG | 0.879 | 0.791 | 0.779 | 0.820 | 0.775 | 0.732 | 0.785 |
| 1313 | GG**Met**G**Glu**GG | 0.818 | 0.641 | 0.799 | 0.664 | 0.848 | 0.615 | 0.732 |
| 1314 | GG**Met**G**Gly**GG | 0.892 | 0.894 | 0.823 | 0.865 | 0.813 | 0.859 | 0.862 |
| 1315 | GG**Met**G**Hip**GG | 0.727 | 0.742 | 0.735 | 0.685 | 0.677 | 0.751 | 0.731 |
| 1316 | GG**Met**G**His**GG | 0.760 | 0.752 | 0.645 | 0.793 | 0.571 | 0.561 | 0.698 |
| 1317 | GG**Met**G**Ile**GG | 0.859 | 0.820 | 0.746 | 0.802 | 0.799 | 0.675 | 0.800 |
| 1318 | GG**Met**G**Leu**GG | 0.781 | 0.675 | 0.804 | 0.632 | 0.757 | 0.699 | 0.728 |
| 1319 | GG**Met**G**Lys**GG | 0.809 | 0.883 | 0.809 | 0.786 | 0.816 | 0.788 | 0.809 |
| 1320 | GG**Met**G**Met**GG | 0.703 | 0.777 | 0.791 | 0.710 | 0.736 | 0.781 | 0.756 |
| 1321 | GG**Met**G**Phe**GG | 0.385 | 0.798 | 0.746 | 0.419 | 0.618 | 0.812 | 0.682 |
| 1322 | GG**Met**G**Pro**GG | 0.892 | 0.866 | 0.849 | 0.871 | 0.858 | 0.838 | 0.862 |
| 1323 | GG**Met**G**Ptr**GG | 0.813 | 0.787 | 0.855 | 0.803 | 0.847 | 0.822 | 0.818 |
| 1324 | GG**Met**G**S1p**GG | 0.779 | 0.717 | 0.785 | 0.735 | 0.824 | 0.825 | 0.782 |
| 1325 | GG**Met**G**Sep**GG | 0.775 | 0.843 | 0.884 | 0.765 | 0.740 | 0.887 | 0.809 |
| 1326 | GG**Met**G**Ser**GG | 0.820 | 0.840 | 0.821 | 0.800 | 0.773 | 0.838 | 0.821 |
| 1327 | GG**Met**G**T1p**GG | 0.801 | 0.717 | 0.690 | 0.715 | 0.795 | 0.617 | 0.716 |
| 1328 | GG**Met**G**Thr**GG | 0.788 | 0.721 | 0.756 | 0.730 | 0.781 | 0.839 | 0.769 |
| 1329 | GG**Met**G**Tpo**GG | 0.827 | 0.404 | 0.555 | 0.344 | 0.604 | 0.714 | 0.579 |
| 1330 | GG**Met**G**Trp**GG | 0.763 | 0.810 | 0.838 | 0.847 | 0.803 | 0.809 | 0.809 |
| 1331 | GG**Met**G**Tyr**GG | 0.807 | 0.781 | 0.738 | 0.843 | 0.808 | 0.839 | 0.808 |
| 1332 | GG**Met**G**Val**GG | 0.799 | 0.742 | 0.688 | 0.767 | 0.649 | 0.686 | 0.715 |
| 1333 | GG**Met**G**Y1p**GG | 0.689 | 0.874 | 0.791 | 0.687 | 0.762 | 0.791 | 0.776 |
| 1334 | GG**Phe**G**Ala**GG | 0.845 | 0.810 | 0.874 | 0.839 | 0.866 | 0.831 | 0.842 |
| 1335 | GG**Phe**G**Arg**GG | 0.671 | 0.790 | 0.731 | 0.792 | 0.811 | 0.798 | 0.791 |
| 1336 | GG**Phe**G**Ash**GG | 0.758 | 0.849 | 0.822 | 0.746 | 0.816 | 0.787 | 0.802 |
| 1337 | GG**Phe**G**Asn**GG | 0.824 | 0.789 | 0.824 | 0.755 | 0.799 | 0.822 | 0.811 |
| 1338 | GG**Phe**G**Asp**GG | 0.815 | 0.826 | 0.851 | 0.694 | 0.791 | 0.818 | 0.816 |
| 1339 | GG**Phe**G**Cys**GG | 0.797 | 0.755 | 0.811 | 0.821 | 0.812 | 0.802 | 0.806 |

This work is shared under a CC BY 4.0 license.

| | | | | | | | | |
|---|---|---|---|---|---|---|---|---|
| 1340 | GG**Phe**G**Glh**GG | 0.795 | 0.585 | 0.864 | 0.703 | 0.767 | 0.576 | 0.735 |
| 1341 | GG**Phe**G**Gln**GG | 0.873 | 0.804 | 0.839 | 0.805 | 0.881 | 0.782 | 0.822 |
| 1342 | GG**Phe**G**Glu**GG | 0.711 | 0.725 | 0.635 | 0.682 | 0.843 | 0.654 | 0.697 |
| 1343 | GG**Phe**G**Gly**GG | 0.912 | 0.858 | 0.862 | 0.860 | 0.864 | 0.846 | 0.861 |
| 1344 | GG**Phe**G**Hip**GG | 0.852 | 0.599 | 0.757 | 0.588 | 0.790 | 0.718 | 0.737 |
| 1345 | GG**Phe**G**His**GG | 0.775 | 0.785 | 0.784 | 0.831 | 0.830 | 0.743 | 0.784 |
| 1346 | GG**Phe**G**Ile**GG | 0.750 | 0.673 | 0.775 | 0.752 | 0.802 | 0.790 | 0.763 |
| 1347 | GG**Phe**G**Leu**GG | 0.631 | 0.819 | 0.745 | 0.803 | 0.814 | 0.858 | 0.808 |
| 1348 | GG**Phe**G**Lys**GG | 0.677 | 0.748 | 0.868 | 0.770 | 0.775 | 0.782 | 0.773 |
| 1349 | GG**Phe**G**Met**GG | 0.794 | 0.757 | 0.721 | 0.783 | 0.780 | 0.794 | 0.781 |

This work is shared under a CC BY 4.0 license.

| **Peptide** | $r_{replica}$**(1,2)** | $r_{replica}$**(1,3)** | $r_{replica}$**(1,4)** | $r_{replica}$**(2,3)** | $r_{replica}$**(2,4)** | $r_{replica}$**(3,4)** | **Median** | |
|---|---|---|---|---|---|---|---|---|
| 1350 | GG**Phe**G**Phe**GG | 0.750 | 0.774 | 0.831 | 0.800 | 0.810 | 0.833 | 0.805 |
| 1351 | GG**Phe**G**Pro**GG | 0.820 | 0.833 | 0.792 | 0.861 | 0.834 | 0.864 | 0.833 |
| 1352 | GG**Phe**G**Ptr**GG | 0.845 | 0.843 | 0.767 | 0.798 | 0.701 | 0.779 | 0.788 |
| 1353 | GG**Phe**G**S1p**GG | 0.736 | 0.803 | 0.835 | 0.760 | 0.751 | 0.829 | 0.781 |
| 1354 | GG**Phe**G**Sep**GG | 0.844 | 0.779 | 0.718 | 0.741 | 0.695 | 0.645 | 0.729 |
| 1355 | GG**Phe**G**Ser**GG | 0.782 | 0.836 | 0.851 | 0.722 | 0.769 | 0.849 | 0.809 |
| 1356 | GG**Phe**G**T1p**GG | 0.684 | 0.309 | 0.323 | 0.622 | 0.679 | 0.876 | 0.651 |
| 1357 | GG**Phe**G**Thr**GG | 0.799 | 0.711 | 0.816 | 0.784 | 0.809 | 0.756 | 0.791 |
| 1358 | GG**Phe**G**Tpo**GG | 0.581 | 0.889 | 0.678 | 0.563 | 0.656 | 0.673 | 0.665 |
| 1359 | GG**Phe**G**Trp**GG | 0.838 | 0.720 | 0.782 | 0.678 | 0.779 | 0.716 | 0.750 |
| 1360 | GG**Phe**G**Tyr**GG | 0.875 | 0.831 | 0.727 | 0.868 | 0.729 | 0.754 | 0.792 |
| 1361 | GG**Phe**G**Val**GG | 0.750 | 0.781 | 0.818 | 0.690 | 0.757 | 0.805 | 0.769 |
| 1362 | GG**Phe**G**Y1p**GG | 0.596 | 0.681 | 0.538 | 0.699 | 0.670 | 0.765 | 0.676 |
| 1363 | GG**Pro**G**Ala**GG | 0.823 | 0.879 | 0.904 | 0.858 | 0.869 | 0.883 | 0.874 |
| 1364 | GG**Pro**G**Arg**GG | 0.714 | 0.802 | 0.802 | 0.732 | 0.793 | 0.829 | 0.798 |
| 1365 | GG**Pro**G**Ash**GG | 0.785 | 0.664 | 0.784 | 0.829 | 0.912 | 0.751 | 0.784 |
| 1366 | GG**Pro**G**Asn**GG | 0.742 | 0.841 | 0.809 | 0.800 | 0.726 | 0.747 | 0.774 |
| 1367 | GG**Pro**G**Asp**GG | 0.881 | 0.710 | 0.870 | 0.554 | 0.847 | 0.665 | 0.778 |
| 1368 | GG**Pro**G**Cys**GG | 0.876 | 0.839 | 0.686 | 0.818 | 0.686 | 0.753 | 0.785 |
| 1369 | GG**Pro**G**Glh**GG | 0.877 | 0.867 | 0.722 | 0.869 | 0.763 | 0.672 | 0.815 |
| 1370 | GG**Pro**G**Gln**GG | 0.802 | 0.841 | 0.789 | 0.802 | 0.759 | 0.669 | 0.795 |
| 1371 | GG**Pro**G**Glu**GG | 0.823 | 0.806 | 0.795 | 0.755 | 0.888 | 0.742 | 0.800 |
| 1372 | GG**Pro**G**Gly**GG | 0.886 | 0.907 | 0.888 | 0.883 | 0.865 | 0.892 | 0.887 |
| 1373 | GG**Pro**G**Hip**GG | 0.757 | 0.824 | 0.806 | 0.785 | 0.820 | 0.822 | 0.813 |
| 1374 | GG**Pro**G**His**GG | 0.668 | 0.489 | 0.731 | 0.663 | 0.842 | 0.594 | 0.666 |
| 1375 | GG**Pro**G**Ile**GG | 0.662 | 0.702 | 0.684 | 0.824 | 0.769 | 0.727 | 0.714 |
| 1376 | GG**Pro**G**Leu**GG | 0.790 | 0.809 | 0.794 | 0.822 | 0.733 | 0.713 | 0.792 |
| 1377 | GG**Pro**G**Lys**GG | 0.808 | 0.864 | 0.868 | 0.786 | 0.770 | 0.825 | 0.816 |
| 1378 | GG**Pro**G**Met**GG | 0.802 | 0.727 | 0.823 | 0.803 | 0.732 | 0.736 | 0.769 |
| 1379 | GG**Pro**G**Phe**GG | 0.677 | 0.846 | 0.790 | 0.728 | 0.619 | 0.758 | 0.743 |
| 1380 | GG**Pro**G**Pro**GG | 0.904 | 0.869 | 0.855 | 0.887 | 0.851 | 0.861 | 0.865 |
| 1381 | GG**Pro**G**Ptr**GG | 0.711 | 0.800 | 0.829 | 0.662 | 0.775 | 0.709 | 0.743 |
| 1382 | GG**Pro**G**S1p**GG | 0.815 | 0.762 | 0.872 | 0.754 | 0.799 | 0.722 | 0.780 |
| 1383 | GG**Pro**G**Sep**GG | 0.862 | 0.779 | 0.856 | 0.748 | 0.881 | 0.835 | 0.846 |
| 1384 | GG**Pro**G**Ser**GG | 0.766 | 0.756 | 0.770 | 0.782 | 0.795 | 0.785 | 0.776 |
| 1385 | GG**Pro**G**T1p**GG | 0.860 | 0.655 | 0.624 | 0.562 | 0.467 | 0.813 | 0.640 |
| 1386 | GG**Pro**G**Thr**GG | 0.822 | 0.731 | 0.792 | 0.827 | 0.829 | 0.813 | 0.818 |
| 1387 | GG**Pro**G**Tpo**GG | 0.776 | 0.810 | 0.592 | 0.734 | 0.590 | 0.432 | 0.663 |
| 1388 | GG**Pro**G**Trp**GG | 0.673 | 0.799 | 0.825 | 0.514 | 0.701 | 0.749 | 0.725 |
| 1389 | GG**Pro**G**Tyr**GG | 0.721 | 0.812 | 0.804 | 0.913 | 0.754 | 0.832 | 0.808 |
| 1390 | GG**Pro**G**Val**GG | 0.832 | 0.814 | 0.886 | 0.849 | 0.855 | 0.790 | 0.840 |
| 1391 | GG**Pro**G**Y1p**GG | 0.698 | 0.493 | 0.672 | 0.519 | 0.714 | 0.598 | 0.635 |
| 1392 | GG**Ptr**G**Ala**GG | 0.801 | 0.866 | 0.841 | 0.825 | 0.862 | 0.871 | 0.852 |
| 1393 | GG**Ptr**G**Arg**GG | 0.387 | 0.326 | 0.864 | 0.368 | 0.319 | 0.323 | 0.347 |
| 1394 | GG**Ptr**G**Ash**GG | 0.795 | 0.795 | 0.761 | 0.730 | 0.752 | 0.808 | 0.778 |
| 1395 | GG**Ptr**G**Asn**GG | 0.836 | 0.801 | 0.827 | 0.809 | 0.823 | 0.818 | 0.820 |
| 1396 | GG**Ptr**G**Asp**GG | 0.752 | 0.678 | 0.830 | 0.647 | 0.798 | 0.674 | 0.715 |
| 1397 | GG**Ptr**G**Cys**GG | 0.627 | 0.765 | 0.575 | 0.685 | 0.731 | 0.699 | 0.692 |
| 1398 | GG**Ptr**G**Glh**GG | 0.629 | 0.706 | 0.596 | 0.715 | 0.501 | 0.525 | 0.613 |
| 1399 | GG**Ptr**G**Gln**GG | 0.640 | 0.687 | 0.368 | 0.706 | 0.544 | 0.520 | 0.592 |

This work is shared under a CC BY 4.0 license.

| | Peptide | $r_{replica}(1,2)$ | $r_{replica}(1,3)$ | $r_{replica}(1,4)$ | $r_{replica}(2,3)$ | $r_{replica}(2,4)$ | $r_{replica}(3,4)$ | Median |
|---|---|---|---|---|---|---|---|---|
| 1400 | GG**Ptr**G**Glu**GG | 0.427 | 0.667 | 0.524 | 0.564 | 0.607 | 0.791 | 0.585 |
| 1401 | GG**Ptr**G**Gly**GG | 0.758 | 0.763 | 0.710 | 0.847 | 0.747 | 0.799 | 0.761 |
| 1402 | GG**Ptr**G**Hip**GG | 0.860 | 0.852 | 0.760 | 0.824 | 0.777 | 0.830 | 0.827 |
| 1403 | GG**Ptr**G**His**GG | 0.419 | 0.540 | 0.611 | 0.475 | 0.519 | 0.604 | 0.529 |
| 1404 | GG**Ptr**G**Ile**GG | 0.739 | 0.671 | 0.662 | 0.678 | 0.601 | 0.697 | 0.674 |
| 1405 | GG**Ptr**G**Leu**GG | 0.636 | 0.704 | 0.626 | 0.720 | 0.693 | 0.605 | 0.664 |
| 1406 | GG**Ptr**G**Lys**GG | 0.765 | 0.786 | 0.707 | 0.834 | 0.669 | 0.807 | 0.775 |
| 1407 | GG**Ptr**G**Met**GG | 0.772 | 0.703 | 0.687 | 0.818 | 0.780 | 0.685 | 0.737 |
| 1408 | GG**Ptr**G**Phe**GG | 0.679 | 0.693 | 0.763 | 0.863 | 0.679 | 0.670 | 0.686 |
| 1409 | GG**Ptr**G**Pro**GG | 0.734 | 0.874 | 0.890 | 0.742 | 0.811 | 0.885 | 0.843 |
| 1410 | GG**Ptr**G**Ptr**GG | 0.731 | 0.553 | 0.684 | 0.478 | 0.789 | 0.513 | 0.619 |
| 1411 | GG**Ptr**G**S1p**GG | 0.868 | 0.688 | 0.807 | 0.746 | 0.793 | 0.578 | 0.770 |
| 1412 | GG**Ptr**G**Sep**GG | 0.823 | 0.792 | 0.837 | 0.813 | 0.705 | 0.708 | 0.802 |
| 1413 | GG**Ptr**G**Ser**GG | 0.751 | 0.511 | 0.591 | 0.588 | 0.581 | 0.599 | 0.590 |
| 1414 | GG**Ptr**G**T1p**GG | 0.714 | 0.616 | 0.741 | 0.530 | 0.865 | 0.555 | 0.665 |
| 1415 | GG**Ptr**G**Thr**GG | 0.657 | 0.481 | 0.812 | 0.688 | 0.735 | 0.546 | 0.672 |
| 1416 | GG**Ptr**G**Tpo**GG | 0.738 | 0.460 | 0.456 | 0.628 | 0.688 | 0.639 | 0.634 |
| 1417 | GG**Ptr**G**Trp**GG | 0.826 | 0.560 | 0.563 | 0.580 | 0.558 | 0.627 | 0.572 |
| 1418 | GG**Ptr**G**Tyr**GG | 0.802 | 0.794 | 0.672 | 0.754 | 0.686 | 0.676 | 0.720 |
| 1419 | GG**Ptr**G**Val**GG | 0.561 | 0.693 | 0.644 | 0.444 | 0.730 | 0.511 | 0.603 |
| 1420 | GG**Ptr**G**Y1p**GG | 0.464 | 0.422 | 0.594 | 0.443 | 0.535 | 0.548 | 0.499 |
| 1421 | GG**S1p**G**Ala**GG | 0.821 | 0.786 | 0.825 | 0.808 | 0.806 | 0.866 | 0.815 |
| 1422 | GG**S1p**G**Arg**GG | 0.557 | 0.434 | 0.336 | 0.664 | 0.619 | 0.721 | 0.588 |
| 1423 | GG**S1p**G**Ash**GG | 0.793 | 0.702 | 0.707 | 0.734 | 0.746 | 0.705 | 0.720 |
| 1424 | GG**S1p**G**Asn**GG | 0.825 | 0.812 | 0.843 | 0.816 | 0.787 | 0.797 | 0.814 |
| 1425 | GG**S1p**G**Asp**GG | 0.845 | 0.779 | 0.797 | 0.792 | 0.750 | 0.728 | 0.785 |
| 1426 | GG**S1p**G**Cys**GG | 0.798 | 0.826 | 0.770 | 0.832 | 0.822 | 0.840 | 0.824 |
| 1427 | GG**S1p**G**Glh**GG | 0.775 | 0.721 | 0.776 | 0.783 | 0.839 | 0.823 | 0.780 |
| 1428 | GG**S1p**G**Gln**GG | 0.674 | 0.705 | 0.655 | 0.715 | 0.662 | 0.649 | 0.668 |
| 1429 | GG**S1p**G**Glu**GG | 0.580 | 0.766 | 0.582 | 0.838 | 0.701 | 0.734 | 0.717 |
| 1430 | GG**S1p**G**Gly**GG | 0.808 | 0.783 | 0.777 | 0.827 | 0.821 | 0.836 | 0.814 |
| 1431 | GG**S1p**G**Hip**GG | 0.723 | 0.805 | 0.395 | 0.791 | 0.541 | 0.512 | 0.632 |
| 1432 | GG**S1p**G**His**GG | 0.713 | 0.655 | 0.741 | 0.701 | 0.677 | 0.613 | 0.689 |
| 1433 | GG**S1p**G**Ile**GG | 0.675 | 0.543 | 0.790 | 0.566 | 0.606 | 0.629 | 0.617 |
| 1434 | GG**S1p**G**Leu**GG | 0.823 | 0.811 | 0.848 | 0.812 | 0.763 | 0.736 | 0.812 |
| 1435 | GG**S1p**G**Lys**GG | 0.737 | 0.762 | 0.775 | 0.807 | 0.720 | 0.739 | 0.751 |
| 1436 | GG**S1p**G**Met**GG | 0.799 | 0.814 | 0.756 | 0.744 | 0.826 | 0.755 | 0.777 |
| 1437 | GG**S1p**G**Phe**GG | 0.769 | 0.786 | 0.761 | 0.711 | 0.792 | 0.766 | 0.767 |
| 1438 | GG**S1p**G**Pro**GG | 0.896 | 0.881 | 0.912 | 0.885 | 0.849 | 0.853 | 0.883 |
| 1439 | GG**S1p**G**Ptr**GG | 0.727 | 0.796 | 0.800 | 0.670 | 0.733 | 0.801 | 0.765 |
| 1440 | GG**S1p**G**S1p**GG | 0.789 | 0.819 | 0.812 | 0.721 | 0.713 | 0.761 | 0.775 |
| 1441 | GG**S1p**G**Sep**GG | 0.830 | 0.885 | 0.856 | 0.828 | 0.811 | 0.832 | 0.831 |
| 1442 | GG**S1p**G**Ser**GG | 0.688 | 0.760 | 0.656 | 0.666 | 0.685 | 0.633 | 0.676 |
| 1443 | GG**S1p**G**T1p**GG | 0.639 | 0.599 | 0.790 | 0.663 | 0.680 | 0.489 | 0.651 |
| 1444 | GG**S1p**G**Thr**GG | 0.814 | 0.732 | 0.810 | 0.717 | 0.792 | 0.764 | 0.778 |
| 1445 | GG**S1p**G**Tpo**GG | 0.547 | 0.398 | 0.905 | 0.940 | 0.338 | 0.161 | 0.473 |
| 1446 | GG**S1p**G**Trp**GG | 0.783 | 0.772 | 0.766 | 0.702 | 0.675 | 0.766 | 0.766 |
| 1447 | GG**S1p**G**Tyr**GG | 0.781 | 0.774 | 0.643 | 0.802 | 0.738 | 0.699 | 0.756 |
| 1448 | GG**S1p**G**Val**GG | 0.601 | 0.525 | 0.602 | 0.338 | 0.567 | 0.711 | 0.584 |
| 1449 | GG**S1p**G**Y1p**GG | 0.559 | 0.671 | 0.739 | 0.460 | 0.542 | 0.737 | 0.615 |

This work is shared under a CC BY 4.0 license.

| | Peptide | $r_{replica}$(1,2) | $r_{replica}$(1,3) | $r_{replica}$(1,4) | $r_{replica}$(2,3) | $r_{replica}$(2,4) | $r_{replica}$(3,4) | Median |
|---|---|---|---|---|---|---|---|---|
| 1450 | GG**Sep**G**Ala**GG | 0.843 | 0.808 | 0.839 | 0.844 | 0.857 | 0.826 | 0.841 |
| 1451 | GG**Sep**G**Arg**GG | 0.740 | 0.798 | 0.527 | 0.868 | 0.705 | 0.745 | 0.743 |
| 1452 | GG**Sep**G**Ash**GG | 0.825 | 0.718 | 0.647 | 0.790 | 0.704 | 0.782 | 0.750 |
| 1453 | GG**Sep**G**Asn**GG | 0.817 | 0.753 | 0.704 | 0.846 | 0.809 | 0.765 | 0.787 |
| 1454 | GG**Sep**G**Asp**GG | 0.856 | 0.831 | 0.816 | 0.829 | 0.853 | 0.832 | 0.831 |
| 1455 | GG**Sep**G**Cys**GG | 0.658 | 0.694 | 0.723 | 0.579 | 0.617 | 0.829 | 0.676 |
| 1456 | GG**Sep**G**Glh**GG | 0.722 | 0.820 | 0.795 | 0.829 | 0.813 | 0.814 | 0.814 |
| 1457 | GG**Sep**G**Gln**GG | 0.750 | 0.735 | 0.837 | 0.806 | 0.758 | 0.725 | 0.754 |
| 1458 | GG**Sep**G**Glu**GG | 0.715 | 0.863 | 0.669 | 0.627 | 0.783 | 0.660 | 0.692 |
| 1459 | GG**Sep**G**Gly**GG | 0.856 | 0.874 | 0.885 | 0.890 | 0.882 | 0.874 | 0.878 |
| 1460 | GG**Sep**G**Hip**GG | 0.804 | 0.786 | 0.599 | 0.836 | 0.761 | 0.576 | 0.774 |
| 1461 | GG**Sep**G**His**GG | 0.815 | 0.707 | 0.608 | 0.689 | 0.556 | 0.761 | 0.698 |
| 1462 | GG**Sep**G**Ile**GG | 0.525 | 0.784 | 0.633 | 0.644 | 0.685 | 0.724 | 0.664 |
| 1463 | GG**Sep**G**Leu**GG | 0.840 | 0.825 | 0.855 | 0.843 | 0.865 | 0.780 | 0.841 |
| 1464 | GG**Sep**G**Lys**GG | 0.659 | 0.762 | 0.620 | 0.655 | 0.636 | 0.622 | 0.646 |
| 1465 | GG**Sep**G**Met**GG | 0.781 | 0.838 | 0.778 | 0.773 | 0.733 | 0.804 | 0.779 |
| 1466 | GG**Sep**G**Phe**GG | 0.665 | 0.746 | 0.762 | 0.800 | 0.742 | 0.804 | 0.754 |
| 1467 | GG**Sep**G**Pro**GG | 0.873 | 0.877 | 0.851 | 0.872 | 0.886 | 0.844 | 0.872 |
| 1468 | GG**Sep**G**Ptr**GG | 0.681 | 0.690 | 0.783 | 0.571 | 0.652 | 0.805 | 0.685 |
| 1469 | GG**Sep**G**S1p**GG | 0.697 | 0.771 | 0.640 | 0.700 | 0.769 | 0.583 | 0.698 |
| 1470 | GG**Sep**G**Sep**GG | 0.838 | 0.768 | 0.808 | 0.776 | 0.848 | 0.779 | 0.794 |
| 1471 | GG**Sep**G**Ser**GG | 0.802 | 0.691 | 0.775 | 0.738 | 0.785 | 0.737 | 0.757 |
| 1472 | GG**Sep**G**T1p**GG | 0.509 | 0.780 | 0.339 | 0.727 | 0.830 | 0.548 | 0.637 |
| 1473 | GG**Sep**G**Thr**GG | 0.647 | 0.730 | 0.660 | 0.738 | 0.655 | 0.585 | 0.658 |
| 1474 | GG**Sep**G**Tpo**GG | 0.435 | 0.324 | 0.413 | 0.866 | 0.794 | 0.800 | 0.615 |
| 1475 | GG**Sep**G**Trp**GG | 0.754 | 0.692 | 0.705 | 0.678 | 0.585 | 0.666 | 0.685 |
| 1476 | GG**Sep**G**Tyr**GG | 0.860 | 0.787 | 0.738 | 0.782 | 0.740 | 0.790 | 0.784 |
| 1477 | GG**Sep**G**Val**GG | 0.790 | 0.749 | 0.698 | 0.783 | 0.712 | 0.799 | 0.766 |
| 1478 | GG**Sep**G**Y1p**GG | 0.827 | 0.758 | 0.831 | 0.763 | 0.818 | 0.761 | 0.790 |
| 1479 | GG**Ser**G**Ala**GG | 0.830 | 0.851 | 0.829 | 0.892 | 0.906 | 0.892 | 0.871 |
| 1480 | GG**Ser**G**Arg**GG | 0.629 | 0.608 | 0.643 | 0.775 | 0.587 | 0.613 | 0.621 |
| 1481 | GG**Ser**G**Ash**GG | 0.824 | 0.818 | 0.756 | 0.805 | 0.797 | 0.825 | 0.812 |
| 1482 | GG**Ser**G**Asn**GG | 0.870 | 0.774 | 0.746 | 0.757 | 0.806 | 0.732 | 0.765 |
| 1483 | GG**Ser**G**Asp**GG | 0.866 | 0.851 | 0.872 | 0.819 | 0.867 | 0.873 | 0.866 |
| 1484 | GG**Ser**G**Cys**GG | 0.839 | 0.841 | 0.748 | 0.830 | 0.770 | 0.801 | 0.815 |
| 1485 | GG**Ser**G**Glh**GG | 0.573 | 0.897 | 0.870 | 0.521 | 0.599 | 0.853 | 0.726 |
| 1486 | GG**Ser**G**Gln**GG | 0.741 | 0.823 | 0.832 | 0.848 | 0.802 | 0.847 | 0.827 |
| 1487 | GG**Ser**G**Glu**GG | 0.822 | 0.812 | 0.828 | 0.830 | 0.863 | 0.837 | 0.829 |
| 1488 | GG**Ser**G**Gly**GG | 0.891 | 0.859 | 0.876 | 0.876 | 0.883 | 0.877 | 0.877 |
| 1489 | GG**Ser**G**Hip**GG | 0.675 | 0.741 | 0.755 | 0.657 | 0.585 | 0.781 | 0.708 |
| 1490 | GG**Ser**G**His**GG | 0.801 | 0.748 | 0.704 | 0.736 | 0.687 | 0.760 | 0.742 |
| 1491 | GG**Ser**G**Ile**GG | 0.622 | 0.817 | 0.763 | 0.582 | 0.708 | 0.771 | 0.735 |
| 1492 | GG**Ser**G**Leu**GG | 0.855 | 0.795 | 0.786 | 0.817 | 0.821 | 0.846 | 0.819 |
| 1493 | GG**Ser**G**Lys**GG | 0.728 | 0.734 | 0.650 | 0.756 | 0.766 | 0.665 | 0.731 |
| 1494 | GG**Ser**G**Met**GG | 0.809 | 0.834 | 0.788 | 0.806 | 0.770 | 0.774 | 0.797 |
| 1495 | GG**Ser**G**Phe**GG | 0.817 | 0.807 | 0.833 | 0.750 | 0.796 | 0.837 | 0.812 |
| 1496 | GG**Ser**G**Pro**GG | 0.860 | 0.842 | 0.874 | 0.812 | 0.838 | 0.841 | 0.842 |
| 1497 | GG**Ser**G**Ptr**GG | 0.834 | 0.826 | 0.828 | 0.810 | 0.832 | 0.815 | 0.827 |
| 1498 | GG**Ser**G**S1p**GG | 0.824 | 0.841 | 0.791 | 0.764 | 0.801 | 0.734 | 0.796 |
| 1499 | GG**Ser**G**Sep**GG | 0.815 | 0.807 | 0.862 | 0.775 | 0.828 | 0.848 | 0.821 |

This work is shared under a CC BY 4.0 license.

| | Peptide | $r_{replica}(1,2)$ | $r_{replica}(1,3)$ | $r_{replica}(1,4)$ | $r_{replica}(2,3)$ | $r_{replica}(2,4)$ | $r_{replica}(3,4)$ | Median |
|---|---|---|---|---|---|---|---|---|
| 1500 | GG**Ser**G**Ser**GG | 0.855 | 0.873 | 0.896 | 0.829 | 0.816 | 0.864 | 0.860 |
| 1501 | GG**Ser**G**T1p**GG | 0.637 | 0.802 | 0.689 | 0.803 | 0.751 | 0.837 | 0.777 |
| 1502 | GG**Ser**G**Thr**GG | 0.744 | 0.788 | 0.724 | 0.878 | 0.747 | 0.764 | 0.756 |
| 1503 | GG**Ser**G**Tpo**GG | 0.823 | 0.810 | 0.743 | 0.812 | 0.667 | 0.727 | 0.777 |
| 1504 | GG**Ser**G**Trp**GG | 0.733 | 0.751 | 0.795 | 0.640 | 0.716 | 0.546 | 0.724 |
| 1505 | GG**Ser**G**Tyr**GG | 0.773 | 0.536 | 0.825 | 0.612 | 0.784 | 0.640 | 0.707 |
| 1506 | GG**Ser**G**Val**GG | 0.835 | 0.794 | 0.828 | 0.767 | 0.766 | 0.730 | 0.781 |
| 1507 | GG**Ser**G**Y1p**GG | 0.694 | 0.767 | 0.780 | 0.653 | 0.763 | 0.697 | 0.730 |
| 1508 | GG**T1p**G**Ala**GG | 0.368 | 0.427 | 0.761 | 0.910 | 0.790 | 0.832 | 0.776 |
| 1509 | GG**T1p**G**Arg**GG | 0.510 | 0.493 | 0.765 | 0.697 | 0.646 | 0.547 | 0.597 |
| 1510 | GG**T1p**G**Ash**GG | 0.778 | 0.735 | 0.787 | 0.777 | 0.673 | 0.678 | 0.756 |
| 1511 | GG**T1p**G**Asn**GG | 0.316 | 0.794 | 0.761 | 0.451 | 0.460 | 0.858 | 0.611 |
| 1512 | GG**T1p**G**Asp**GG | 0.636 | 0.651 | 0.765 | 0.632 | 0.612 | 0.656 | 0.644 |
| 1513 | GG**T1p**G**Cys**GG | 0.787 | 0.558 | 0.748 | 0.393 | 0.762 | 0.684 | 0.716 |
| 1514 | GG**T1p**G**Glh**GG | 0.590 | 0.778 | 0.615 | 0.637 | 0.581 | 0.542 | 0.602 |
| 1515 | GG**T1p**G**Gln**GG | 0.675 | 0.515 | 0.667 | 0.669 | 0.594 | 0.589 | 0.630 |
| 1516 | GG**T1p**G**Glu**GG | 0.666 | 0.834 | 0.735 | 0.452 | 0.892 | 0.555 | 0.701 |
| 1517 | GG**T1p**G**Gly**GG | 0.507 | 0.748 | 0.833 | 0.777 | 0.680 | 0.834 | 0.762 |
| 1518 | GG**T1p**G**Hip**GG | 0.912 | 0.708 | 0.710 | 0.521 | 0.518 | 0.846 | 0.709 |
| 1519 | GG**T1p**G**His**GG | 0.423 | 0.834 | 0.787 | 0.591 | 0.370 | 0.811 | 0.689 |
| 1520 | GG**T1p**G**Ile**GG | 0.837 | 0.633 | 0.819 | 0.549 | 0.849 | 0.538 | 0.726 |
| 1521 | GG**T1p**G**Leu**GG | 0.542 | 0.693 | 0.441 | 0.514 | 0.513 | 0.480 | 0.513 |
| 1522 | GG**T1p**G**Lys**GG | 0.749 | 0.764 | 0.760 | 0.607 | 0.778 | 0.537 | 0.755 |
| 1523 | GG**T1p**G**Met**GG | 0.726 | 0.536 | 0.718 | 0.697 | 0.814 | 0.700 | 0.709 |
| 1524 | GG**T1p**G**Phe**GG | 0.463 | 0.694 | 0.584 | 0.770 | 0.843 | 0.860 | 0.732 |
| 1525 | GG**T1p**G**Pro**GG | 0.863 | 0.483 | 0.513 | 0.747 | 0.732 | 0.956 | 0.739 |
| 1526 | GG**T1p**G**Ptr**GG | 0.802 | 0.880 | 0.524 | 0.715 | 0.651 | 0.325 | 0.683 |
| 1527 | GG**T1p**G**S1p**GG | 0.646 | 0.498 | 0.466 | 0.475 | 0.574 | 0.594 | 0.536 |
| 1528 | GG**T1p**G**Sep**GG | 0.428 | 0.408 | 0.438 | 0.751 | 0.814 | 0.707 | 0.572 |
| 1529 | GG**T1p**G**Ser**GG | 0.534 | 0.767 | 0.459 | 0.550 | 0.790 | 0.504 | 0.542 |
| 1530 | GG**T1p**G**T1p**GG | 0.388 | 0.417 | 0.497 | 0.553 | 0.432 | 0.638 | 0.465 |
| 1531 | GG**T1p**G**Thr**GG | 0.700 | 0.857 | 0.882 | 0.542 | 0.565 | 0.873 | 0.778 |
| 1532 | GG**T1p**G**Tpo**GG | 0.746 | 0.582 | 0.406 | 0.755 | 0.382 | 0.433 | 0.508 |
| 1533 | GG**T1p**G**Trp**GG | 0.769 | 0.767 | 0.704 | 0.851 | 0.575 | 0.650 | 0.736 |
| 1534 | GG**T1p**G**Tyr**GG | 0.581 | 0.794 | 0.695 | 0.696 | 0.771 | 0.746 | 0.721 |
| 1535 | GG**T1p**G**Val**GG | 0.635 | 0.644 | 0.727 | 0.781 | 0.524 | 0.664 | 0.654 |
| 1536 | GG**T1p**G**Y1p**GG | 0.762 | 0.588 | 0.798 | 0.736 | 0.790 | 0.708 | 0.749 |
| 1537 | GG**Thr**G**Ala**GG | 0.816 | 0.870 | 0.822 | 0.873 | 0.866 | 0.868 | 0.867 |
| 1538 | GG**Thr**G**Arg**GG | 0.687 | 0.639 | 0.635 | 0.737 | 0.706 | 0.666 | 0.676 |
| 1539 | GG**Thr**G**Ash**GG | 0.710 | 0.816 | 0.752 | 0.821 | 0.869 | 0.855 | 0.818 |
| 1540 | GG**Thr**G**Asn**GG | 0.836 | 0.808 | 0.805 | 0.825 | 0.800 | 0.755 | 0.806 |
| 1541 | GG**Thr**G**Asp**GG | 0.850 | 0.820 | 0.566 | 0.827 | 0.586 | 0.531 | 0.703 |
| 1542 | GG**Thr**G**Cys**GG | 0.816 | 0.813 | 0.771 | 0.859 | 0.828 | 0.836 | 0.822 |
| 1543 | GG**Thr**G**Glh**GG | 0.847 | 0.792 | 0.687 | 0.828 | 0.622 | 0.606 | 0.739 |
| 1544 | GG**Thr**G**Gln**GG | 0.817 | 0.777 | 0.730 | 0.808 | 0.735 | 0.764 | 0.770 |
| 1545 | GG**Thr**G**Glu**GG | 0.751 | 0.782 | 0.768 | 0.787 | 0.752 | 0.758 | 0.763 |
| 1546 | GG**Thr**G**Gly**GG | 0.871 | 0.879 | 0.836 | 0.876 | 0.863 | 0.850 | 0.867 |
| 1547 | GG**Thr**G**Hip**GG | 0.612 | 0.637 | 0.542 | 0.632 | 0.579 | 0.611 | 0.611 |
| 1548 | GG**Thr**G**His**GG | 0.711 | 0.798 | 0.595 | 0.737 | 0.682 | 0.629 | 0.696 |
| 1549 | GG**Thr**G**Ile**GG | 0.799 | 0.825 | 0.857 | 0.839 | 0.761 | 0.773 | 0.812 |

This work is shared under a CC BY 4.0 license.

| | Peptide | $r_{replica}(1,2)$ | $r_{replica}(1,3)$ | $r_{replica}(1,4)$ | $r_{replica}(2,3)$ | $r_{replica}(2,4)$ | $r_{replica}(3,4)$ | Median |
|---|---|---|---|---|---|---|---|---|
| 1550 | GG**Thr**G**Leu**GG | 0.752 | 0.733 | 0.807 | 0.773 | 0.770 | 0.805 | 0.772 |
| 1551 | GG**Thr**G**Lys**GG | 0.832 | 0.839 | 0.870 | 0.777 | 0.807 | 0.830 | 0.831 |
| 1552 | GG**Thr**G**Met**GG | 0.748 | 0.797 | 0.696 | 0.801 | 0.848 | 0.715 | 0.773 |
| 1553 | GG**Thr**G**Phe**GG | 0.875 | 0.691 | 0.738 | 0.735 | 0.769 | 0.750 | 0.744 |
| 1554 | GG**Thr**G**Pro**GG | 0.835 | 0.853 | 0.848 | 0.850 | 0.831 | 0.855 | 0.849 |
| 1555 | GG**Thr**G**Ptr**GG | 0.828 | 0.784 | 0.801 | 0.825 | 0.853 | 0.831 | 0.826 |
| 1556 | GG**Thr**G**S1p**GG | 0.792 | 0.747 | 0.802 | 0.722 | 0.731 | 0.790 | 0.769 |
| 1557 | GG**Thr**G**Sep**GG | 0.791 | 0.775 | 0.749 | 0.835 | 0.783 | 0.760 | 0.779 |
| 1558 | GG**Thr**G**Ser**GG | 0.790 | 0.789 | 0.776 | 0.834 | 0.866 | 0.834 | 0.812 |
| 1559 | GG**Thr**G**T1p**GG | 0.647 | 0.722 | 0.729 | 0.782 | 0.642 | 0.704 | 0.713 |
| 1560 | GG**Thr**G**Thr**GG | 0.771 | 0.739 | 0.694 | 0.798 | 0.813 | 0.769 | 0.770 |
| 1561 | GG**Thr**G**Tpo**GG | 0.793 | 0.436 | 0.435 | 0.533 | 0.589 | 0.891 | 0.561 |
| 1562 | GG**Thr**G**Trp**GG | 0.641 | 0.700 | 0.663 | 0.604 | 0.561 | 0.738 | 0.652 |
| 1563 | GG**Thr**G**Tyr**GG | 0.819 | 0.783 | 0.740 | 0.791 | 0.706 | 0.755 | 0.769 |
| 1564 | GG**Thr**G**Val**GG | 0.759 | 0.797 | 0.769 | 0.760 | 0.744 | 0.776 | 0.765 |
| 1565 | GG**Thr**G**Y1p**GG | 0.758 | 0.803 | 0.623 | 0.715 | 0.599 | 0.686 | 0.701 |
| 1566 | GG**Tpo**G**Ala**GG | 0.865 | 0.846 | 0.619 | 0.638 | 0.786 | 0.239 | 0.712 |
| 1567 | GG**Tpo**G**Arg**GG | 0.830 | 0.883 | 0.824 | 0.873 | 0.856 | 0.857 | 0.856 |
| 1568 | GG**Tpo**G**Ash**GG | 0.302 | 0.872 | 0.863 | 0.322 | 0.325 | 0.920 | 0.594 |
| 1569 | GG**Tpo**G**Asn**GG | 0.440 | 0.190 | 0.549 | 0.847 | 0.886 | 0.811 | 0.680 |
| 1570 | GG**Tpo**G**Asp**GG | 0.446 | 0.173 | 0.879 | 0.840 | 0.597 | 0.320 | 0.521 |
| 1571 | GG**Tpo**G**Cys**GG | 0.664 | 0.832 | 0.502 | 0.748 | 0.363 | 0.530 | 0.597 |
| 1572 | GG**Tpo**G**Glh**GG | 0.810 | 0.772 | 0.821 | 0.589 | 0.758 | 0.758 | 0.765 |
| 1573 | GG**Tpo**G**Gln**GG | 0.677 | 0.731 | 0.617 | 0.760 | 0.787 | 0.679 | 0.705 |
| 1574 | GG**Tpo**G**Glu**GG | 0.925 | 0.897 | 0.232 | 0.922 | 0.344 | 0.423 | 0.660 |
| 1575 | GG**Tpo**G**Gly**GG | 0.823 | 0.767 | 0.672 | 0.677 | 0.445 | 0.809 | 0.722 |
| 1576 | GG**Tpo**G**Hip**GG | 0.797 | 0.457 | 0.792 | 0.696 | 0.886 | 0.784 | 0.788 |
| 1577 | GG**Tpo**G**His**GG | 0.424 | 0.720 | 0.693 | 0.536 | 0.442 | 0.625 | 0.581 |
| 1578 | GG**Tpo**G**Ile**GG | 0.638 | 0.766 | 0.473 | 0.815 | 0.857 | 0.774 | 0.770 |
| 1579 | GG**Tpo**G**Leu**GG | 0.486 | 0.768 | 0.651 | 0.647 | 0.394 | 0.538 | 0.593 |
| 1580 | GG**Tpo**G**Lys**GG | 0.323 | -0.038 | 0.249 | 0.842 | 0.278 | 0.212 | 0.263 |
| 1581 | GG**Tpo**G**Met**GG | 0.473 | 0.592 | 0.893 | 0.881 | 0.332 | 0.454 | 0.533 |
| 1582 | GG**Tpo**G**Phe**GG | 0.793 | 0.546 | 0.617 | 0.484 | 0.653 | 0.466 | 0.581 |
| 1583 | GG**Tpo**G**Pro**GG | 0.877 | 0.875 | 0.845 | 0.838 | 0.799 | 0.622 | 0.842 |
| 1584 | GG**Tpo**G**Ptr**GG | 0.846 | 0.781 | 0.432 | 0.738 | 0.525 | 0.752 | 0.745 |
| 1585 | GG**Tpo**G**S1p**GG | 0.738 | 0.837 | 0.640 | 0.733 | 0.177 | 0.475 | 0.686 |
| 1586 | GG**Tpo**G**Sep**GG | 0.725 | 0.813 | 0.753 | 0.822 | 0.565 | 0.596 | 0.739 |
| 1587 | GG**Tpo**G**Ser**GG | 0.771 | 0.562 | 0.768 | 0.645 | 0.767 | 0.640 | 0.706 |
| 1588 | GG**Tpo**G**T1p**GG | 0.894 | 0.362 | 0.286 | 0.199 | 0.133 | 0.850 | 0.324 |
| 1589 | GG**Tpo**G**Thr**GG | 0.838 | 0.904 | 0.310 | 0.901 | 0.543 | 0.388 | 0.691 |
| 1590 | GG**Tpo**G**Tpo**GG | 0.604 | 0.592 | 0.599 | 0.478 | 0.865 | 0.457 | 0.596 |
| 1591 | GG**Tpo**G**Trp**GG | 0.677 | 0.406 | 0.885 | 0.705 | 0.788 | 0.479 | 0.691 |
| 1592 | GG**Tpo**G**Tyr**GG | 0.821 | 0.443 | 0.636 | 0.329 | 0.531 | 0.727 | 0.584 |
| 1593 | GG**Tpo**G**Val**GG | 0.832 | 0.908 | 0.516 | 0.862 | 0.213 | 0.364 | 0.674 |
| 1594 | GG**Tpo**G**Y1p**GG | 0.521 | 0.701 | 0.323 | 0.792 | 0.827 | 0.776 | 0.738 |
| 1595 | GG**Trp**G**Ala**GG | 0.805 | 0.792 | 0.788 | 0.799 | 0.793 | 0.860 | 0.796 |
| 1596 | GG**Trp**G**Arg**GG | 0.753 | 0.719 | 0.677 | 0.725 | 0.749 | 0.714 | 0.722 |
| 1597 | GG**Trp**G**Ash**GG | 0.527 | 0.600 | 0.511 | 0.854 | 0.611 | 0.660 | 0.606 |
| 1598 | GG**Trp**G**Asn**GG | 0.896 | 0.755 | 0.814 | 0.739 | 0.783 | 0.803 | 0.793 |
| 1599 | GG**Trp**G**Asp**GG | 0.834 | 0.401 | 0.479 | 0.472 | 0.448 | 0.774 | 0.476 |

This work is shared under a CC BY 4.0 license.

| | Peptide | $r_{replica}$(1,2) | $r_{replica}$(1,3) | $r_{replica}$(1,4) | $r_{replica}$(2,3) | $r_{replica}$(2,4) | $r_{replica}$(3,4) | Median |
|---|---|---|---|---|---|---|---|---|
| 1600 | GG**Trp**G**Cys**GG | 0.844 | 0.793 | 0.821 | 0.833 | 0.833 | 0.845 | 0.833 |
| 1601 | GG**Trp**G**Glh**GG | 0.769 | 0.760 | 0.746 | 0.807 | 0.755 | 0.807 | 0.764 |
| 1602 | GG**Trp**G**Gln**GG | 0.705 | 0.585 | 0.847 | 0.736 | 0.763 | 0.649 | 0.721 |
| 1603 | GG**Trp**G**Glu**GG | 0.533 | 0.783 | 0.807 | 0.569 | 0.620 | 0.706 | 0.663 |
| 1604 | GG**Trp**G**Gly**GG | 0.778 | 0.848 | 0.824 | 0.811 | 0.902 | 0.878 | 0.836 |
| 1605 | GG**Trp**G**Hip**GG | 0.679 | 0.728 | 0.770 | 0.697 | 0.668 | 0.782 | 0.712 |
| 1606 | GG**Trp**G**His**GG | 0.809 | 0.836 | 0.641 | 0.797 | 0.592 | 0.551 | 0.719 |
| 1607 | GG**Trp**G**Ile**GG | 0.665 | 0.591 | 0.656 | 0.627 | 0.766 | 0.769 | 0.660 |
| 1608 | GG**Trp**G**Leu**GG | 0.621 | 0.640 | 0.649 | 0.667 | 0.688 | 0.818 | 0.658 |
| 1609 | GG**Trp**G**Lys**GG | 0.766 | 0.702 | 0.669 | 0.696 | 0.740 | 0.698 | 0.700 |
| 1610 | GG**Trp**G**Met**GG | 0.726 | 0.688 | 0.718 | 0.684 | 0.606 | 0.754 | 0.703 |
| 1611 | GG**Trp**G**Phe**GG | 0.804 | 0.775 | 0.827 | 0.774 | 0.895 | 0.780 | 0.792 |
| 1612 | GG**Trp**G**Pro**GG | 0.745 | 0.703 | 0.736 | 0.758 | 0.783 | 0.710 | 0.740 |
| 1613 | GG**Trp**G**Ptr**GG | 0.461 | 0.728 | 0.773 | 0.332 | 0.325 | 0.822 | 0.595 |
| 1614 | GG**Trp**G**S1p**GG | 0.692 | 0.777 | 0.659 | 0.807 | 0.705 | 0.709 | 0.707 |
| 1615 | GG**Trp**G**Sep**GG | 0.535 | 0.688 | 0.421 | 0.811 | 0.772 | 0.610 | 0.649 |
| 1616 | GG**Trp**G**Ser**GG | 0.791 | 0.797 | 0.793 | 0.770 | 0.830 | 0.798 | 0.795 |
| 1617 | GG**Trp**G**T1p**GG | 0.700 | 0.416 | 0.720 | 0.669 | 0.782 | 0.659 | 0.685 |
| 1618 | GG**Trp**G**Thr**GG | 0.570 | 0.645 | 0.776 | 0.722 | 0.745 | 0.694 | 0.708 |
| 1619 | GG**Trp**G**Tpo**GG | 0.812 | 0.423 | 0.380 | 0.468 | 0.523 | 0.870 | 0.495 |
| 1620 | GG**Trp**G**Trp**GG | 0.679 | 0.630 | 0.655 | 0.638 | 0.593 | 0.634 | 0.636 |
| 1621 | GG**Trp**G**Tyr**GG | 0.669 | 0.759 | 0.729 | 0.705 | 0.701 | 0.837 | 0.717 |
| 1622 | GG**Trp**G**Val**GG | 0.787 | 0.722 | 0.836 | 0.724 | 0.786 | 0.733 | 0.760 |
| 1623 | GG**Trp**G**Y1p**GG | 0.174 | 0.904 | 0.769 | 0.142 | 0.152 | 0.785 | 0.471 |
| 1624 | GG**Tyr**G**Ala**GG | 0.817 | 0.804 | 0.733 | 0.769 | 0.738 | 0.726 | 0.754 |
| 1625 | GG**Tyr**G**Arg**GG | 0.485 | 0.380 | 0.468 | 0.599 | 0.679 | 0.652 | 0.542 |
| 1626 | GG**Tyr**G**Ash**GG | 0.846 | 0.826 | 0.850 | 0.875 | 0.841 | 0.853 | 0.848 |
| 1627 | GG**Tyr**G**Asn**GG | 0.822 | 0.777 | 0.810 | 0.683 | 0.761 | 0.783 | 0.780 |
| 1628 | GG**Tyr**G**Asp**GG | 0.806 | 0.839 | 0.830 | 0.804 | 0.833 | 0.844 | 0.831 |
| 1629 | GG**Tyr**G**Cys**GG | 0.855 | 0.826 | 0.857 | 0.825 | 0.867 | 0.856 | 0.855 |
| 1630 | GG**Tyr**G**Glh**GG | 0.847 | 0.789 | 0.851 | 0.800 | 0.896 | 0.813 | 0.830 |
| 1631 | GG**Tyr**G**Gln**GG | 0.862 | 0.716 | 0.853 | 0.710 | 0.880 | 0.765 | 0.809 |
| 1632 | GG**Tyr**G**Glu**GG | 0.667 | 0.693 | 0.674 | 0.835 | 0.851 | 0.837 | 0.764 |
| 1633 | GG**Tyr**G**Gly**GG | 0.899 | 0.895 | 0.875 | 0.924 | 0.897 | 0.876 | 0.896 |
| 1634 | GG**Tyr**G**Hip**GG | 0.805 | 0.781 | 0.790 | 0.773 | 0.813 | 0.817 | 0.797 |
| 1635 | GG**Tyr**G**His**GG | 0.687 | 0.807 | 0.753 | 0.750 | 0.666 | 0.722 | 0.736 |
| 1636 | GG**Tyr**G**Ile**GG | 0.767 | 0.725 | 0.711 | 0.795 | 0.821 | 0.749 | 0.758 |
| 1637 | GG**Tyr**G**Leu**GG | 0.837 | 0.796 | 0.534 | 0.821 | 0.639 | 0.599 | 0.718 |
| 1638 | GG**Tyr**G**Lys**GG | 0.781 | 0.879 | 0.665 | 0.770 | 0.768 | 0.669 | 0.769 |
| 1639 | GG**Tyr**G**Met**GG | 0.828 | 0.647 | 0.803 | 0.672 | 0.743 | 0.722 | 0.732 |
| 1640 | GG**Tyr**G**Phe**GG | 0.743 | 0.815 | 0.880 | 0.782 | 0.751 | 0.788 | 0.785 |
| 1641 | GG**Tyr**G**Pro**GG | 0.769 | 0.717 | 0.721 | 0.772 | 0.789 | 0.810 | 0.771 |
| 1642 | GG**Tyr**G**Ptr**GG | 0.761 | 0.808 | 0.754 | 0.756 | 0.664 | 0.748 | 0.755 |
| 1643 | GG**Tyr**G**S1p**GG | 0.827 | 0.772 | 0.859 | 0.728 | 0.833 | 0.829 | 0.828 |
| 1644 | GG**Tyr**G**Sep**GG | 0.639 | 0.698 | 0.655 | 0.803 | 0.830 | 0.785 | 0.742 |
| 1645 | GG**Tyr**G**Ser**GG | 0.788 | 0.819 | 0.803 | 0.779 | 0.811 | 0.803 | 0.803 |
| 1646 | GG**Tyr**G**T1p**GG | 0.798 | 0.797 | 0.738 | 0.812 | 0.842 | 0.771 | 0.797 |
| 1647 | GG**Tyr**G**Thr**GG | 0.776 | 0.805 | 0.865 | 0.768 | 0.779 | 0.790 | 0.785 |
| 1648 | GG**Tyr**G**Tpo**GG | 0.627 | 0.764 | 0.613 | 0.740 | 0.382 | 0.505 | 0.620 |
| 1649 | GG**Tyr**G**Trp**GG | 0.865 | 0.892 | 0.795 | 0.809 | 0.777 | 0.845 | 0.827 |

This work is shared under a CC BY 4.0 license.

| | Peptide | $r_{replica}$(1,2) | $r_{replica}$(1,3) | $r_{replica}$(1,4) | $r_{replica}$(2,3) | $r_{replica}$(2,4) | $r_{replica}$(3,4) | Median |
|---|---|---|---|---|---|---|---|---|
| 1650 | GG**Tyr**G**Tyr**GG | 0.849 | 0.782 | 0.859 | 0.764 | 0.836 | 0.786 | 0.811 |
| 1651 | GG**Tyr**G**Val**GG | 0.686 | 0.720 | 0.716 | 0.730 | 0.685 | 0.801 | 0.718 |
| 1652 | GG**Tyr**G**Y1p**GG | 0.801 | 0.750 | 0.743 | 0.878 | 0.857 | 0.895 | 0.829 |
| 1653 | GG**Val**G**Ala**GG | 0.858 | 0.872 | 0.750 | 0.880 | 0.769 | 0.788 | 0.823 |
| 1654 | GG**Val**G**Arg**GG | 0.596 | 0.662 | 0.775 | 0.640 | 0.531 | 0.699 | 0.651 |
| 1655 | GG**Val**G**Ash**GG | 0.758 | 0.834 | 0.738 | 0.777 | 0.759 | 0.739 | 0.759 |
| 1656 | GG**Val**G**Asn**GG | 0.807 | 0.792 | 0.766 | 0.777 | 0.774 | 0.738 | 0.775 |
| 1657 | GG**Val**G**Asp**GG | 0.699 | 0.657 | 0.720 | 0.744 | 0.757 | 0.726 | 0.723 |
| 1658 | GG**Val**G**Cys**GG | 0.705 | 0.828 | 0.750 | 0.763 | 0.750 | 0.789 | 0.757 |
| 1659 | GG**Val**G**Glh**GG | 0.782 | 0.696 | 0.735 | 0.781 | 0.798 | 0.841 | 0.781 |
| 1660 | GG**Val**G**Gln**GG | 0.750 | 0.848 | 0.790 | 0.763 | 0.744 | 0.801 | 0.776 |
| 1661 | GG**Val**G**Glu**GG | 0.804 | 0.764 | 0.843 | 0.815 | 0.798 | 0.775 | 0.801 |
| 1662 | GG**Val**G**Gly**GG | 0.858 | 0.867 | 0.828 | 0.878 | 0.860 | 0.876 | 0.864 |
| 1663 | GG**Val**G**Hip**GG | 0.695 | 0.728 | 0.732 | 0.783 | 0.560 | 0.659 | 0.712 |
| 1664 | GG**Val**G**His**GG | 0.680 | 0.827 | 0.804 | 0.690 | 0.735 | 0.791 | 0.763 |
| 1665 | GG**Val**G**Ile**GG | 0.482 | 0.534 | 0.715 | 0.707 | 0.575 | 0.768 | 0.641 |
| 1666 | GG**Val**G**Leu**GG | 0.754 | 0.693 | 0.707 | 0.839 | 0.848 | 0.855 | 0.797 |
| 1667 | GG**Val**G**Lys**GG | 0.692 | 0.746 | 0.553 | 0.761 | 0.660 | 0.668 | 0.680 |
| 1668 | GG**Val**G**Met**GG | 0.688 | 0.752 | 0.717 | 0.789 | 0.778 | 0.816 | 0.765 |
| 1669 | GG**Val**G**Phe**GG | 0.813 | 0.844 | 0.796 | 0.803 | 0.814 | 0.787 | 0.808 |
| 1670 | GG**Val**G**Pro**GG | 0.837 | 0.792 | 0.857 | 0.852 | 0.812 | 0.833 | 0.835 |
| 1671 | GG**Val**G**Ptr**GG | 0.767 | 0.744 | 0.699 | 0.717 | 0.655 | 0.677 | 0.708 |
| 1672 | GG**Val**G**S1p**GG | 0.840 | 0.824 | 0.731 | 0.826 | 0.695 | 0.685 | 0.778 |
| 1673 | GG**Val**G**Sep**GG | 0.771 | 0.801 | 0.732 | 0.816 | 0.829 | 0.776 | 0.789 |
| 1674 | GG**Val**G**Ser**GG | 0.761 | 0.753 | 0.777 | 0.737 | 0.766 | 0.670 | 0.757 |
| 1675 | GG**Val**G**T1p**GG | 0.844 | 0.746 | 0.757 | 0.675 | 0.727 | 0.869 | 0.751 |
| 1676 | GG**Val**G**Thr**GG | 0.781 | 0.757 | 0.705 | 0.769 | 0.766 | 0.626 | 0.762 |
| 1677 | GG**Val**G**Tpo**GG | 0.334 | 0.830 | 0.890 | 0.479 | 0.360 | 0.881 | 0.654 |
| 1678 | GG**Val**G**Trp**GG | 0.851 | 0.755 | 0.730 | 0.611 | 0.642 | 0.824 | 0.743 |
| 1679 | GG**Val**G**Tyr**GG | 0.780 | 0.805 | 0.833 | 0.754 | 0.822 | 0.763 | 0.793 |
| 1680 | GG**Val**G**Val**GG | 0.778 | 0.741 | 0.769 | 0.765 | 0.764 | 0.812 | 0.767 |
| 1681 | GG**Val**G**Y1p**GG | 0.777 | 0.775 | 0.688 | 0.743 | 0.659 | 0.820 | 0.759 |
| 1682 | GG**Y1p**G**Ala**GG | 0.837 | 0.850 | 0.876 | 0.867 | 0.837 | 0.889 | 0.859 |
| 1683 | GG**Y1p**G**Arg**GG | 0.734 | 0.667 | 0.488 | 0.756 | 0.623 | 0.604 | 0.645 |
| 1684 | GG**Y1p**G**Ash**GG | 0.801 | 0.845 | 0.876 | 0.743 | 0.771 | 0.840 | 0.821 |
| 1685 | GG**Y1p**G**Asn**GG | 0.885 | 0.774 | 0.782 | 0.766 | 0.747 | 0.750 | 0.770 |
| 1686 | GG**Y1p**G**Asp**GG | 0.724 | 0.859 | 0.594 | 0.822 | 0.605 | 0.679 | 0.701 |
| 1687 | GG**Y1p**G**Cys**GG | 0.828 | 0.719 | 0.794 | 0.711 | 0.825 | 0.811 | 0.802 |
| 1688 | GG**Y1p**G**Glh**GG | 0.757 | 0.773 | 0.602 | 0.714 | 0.670 | 0.607 | 0.692 |
| 1689 | GG**Y1p**G**Gln**GG | 0.853 | 0.759 | 0.762 | 0.747 | 0.760 | 0.685 | 0.759 |
| 1690 | GG**Y1p**G**Glu**GG | 0.698 | 0.672 | 0.579 | 0.517 | 0.499 | 0.387 | 0.548 |
| 1691 | GG**Y1p**G**Gly**GG | 0.876 | 0.823 | 0.842 | 0.870 | 0.857 | 0.781 | 0.850 |
| 1692 | GG**Y1p**G**Hip**GG | 0.706 | 0.679 | 0.459 | 0.829 | 0.622 | 0.677 | 0.678 |
| 1693 | GG**Y1p**G**His**GG | 0.825 | 0.837 | 0.796 | 0.817 | 0.814 | 0.793 | 0.816 |
| 1694 | GG**Y1p**G**Ile**GG | 0.783 | 0.635 | 0.619 | 0.548 | 0.564 | 0.683 | 0.627 |
| 1695 | GG**Y1p**G**Leu**GG | 0.634 | 0.787 | 0.663 | 0.685 | 0.659 | 0.698 | 0.674 |
| 1696 | GG**Y1p**G**Lys**GG | 0.742 | 0.704 | 0.763 | 0.695 | 0.782 | 0.805 | 0.752 |
| 1697 | GG**Y1p**G**Met**GG | 0.769 | 0.656 | 0.718 | 0.750 | 0.671 | 0.495 | 0.694 |
| 1698 | GG**Y1p**G**Phe**GG | 0.692 | 0.715 | 0.482 | 0.773 | 0.761 | 0.615 | 0.704 |
| 1699 | GG**Y1p**G**Pro**GG | 0.901 | 0.862 | 0.893 | 0.829 | 0.886 | 0.852 | 0.874 |

This work is shared under a CC BY 4.0 license.

| | Peptide | $r_{replica}(1,2)$ | $r_{replica}(1,3)$ | $r_{replica}(1,4)$ | $r_{replica}(2,3)$ | $r_{replica}(2,4)$ | $r_{replica}(3,4)$ | Median |
|---|---|---|---|---|---|---|---|---|
| 1700 | GGY1pGPtrGG | 0.633 | 0.567 | 0.621 | 0.584 | 0.594 | 0.663 | 0.607 |
| 1701 | GGY1pGS1pGG | 0.798 | 0.710 | 0.775 | 0.636 | 0.627 | 0.735 | 0.722 |
| 1702 | GGY1pGSepGG | 0.777 | 0.745 | 0.798 | 0.740 | 0.715 | 0.723 | 0.743 |
| 1703 | GGY1pGSerGG | 0.635 | 0.840 | 0.855 | 0.639 | 0.643 | 0.853 | 0.742 |
| 1704 | GGY1pGT1pGG | 0.755 | 0.602 | 0.816 | 0.367 | 0.797 | 0.628 | 0.691 |
| 1705 | GGY1pGThrGG | 0.838 | 0.785 | 0.824 | 0.797 | 0.787 | 0.795 | 0.796 |
| 1706 | GGY1pGTpoGG | 0.614 | 0.696 | 0.605 | 0.407 | 0.314 | 0.599 | 0.602 |
| 1707 | GGY1pGTrpGG | 0.617 | 0.644 | 0.548 | 0.624 | 0.615 | 0.439 | 0.616 |
| 1708 | GGY1pGTyrGG | 0.779 | 0.799 | 0.742 | 0.854 | 0.812 | 0.834 | 0.805 |
| 1709 | GGY1pGValGG | 0.517 | 0.739 | 0.695 | 0.538 | 0.483 | 0.787 | 0.617 |
| 1710 | GGY1pGY1pGG | 0.824 | 0.670 | 0.720 | 0.702 | 0.729 | 0.713 | 0.716 |
| 1711 | GGAlaGGAlaGG | 0.896 | 0.884 | 0.870 | 0.919 | 0.908 | 0.917 | 0.902 |
| 1712 | GGAlaGGArgGG | 0.784 | 0.816 | 0.776 | 0.805 | 0.754 | 0.830 | 0.795 |
| 1713 | GGAlaGGAshGG | 0.796 | 0.853 | 0.874 | 0.864 | 0.858 | 0.906 | 0.861 |
| 1714 | GGAlaGGAsnGG | 0.827 | 0.824 | 0.887 | 0.769 | 0.875 | 0.818 | 0.825 |
| 1715 | GGAlaGGAspGG | 0.758 | 0.732 | 0.810 | 0.822 | 0.861 | 0.854 | 0.816 |
| 1716 | GGAlaGGCysGG | 0.829 | 0.859 | 0.864 | 0.864 | 0.841 | 0.896 | 0.861 |
| 1717 | GGAlaGGGlhGG | 0.821 | 0.794 | 0.842 | 0.843 | 0.876 | 0.887 | 0.842 |
| 1718 | GGAlaGGGlnGG | 0.875 | 0.865 | 0.897 | 0.865 | 0.867 | 0.849 | 0.866 |
| 1719 | GGAlaGGGluGG | 0.867 | 0.914 | 0.876 | 0.840 | 0.865 | 0.842 | 0.866 |
| 1720 | GGAlaGGGlyGG | 0.906 | 0.931 | 0.867 | 0.929 | 0.862 | 0.889 | 0.897 |
| 1721 | GGAlaGGHipGG | 0.767 | 0.820 | 0.733 | 0.731 | 0.630 | 0.759 | 0.746 |
| 1722 | GGAlaGGHisGG | 0.709 | 0.691 | 0.784 | 0.548 | 0.794 | 0.641 | 0.700 |
| 1723 | GGAlaGGIleGG | 0.890 | 0.908 | 0.828 | 0.865 | 0.812 | 0.840 | 0.852 |
| 1724 | GGAlaGGLeuGG | 0.800 | 0.827 | 0.811 | 0.839 | 0.823 | 0.836 | 0.825 |
| 1725 | GGAlaGGLysGG | 0.699 | 0.714 | 0.736 | 0.644 | 0.653 | 0.684 | 0.692 |
| 1726 | GGAlaGGMetGG | 0.787 | 0.771 | 0.776 | 0.808 | 0.814 | 0.786 | 0.786 |
| 1727 | GGAlaGGPheGG | 0.916 | 0.892 | 0.866 | 0.876 | 0.886 | 0.896 | 0.889 |
| 1728 | GGAlaGGProGG | 0.917 | 0.901 | 0.861 | 0.915 | 0.901 | 0.927 | 0.908 |
| 1729 | GGAlaGGPtrGG | 0.903 | 0.882 | 0.844 | 0.895 | 0.863 | 0.870 | 0.876 |
| 1730 | GGAlaGGS1pGG | 0.866 | 0.861 | 0.843 | 0.883 | 0.804 | 0.856 | 0.858 |
| 1731 | GGAlaGGSepGG | 0.846 | 0.784 | 0.833 | 0.730 | 0.780 | 0.801 | 0.793 |
| 1732 | GGAlaGGSerGG | 0.824 | 0.791 | 0.805 | 0.806 | 0.812 | 0.918 | 0.809 |
| 1733 | GGAlaGGT1pGG | 0.832 | 0.699 | 0.750 | 0.720 | 0.759 | 0.844 | 0.755 |
| 1734 | GGAlaGGThrGG | 0.828 | 0.830 | 0.884 | 0.869 | 0.886 | 0.870 | 0.869 |
| 1735 | GGAlaGGTpoGG | 0.140 | 0.955 | 0.773 | 0.273 | 0.665 | 0.831 | 0.719 |
| 1736 | GGAlaGGTrpGG | 0.785 | 0.703 | 0.723 | 0.802 | 0.776 | 0.744 | 0.760 |
| 1737 | GGAlaGGTyrGG | 0.894 | 0.903 | 0.896 | 0.914 | 0.882 | 0.901 | 0.898 |
| 1738 | GGAlaGGValGG | 0.764 | 0.836 | 0.827 | 0.789 | 0.770 | 0.806 | 0.797 |
| 1739 | GGAlaGGY1pGG | 0.866 | 0.872 | 0.862 | 0.893 | 0.884 | 0.872 | 0.872 |
| 1740 | GGArgGGAlaGG | 0.704 | 0.784 | 0.744 | 0.807 | 0.666 | 0.713 | 0.728 |
| 1741 | GGArgGGArgGG | 0.750 | 0.592 | 0.739 | 0.645 | 0.713 | 0.482 | 0.679 |
| 1742 | GGArgGGAshGG | 0.853 | 0.840 | 0.870 | 0.881 | 0.846 | 0.837 | 0.849 |
| 1743 | GGArgGGAsnGG | 0.814 | 0.822 | 0.771 | 0.867 | 0.800 | 0.823 | 0.818 |
| 1744 | GGArgGGAspGG | 0.748 | 0.770 | 0.805 | 0.794 | 0.809 | 0.870 | 0.799 |
| 1745 | GGArgGGCysGG | 0.769 | 0.747 | 0.781 | 0.747 | 0.782 | 0.735 | 0.758 |
| 1746 | GGArgGGGlhGG | 0.663 | 0.676 | 0.718 | 0.525 | 0.718 | 0.686 | 0.681 |
| 1747 | GGArgGGGlnGG | 0.745 | 0.653 | 0.780 | 0.768 | 0.684 | 0.601 | 0.715 |
| 1748 | GGArgGGGluGG | 0.724 | 0.619 | 0.581 | 0.596 | 0.572 | 0.756 | 0.607 |
| 1749 | GGArgGGGlyGG | 0.862 | 0.878 | 0.851 | 0.835 | 0.855 | 0.817 | 0.853 |

This work is shared under a CC BY 4.0 license.

| | Peptide | $r_{replica}(1,2)$ | $r_{replica}(1,3)$ | $r_{replica}(1,4)$ | $r_{replica}(2,3)$ | $r_{replica}(2,4)$ | $r_{replica}(3,4)$ | Median |
|---|---|---|---|---|---|---|---|---|
| 1750 | GG**Arg**GG**Hip**GG | 0.672 | 0.685 | 0.744 | 0.762 | 0.614 | 0.631 | 0.678 |
| 1751 | GG**Arg**GG**His**GG | 0.816 | 0.706 | 0.706 | 0.705 | 0.742 | 0.782 | 0.724 |
| 1752 | GG**Arg**GG**Ile**GG | 0.745 | 0.796 | 0.690 | 0.758 | 0.759 | 0.749 | 0.754 |
| 1753 | GG**Arg**GG**Leu**GG | 0.656 | 0.703 | 0.708 | 0.838 | 0.810 | 0.823 | 0.759 |
| 1754 | GG**Arg**GG**Lys**GG | 0.797 | 0.695 | 0.724 | 0.711 | 0.751 | 0.723 | 0.723 |
| 1755 | GG**Arg**GG**Met**GG | 0.405 | 0.425 | 0.611 | 0.804 | 0.824 | 0.817 | 0.708 |
| 1756 | GG**Arg**GG**Phe**GG | 0.844 | 0.869 | 0.856 | 0.872 | 0.816 | 0.867 | 0.862 |
| 1757 | GG**Arg**GG**Pro**GG | 0.778 | 0.780 | 0.770 | 0.704 | 0.885 | 0.754 | 0.774 |
| 1758 | GG**Arg**GG**Ptr**GG | 0.660 | 0.718 | 0.687 | 0.864 | 0.750 | 0.776 | 0.734 |
| 1759 | GG**Arg**GG**S1p**GG | 0.742 | 0.647 | 0.671 | 0.644 | 0.640 | 0.730 | 0.659 |
| 1760 | GG**Arg**GG**Sep**GG | 0.742 | 0.700 | 0.711 | 0.757 | 0.729 | 0.706 | 0.720 |
| 1761 | GG**Arg**GG**Ser**GG | 0.774 | 0.813 | 0.800 | 0.860 | 0.870 | 0.858 | 0.835 |
| 1762 | GG**Arg**GG**T1p**GG | 0.372 | 0.534 | 0.719 | 0.775 | 0.372 | 0.483 | 0.509 |
| 1763 | GG**Arg**GG**Thr**GG | 0.692 | 0.823 | 0.744 | 0.686 | 0.740 | 0.761 | 0.742 |
| 1764 | GG**Arg**GG**Tpo**GG | 0.084 | 0.506 | 0.859 | 0.548 | 0.159 | 0.600 | 0.527 |
| 1765 | GG**Arg**GG**Trp**GG | 0.805 | 0.723 | 0.822 | 0.775 | 0.779 | 0.655 | 0.777 |
| 1766 | GG**Arg**GG**Tyr**GG | 0.700 | 0.716 | 0.679 | 0.650 | 0.587 | 0.611 | 0.664 |
| 1767 | GG**Arg**GG**Val**GG | 0.764 | 0.707 | 0.746 | 0.736 | 0.796 | 0.735 | 0.741 |
| 1768 | GG**Arg**GG**Y1p**GG | 0.607 | 0.732 | 0.730 | 0.699 | 0.614 | 0.712 | 0.706 |
| 1769 | GG**Ash**GG**Ala**GG | 0.867 | 0.850 | 0.833 | 0.889 | 0.860 | 0.863 | 0.862 |
| 1770 | GG**Ash**GG**Arg**GG | 0.595 | 0.590 | 0.646 | 0.675 | 0.760 | 0.762 | 0.661 |
| 1771 | GG**Ash**GG**Ash**GG | 0.755 | 0.796 | 0.752 | 0.825 | 0.782 | 0.814 | 0.789 |
| 1772 | GG**Ash**GG**Asn**GG | 0.878 | 0.826 | 0.905 | 0.770 | 0.871 | 0.851 | 0.861 |
| 1773 | GG**Ash**GG**Asp**GG | 0.641 | 0.620 | 0.677 | 0.804 | 0.881 | 0.827 | 0.741 |
| 1774 | GG**Ash**GG**Cys**GG | 0.848 | 0.846 | 0.800 | 0.841 | 0.832 | 0.890 | 0.844 |
| 1775 | GG**Ash**GG**Glh**GG | 0.761 | 0.736 | 0.877 | 0.787 | 0.786 | 0.789 | 0.787 |
| 1776 | GG**Ash**GG**Gln**GG | 0.746 | 0.728 | 0.756 | 0.702 | 0.669 | 0.832 | 0.737 |
| 1777 | GG**Ash**GG**Glu**GG | 0.776 | 0.850 | 0.770 | 0.793 | 0.832 | 0.789 | 0.791 |
| 1778 | GG**Ash**GG**Gly**GG | 0.848 | 0.927 | 0.911 | 0.874 | 0.902 | 0.927 | 0.906 |
| 1779 | GG**Ash**GG**Hip**GG | 0.663 | 0.584 | 0.878 | 0.693 | 0.678 | 0.619 | 0.671 |
| 1780 | GG**Ash**GG**His**GG | 0.771 | 0.776 | 0.732 | 0.770 | 0.775 | 0.797 | 0.773 |
| 1781 | GG**Ash**GG**Ile**GG | 0.670 | 0.788 | 0.846 | 0.733 | 0.687 | 0.818 | 0.761 |
| 1782 | GG**Ash**GG**Leu**GG | 0.802 | 0.820 | 0.756 | 0.851 | 0.784 | 0.823 | 0.811 |
| 1783 | GG**Ash**GG**Lys**GG | 0.527 | 0.499 | 0.551 | 0.818 | 0.811 | 0.817 | 0.681 |
| 1784 | GG**Ash**GG**Met**GG | 0.814 | 0.825 | 0.817 | 0.830 | 0.768 | 0.809 | 0.816 |
| 1785 | GG**Ash**GG**Phe**GG | 0.845 | 0.865 | 0.786 | 0.815 | 0.766 | 0.783 | 0.800 |
| 1786 | GG**Ash**GG**Pro**GG | 0.897 | 0.881 | 0.791 | 0.846 | 0.807 | 0.809 | 0.828 |
| 1787 | GG**Ash**GG**Ptr**GG | 0.646 | 0.687 | 0.621 | 0.838 | 0.844 | 0.809 | 0.748 |
| 1788 | GG**Ash**GG**S1p**GG | 0.783 | 0.812 | 0.770 | 0.817 | 0.808 | 0.771 | 0.795 |
| 1789 | GG**Ash**GG**Sep**GG | 0.606 | 0.800 | 0.661 | 0.697 | 0.758 | 0.725 | 0.711 |
| 1790 | GG**Ash**GG**Ser**GG | 0.804 | 0.883 | 0.831 | 0.813 | 0.765 | 0.835 | 0.822 |
| 1791 | GG**Ash**GG**T1p**GG | 0.614 | 0.309 | 0.783 | 0.652 | 0.652 | 0.433 | 0.633 |
| 1792 | GG**Ash**GG**Thr**GG | 0.784 | 0.804 | 0.817 | 0.763 | 0.778 | 0.804 | 0.794 |
| 1793 | GG**Ash**GG**Tpo**GG | 0.795 | 0.888 | 0.874 | 0.809 | 0.767 | 0.934 | 0.842 |
| 1794 | GG**Ash**GG**Trp**GG | 0.757 | 0.770 | 0.780 | 0.771 | 0.764 | 0.761 | 0.767 |
| 1795 | GG**Ash**GG**Tyr**GG | 0.873 | 0.832 | 0.849 | 0.813 | 0.821 | 0.838 | 0.835 |
| 1796 | GG**Ash**GG**Val**GG | 0.775 | 0.830 | 0.788 | 0.782 | 0.737 | 0.810 | 0.785 |
| 1797 | GG**Ash**GG**Y1p**GG | 0.743 | 0.714 | 0.706 | 0.780 | 0.789 | 0.790 | 0.761 |
| 1798 | GG**Asn**GG**Ala**GG | 0.867 | 0.870 | 0.844 | 0.889 | 0.873 | 0.877 | 0.871 |
| 1799 | GG**Asn**GG**Arg**GG | 0.853 | 0.850 | 0.765 | 0.824 | 0.722 | 0.758 | 0.795 |

This work is shared under a CC BY 4.0 license.

| | Peptide | $r_{replica}(1,2)$ | $r_{replica}(1,3)$ | $r_{replica}(1,4)$ | $r_{replica}(2,3)$ | $r_{replica}(2,4)$ | $r_{replica}(3,4)$ | Median |
|---|---|---|---|---|---|---|---|---|
| 1800 | GG**Asn**GG**Ash**GG | 0.814 | 0.809 | 0.826 | 0.855 | 0.826 | 0.803 | 0.820 |
| 1801 | GG**Asn**GG**Asn**GG | 0.893 | 0.913 | 0.921 | 0.882 | 0.918 | 0.895 | 0.904 |
| 1802 | GG**Asn**GG**Asp**GG | 0.836 | 0.786 | 0.865 | 0.791 | 0.807 | 0.827 | 0.817 |
| 1803 | GG**Asn**GG**Cys**GG | 0.895 | 0.869 | 0.805 | 0.884 | 0.832 | 0.856 | 0.862 |
| 1804 | GG**Asn**GG**Glh**GG | 0.819 | 0.838 | 0.853 | 0.824 | 0.829 | 0.839 | 0.833 |
| 1805 | GG**Asn**GG**Gln**GG | 0.812 | 0.829 | 0.803 | 0.836 | 0.855 | 0.843 | 0.833 |
| 1806 | GG**Asn**GG**Glu**GG | 0.826 | 0.796 | 0.773 | 0.808 | 0.852 | 0.813 | 0.810 |
| 1807 | GG**Asn**GG**Gly**GG | 0.890 | 0.922 | 0.922 | 0.912 | 0.865 | 0.922 | 0.917 |
| 1808 | GG**Asn**GG**Hip**GG | 0.725 | 0.759 | 0.715 | 0.731 | 0.755 | 0.795 | 0.743 |
| 1809 | GG**Asn**GG**His**GG | 0.835 | 0.853 | 0.807 | 0.867 | 0.796 | 0.787 | 0.821 |
| 1810 | GG**Asn**GG**Ile**GG | 0.763 | 0.822 | 0.764 | 0.831 | 0.819 | 0.812 | 0.816 |
| 1811 | GG**Asn**GG**Leu**GG | 0.888 | 0.810 | 0.775 | 0.836 | 0.855 | 0.839 | 0.837 |
| 1812 | GG**Asn**GG**Lys**GG | 0.819 | 0.772 | 0.800 | 0.821 | 0.846 | 0.832 | 0.820 |
| 1813 | GG**Asn**GG**Met**GG | 0.748 | 0.783 | 0.793 | 0.779 | 0.874 | 0.841 | 0.788 |
| 1814 | GG**Asn**GG**Phe**GG | 0.759 | 0.746 | 0.760 | 0.859 | 0.902 | 0.873 | 0.809 |
| 1815 | GG**Asn**GG**Pro**GG | 0.901 | 0.885 | 0.754 | 0.894 | 0.773 | 0.823 | 0.854 |
| 1816 | GG**Asn**GG**Ptr**GG | 0.867 | 0.852 | 0.720 | 0.847 | 0.735 | 0.736 | 0.791 |
| 1817 | GG**Asn**GG**S1p**GG | 0.829 | 0.868 | 0.858 | 0.818 | 0.823 | 0.855 | 0.842 |
| 1818 | GG**Asn**GG**Sep**GG | 0.609 | 0.530 | 0.621 | 0.819 | 0.793 | 0.826 | 0.707 |
| 1819 | GG**Asn**GG**Ser**GG | 0.853 | 0.847 | 0.884 | 0.828 | 0.834 | 0.827 | 0.841 |
| 1820 | GG**Asn**GG**T1p**GG | 0.786 | 0.592 | 0.801 | 0.688 | 0.813 | 0.656 | 0.737 |
| 1821 | GG**Asn**GG**Thr**GG | 0.863 | 0.867 | 0.858 | 0.897 | 0.858 | 0.865 | 0.864 |
| 1822 | GG**Asn**GG**Tpo**GG | 0.886 | 0.876 | 0.830 | 0.866 | 0.782 | 0.821 | 0.848 |
| 1823 | GG**Asn**GG**Trp**GG | 0.819 | 0.801 | 0.732 | 0.830 | 0.761 | 0.704 | 0.781 |
| 1824 | GG**Asn**GG**Tyr**GG | 0.827 | 0.829 | 0.861 | 0.844 | 0.869 | 0.891 | 0.853 |
| 1825 | GG**Asn**GG**Val**GG | 0.852 | 0.713 | 0.832 | 0.648 | 0.780 | 0.741 | 0.760 |
| 1826 | GG**Asn**GG**Y1p**GG | 0.818 | 0.806 | 0.906 | 0.817 | 0.823 | 0.826 | 0.821 |
| 1827 | GG**Asp**GG**Ala**GG | 0.779 | 0.738 | 0.722 | 0.867 | 0.866 | 0.817 | 0.798 |
| 1828 | GG**Asp**GG**Arg**GG | 0.784 | 0.784 | 0.650 | 0.751 | 0.643 | 0.605 | 0.701 |
| 1829 | GG**Asp**GG**Ash**GG | 0.792 | 0.803 | 0.793 | 0.780 | 0.721 | 0.728 | 0.786 |
| 1830 | GG**Asp**GG**Asn**GG | 0.849 | 0.821 | 0.773 | 0.732 | 0.728 | 0.671 | 0.753 |
| 1831 | GG**Asp**GG**Asp**GG | 0.595 | 0.608 | 0.530 | 0.748 | 0.743 | 0.842 | 0.676 |
| 1832 | GG**Asp**GG**Cys**GG | 0.826 | 0.773 | 0.788 | 0.774 | 0.818 | 0.792 | 0.790 |
| 1833 | GG**Asp**GG**Glh**GG | 0.840 | 0.770 | 0.770 | 0.767 | 0.781 | 0.782 | 0.776 |
| 1834 | GG**Asp**GG**Gln**GG | 0.797 | 0.805 | 0.771 | 0.773 | 0.753 | 0.744 | 0.772 |
| 1835 | GG**Asp**GG**Glu**GG | 0.636 | 0.770 | 0.715 | 0.633 | 0.620 | 0.743 | 0.675 |
| 1836 | GG**Asp**GG**Gly**GG | 0.873 | 0.869 | 0.851 | 0.840 | 0.843 | 0.846 | 0.849 |
| 1837 | GG**Asp**GG**Hip**GG | 0.625 | 0.555 | 0.711 | 0.569 | 0.722 | 0.625 | 0.625 |
| 1838 | GG**Asp**GG**His**GG | 0.653 | 0.506 | 0.596 | 0.737 | 0.783 | 0.779 | 0.695 |
| 1839 | GG**Asp**GG**Ile**GG | 0.771 | 0.710 | 0.832 | 0.735 | 0.738 | 0.728 | 0.737 |
| 1840 | GG**Asp**GG**Leu**GG | 0.747 | 0.735 | 0.717 | 0.780 | 0.803 | 0.777 | 0.762 |
| 1841 | GG**Asp**GG**Lys**GG | 0.748 | 0.655 | 0.712 | 0.667 | 0.671 | 0.665 | 0.669 |
| 1842 | GG**Asp**GG**Met**GG | 0.750 | 0.738 | 0.755 | 0.694 | 0.644 | 0.721 | 0.730 |
| 1843 | GG**Asp**GG**Phe**GG | 0.828 | 0.818 | 0.817 | 0.763 | 0.811 | 0.820 | 0.817 |
| 1844 | GG**Asp**GG**Pro**GG | 0.790 | 0.786 | 0.779 | 0.705 | 0.809 | 0.703 | 0.783 |
| 1845 | GG**Asp**GG**Ptr**GG | 0.802 | 0.855 | 0.781 | 0.795 | 0.839 | 0.791 | 0.798 |
| 1846 | GG**Asp**GG**S1p**GG | 0.674 | 0.732 | 0.702 | 0.818 | 0.808 | 0.821 | 0.770 |
| 1847 | GG**Asp**GG**Sep**GG | 0.544 | 0.786 | 0.767 | 0.447 | 0.456 | 0.847 | 0.656 |
| 1848 | GG**Asp**GG**Ser**GG | 0.765 | 0.801 | 0.679 | 0.764 | 0.666 | 0.748 | 0.756 |
| 1849 | GG**Asp**GG**T1p**GG | 0.343 | 0.830 | 0.704 | 0.398 | 0.682 | 0.672 | 0.677 |

This work is shared under a CC BY 4.0 license.

| | Peptide | $r_{replica}(1,2)$ | $r_{replica}(1,3)$ | $r_{replica}(1,4)$ | $r_{replica}(2,3)$ | $r_{replica}(2,4)$ | $r_{replica}(3,4)$ | Median |
|---|---|---|---|---|---|---|---|---|
| 1850 | GG**Asp**GG**Thr**GG | 0.706 | 0.760 | 0.742 | 0.723 | 0.763 | 0.697 | 0.733 |
| 1851 | GG**Asp**GG**Tpo**GG | 0.280 | 0.892 | 0.862 | 0.260 | 0.591 | 0.840 | 0.715 |
| 1852 | GG**Asp**GG**Trp**GG | 0.688 | 0.788 | 0.750 | 0.674 | 0.699 | 0.719 | 0.709 |
| 1853 | GG**Asp**GG**Tyr**GG | 0.744 | 0.827 | 0.847 | 0.742 | 0.737 | 0.813 | 0.778 |
| 1854 | GG**Asp**GG**Val**GG | 0.779 | 0.715 | 0.730 | 0.728 | 0.827 | 0.745 | 0.737 |
| 1855 | GG**Asp**GG**Y1p**GG | 0.735 | 0.721 | 0.735 | 0.786 | 0.767 | 0.824 | 0.751 |
| 1856 | GG**Cys**GG**Ala**GG | 0.828 | 0.845 | 0.836 | 0.883 | 0.869 | 0.870 | 0.857 |
| 1857 | GG**Cys**GG**Arg**GG | 0.716 | 0.790 | 0.679 | 0.742 | 0.742 | 0.789 | 0.742 |
| 1858 | GG**Cys**GG**Ash**GG | 0.836 | 0.825 | 0.783 | 0.827 | 0.752 | 0.785 | 0.805 |
| 1859 | GG**Cys**GG**Asn**GG | 0.846 | 0.823 | 0.812 | 0.816 | 0.804 | 0.864 | 0.820 |
| 1860 | GG**Cys**GG**Asp**GG | 0.842 | 0.755 | 0.791 | 0.761 | 0.845 | 0.774 | 0.783 |
| 1861 | GG**Cys**GG**Cys**GG | 0.749 | 0.818 | 0.826 | 0.819 | 0.782 | 0.820 | 0.819 |
| 1862 | GG**Cys**GG**Glh**GG | 0.817 | 0.822 | 0.737 | 0.857 | 0.758 | 0.764 | 0.791 |
| 1863 | GG**Cys**GG**Gln**GG | 0.809 | 0.815 | 0.853 | 0.818 | 0.850 | 0.802 | 0.817 |
| 1864 | GG**Cys**GG**Glu**GG | 0.767 | 0.757 | 0.589 | 0.788 | 0.708 | 0.779 | 0.762 |
| 1865 | GG**Cys**GG**Gly**GG | 0.911 | 0.931 | 0.897 | 0.929 | 0.902 | 0.918 | 0.914 |
| 1866 | GG**Cys**GG**Hip**GG | 0.728 | 0.731 | 0.783 | 0.825 | 0.829 | 0.824 | 0.804 |
| 1867 | GG**Cys**GG**His**GG | 0.852 | 0.796 | 0.826 | 0.840 | 0.837 | 0.775 | 0.831 |
| 1868 | GG**Cys**GG**Ile**GG | 0.784 | 0.780 | 0.742 | 0.684 | 0.775 | 0.688 | 0.758 |
| 1869 | GG**Cys**GG**Leu**GG | 0.844 | 0.888 | 0.835 | 0.856 | 0.853 | 0.869 | 0.855 |
| 1870 | GG**Cys**GG**Lys**GG | 0.847 | 0.813 | 0.818 | 0.770 | 0.825 | 0.707 | 0.815 |
| 1871 | GG**Cys**GG**Met**GG | 0.767 | 0.767 | 0.774 | 0.804 | 0.795 | 0.888 | 0.785 |
| 1872 | GG**Cys**GG**Phe**GG | 0.724 | 0.776 | 0.878 | 0.657 | 0.758 | 0.790 | 0.767 |
| 1873 | GG**Cys**GG**Pro**GG | 0.863 | 0.880 | 0.851 | 0.844 | 0.873 | 0.839 | 0.857 |
| 1874 | GG**Cys**GG**Ptr**GG | 0.682 | 0.678 | 0.741 | 0.790 | 0.758 | 0.721 | 0.731 |
| 1875 | GG**Cys**GG**S1p**GG | 0.775 | 0.703 | 0.811 | 0.787 | 0.821 | 0.770 | 0.781 |
| 1876 | GG**Cys**GG**Sep**GG | 0.844 | 0.822 | 0.845 | 0.753 | 0.848 | 0.790 | 0.833 |
| 1877 | GG**Cys**GG**Ser**GG | 0.867 | 0.827 | 0.845 | 0.817 | 0.835 | 0.805 | 0.831 |
| 1878 | GG**Cys**GG**T1p**GG | 0.477 | 0.819 | 0.537 | 0.553 | 0.807 | 0.644 | 0.598 |
| 1879 | GG**Cys**GG**Thr**GG | 0.843 | 0.829 | 0.866 | 0.827 | 0.841 | 0.837 | 0.839 |
| 1880 | GG**Cys**GG**Tpo**GG | 0.720 | 0.781 | 0.810 | 0.271 | 0.311 | 0.960 | 0.751 |
| 1881 | GG**Cys**GG**Trp**GG | 0.780 | 0.847 | 0.844 | 0.863 | 0.833 | 0.877 | 0.845 |
| 1882 | GG**Cys**GG**Tyr**GG | 0.882 | 0.862 | 0.895 | 0.852 | 0.885 | 0.848 | 0.872 |
| 1883 | GG**Cys**GG**Val**GG | 0.817 | 0.764 | 0.803 | 0.770 | 0.790 | 0.824 | 0.797 |
| 1884 | GG**Cys**GG**Y1p**GG | 0.721 | 0.741 | 0.693 | 0.803 | 0.832 | 0.791 | 0.766 |
| 1885 | GG**Glh**GG**Ala**GG | 0.849 | 0.850 | 0.843 | 0.813 | 0.884 | 0.814 | 0.846 |
| 1886 | GG**Glh**GG**Arg**GG | 0.736 | 0.597 | 0.668 | 0.608 | 0.636 | 0.602 | 0.622 |
| 1887 | GG**Glh**GG**Ash**GG | 0.847 | 0.854 | 0.860 | 0.864 | 0.857 | 0.859 | 0.858 |
| 1888 | GG**Glh**GG**Asn**GG | 0.851 | 0.859 | 0.864 | 0.849 | 0.845 | 0.854 | 0.852 |
| 1889 | GG**Glh**GG**Asp**GG | 0.883 | 0.879 | 0.825 | 0.900 | 0.788 | 0.817 | 0.852 |
| 1890 | GG**Glh**GG**Cys**GG | 0.809 | 0.813 | 0.801 | 0.839 | 0.853 | 0.863 | 0.826 |
| 1891 | GG**Glh**GG**Glh**GG | 0.470 | 0.799 | 0.808 | 0.454 | 0.483 | 0.828 | 0.641 |
| 1892 | GG**Glh**GG**Gln**GG | 0.806 | 0.835 | 0.874 | 0.744 | 0.787 | 0.872 | 0.821 |
| 1893 | GG**Glh**GG**Glu**GG | 0.737 | 0.791 | 0.708 | 0.826 | 0.749 | 0.849 | 0.770 |
| 1894 | GG**Glh**GG**Gly**GG | 0.864 | 0.916 | 0.890 | 0.876 | 0.878 | 0.876 | 0.877 |
| 1895 | GG**Glh**GG**Hip**GG | 0.626 | 0.639 | 0.622 | 0.654 | 0.534 | 0.574 | 0.624 |
| 1896 | GG**Glh**GG**His**GG | 0.859 | 0.859 | 0.838 | 0.836 | 0.819 | 0.844 | 0.841 |
| 1897 | GG**Glh**GG**Ile**GG | 0.814 | 0.841 | 0.843 | 0.829 | 0.789 | 0.834 | 0.831 |
| 1898 | GG**Glh**GG**Leu**GG | 0.798 | 0.763 | 0.714 | 0.800 | 0.699 | 0.764 | 0.763 |
| 1899 | GG**Glh**GG**Lys**GG | 0.790 | 0.803 | 0.764 | 0.785 | 0.797 | 0.774 | 0.788 |

This work is shared under a CC BY 4.0 license.

| | Peptide | $r_{replica}$(1,2) | $r_{replica}$(1,3) | $r_{replica}$(1,4) | $r_{replica}$(2,3) | $r_{replica}$(2,4) | $r_{replica}$(3,4) | Median |
|---|---|---|---|---|---|---|---|---|
| 1900 | GG**Glh**GG**Met**GG | 0.900 | 0.889 | 0.871 | 0.874 | 0.893 | 0.896 | 0.891 |
| 1901 | GG**Glh**GG**Phe**GG | 0.744 | 0.766 | 0.798 | 0.796 | 0.834 | 0.789 | 0.793 |
| 1902 | GG**Glh**GG**Pro**GG | 0.834 | 0.871 | 0.850 | 0.874 | 0.820 | 0.854 | 0.852 |
| 1903 | GG**Glh**GG**Ptr**GG | 0.598 | 0.832 | 0.851 | 0.715 | 0.714 | 0.862 | 0.774 |
| 1904 | GG**Glh**GG**S1p**GG | 0.646 | 0.754 | 0.658 | 0.682 | 0.720 | 0.774 | 0.701 |
| 1905 | GG**Glh**GG**Sep**GG | 0.779 | 0.770 | 0.765 | 0.801 | 0.853 | 0.786 | 0.783 |
| 1906 | GG**Glh**GG**Ser**GG | 0.881 | 0.831 | 0.828 | 0.834 | 0.822 | 0.865 | 0.833 |
| 1907 | GG**Glh**GG**T1p**GG | 0.502 | 0.678 | 0.557 | 0.883 | 0.841 | 0.860 | 0.760 |
| 1908 | GG**Glh**GG**Thr**GG | 0.789 | 0.800 | 0.826 | 0.816 | 0.814 | 0.823 | 0.815 |
| 1909 | GG**Glh**GG**Tpo**GG | 0.881 | 0.808 | 0.881 | 0.778 | 0.874 | 0.804 | 0.841 |
| 1910 | GG**Glh**GG**Trp**GG | 0.729 | 0.860 | 0.858 | 0.697 | 0.736 | 0.895 | 0.797 |
| 1911 | GG**Glh**GG**Tyr**GG | 0.869 | 0.892 | 0.799 | 0.861 | 0.787 | 0.775 | 0.830 |
| 1912 | GG**Glh**GG**Val**GG | 0.745 | 0.794 | 0.848 | 0.814 | 0.795 | 0.821 | 0.804 |
| 1913 | GG**Glh**GG**Y1p**GG | 0.817 | 0.860 | 0.767 | 0.827 | 0.851 | 0.795 | 0.822 |
| 1914 | GG**Gln**GG**Ala**GG | 0.890 | 0.883 | 0.854 | 0.886 | 0.859 | 0.864 | 0.874 |
| 1915 | GG**Gln**GG**Arg**GG | 0.769 | 0.721 | 0.796 | 0.729 | 0.843 | 0.745 | 0.757 |
| 1916 | GG**Gln**GG**Ash**GG | 0.893 | 0.849 | 0.874 | 0.885 | 0.888 | 0.853 | 0.880 |
| 1917 | GG**Gln**GG**Asn**GG | 0.860 | 0.780 | 0.833 | 0.877 | 0.781 | 0.771 | 0.807 |
| 1918 | GG**Gln**GG**Asp**GG | 0.623 | 0.775 | 0.835 | 0.656 | 0.623 | 0.796 | 0.716 |
| 1919 | GG**Gln**GG**Cys**GG | 0.817 | 0.883 | 0.853 | 0.803 | 0.813 | 0.884 | 0.835 |
| 1920 | GG**Gln**GG**Glh**GG | 0.705 | 0.778 | 0.822 | 0.748 | 0.669 | 0.734 | 0.741 |
| 1921 | GG**Gln**GG**Gln**GG | 0.716 | 0.803 | 0.713 | 0.805 | 0.843 | 0.779 | 0.791 |
| 1922 | GG**Gln**GG**Glu**GG | 0.768 | 0.838 | 0.841 | 0.775 | 0.765 | 0.823 | 0.799 |
| 1923 | GG**Gln**GG**Gly**GG | 0.873 | 0.887 | 0.873 | 0.888 | 0.842 | 0.859 | 0.873 |
| 1924 | GG**Gln**GG**Hip**GG | 0.791 | 0.773 | 0.465 | 0.819 | 0.507 | 0.510 | 0.642 |
| 1925 | GG**Gln**GG**His**GG | 0.753 | 0.803 | 0.727 | 0.723 | 0.708 | 0.651 | 0.725 |
| 1926 | GG**Gln**GG**Ile**GG | 0.837 | 0.732 | 0.924 | 0.697 | 0.840 | 0.728 | 0.784 |
| 1927 | GG**Gln**GG**Leu**GG | 0.704 | 0.610 | 0.682 | 0.719 | 0.765 | 0.708 | 0.706 |
| 1928 | GG**Gln**GG**Lys**GG | 0.764 | 0.617 | 0.778 | 0.683 | 0.832 | 0.658 | 0.723 |
| 1929 | GG**Gln**GG**Met**GG | 0.829 | 0.839 | 0.846 | 0.859 | 0.811 | 0.839 | 0.839 |
| 1930 | GG**Gln**GG**Phe**GG | 0.804 | 0.861 | 0.906 | 0.771 | 0.762 | 0.852 | 0.828 |
| 1931 | GG**Gln**GG**Pro**GG | 0.809 | 0.836 | 0.803 | 0.840 | 0.847 | 0.873 | 0.838 |
| 1932 | GG**Gln**GG**Ptr**GG | 0.771 | 0.733 | 0.771 | 0.834 | 0.828 | 0.840 | 0.800 |
| 1933 | GG**Gln**GG**S1p**GG | 0.830 | 0.847 | 0.746 | 0.816 | 0.828 | 0.787 | 0.822 |
| 1934 | GG**Gln**GG**Sep**GG | 0.717 | 0.795 | 0.552 | 0.765 | 0.720 | 0.594 | 0.718 |
| 1935 | GG**Gln**GG**Ser**GG | 0.857 | 0.823 | 0.822 | 0.828 | 0.865 | 0.835 | 0.831 |
| 1936 | GG**Gln**GG**T1p**GG | 0.749 | 0.834 | 0.473 | 0.777 | 0.618 | 0.568 | 0.683 |
| 1937 | GG**Gln**GG**Thr**GG | 0.836 | 0.826 | 0.853 | 0.823 | 0.802 | 0.795 | 0.825 |
| 1938 | GG**Gln**GG**Tpo**GG | 0.873 | 0.555 | 0.852 | 0.630 | 0.873 | 0.579 | 0.741 |
| 1939 | GG**Gln**GG**Trp**GG | 0.833 | 0.815 | 0.815 | 0.837 | 0.774 | 0.752 | 0.815 |
| 1940 | GG**Gln**GG**Tyr**GG | 0.548 | 0.549 | 0.563 | 0.853 | 0.917 | 0.875 | 0.708 |
| 1941 | GG**Gln**GG**Val**GG | 0.678 | 0.763 | 0.674 | 0.758 | 0.659 | 0.720 | 0.699 |
| 1942 | GG**Gln**GG**Y1p**GG | 0.798 | 0.730 | 0.840 | 0.690 | 0.781 | 0.732 | 0.756 |
| 1943 | GG**Glu**GG**Ala**GG | 0.845 | 0.837 | 0.854 | 0.787 | 0.814 | 0.823 | 0.830 |
| 1944 | GG**Glu**GG**Arg**GG | 0.692 | 0.644 | 0.717 | 0.641 | 0.686 | 0.575 | 0.665 |
| 1945 | GG**Glu**GG**Ash**GG | 0.661 | 0.689 | 0.793 | 0.714 | 0.704 | 0.815 | 0.709 |
| 1946 | GG**Glu**GG**Asn**GG | 0.866 | 0.842 | 0.882 | 0.844 | 0.848 | 0.824 | 0.846 |
| 1947 | GG**Glu**GG**Asp**GG | 0.731 | 0.769 | 0.744 | 0.762 | 0.801 | 0.775 | 0.766 |
| 1948 | GG**Glu**GG**Cys**GG | 0.555 | 0.754 | 0.643 | 0.653 | 0.710 | 0.731 | 0.681 |
| 1949 | GG**Glu**GG**Glh**GG | 0.777 | 0.700 | 0.802 | 0.760 | 0.842 | 0.782 | 0.779 |

This work is shared under a CC BY 4.0 license.

| | Peptide | $r_{replica}(1,2)$ | $r_{replica}(1,3)$ | $r_{replica}(1,4)$ | $r_{replica}(2,3)$ | $r_{replica}(2,4)$ | $r_{replica}(3,4)$ | Median |
|---|---|---|---|---|---|---|---|---|
| 1950 | GG**Glu**GG**Gln**GG | 0.757 | 0.869 | 0.706 | 0.775 | 0.749 | 0.758 | 0.757 |
| 1951 | GG**Glu**GG**Glu**GG | 0.808 | 0.787 | 0.835 | 0.754 | 0.711 | 0.693 | 0.771 |
| 1952 | GG**Glu**GG**Gly**GG | 0.897 | 0.891 | 0.907 | 0.861 | 0.875 | 0.866 | 0.883 |
| 1953 | GG**Glu**GG**Hip**GG | 0.647 | 0.721 | 0.661 | 0.785 | 0.725 | 0.702 | 0.712 |
| 1954 | GG**Glu**GG**His**GG | 0.837 | 0.556 | 0.586 | 0.544 | 0.630 | 0.767 | 0.608 |
| 1955 | GG**Glu**GG**Ile**GG | 0.603 | 0.799 | 0.779 | 0.651 | 0.668 | 0.800 | 0.724 |
| 1956 | GG**Glu**GG**Leu**GG | 0.859 | 0.856 | 0.823 | 0.840 | 0.869 | 0.866 | 0.857 |
| 1957 | GG**Glu**GG**Lys**GG | 0.674 | 0.760 | 0.735 | 0.772 | 0.742 | 0.765 | 0.751 |
| 1958 | GG**Glu**GG**Met**GG | 0.776 | 0.692 | 0.666 | 0.785 | 0.750 | 0.745 | 0.748 |
| 1959 | GG**Glu**GG**Phe**GG | 0.830 | 0.760 | 0.837 | 0.772 | 0.912 | 0.767 | 0.801 |
| 1960 | GG**Glu**GG**Pro**GG | 0.846 | 0.829 | 0.828 | 0.844 | 0.895 | 0.867 | 0.845 |
| 1961 | GG**Glu**GG**Ptr**GG | 0.545 | 0.575 | 0.763 | 0.835 | 0.669 | 0.746 | 0.708 |
| 1962 | GG**Glu**GG**S1p**GG | 0.778 | 0.759 | 0.820 | 0.766 | 0.792 | 0.754 | 0.772 |
| 1963 | GG**Glu**GG**Sep**GG | 0.770 | 0.768 | 0.762 | 0.768 | 0.806 | 0.761 | 0.768 |
| 1964 | GG**Glu**GG**Ser**GG | 0.886 | 0.894 | 0.897 | 0.862 | 0.897 | 0.893 | 0.893 |
| 1965 | GG**Glu**GG**T1p**GG | 0.909 | 0.822 | 0.861 | 0.890 | 0.919 | 0.901 | 0.896 |
| 1966 | GG**Glu**GG**Thr**GG | 0.836 | 0.861 | 0.803 | 0.856 | 0.793 | 0.796 | 0.820 |
| 1967 | GG**Glu**GG**Tpo**GG | 0.805 | 0.766 | 0.639 | 0.766 | 0.647 | 0.781 | 0.766 |
| 1968 | GG**Glu**GG**Trp**GG | 0.764 | 0.753 | 0.792 | 0.820 | 0.849 | 0.754 | 0.778 |
| 1969 | GG**Glu**GG**Tyr**GG | 0.769 | 0.827 | 0.792 | 0.656 | 0.657 | 0.834 | 0.780 |
| 1970 | GG**Glu**GG**Val**GG | 0.829 | 0.747 | 0.789 | 0.801 | 0.808 | 0.763 | 0.795 |
| 1971 | GG**Glu**GG**Y1p**GG | 0.835 | 0.826 | 0.833 | 0.801 | 0.841 | 0.851 | 0.834 |
| 1972 | GG**Gly**GG**Ala**GG | 0.914 | 0.912 | 0.935 | 0.886 | 0.932 | 0.893 | 0.913 |
| 1973 | GG**Gly**GG**Arg**GG | 0.828 | 0.763 | 0.835 | 0.719 | 0.769 | 0.793 | 0.781 |
| 1974 | GG**Gly**GG**Ash**GG | 0.868 | 0.881 | 0.902 | 0.906 | 0.883 | 0.904 | 0.893 |
| 1975 | GG**Gly**GG**Asn**GG | 0.864 | 0.863 | 0.797 | 0.903 | 0.863 | 0.864 | 0.863 |
| 1976 | GG**Gly**GG**Asp**GG | 0.833 | 0.863 | 0.827 | 0.749 | 0.918 | 0.712 | 0.830 |
| 1977 | GG**Gly**GG**Cys**GG | 0.899 | 0.910 | 0.912 | 0.896 | 0.899 | 0.907 | 0.903 |
| 1978 | GG**Gly**GG**Glh**GG | 0.884 | 0.896 | 0.851 | 0.910 | 0.867 | 0.877 | 0.881 |
| 1979 | GG**Gly**GG**Gln**GG | 0.853 | 0.873 | 0.903 | 0.894 | 0.884 | 0.897 | 0.889 |
| 1980 | GG**Gly**GG**Glu**GG | 0.924 | 0.904 | 0.864 | 0.895 | 0.856 | 0.892 | 0.894 |
| 1981 | GG**Gly**GG**Gly**GG | 0.941 | 0.934 | 0.917 | 0.931 | 0.918 | 0.923 | 0.927 |
| 1982 | GG**Gly**GG**Hip**GG | 0.837 | 0.800 | 0.860 | 0.808 | 0.831 | 0.833 | 0.832 |
| 1983 | GG**Gly**GG**His**GG | 0.839 | 0.855 | 0.884 | 0.891 | 0.891 | 0.892 | 0.887 |
| 1984 | GG**Gly**GG**Ile**GG | 0.843 | 0.840 | 0.853 | 0.874 | 0.899 | 0.881 | 0.864 |
| 1985 | GG**Gly**GG**Leu**GG | 0.869 | 0.868 | 0.893 | 0.878 | 0.906 | 0.888 | 0.883 |
| 1986 | GG**Gly**GG**Lys**GG | 0.847 | 0.845 | 0.813 | 0.868 | 0.868 | 0.835 | 0.846 |
| 1987 | GG**Gly**GG**Met**GG | 0.921 | 0.930 | 0.910 | 0.916 | 0.909 | 0.910 | 0.913 |
| 1988 | GG**Gly**GG**Phe**GG | 0.888 | 0.900 | 0.901 | 0.872 | 0.877 | 0.909 | 0.894 |
| 1989 | GG**Gly**GG**Pro**GG | 0.897 | 0.907 | 0.914 | 0.913 | 0.882 | 0.897 | 0.902 |
| 1990 | GG**Gly**GG**Ptr**GG | 0.898 | 0.869 | 0.912 | 0.903 | 0.911 | 0.921 | 0.907 |
| 1991 | GG**Gly**GG**S1p**GG | 0.860 | 0.762 | 0.791 | 0.827 | 0.862 | 0.900 | 0.844 |
| 1992 | GG**Gly**GG**Sep**GG | 0.845 | 0.821 | 0.851 | 0.848 | 0.867 | 0.846 | 0.847 |
| 1993 | GG**Gly**GG**Ser**GG | 0.886 | 0.913 | 0.897 | 0.907 | 0.918 | 0.915 | 0.910 |
| 1994 | GG**Gly**GG**T1p**GG | 0.929 | 0.914 | 0.927 | 0.908 | 0.900 | 0.926 | 0.920 |
| 1995 | GG**Gly**GG**Thr**GG | 0.921 | 0.935 | 0.922 | 0.929 | 0.932 | 0.912 | 0.926 |
| 1996 | GG**Gly**GG**Tpo**GG | 0.571 | 0.943 | 0.330 | 0.689 | 0.885 | 0.471 | 0.630 |
| 1997 | GG**Gly**GG**Trp**GG | 0.747 | 0.731 | 0.726 | 0.731 | 0.776 | 0.794 | 0.739 |
| 1998 | GG**Gly**GG**Tyr**GG | 0.864 | 0.854 | 0.889 | 0.806 | 0.885 | 0.857 | 0.860 |
| 1999 | GG**Gly**GG**Val**GG | 0.805 | 0.839 | 0.856 | 0.843 | 0.828 | 0.868 | 0.841 |

This work is shared under a CC BY 4.0 license.

| | Peptide | $r_{replica}$(1,2) | $r_{replica}$(1,3) | $r_{replica}$(1,4) | $r_{replica}$(2,3) | $r_{replica}$(2,4) | $r_{replica}$(3,4) | Median |
|---|---|---|---|---|---|---|---|---|
| 2000 | GG**Gly**GG**Y1p**GG | 0.841 | 0.847 | 0.870 | 0.870 | 0.831 | 0.864 | 0.855 |
| 2001 | GG**Hip**GG**Ala**GG | 0.821 | 0.823 | 0.797 | 0.852 | 0.801 | 0.816 | 0.818 |
| 2002 | GG**Hip**GG**Arg**GG | 0.692 | 0.724 | 0.695 | 0.831 | 0.813 | 0.831 | 0.768 |
| 2003 | GG**Hip**GG**Ash**GG | 0.891 | 0.875 | 0.480 | 0.906 | 0.490 | 0.463 | 0.682 |
| 2004 | GG**Hip**GG**Asn**GG | 0.880 | 0.838 | 0.861 | 0.819 | 0.903 | 0.801 | 0.849 |
| 2005 | GG**Hip**GG**Asp**GG | 0.844 | 0.863 | 0.852 | 0.846 | 0.806 | 0.832 | 0.845 |
| 2006 | GG**Hip**GG**Cys**GG | 0.752 | 0.795 | 0.780 | 0.753 | 0.718 | 0.788 | 0.766 |
| 2007 | GG**Hip**GG**Glh**GG | 0.765 | 0.826 | 0.749 | 0.812 | 0.790 | 0.831 | 0.801 |
| 2008 | GG**Hip**GG**Gln**GG | 0.810 | 0.781 | 0.806 | 0.811 | 0.776 | 0.715 | 0.794 |
| 2009 | GG**Hip**GG**Glu**GG | 0.724 | 0.784 | 0.814 | 0.793 | 0.788 | 0.781 | 0.786 |
| 2010 | GG**Hip**GG**Gly**GG | 0.835 | 0.818 | 0.816 | 0.790 | 0.816 | 0.676 | 0.816 |
| 2011 | GG**Hip**GG**Hip**GG | 0.814 | 0.835 | 0.795 | 0.880 | 0.848 | 0.811 | 0.825 |
| 2012 | GG**Hip**GG**His**GG | 0.814 | 0.798 | 0.805 | 0.804 | 0.835 | 0.720 | 0.805 |
| 2013 | GG**Hip**GG**Ile**GG | 0.804 | 0.866 | 0.839 | 0.813 | 0.755 | 0.836 | 0.824 |
| 2014 | GG**Hip**GG**Leu**GG | 0.822 | 0.856 | 0.792 | 0.850 | 0.815 | 0.837 | 0.830 |
| 2015 | GG**Hip**GG**Lys**GG | 0.816 | 0.810 | 0.868 | 0.777 | 0.815 | 0.802 | 0.813 |
| 2016 | GG**Hip**GG**Met**GG | 0.761 | 0.812 | 0.670 | 0.783 | 0.652 | 0.707 | 0.734 |
| 2017 | GG**Hip**GG**Phe**GG | 0.818 | 0.728 | 0.831 | 0.832 | 0.839 | 0.760 | 0.825 |
| 2018 | GG**Hip**GG**Pro**GG | 0.841 | 0.787 | 0.778 | 0.727 | 0.773 | 0.785 | 0.782 |
| 2019 | GG**Hip**GG**Ptr**GG | 0.688 | 0.514 | 0.724 | 0.746 | 0.734 | 0.664 | 0.706 |
| 2020 | GG**Hip**GG**S1p**GG | 0.816 | 0.828 | 0.822 | 0.826 | 0.759 | 0.810 | 0.819 |
| 2021 | GG**Hip**GG**Sep**GG | 0.806 | 0.457 | 0.763 | 0.413 | 0.790 | 0.516 | 0.640 |
| 2022 | GG**Hip**GG**Ser**GG | 0.716 | 0.730 | 0.854 | 0.784 | 0.833 | 0.816 | 0.800 |
| 2023 | GG**Hip**GG**T1p**GG | 0.747 | 0.673 | 0.493 | 0.640 | 0.478 | 0.741 | 0.657 |
| 2024 | GG**Hip**GG**Thr**GG | 0.865 | 0.871 | 0.859 | 0.858 | 0.868 | 0.850 | 0.862 |
| 2025 | GG**Hip**GG**Tpo**GG | 0.887 | 0.717 | 0.690 | 0.759 | 0.632 | 0.513 | 0.703 |
| 2026 | GG**Hip**GG**Trp**GG | 0.664 | 0.665 | 0.702 | 0.738 | 0.756 | 0.758 | 0.720 |
| 2027 | GG**Hip**GG**Tyr**GG | 0.748 | 0.767 | 0.839 | 0.785 | 0.643 | 0.728 | 0.758 |
| 2028 | GG**Hip**GG**Val**GG | 0.845 | 0.866 | 0.799 | 0.881 | 0.819 | 0.820 | 0.832 |
| 2029 | GG**Hip**GG**Y1p**GG | 0.825 | 0.850 | 0.846 | 0.786 | 0.745 | 0.819 | 0.822 |
| 2030 | GG**His**GG**Ala**GG | 0.850 | 0.818 | 0.867 | 0.792 | 0.829 | 0.803 | 0.823 |
| 2031 | GG**His**GG**Arg**GG | 0.769 | 0.749 | 0.602 | 0.774 | 0.620 | 0.606 | 0.684 |
| 2032 | GG**His**GG**Ash**GG | 0.832 | 0.680 | 0.851 | 0.719 | 0.874 | 0.730 | 0.781 |
| 2033 | GG**His**GG**Asn**GG | 0.690 | 0.810 | 0.815 | 0.566 | 0.645 | 0.747 | 0.719 |
| 2034 | GG**His**GG**Asp**GG | 0.628 | 0.778 | 0.678 | 0.704 | 0.691 | 0.732 | 0.697 |
| 2035 | GG**His**GG**Cys**GG | 0.797 | 0.654 | 0.822 | 0.645 | 0.781 | 0.703 | 0.742 |
| 2036 | GG**His**GG**Glh**GG | 0.580 | 0.617 | 0.684 | 0.628 | 0.574 | 0.585 | 0.601 |
| 2037 | GG**His**GG**Gln**GG | 0.773 | 0.762 | 0.704 | 0.839 | 0.740 | 0.762 | 0.762 |
| 2038 | GG**His**GG**Glu**GG | 0.725 | 0.755 | 0.711 | 0.640 | 0.762 | 0.654 | 0.718 |
| 2039 | GG**His**GG**Gly**GG | 0.755 | 0.783 | 0.803 | 0.861 | 0.744 | 0.771 | 0.777 |
| 2040 | GG**His**GG**Hip**GG | 0.626 | 0.629 | 0.710 | 0.655 | 0.629 | 0.661 | 0.642 |
| 2041 | GG**His**GG**His**GG | 0.774 | 0.818 | 0.765 | 0.806 | 0.728 | 0.677 | 0.769 |
| 2042 | GG**His**GG**Ile**GG | 0.714 | 0.770 | 0.741 | 0.809 | 0.810 | 0.805 | 0.788 |
| 2043 | GG**His**GG**Leu**GG | 0.814 | 0.820 | 0.838 | 0.821 | 0.819 | 0.741 | 0.819 |
| 2044 | GG**His**GG**Lys**GG | 0.755 | 0.705 | 0.521 | 0.749 | 0.600 | 0.633 | 0.669 |
| 2045 | GG**His**GG**Met**GG | 0.819 | 0.749 | 0.784 | 0.836 | 0.827 | 0.759 | 0.802 |
| 2046 | GG**His**GG**Phe**GG | 0.749 | 0.830 | 0.808 | 0.676 | 0.748 | 0.808 | 0.778 |
| 2047 | GG**His**GG**Pro**GG | 0.832 | 0.793 | 0.813 | 0.754 | 0.773 | 0.739 | 0.783 |
| 2048 | GG**His**GG**Ptr**GG | 0.840 | 0.446 | 0.791 | 0.422 | 0.756 | 0.424 | 0.601 |
| 2049 | GG**His**GG**S1p**GG | 0.764 | 0.772 | 0.442 | 0.689 | 0.503 | 0.427 | 0.596 |

This work is shared under a CC BY 4.0 license.

| | Peptide | $r_{replica}(1,2)$ | $r_{replica}(1,3)$ | $r_{replica}(1,4)$ | $r_{replica}(2,3)$ | $r_{replica}(2,4)$ | $r_{replica}(3,4)$ | Median |
|---|---|---|---|---|---|---|---|---|
| 2050 | GG**His**GG**Sep**GG | 0.840 | 0.820 | 0.736 | 0.807 | 0.616 | 0.672 | 0.771 |
| 2051 | GG**His**GG**Ser**GG | 0.752 | 0.745 | 0.627 | 0.786 | 0.756 | 0.741 | 0.749 |
| 2052 | GG**His**GG**T1p**GG | 0.394 | 0.620 | 0.670 | 0.532 | 0.607 | 0.726 | 0.613 |
| 2053 | GG**His**GG**Thr**GG | 0.853 | 0.806 | 0.852 | 0.786 | 0.794 | 0.856 | 0.829 |
| 2054 | GG**His**GG**Tpo**GG | 0.924 | 0.873 | 0.380 | 0.921 | 0.318 | 0.389 | 0.631 |
| 2055 | GG**His**GG**Trp**GG | 0.745 | 0.713 | 0.818 | 0.773 | 0.804 | 0.712 | 0.759 |
| 2056 | GG**His**GG**Tyr**GG | 0.751 | 0.645 | 0.657 | 0.707 | 0.800 | 0.680 | 0.694 |
| 2057 | GG**His**GG**Val**GG | 0.735 | 0.675 | 0.775 | 0.660 | 0.784 | 0.686 | 0.711 |
| 2058 | GG**His**GG**Y1p**GG | 0.729 | 0.676 | 0.689 | 0.837 | 0.751 | 0.696 | 0.713 |
| 2059 | GG**Ile**GG**Ala**GG | 0.769 | 0.764 | 0.727 | 0.755 | 0.792 | 0.796 | 0.766 |
| 2060 | GG**Ile**GG**Arg**GG | 0.749 | 0.694 | 0.526 | 0.739 | 0.460 | 0.452 | 0.610 |
| 2061 | GG**Ile**GG**Ash**GG | 0.777 | 0.751 | 0.711 | 0.842 | 0.826 | 0.780 | 0.779 |
| 2062 | GG**Ile**GG**Asn**GG | 0.901 | 0.884 | 0.903 | 0.889 | 0.913 | 0.914 | 0.902 |
| 2063 | GG**Ile**GG**Asp**GG | 0.852 | 0.895 | 0.893 | 0.851 | 0.914 | 0.876 | 0.885 |
| 2064 | GG**Ile**GG**Cys**GG | 0.742 | 0.806 | 0.773 | 0.723 | 0.724 | 0.786 | 0.758 |
| 2065 | GG**Ile**GG**Glh**GG | 0.816 | 0.765 | 0.781 | 0.811 | 0.774 | 0.781 | 0.781 |
| 2066 | GG**Ile**GG**Gln**GG | 0.805 | 0.875 | 0.869 | 0.818 | 0.777 | 0.844 | 0.831 |
| 2067 | GG**Ile**GG**Glu**GG | 0.801 | 0.744 | 0.768 | 0.733 | 0.887 | 0.719 | 0.756 |
| 2068 | GG**Ile**GG**Gly**GG | 0.871 | 0.883 | 0.915 | 0.875 | 0.916 | 0.896 | 0.889 |
| 2069 | GG**Ile**GG**Hip**GG | 0.785 | 0.620 | 0.657 | 0.746 | 0.745 | 0.823 | 0.746 |
| 2070 | GG**Ile**GG**His**GG | 0.622 | 0.761 | 0.725 | 0.696 | 0.659 | 0.777 | 0.710 |
| 2071 | GG**Ile**GG**Ile**GG | 0.823 | 0.821 | 0.833 | 0.761 | 0.717 | 0.782 | 0.801 |
| 2072 | GG**Ile**GG**Leu**GG | 0.736 | 0.756 | 0.743 | 0.726 | 0.761 | 0.765 | 0.750 |
| 2073 | GG**Ile**GG**Lys**GG | 0.694 | 0.792 | 0.711 | 0.708 | 0.710 | 0.764 | 0.710 |
| 2074 | GG**Ile**GG**Met**GG | 0.744 | 0.725 | 0.760 | 0.784 | 0.784 | 0.761 | 0.760 |
| 2075 | GG**Ile**GG**Phe**GG | 0.873 | 0.842 | 0.886 | 0.855 | 0.891 | 0.850 | 0.864 |
| 2076 | GG**Ile**GG**Pro**GG | 0.869 | 0.858 | 0.874 | 0.854 | 0.833 | 0.907 | 0.864 |
| 2077 | GG**Ile**GG**Ptr**GG | 0.855 | 0.775 | 0.844 | 0.855 | 0.857 | 0.804 | 0.849 |
| 2078 | GG**Ile**GG**S1p**GG | 0.803 | 0.774 | 0.784 | 0.803 | 0.844 | 0.847 | 0.803 |
| 2079 | GG**Ile**GG**Sep**GG | 0.693 | 0.741 | 0.830 | 0.692 | 0.666 | 0.706 | 0.699 |
| 2080 | GG**Ile**GG**Ser**GG | 0.799 | 0.768 | 0.829 | 0.803 | 0.841 | 0.814 | 0.808 |
| 2081 | GG**Ile**GG**T1p**GG | 0.880 | 0.603 | 0.835 | 0.618 | 0.865 | 0.739 | 0.787 |
| 2082 | GG**Ile**GG**Thr**GG | 0.654 | 0.676 | 0.678 | 0.807 | 0.871 | 0.806 | 0.742 |
| 2083 | GG**Ile**GG**Tpo**GG | 0.566 | 0.520 | 0.668 | 0.688 | 0.707 | 0.709 | 0.678 |
| 2084 | GG**Ile**GG**Trp**GG | 0.728 | 0.538 | 0.685 | 0.596 | 0.761 | 0.641 | 0.663 |
| 2085 | GG**Ile**GG**Tyr**GG | 0.795 | 0.864 | 0.869 | 0.808 | 0.783 | 0.909 | 0.836 |
| 2086 | GG**Ile**GG**Val**GG | 0.782 | 0.809 | 0.780 | 0.803 | 0.744 | 0.796 | 0.789 |
| 2087 | GG**Ile**GG**Y1p**GG | 0.800 | 0.834 | 0.809 | 0.836 | 0.782 | 0.783 | 0.805 |
| 2088 | GG**Leu**GG**Ala**GG | 0.782 | 0.841 | 0.863 | 0.784 | 0.784 | 0.834 | 0.809 |
| 2089 | GG**Leu**GG**Arg**GG | 0.689 | 0.686 | 0.707 | 0.711 | 0.697 | 0.743 | 0.702 |
| 2090 | GG**Leu**GG**Ash**GG | 0.837 | 0.824 | 0.826 | 0.833 | 0.850 | 0.811 | 0.829 |
| 2091 | GG**Leu**GG**Asn**GG | 0.687 | 0.863 | 0.846 | 0.660 | 0.764 | 0.822 | 0.793 |
| 2092 | GG**Leu**GG**Asp**GG | 0.823 | 0.863 | 0.805 | 0.809 | 0.709 | 0.796 | 0.807 |
| 2093 | GG**Leu**GG**Cys**GG | 0.839 | 0.865 | 0.857 | 0.778 | 0.821 | 0.846 | 0.842 |
| 2094 | GG**Leu**GG**Glh**GG | 0.680 | 0.663 | 0.677 | 0.810 | 0.800 | 0.752 | 0.716 |
| 2095 | GG**Leu**GG**Gln**GG | 0.784 | 0.806 | 0.795 | 0.803 | 0.881 | 0.821 | 0.805 |
| 2096 | GG**Leu**GG**Glu**GG | 0.805 | 0.797 | 0.649 | 0.872 | 0.669 | 0.665 | 0.733 |
| 2097 | GG**Leu**GG**Gly**GG | 0.895 | 0.880 | 0.892 | 0.897 | 0.917 | 0.891 | 0.893 |
| 2098 | GG**Leu**GG**Hip**GG | 0.705 | 0.725 | 0.713 | 0.761 | 0.690 | 0.675 | 0.709 |
| 2099 | GG**Leu**GG**His**GG | 0.663 | 0.733 | 0.753 | 0.783 | 0.844 | 0.810 | 0.768 |

This work is shared under a CC BY 4.0 license.

| | Peptide | $r_{replica}(1,2)$ | $r_{replica}(1,3)$ | $r_{replica}(1,4)$ | $r_{replica}(2,3)$ | $r_{replica}(2,4)$ | $r_{replica}(3,4)$ | Median |
|---|---|---|---|---|---|---|---|---|
| 2100 | GG**Leu**GG**Ile**GG | 0.718 | 0.838 | 0.764 | 0.736 | 0.758 | 0.785 | 0.761 |
| 2101 | GG**Leu**GG**Leu**GG | 0.862 | 0.799 | 0.845 | 0.863 | 0.808 | 0.794 | 0.827 |
| 2102 | GG**Leu**GG**Lys**GG | 0.837 | 0.767 | 0.701 | 0.821 | 0.752 | 0.724 | 0.759 |
| 2103 | GG**Leu**GG**Met**GG | 0.755 | 0.811 | 0.803 | 0.734 | 0.790 | 0.824 | 0.796 |
| 2104 | GG**Leu**GG**Phe**GG | 0.845 | 0.868 | 0.851 | 0.894 | 0.868 | 0.855 | 0.861 |
| 2105 | GG**Leu**GG**Pro**GG | 0.807 | 0.851 | 0.848 | 0.839 | 0.782 | 0.836 | 0.837 |
| 2106 | GG**Leu**GG**Ptr**GG | 0.786 | 0.756 | 0.813 | 0.819 | 0.865 | 0.813 | 0.813 |
| 2107 | GG**Leu**GG**S1p**GG | 0.826 | 0.815 | 0.771 | 0.850 | 0.827 | 0.804 | 0.820 |
| 2108 | GG**Leu**GG**Sep**GG | 0.792 | 0.780 | 0.783 | 0.794 | 0.792 | 0.803 | 0.792 |
| 2109 | GG**Leu**GG**Ser**GG | 0.788 | 0.819 | 0.841 | 0.745 | 0.853 | 0.801 | 0.810 |
| 2110 | GG**Leu**GG**T1p**GG | 0.720 | 0.674 | 0.460 | 0.859 | 0.399 | 0.353 | 0.567 |
| 2111 | GG**Leu**GG**Thr**GG | 0.830 | 0.821 | 0.815 | 0.872 | 0.867 | 0.858 | 0.844 |
| 2112 | GG**Leu**GG**Tpo**GG | 0.578 | 0.832 | 0.848 | 0.808 | 0.281 | 0.616 | 0.712 |
| 2113 | GG**Leu**GG**Trp**GG | 0.815 | 0.733 | 0.777 | 0.683 | 0.756 | 0.658 | 0.745 |
| 2114 | GG**Leu**GG**Tyr**GG | 0.891 | 0.866 | 0.890 | 0.835 | 0.884 | 0.885 | 0.885 |
| 2115 | GG**Leu**GG**Val**GG | 0.392 | 0.762 | 0.785 | 0.402 | 0.304 | 0.777 | 0.582 |
| 2116 | GG**Leu**GG**Y1p**GG | 0.662 | 0.736 | 0.726 | 0.624 | 0.640 | 0.685 | 0.674 |
| 2117 | GG**Lys**GG**Ala**GG | 0.789 | 0.799 | 0.800 | 0.857 | 0.839 | 0.884 | 0.819 |
| 2118 | GG**Lys**GG**Arg**GG | 0.689 | 0.805 | 0.755 | 0.765 | 0.651 | 0.803 | 0.760 |
| 2119 | GG**Lys**GG**Ash**GG | 0.907 | 0.886 | 0.852 | 0.895 | 0.876 | 0.815 | 0.881 |
| 2120 | GG**Lys**GG**Asn**GG | 0.866 | 0.858 | 0.862 | 0.860 | 0.858 | 0.870 | 0.861 |
| 2121 | GG**Lys**GG**Asp**GG | 0.687 | 0.655 | 0.672 | 0.756 | 0.742 | 0.709 | 0.698 |
| 2122 | GG**Lys**GG**Cys**GG | 0.819 | 0.774 | 0.881 | 0.703 | 0.809 | 0.798 | 0.804 |
| 2123 | GG**Lys**GG**Glh**GG | 0.635 | 0.684 | 0.614 | 0.787 | 0.709 | 0.801 | 0.696 |
| 2124 | GG**Lys**GG**Gln**GG | 0.769 | 0.775 | 0.801 | 0.743 | 0.813 | 0.772 | 0.773 |
| 2125 | GG**Lys**GG**Glu**GG | 0.711 | 0.785 | 0.685 | 0.864 | 0.806 | 0.802 | 0.794 |
| 2126 | GG**Lys**GG**Gly**GG | 0.797 | 0.901 | 0.887 | 0.789 | 0.834 | 0.865 | 0.850 |
| 2127 | GG**Lys**GG**Hip**GG | 0.710 | 0.718 | 0.694 | 0.754 | 0.752 | 0.841 | 0.735 |
| 2128 | GG**Lys**GG**His**GG | 0.777 | 0.841 | 0.773 | 0.755 | 0.643 | 0.814 | 0.775 |
| 2129 | GG**Lys**GG**Ile**GG | 0.857 | 0.853 | 0.857 | 0.875 | 0.873 | 0.894 | 0.865 |
| 2130 | GG**Lys**GG**Leu**GG | 0.679 | 0.735 | 0.754 | 0.784 | 0.813 | 0.867 | 0.769 |
| 2131 | GG**Lys**GG**Lys**GG | 0.773 | 0.811 | 0.768 | 0.809 | 0.815 | 0.824 | 0.810 |
| 2132 | GG**Lys**GG**Met**GG | 0.815 | 0.789 | 0.774 | 0.804 | 0.824 | 0.842 | 0.809 |
| 2133 | GG**Lys**GG**Phe**GG | 0.824 | 0.849 | 0.823 | 0.894 | 0.886 | 0.894 | 0.867 |
| 2134 | GG**Lys**GG**Pro**GG | 0.789 | 0.742 | 0.803 | 0.854 | 0.892 | 0.882 | 0.828 |
| 2135 | GG**Lys**GG**Ptr**GG | 0.779 | 0.812 | 0.803 | 0.716 | 0.774 | 0.764 | 0.777 |
| 2136 | GG**Lys**GG**S1p**GG | 0.560 | 0.661 | 0.604 | 0.640 | 0.652 | 0.752 | 0.646 |
| 2137 | GG**Lys**GG**Sep**GG | 0.850 | 0.811 | 0.830 | 0.763 | 0.821 | 0.768 | 0.816 |
| 2138 | GG**Lys**GG**Ser**GG | 0.901 | 0.882 | 0.847 | 0.877 | 0.894 | 0.918 | 0.888 |
| 2139 | GG**Lys**GG**T1p**GG | 0.883 | 0.860 | 0.862 | 0.795 | 0.822 | 0.875 | 0.861 |
| 2140 | GG**Lys**GG**Thr**GG | 0.873 | 0.857 | 0.902 | 0.878 | 0.899 | 0.891 | 0.884 |
| 2141 | GG**Lys**GG**Tpo**GG | 0.726 | 0.143 | 0.830 | 0.667 | 0.839 | 0.504 | 0.697 |
| 2142 | GG**Lys**GG**Trp**GG | 0.830 | 0.828 | 0.847 | 0.739 | 0.852 | 0.799 | 0.829 |
| 2143 | GG**Lys**GG**Tyr**GG | 0.798 | 0.879 | 0.827 | 0.845 | 0.852 | 0.877 | 0.848 |
| 2144 | GG**Lys**GG**Val**GG | 0.827 | 0.845 | 0.798 | 0.846 | 0.833 | 0.821 | 0.830 |
| 2145 | GG**Lys**GG**Y1p**GG | 0.743 | 0.708 | 0.748 | 0.667 | 0.706 | 0.714 | 0.711 |
| 2146 | GG**Met**GG**Ala**GG | 0.808 | 0.837 | 0.791 | 0.881 | 0.728 | 0.767 | 0.800 |
| 2147 | GG**Met**GG**Arg**GG | 0.582 | 0.748 | 0.746 | 0.609 | 0.594 | 0.619 | 0.614 |
| 2148 | GG**Met**GG**Ash**GG | 0.843 | 0.886 | 0.853 | 0.828 | 0.833 | 0.859 | 0.848 |
| 2149 | GG**Met**GG**Asn**GG | 0.828 | 0.809 | 0.802 | 0.773 | 0.789 | 0.771 | 0.796 |

This work is shared under a CC BY 4.0 license.

| | Peptide | $r_{replica}(1,2)$ | $r_{replica}(1,3)$ | $r_{replica}(1,4)$ | $r_{replica}(2,3)$ | $r_{replica}(2,4)$ | $r_{replica}(3,4)$ | Median |
|---|---|---|---|---|---|---|---|---|
| 2150 | GG**Met**GG**Asp**GG | 0.880 | 0.848 | 0.789 | 0.831 | 0.831 | 0.798 | 0.831 |
| 2151 | GG**Met**GG**Cys**GG | 0.821 | 0.784 | 0.708 | 0.771 | 0.725 | 0.667 | 0.748 |
| 2152 | GG**Met**GG**Glh**GG | 0.868 | 0.789 | 0.766 | 0.794 | 0.701 | 0.684 | 0.777 |
| 2153 | GG**Met**GG**Gln**GG | 0.882 | 0.845 | 0.822 | 0.846 | 0.844 | 0.792 | 0.845 |
| 2154 | GG**Met**GG**Glu**GG | 0.814 | 0.781 | 0.754 | 0.778 | 0.767 | 0.824 | 0.780 |
| 2155 | GG**Met**GG**Gly**GG | 0.831 | 0.898 | 0.884 | 0.871 | 0.833 | 0.869 | 0.870 |
| 2156 | GG**Met**GG**Hip**GG | 0.796 | 0.755 | 0.782 | 0.806 | 0.805 | 0.760 | 0.789 |
| 2157 | GG**Met**GG**His**GG | 0.785 | 0.829 | 0.790 | 0.769 | 0.794 | 0.761 | 0.788 |
| 2158 | GG**Met**GG**Ile**GG | 0.758 | 0.769 | 0.814 | 0.858 | 0.827 | 0.813 | 0.814 |
| 2159 | GG**Met**GG**Leu**GG | 0.667 | 0.750 | 0.731 | 0.743 | 0.685 | 0.781 | 0.737 |
| 2160 | GG**Met**GG**Lys**GG | 0.778 | 0.690 | 0.732 | 0.792 | 0.741 | 0.763 | 0.752 |
| 2161 | GG**Met**GG**Met**GG | 0.874 | 0.855 | 0.857 | 0.872 | 0.827 | 0.843 | 0.856 |
| 2162 | GG**Met**GG**Phe**GG | 0.734 | 0.855 | 0.811 | 0.820 | 0.812 | 0.872 | 0.816 |
| 2163 | GG**Met**GG**Pro**GG | 0.810 | 0.807 | 0.808 | 0.761 | 0.820 | 0.842 | 0.809 |
| 2164 | GG**Met**GG**Ptr**GG | 0.839 | 0.861 | 0.845 | 0.834 | 0.836 | 0.813 | 0.837 |
| 2165 | GG**Met**GG**S1p**GG | 0.765 | 0.772 | 0.724 | 0.775 | 0.763 | 0.709 | 0.764 |
| 2166 | GG**Met**GG**Sep**GG | 0.851 | 0.868 | 0.864 | 0.868 | 0.875 | 0.870 | 0.868 |
| 2167 | GG**Met**GG**Ser**GG | 0.850 | 0.853 | 0.813 | 0.846 | 0.824 | 0.812 | 0.835 |
| 2168 | GG**Met**GG**T1p**GG | 0.715 | 0.788 | 0.796 | 0.755 | 0.806 | 0.849 | 0.792 |
| 2169 | GG**Met**GG**Thr**GG | 0.847 | 0.856 | 0.759 | 0.838 | 0.699 | 0.704 | 0.798 |
| 2170 | GG**Met**GG**Tpo**GG | 0.547 | 0.817 | 0.828 | 0.328 | 0.496 | 0.842 | 0.682 |
| 2171 | GG**Met**GG**Trp**GG | 0.838 | 0.777 | 0.765 | 0.861 | 0.831 | 0.762 | 0.804 |
| 2172 | GG**Met**GG**Tyr**GG | 0.717 | 0.820 | 0.753 | 0.839 | 0.825 | 0.867 | 0.822 |
| 2173 | GG**Met**GG**Val**GG | 0.663 | 0.678 | 0.660 | 0.835 | 0.808 | 0.812 | 0.743 |
| 2174 | GG**Met**GG**Y1p**GG | 0.790 | 0.783 | 0.808 | 0.857 | 0.872 | 0.829 | 0.818 |
| 2175 | GG**Phe**GG**Ala**GG | 0.806 | 0.819 | 0.875 | 0.871 | 0.853 | 0.859 | 0.856 |
| 2176 | GG**Phe**GG**Arg**GG | 0.830 | 0.862 | 0.534 | 0.853 | 0.575 | 0.487 | 0.702 |
| 2177 | GG**Phe**GG**Ash**GG | 0.933 | 0.743 | 0.835 | 0.770 | 0.845 | 0.735 | 0.803 |
| 2178 | GG**Phe**GG**Asn**GG | 0.870 | 0.860 | 0.838 | 0.819 | 0.832 | 0.808 | 0.835 |
| 2179 | GG**Phe**GG**Asp**GG | 0.858 | 0.844 | 0.791 | 0.855 | 0.864 | 0.781 | 0.850 |
| 2180 | GG**Phe**GG**Cys**GG | 0.795 | 0.829 | 0.852 | 0.854 | 0.874 | 0.871 | 0.853 |
| 2181 | GG**Phe**GG**Glh**GG | 0.875 | 0.857 | 0.894 | 0.846 | 0.863 | 0.875 | 0.869 |
| 2182 | GG**Phe**GG**Gln**GG | 0.877 | 0.877 | 0.768 | 0.856 | 0.806 | 0.712 | 0.831 |
| 2183 | GG**Phe**GG**Glu**GG | 0.730 | 0.880 | 0.856 | 0.681 | 0.632 | 0.819 | 0.774 |
| 2184 | GG**Phe**GG**Gly**GG | 0.849 | 0.889 | 0.869 | 0.888 | 0.909 | 0.890 | 0.889 |
| 2185 | GG**Phe**GG**Hip**GG | 0.699 | 0.765 | 0.745 | 0.832 | 0.681 | 0.656 | 0.722 |
| 2186 | GG**Phe**GG**His**GG | 0.870 | 0.852 | 0.810 | 0.815 | 0.808 | 0.767 | 0.812 |
| 2187 | GG**Phe**GG**Ile**GG | 0.883 | 0.894 | 0.795 | 0.882 | 0.782 | 0.784 | 0.839 |
| 2188 | GG**Phe**GG**Leu**GG | 0.767 | 0.825 | 0.833 | 0.825 | 0.744 | 0.836 | 0.825 |
| 2189 | GG**Phe**GG**Lys**GG | 0.713 | 0.790 | 0.718 | 0.788 | 0.774 | 0.793 | 0.781 |
| 2190 | GG**Phe**GG**Met**GG | 0.836 | 0.818 | 0.843 | 0.823 | 0.828 | 0.870 | 0.832 |
| 2191 | GG**Phe**GG**Phe**GG | 0.876 | 0.880 | 0.839 | 0.822 | 0.860 | 0.791 | 0.850 |
| 2192 | GG**Phe**GG**Pro**GG | 0.909 | 0.826 | 0.795 | 0.825 | 0.831 | 0.810 | 0.825 |
| 2193 | GG**Phe**GG**Ptr**GG | 0.835 | 0.844 | 0.789 | 0.808 | 0.734 | 0.753 | 0.799 |
| 2194 | GG**Phe**GG**S1p**GG | 0.726 | 0.805 | 0.762 | 0.867 | 0.864 | 0.874 | 0.835 |
| 2195 | GG**Phe**GG**Sep**GG | 0.813 | 0.757 | 0.725 | 0.747 | 0.705 | 0.648 | 0.736 |
| 2196 | GG**Phe**GG**Ser**GG | 0.827 | 0.889 | 0.832 | 0.778 | 0.773 | 0.813 | 0.820 |
| 2197 | GG**Phe**GG**T1p**GG | 0.830 | 0.483 | 0.727 | 0.568 | 0.784 | 0.649 | 0.688 |
| 2198 | GG**Phe**GG**Thr**GG | 0.824 | 0.745 | 0.739 | 0.770 | 0.740 | 0.760 | 0.752 |
| 2199 | GG**Phe**GG**Tpo**GG | 0.699 | 0.832 | 0.848 | 0.744 | 0.735 | 0.834 | 0.788 |

This work is shared under a CC BY 4.0 license.

| | Peptide | $r_{replica}$(1,2) | $r_{replica}$(1,3) | $r_{replica}$(1,4) | $r_{replica}$(2,3) | $r_{replica}$(2,4) | $r_{replica}$(3,4) | Median |
|---|---|---|---|---|---|---|---|---|
| 2200 | GG**Phe**GG**Trp**GG | 0.852 | 0.837 | 0.844 | 0.785 | 0.810 | 0.851 | 0.841 |
| 2201 | GG**Phe**GG**Tyr**GG | 0.819 | 0.876 | 0.911 | 0.809 | 0.789 | 0.893 | 0.847 |
| 2202 | GG**Phe**GG**Val**GG | 0.799 | 0.834 | 0.771 | 0.823 | 0.761 | 0.799 | 0.799 |
| 2203 | GG**Phe**GG**Y1p**GG | 0.915 | 0.873 | 0.764 | 0.891 | 0.772 | 0.805 | 0.839 |
| 2204 | GG**Pro**GG**Ala**GG | 0.834 | 0.728 | 0.839 | 0.682 | 0.825 | 0.711 | 0.776 |
| 2205 | GG**Pro**GG**Arg**GG | 0.654 | 0.648 | 0.726 | 0.731 | 0.768 | 0.706 | 0.716 |
| 2206 | GG**Pro**GG**Ash**GG | 0.741 | 0.876 | 0.902 | 0.596 | 0.613 | 0.917 | 0.808 |
| 2207 | GG**Pro**GG**Asn**GG | 0.830 | 0.893 | 0.891 | 0.814 | 0.828 | 0.883 | 0.856 |
| 2208 | GG**Pro**GG**Asp**GG | 0.875 | 0.748 | 0.878 | 0.803 | 0.845 | 0.756 | 0.824 |
| 2209 | GG**Pro**GG**Cys**GG | 0.862 | 0.842 | 0.894 | 0.814 | 0.847 | 0.800 | 0.844 |
| 2210 | GG**Pro**GG**Glh**GG | 0.811 | 0.808 | 0.797 | 0.852 | 0.812 | 0.838 | 0.812 |
| 2211 | GG**Pro**GG**Gln**GG | 0.831 | 0.849 | 0.759 | 0.811 | 0.728 | 0.815 | 0.813 |
| 2212 | GG**Pro**GG**Glu**GG | 0.803 | 0.847 | 0.703 | 0.884 | 0.750 | 0.717 | 0.776 |
| 2213 | GG**Pro**GG**Gly**GG | 0.838 | 0.909 | 0.888 | 0.855 | 0.876 | 0.879 | 0.878 |
| 2214 | GG**Pro**GG**Hip**GG | 0.864 | 0.797 | 0.757 | 0.825 | 0.831 | 0.812 | 0.819 |
| 2215 | GG**Pro**GG**His**GG | 0.847 | 0.768 | 0.825 | 0.817 | 0.845 | 0.784 | 0.821 |
| 2216 | GG**Pro**GG**Ile**GG | 0.786 | 0.810 | 0.758 | 0.868 | 0.827 | 0.865 | 0.818 |
| 2217 | GG**Pro**GG**Leu**GG | 0.749 | 0.804 | 0.828 | 0.822 | 0.786 | 0.834 | 0.813 |
| 2218 | GG**Pro**GG**Lys**GG | 0.830 | 0.885 | 0.852 | 0.826 | 0.794 | 0.826 | 0.828 |
| 2219 | GG**Pro**GG**Met**GG | 0.741 | 0.768 | 0.760 | 0.788 | 0.736 | 0.783 | 0.764 |
| 2220 | GG**Pro**GG**Phe**GG | 0.773 | 0.857 | 0.847 | 0.781 | 0.767 | 0.834 | 0.807 |
| 2221 | GG**Pro**GG**Pro**GG | 0.920 | 0.849 | 0.863 | 0.852 | 0.861 | 0.897 | 0.862 |
| 2222 | GG**Pro**GG**Ptr**GG | 0.827 | 0.821 | 0.823 | 0.678 | 0.743 | 0.774 | 0.797 |
| 2223 | GG**Pro**GG**S1p**GG | 0.861 | 0.776 | 0.790 | 0.817 | 0.817 | 0.810 | 0.813 |
| 2224 | GG**Pro**GG**Sep**GG | 0.869 | 0.851 | 0.854 | 0.894 | 0.872 | 0.880 | 0.870 |
| 2225 | GG**Pro**GG**Ser**GG | 0.887 | 0.896 | 0.815 | 0.868 | 0.780 | 0.770 | 0.841 |
| 2226 | GG**Pro**GG**T1p**GG | 0.821 | 0.740 | 0.724 | 0.765 | 0.710 | 0.916 | 0.753 |
| 2227 | GG**Pro**GG**Thr**GG | 0.855 | 0.894 | 0.891 | 0.892 | 0.844 | 0.909 | 0.892 |
| 2228 | GG**Pro**GG**Tpo**GG | 0.364 | 0.855 | 0.932 | 0.646 | 0.494 | 0.872 | 0.750 |
| 2229 | GG**Pro**GG**Trp**GG | 0.708 | 0.518 | 0.575 | 0.680 | 0.822 | 0.705 | 0.693 |
| 2230 | GG**Pro**GG**Tyr**GG | 0.858 | 0.866 | 0.846 | 0.863 | 0.899 | 0.862 | 0.862 |
| 2231 | GG**Pro**GG**Val**GG | 0.721 | 0.780 | 0.823 | 0.811 | 0.634 | 0.750 | 0.765 |
| 2232 | GG**Pro**GG**Y1p**GG | 0.844 | 0.826 | 0.674 | 0.799 | 0.627 | 0.640 | 0.737 |
| 2233 | GG**Ptr**GG**Ala**GG | 0.808 | 0.840 | 0.729 | 0.821 | 0.724 | 0.827 | 0.814 |
| 2234 | GG**Ptr**GG**Arg**GG | 0.479 | 0.377 | 0.452 | 0.452 | 0.346 | 0.586 | 0.452 |
| 2235 | GG**Ptr**GG**Ash**GG | 0.667 | 0.825 | 0.784 | 0.772 | 0.762 | 0.848 | 0.778 |
| 2236 | GG**Ptr**GG**Asn**GG | 0.729 | 0.714 | 0.738 | 0.775 | 0.745 | 0.760 | 0.742 |
| 2237 | GG**Ptr**GG**Asp**GG | 0.712 | 0.737 | 0.732 | 0.693 | 0.696 | 0.703 | 0.708 |
| 2238 | GG**Ptr**GG**Cys**GG | 0.762 | 0.748 | 0.754 | 0.664 | 0.680 | 0.802 | 0.751 |
| 2239 | GG**Ptr**GG**Glh**GG | 0.638 | 0.690 | 0.505 | 0.745 | 0.634 | 0.541 | 0.636 |
| 2240 | GG**Ptr**GG**Gln**GG | 0.718 | 0.767 | 0.745 | 0.746 | 0.674 | 0.716 | 0.732 |
| 2241 | GG**Ptr**GG**Glu**GG | 0.693 | 0.623 | 0.785 | 0.772 | 0.604 | 0.546 | 0.658 |
| 2242 | GG**Ptr**GG**Gly**GG | 0.876 | 0.879 | 0.888 | 0.845 | 0.838 | 0.851 | 0.864 |
| 2243 | GG**Ptr**GG**Hip**GG | 0.502 | 0.430 | 0.546 | 0.338 | 0.487 | 0.747 | 0.495 |
| 2244 | GG**Ptr**GG**His**GG | 0.763 | 0.777 | 0.731 | 0.790 | 0.795 | 0.786 | 0.782 |
| 2245 | GG**Ptr**GG**Ile**GG | 0.392 | 0.308 | 0.334 | 0.612 | 0.684 | 0.669 | 0.502 |
| 2246 | GG**Ptr**GG**Leu**GG | 0.789 | 0.714 | 0.801 | 0.728 | 0.781 | 0.710 | 0.754 |
| 2247 | GG**Ptr**GG**Lys**GG | 0.703 | 0.705 | 0.711 | 0.762 | 0.716 | 0.650 | 0.708 |
| 2248 | GG**Ptr**GG**Met**GG | 0.714 | 0.637 | 0.771 | 0.584 | 0.818 | 0.685 | 0.700 |
| 2249 | GG**Ptr**GG**Phe**GG | 0.691 | 0.652 | 0.479 | 0.729 | 0.584 | 0.754 | 0.671 |

This work is shared under a CC BY 4.0 license.

| | Peptide | $r_{replica}$(1,2) | $r_{replica}$(1,3) | $r_{replica}$(1,4) | $r_{replica}$(2,3) | $r_{replica}$(2,4) | $r_{replica}$(3,4) | Median |
|---|---|---|---|---|---|---|---|---|
| 2250 | GG**Ptr**GG**Pro**GG | 0.832 | 0.828 | 0.860 | 0.855 | 0.870 | 0.829 | 0.843 |
| 2251 | GG**Ptr**GG**Ptr**GG | 0.742 | 0.701 | 0.489 | 0.784 | 0.421 | 0.371 | 0.595 |
| 2252 | GG**Ptr**GG**S1p**GG | 0.663 | 0.646 | 0.716 | 0.576 | 0.594 | 0.644 | 0.645 |
| 2253 | GG**Ptr**GG**Sep**GG | 0.484 | 0.604 | 0.505 | 0.693 | 0.829 | 0.725 | 0.648 |
| 2254 | GG**Ptr**GG**Ser**GG | 0.718 | 0.825 | 0.852 | 0.740 | 0.704 | 0.813 | 0.776 |
| 2255 | GG**Ptr**GG**T1p**GG | 0.721 | 0.513 | 0.692 | 0.556 | 0.753 | 0.523 | 0.624 |
| 2256 | GG**Ptr**GG**Thr**GG | 0.853 | 0.811 | 0.893 | 0.759 | 0.840 | 0.798 | 0.826 |
| 2257 | GG**Ptr**GG**Tpo**GG | 0.701 | 0.375 | 0.843 | 0.810 | 0.859 | 0.699 | 0.756 |
| 2258 | GG**Ptr**GG**Trp**GG | 0.667 | 0.711 | 0.526 | 0.697 | 0.563 | 0.586 | 0.626 |
| 2259 | GG**Ptr**GG**Tyr**GG | 0.766 | 0.738 | 0.833 | 0.768 | 0.783 | 0.771 | 0.770 |
| 2260 | GG**Ptr**GG**Val**GG | 0.550 | 0.478 | 0.521 | 0.674 | 0.702 | 0.593 | 0.572 |
| 2261 | GG**Ptr**GG**Y1p**GG | 0.764 | 0.713 | 0.695 | 0.724 | 0.664 | 0.771 | 0.718 |
| 2262 | GG**S1p**GG**Ala**GG | 0.864 | 0.856 | 0.831 | 0.898 | 0.853 | 0.867 | 0.860 |
| 2263 | GG**S1p**GG**Arg**GG | 0.754 | 0.749 | 0.663 | 0.747 | 0.597 | 0.685 | 0.716 |
| 2264 | GG**S1p**GG**Ash**GG | 0.797 | 0.689 | 0.803 | 0.702 | 0.807 | 0.720 | 0.759 |
| 2265 | GG**S1p**GG**Asn**GG | 0.747 | 0.762 | 0.820 | 0.782 | 0.774 | 0.797 | 0.778 |
| 2266 | GG**S1p**GG**Asp**GG | 0.730 | 0.822 | 0.769 | 0.675 | 0.823 | 0.759 | 0.764 |
| 2267 | GG**S1p**GG**Cys**GG | 0.780 | 0.746 | 0.849 | 0.723 | 0.802 | 0.731 | 0.763 |
| 2268 | GG**S1p**GG**Glh**GG | 0.748 | 0.764 | 0.690 | 0.799 | 0.767 | 0.747 | 0.756 |
| 2269 | GG**S1p**GG**Gln**GG | 0.758 | 0.797 | 0.748 | 0.772 | 0.768 | 0.723 | 0.763 |
| 2270 | GG**S1p**GG**Glu**GG | 0.752 | 0.814 | 0.696 | 0.764 | 0.716 | 0.732 | 0.742 |
| 2271 | GG**S1p**GG**Gly**GG | 0.846 | 0.824 | 0.834 | 0.784 | 0.849 | 0.850 | 0.840 |
| 2272 | GG**S1p**GG**Hip**GG | 0.531 | 0.724 | 0.719 | 0.690 | 0.728 | 0.832 | 0.721 |
| 2273 | GG**S1p**GG**His**GG | 0.742 | 0.821 | 0.644 | 0.742 | 0.609 | 0.753 | 0.742 |
| 2274 | GG**S1p**GG**Ile**GG | 0.779 | 0.736 | 0.802 | 0.716 | 0.777 | 0.784 | 0.778 |
| 2275 | GG**S1p**GG**Leu**GG | 0.799 | 0.793 | 0.808 | 0.754 | 0.797 | 0.829 | 0.798 |
| 2276 | GG**S1p**GG**Lys**GG | 0.743 | 0.700 | 0.857 | 0.705 | 0.792 | 0.732 | 0.737 |
| 2277 | GG**S1p**GG**Met**GG | 0.864 | 0.808 | 0.815 | 0.823 | 0.811 | 0.817 | 0.816 |
| 2278 | GG**S1p**GG**Phe**GG | 0.810 | 0.674 | 0.789 | 0.669 | 0.767 | 0.649 | 0.720 |
| 2279 | GG**S1p**GG**Pro**GG | 0.817 | 0.794 | 0.794 | 0.844 | 0.823 | 0.814 | 0.816 |
| 2280 | GG**S1p**GG**Ptr**GG | 0.855 | 0.795 | 0.828 | 0.815 | 0.812 | 0.782 | 0.813 |
| 2281 | GG**S1p**GG**S1p**GG | 0.771 | 0.714 | 0.768 | 0.780 | 0.843 | 0.756 | 0.770 |
| 2282 | GG**S1p**GG**Sep**GG | 0.795 | 0.815 | 0.767 | 0.851 | 0.871 | 0.840 | 0.827 |
| 2283 | GG**S1p**GG**Ser**GG | 0.814 | 0.575 | 0.835 | 0.572 | 0.769 | 0.536 | 0.672 |
| 2284 | GG**S1p**GG**T1p**GG | 0.464 | 0.647 | 0.527 | 0.677 | 0.785 | 0.662 | 0.655 |
| 2285 | GG**S1p**GG**Thr**GG | 0.784 | 0.848 | 0.835 | 0.799 | 0.831 | 0.837 | 0.833 |
| 2286 | GG**S1p**GG**Tpo**GG | 0.829 | 0.748 | 0.505 | 0.783 | 0.553 | 0.577 | 0.663 |
| 2287 | GG**S1p**GG**Trp**GG | 0.807 | 0.761 | 0.850 | 0.777 | 0.794 | 0.746 | 0.786 |
| 2288 | GG**S1p**GG**Tyr**GG | 0.793 | 0.718 | 0.749 | 0.747 | 0.804 | 0.806 | 0.771 |
| 2289 | GG**S1p**GG**Val**GG | 0.736 | 0.773 | 0.752 | 0.749 | 0.782 | 0.790 | 0.763 |
| 2290 | GG**S1p**GG**Y1p**GG | 0.821 | 0.817 | 0.781 | 0.839 | 0.709 | 0.691 | 0.799 |
| 2291 | GG**Sep**GG**Ala**GG | 0.720 | 0.748 | 0.702 | 0.825 | 0.757 | 0.810 | 0.752 |
| 2292 | GG**Sep**GG**Arg**GG | 0.733 | 0.696 | 0.693 | 0.741 | 0.770 | 0.683 | 0.714 |
| 2293 | GG**Sep**GG**Ash**GG | 0.755 | 0.786 | 0.818 | 0.771 | 0.810 | 0.865 | 0.798 |
| 2294 | GG**Sep**GG**Asn**GG | 0.717 | 0.788 | 0.790 | 0.800 | 0.813 | 0.851 | 0.795 |
| 2295 | GG**Sep**GG**Asp**GG | 0.736 | 0.816 | 0.796 | 0.716 | 0.708 | 0.818 | 0.766 |
| 2296 | GG**Sep**GG**Cys**GG | 0.831 | 0.796 | 0.717 | 0.837 | 0.757 | 0.776 | 0.786 |
| 2297 | GG**Sep**GG**Glh**GG | 0.781 | 0.804 | 0.771 | 0.758 | 0.830 | 0.786 | 0.783 |
| 2298 | GG**Sep**GG**Gln**GG | 0.738 | 0.791 | 0.728 | 0.800 | 0.765 | 0.794 | 0.778 |
| 2299 | GG**Sep**GG**Glu**GG | 0.808 | 0.811 | 0.822 | 0.805 | 0.796 | 0.851 | 0.809 |

This work is shared under a CC BY 4.0 license.

| | Peptide | $r_{replica}$(1,2) | $r_{replica}$(1,3) | $r_{replica}$(1,4) | $r_{replica}$(2,3) | $r_{replica}$(2,4) | $r_{replica}$(3,4) | Median |
|---|---|---|---|---|---|---|---|---|
| 2300 | GG**Sep**GG**Gly**GG | 0.847 | 0.842 | 0.872 | 0.821 | 0.831 | 0.872 | 0.844 |
| 2301 | GG**Sep**GG**Hip**GG | 0.691 | 0.758 | 0.759 | 0.641 | 0.601 | 0.708 | 0.699 |
| 2302 | GG**Sep**GG**His**GG | 0.765 | 0.795 | 0.827 | 0.772 | 0.796 | 0.826 | 0.795 |
| 2303 | GG**Sep**GG**Ile**GG | 0.646 | 0.779 | 0.705 | 0.656 | 0.608 | 0.747 | 0.680 |
| 2304 | GG**Sep**GG**Leu**GG | 0.683 | 0.764 | 0.692 | 0.717 | 0.800 | 0.740 | 0.729 |
| 2305 | GG**Sep**GG**Lys**GG | 0.440 | 0.484 | 0.454 | 0.388 | 0.324 | 0.617 | 0.447 |
| 2306 | GG**Sep**GG**Met**GG | 0.843 | 0.769 | 0.820 | 0.836 | 0.841 | 0.813 | 0.828 |
| 2307 | GG**Sep**GG**Phe**GG | 0.801 | 0.800 | 0.819 | 0.765 | 0.768 | 0.749 | 0.784 |
| 2308 | GG**Sep**GG**Pro**GG | 0.779 | 0.731 | 0.801 | 0.828 | 0.809 | 0.808 | 0.804 |
| 2309 | GG**Sep**GG**Ptr**GG | 0.541 | 0.659 | 0.712 | 0.770 | 0.703 | 0.825 | 0.708 |
| 2310 | GG**Sep**GG**S1p**GG | 0.784 | 0.794 | 0.716 | 0.803 | 0.750 | 0.801 | 0.789 |
| 2311 | GG**Sep**GG**Sep**GG | 0.754 | 0.687 | 0.747 | 0.748 | 0.755 | 0.714 | 0.747 |
| 2312 | GG**Sep**GG**Ser**GG | 0.595 | 0.721 | 0.768 | 0.664 | 0.623 | 0.781 | 0.693 |
| 2313 | GG**Sep**GG**T1p**GG | 0.771 | 0.884 | 0.757 | 0.824 | 0.691 | 0.756 | 0.764 |
| 2314 | GG**Sep**GG**Thr**GG | 0.902 | 0.891 | 0.923 | 0.862 | 0.901 | 0.916 | 0.902 |
| 2315 | GG**Sep**GG**Tpo**GG | 0.132 | 0.853 | 0.136 | 0.437 | 0.967 | 0.453 | 0.445 |
| 2316 | GG**Sep**GG**Trp**GG | 0.651 | 0.713 | 0.753 | 0.739 | 0.738 | 0.740 | 0.739 |
| 2317 | GG**Sep**GG**Tyr**GG | 0.808 | 0.787 | 0.789 | 0.846 | 0.794 | 0.816 | 0.801 |
| 2318 | GG**Sep**GG**Val**GG | 0.639 | 0.764 | 0.680 | 0.631 | 0.532 | 0.643 | 0.641 |
| 2319 | GG**Sep**GG**Y1p**GG | 0.691 | 0.782 | 0.764 | 0.766 | 0.820 | 0.822 | 0.774 |
| 2320 | GG**Ser**GG**Ala**GG | 0.755 | 0.761 | 0.734 | 0.862 | 0.887 | 0.866 | 0.812 |
| 2321 | GG**Ser**GG**Arg**GG | 0.794 | 0.703 | 0.750 | 0.758 | 0.809 | 0.762 | 0.760 |
| 2322 | GG**Ser**GG**Ash**GG | 0.904 | 0.891 | 0.768 | 0.889 | 0.831 | 0.813 | 0.860 |
| 2323 | GG**Ser**GG**Asn**GG | 0.745 | 0.785 | 0.809 | 0.774 | 0.770 | 0.859 | 0.780 |
| 2324 | GG**Ser**GG**Asp**GG | 0.808 | 0.773 | 0.820 | 0.821 | 0.841 | 0.820 | 0.820 |
| 2325 | GG**Ser**GG**Cys**GG | 0.836 | 0.850 | 0.840 | 0.906 | 0.922 | 0.883 | 0.867 |
| 2326 | GG**Ser**GG**Glh**GG | 0.810 | 0.783 | 0.832 | 0.819 | 0.829 | 0.855 | 0.824 |
| 2327 | GG**Ser**GG**Gln**GG | 0.880 | 0.881 | 0.848 | 0.891 | 0.830 | 0.834 | 0.864 |
| 2328 | GG**Ser**GG**Glu**GG | 0.779 | 0.767 | 0.725 | 0.845 | 0.799 | 0.848 | 0.789 |
| 2329 | GG**Ser**GG**Gly**GG | 0.883 | 0.901 | 0.868 | 0.893 | 0.879 | 0.893 | 0.888 |
| 2330 | GG**Ser**GG**Hip**GG | 0.765 | 0.777 | 0.707 | 0.676 | 0.753 | 0.763 | 0.758 |
| 2331 | GG**Ser**GG**His**GG | 0.873 | 0.780 | 0.802 | 0.751 | 0.803 | 0.790 | 0.796 |
| 2332 | GG**Ser**GG**Ile**GG | 0.776 | 0.718 | 0.784 | 0.694 | 0.718 | 0.829 | 0.747 |
| 2333 | GG**Ser**GG**Leu**GG | 0.755 | 0.838 | 0.795 | 0.743 | 0.822 | 0.812 | 0.804 |
| 2334 | GG**Ser**GG**Lys**GG | 0.769 | 0.736 | 0.725 | 0.700 | 0.694 | 0.708 | 0.717 |
| 2335 | GG**Ser**GG**Met**GG | 0.832 | 0.783 | 0.802 | 0.791 | 0.852 | 0.767 | 0.797 |
| 2336 | GG**Ser**GG**Phe**GG | 0.881 | 0.902 | 0.827 | 0.902 | 0.874 | 0.871 | 0.878 |
| 2337 | GG**Ser**GG**Pro**GG | 0.827 | 0.779 | 0.831 | 0.843 | 0.896 | 0.844 | 0.837 |
| 2338 | GG**Ser**GG**Ptr**GG | 0.745 | 0.792 | 0.818 | 0.751 | 0.819 | 0.797 | 0.795 |
| 2339 | GG**Ser**GG**S1p**GG | 0.835 | 0.821 | 0.879 | 0.806 | 0.847 | 0.865 | 0.841 |
| 2340 | GG**Ser**GG**Sep**GG | 0.810 | 0.790 | 0.817 | 0.833 | 0.819 | 0.862 | 0.818 |
| 2341 | GG**Ser**GG**Ser**GG | 0.887 | 0.914 | 0.911 | 0.898 | 0.910 | 0.942 | 0.911 |
| 2342 | GG**Ser**GG**T1p**GG | 0.856 | 0.665 | 0.465 | 0.859 | 0.713 | 0.907 | 0.785 |
| 2343 | GG**Ser**GG**Thr**GG | 0.873 | 0.911 | 0.892 | 0.898 | 0.891 | 0.879 | 0.891 |
| 2344 | GG**Ser**GG**Tpo**GG | 0.448 | 0.406 | 0.915 | 0.924 | 0.413 | 0.372 | 0.430 |
| 2345 | GG**Ser**GG**Trp**GG | 0.745 | 0.851 | 0.893 | 0.719 | 0.731 | 0.854 | 0.798 |
| 2346 | GG**Ser**GG**Tyr**GG | 0.819 | 0.868 | 0.873 | 0.858 | 0.838 | 0.891 | 0.863 |
| 2347 | GG**Ser**GG**Val**GG | 0.834 | 0.852 | 0.819 | 0.845 | 0.806 | 0.788 | 0.826 |
| 2348 | GG**Ser**GG**Y1p**GG | 0.833 | 0.761 | 0.689 | 0.781 | 0.666 | 0.686 | 0.725 |
| 2349 | GG**T1p**GG**Ala**GG | 0.471 | 0.720 | 0.788 | 0.770 | 0.579 | 0.733 | 0.727 |

This work is shared under a CC BY 4.0 license.

| | Peptide | $r_{replica}$(1,2) | $r_{replica}$(1,3) | $r_{replica}$(1,4) | $r_{replica}$(2,3) | $r_{replica}$(2,4) | $r_{replica}$(3,4) | Median |
|---|---|---|---|---|---|---|---|---|
| 2350 | GG**T1p**GG**Arg**GG | 0.637 | 0.824 | 0.567 | 0.573 | 0.757 | 0.481 | 0.605 |
| 2351 | GG**T1p**GG**Ash**GG | 0.858 | 0.869 | 0.760 | 0.820 | 0.785 | 0.750 | 0.803 |
| 2352 | GG**T1p**GG**Asn**GG | 0.532 | 0.475 | 0.495 | 0.879 | 0.934 | 0.809 | 0.670 |
| 2353 | GG**T1p**GG**Asp**GG | 0.866 | 0.811 | 0.844 | 0.843 | 0.821 | 0.870 | 0.843 |
| 2354 | GG**T1p**GG**Cys**GG | 0.824 | 0.889 | 0.654 | 0.844 | 0.650 | 0.571 | 0.739 |
| 2355 | GG**T1p**GG**Glh**GG | 0.849 | 0.732 | 0.881 | 0.794 | 0.848 | 0.762 | 0.821 |
| 2356 | GG**T1p**GG**Gln**GG | 0.654 | 0.710 | 0.621 | 0.495 | 0.316 | 0.703 | 0.637 |
| 2357 | GG**T1p**GG**Glu**GG | 0.777 | 0.757 | 0.720 | 0.768 | 0.727 | 0.856 | 0.762 |
| 2358 | GG**T1p**GG**Gly**GG | 0.521 | 0.731 | 0.436 | 0.925 | 0.955 | 0.897 | 0.814 |
| 2359 | GG**T1p**GG**Hip**GG | 0.783 | 0.568 | 0.507 | 0.778 | 0.768 | 0.864 | 0.773 |
| 2360 | GG**T1p**GG**His**GG | 0.612 | 0.759 | 0.630 | 0.551 | 0.757 | 0.591 | 0.621 |
| 2361 | GG**T1p**GG**Ile**GG | 0.336 | 0.568 | 0.518 | 0.708 | 0.509 | 0.600 | 0.543 |
| 2362 | GG**T1p**GG**Leu**GG | 0.656 | 0.670 | 0.527 | 0.690 | 0.692 | 0.674 | 0.672 |
| 2363 | GG**T1p**GG**Lys**GG | 0.624 | 0.695 | 0.729 | 0.551 | 0.512 | 0.817 | 0.659 |
| 2364 | GG**T1p**GG**Met**GG | 0.608 | 0.372 | 0.722 | 0.791 | 0.729 | 0.632 | 0.677 |
| 2365 | GG**T1p**GG**Phe**GG | 0.601 | 0.555 | 0.402 | 0.753 | 0.664 | 0.818 | 0.632 |
| 2366 | GG**T1p**GG**Pro**GG | 0.717 | 0.825 | 0.856 | 0.649 | 0.798 | 0.850 | 0.811 |
| 2367 | GG**T1p**GG**Ptr**GG | 0.632 | 0.873 | 0.833 | 0.643 | 0.738 | 0.830 | 0.784 |
| 2368 | GG**T1p**GG**S1p**GG | 0.876 | 0.748 | 0.876 | 0.626 | 0.901 | 0.643 | 0.812 |
| 2369 | GG**T1p**GG**Sep**GG | 0.837 | 0.809 | 0.854 | 0.673 | 0.718 | 0.764 | 0.787 |
| 2370 | GG**T1p**GG**Ser**GG | 0.717 | 0.822 | 0.326 | 0.690 | 0.733 | 0.363 | 0.704 |
| 2371 | GG**T1p**GG**T1p**GG | 0.717 | 0.581 | 0.590 | 0.479 | 0.552 | 0.883 | 0.586 |
| 2372 | GG**T1p**GG**Thr**GG | 0.870 | 0.871 | 0.516 | 0.918 | 0.775 | 0.736 | 0.823 |
| 2373 | GG**T1p**GG**Tpo**GG | 0.716 | 0.661 | 0.551 | 0.689 | 0.660 | 0.426 | 0.661 |
| 2374 | GG**T1p**GG**Trp**GG | 0.744 | 0.791 | 0.422 | 0.697 | 0.446 | 0.405 | 0.571 |
| 2375 | GG**T1p**GG**Tyr**GG | 0.738 | 0.380 | 0.708 | 0.491 | 0.714 | 0.405 | 0.599 |
| 2376 | GG**T1p**GG**Val**GG | 0.812 | 0.717 | 0.702 | 0.718 | 0.702 | 0.790 | 0.717 |
| 2377 | GG**T1p**GG**Y1p**GG | 0.903 | 0.772 | 0.849 | 0.683 | 0.778 | 0.865 | 0.814 |
| 2378 | GG**Thr**GG**Ala**GG | 0.863 | 0.883 | 0.840 | 0.880 | 0.880 | 0.818 | 0.871 |
| 2379 | GG**Thr**GG**Arg**GG | 0.763 | 0.805 | 0.724 | 0.791 | 0.724 | 0.767 | 0.765 |
| 2380 | GG**Thr**GG**Ash**GG | 0.849 | 0.781 | 0.778 | 0.801 | 0.819 | 0.809 | 0.805 |
| 2381 | GG**Thr**GG**Asn**GG | 0.829 | 0.863 | 0.849 | 0.848 | 0.847 | 0.851 | 0.848 |
| 2382 | GG**Thr**GG**Asp**GG | 0.815 | 0.837 | 0.768 | 0.828 | 0.770 | 0.820 | 0.818 |
| 2383 | GG**Thr**GG**Cys**GG | 0.839 | 0.730 | 0.819 | 0.688 | 0.835 | 0.682 | 0.774 |
| 2384 | GG**Thr**GG**Glh**GG | 0.729 | 0.626 | 0.822 | 0.814 | 0.820 | 0.706 | 0.771 |
| 2385 | GG**Thr**GG**Gln**GG | 0.828 | 0.825 | 0.790 | 0.822 | 0.835 | 0.821 | 0.823 |
| 2386 | GG**Thr**GG**Glu**GG | 0.791 | 0.780 | 0.857 | 0.740 | 0.833 | 0.777 | 0.786 |
| 2387 | GG**Thr**GG**Gly**GG | 0.896 | 0.875 | 0.861 | 0.892 | 0.852 | 0.880 | 0.877 |
| 2388 | GG**Thr**GG**Hip**GG | 0.762 | 0.791 | 0.774 | 0.742 | 0.779 | 0.794 | 0.777 |
| 2389 | GG**Thr**GG**His**GG | 0.826 | 0.827 | 0.801 | 0.862 | 0.829 | 0.862 | 0.828 |
| 2390 | GG**Thr**GG**Ile**GG | 0.767 | 0.754 | 0.832 | 0.856 | 0.784 | 0.836 | 0.808 |
| 2391 | GG**Thr**GG**Leu**GG | 0.838 | 0.818 | 0.750 | 0.809 | 0.795 | 0.796 | 0.803 |
| 2392 | GG**Thr**GG**Lys**GG | 0.828 | 0.834 | 0.761 | 0.812 | 0.761 | 0.806 | 0.809 |
| 2393 | GG**Thr**GG**Met**GG | 0.849 | 0.841 | 0.837 | 0.851 | 0.804 | 0.843 | 0.842 |
| 2394 | GG**Thr**GG**Phe**GG | 0.823 | 0.742 | 0.843 | 0.715 | 0.806 | 0.779 | 0.793 |
| 2395 | GG**Thr**GG**Pro**GG | 0.847 | 0.858 | 0.852 | 0.837 | 0.855 | 0.863 | 0.854 |
| 2396 | GG**Thr**GG**Ptr**GG | 0.689 | 0.763 | 0.710 | 0.813 | 0.820 | 0.797 | 0.780 |
| 2397 | GG**Thr**GG**S1p**GG | 0.832 | 0.823 | 0.838 | 0.821 | 0.826 | 0.817 | 0.825 |
| 2398 | GG**Thr**GG**Sep**GG | 0.774 | 0.760 | 0.620 | 0.786 | 0.698 | 0.757 | 0.759 |
| 2399 | GG**Thr**GG**Ser**GG | 0.808 | 0.854 | 0.791 | 0.894 | 0.842 | 0.864 | 0.848 |

This work is shared under a CC BY 4.0 license.

| | **Peptide** | $r_{replica}(1,2)$ | $r_{replica}(1,3)$ | $r_{replica}(1,4)$ | $r_{replica}(2,3)$ | $r_{replica}(2,4)$ | $r_{replica}(3,4)$ | **Median** |
|---|---|---|---|---|---|---|---|---|
| 2400 | GG**Thr**GG**T1p**GG | 0.573 | 0.778 | 0.932 | 0.879 | 0.576 | 0.774 | 0.776 |
| 2401 | GG**Thr**GG**Thr**GG | 0.836 | 0.833 | 0.839 | 0.813 | 0.835 | 0.836 | 0.835 |
| 2402 | GG**Thr**GG**Tpo**GG | 0.561 | 0.575 | 0.448 | 0.815 | 0.835 | 0.727 | 0.651 |
| 2403 | GG**Thr**GG**Trp**GG | 0.759 | 0.815 | 0.829 | 0.758 | 0.765 | 0.839 | 0.790 |
| 2404 | GG**Thr**GG**Tyr**GG | 0.864 | 0.771 | 0.822 | 0.834 | 0.852 | 0.802 | 0.828 |
| 2405 | GG**Thr**GG**Val**GG | 0.855 | 0.818 | 0.804 | 0.818 | 0.850 | 0.792 | 0.818 |
| 2406 | GG**Thr**GG**Y1p**GG | 0.853 | 0.888 | 0.879 | 0.850 | 0.861 | 0.860 | 0.861 |
| 2407 | GG**Tpo**GG**Ala**GG | 0.564 | 0.354 | 0.927 | 0.895 | 0.388 | 0.155 | 0.476 |
| 2408 | GG**Tpo**GG**Arg**GG | 0.377 | 0.777 | 0.427 | 0.049 | 0.911 | 0.139 | 0.402 |
| 2409 | GG**Tpo**GG**Ash**GG | 0.481 | 0.411 | 0.303 | 0.558 | 0.442 | 0.754 | 0.462 |
| 2410 | GG**Tpo**GG**Asn**GG | 0.750 | 0.868 | 0.735 | 0.647 | 0.778 | 0.649 | 0.742 |
| 2411 | GG**Tpo**GG**Asp**GG | 0.536 | 0.876 | 0.277 | 0.681 | 0.842 | 0.453 | 0.609 |
| 2412 | GG**Tpo**GG**Cys**GG | 0.684 | 0.665 | 0.272 | 0.530 | 0.322 | 0.517 | 0.523 |
| 2413 | GG**Tpo**GG**Glh**GG | 0.778 | 0.468 | 0.414 | 0.741 | 0.698 | 0.877 | 0.720 |
| 2414 | GG**Tpo**GG**Gln**GG | 0.581 | 0.660 | 0.485 | 0.823 | 0.828 | 0.705 | 0.683 |
| 2415 | GG**Tpo**GG**Glu**GG | 0.726 | 0.852 | 0.754 | 0.582 | 0.525 | 0.893 | 0.740 |
| 2416 | GG**Tpo**GG**Gly**GG | 0.773 | 0.839 | 0.840 | 0.873 | 0.831 | 0.888 | 0.840 |
| 2417 | GG**Tpo**GG**Hip**GG | 0.606 | 0.739 | 0.645 | 0.600 | 0.553 | 0.644 | 0.625 |
| 2418 | GG**Tpo**GG**His**GG | 0.642 | 0.816 | 0.334 | 0.715 | 0.729 | 0.488 | 0.679 |
| 2419 | GG**Tpo**GG**Ile**GG | 0.777 | 0.791 | 0.762 | 0.624 | 0.840 | 0.572 | 0.769 |
| 2420 | GG**Tpo**GG**Leu**GG | 0.695 | 0.514 | 0.786 | 0.689 | 0.837 | 0.721 | 0.708 |
| 2421 | GG**Tpo**GG**Lys**GG | 0.720 | 0.706 | 0.629 | 0.827 | 0.694 | 0.690 | 0.700 |
| 2422 | GG**Tpo**GG**Met**GG | 0.766 | 0.708 | 0.777 | 0.779 | 0.810 | 0.632 | 0.772 |
| 2423 | GG**Tpo**GG**Phe**GG | 0.399 | 0.383 | 0.895 | 0.897 | 0.501 | 0.480 | 0.491 |
| 2424 | GG**Tpo**GG**Pro**GG | 0.781 | 0.585 | 0.940 | 0.840 | 0.688 | 0.450 | 0.734 |
| 2425 | GG**Tpo**GG**Ptr**GG | 0.698 | 0.796 | 0.855 | 0.878 | 0.665 | 0.803 | 0.799 |
| 2426 | GG**Tpo**GG**S1p**GG | 0.467 | 0.656 | 0.888 | 0.820 | 0.561 | 0.699 | 0.677 |
| 2427 | GG**Tpo**GG**Sep**GG | 0.791 | 0.744 | 0.757 | 0.529 | 0.874 | 0.452 | 0.750 |
| 2428 | GG**Tpo**GG**Ser**GG | 0.917 | 0.720 | 0.895 | 0.777 | 0.878 | 0.618 | 0.827 |
| 2429 | GG**Tpo**GG**T1p**GG | 0.396 | 0.200 | 0.611 | 0.743 | 0.591 | 0.619 | 0.601 |
| 2430 | GG**Tpo**GG**Thr**GG | 0.760 | 0.748 | 0.814 | 0.547 | 0.727 | 0.711 | 0.737 |
| 2431 | GG**Tpo**GG**Tpo**GG | 0.651 | 0.518 | 0.652 | 0.424 | 0.760 | 0.460 | 0.585 |
| 2432 | GG**Tpo**GG**Trp**GG | 0.505 | 0.815 | 0.499 | 0.483 | 0.819 | 0.471 | 0.502 |
| 2433 | GG**Tpo**GG**Tyr**GG | 0.797 | 0.745 | 0.807 | 0.710 | 0.787 | 0.704 | 0.766 |
| 2434 | GG**Tpo**GG**Val**GG | 0.765 | 0.737 | 0.773 | 0.740 | 0.816 | 0.658 | 0.752 |
| 2435 | GG**Tpo**GG**Y1p**GG | 0.632 | 0.764 | 0.847 | 0.639 | 0.448 | 0.821 | 0.701 |
| 2436 | GG**Trp**GG**Ala**GG | 0.703 | 0.772 | 0.808 | 0.849 | 0.799 | 0.827 | 0.804 |
| 2437 | GG**Trp**GG**Arg**GG | 0.731 | 0.700 | 0.823 | 0.690 | 0.674 | 0.645 | 0.695 |
| 2438 | GG**Trp**GG**Ash**GG | 0.653 | 0.823 | 0.673 | 0.612 | 0.741 | 0.590 | 0.663 |
| 2439 | GG**Trp**GG**Asn**GG | 0.800 | 0.843 | 0.825 | 0.846 | 0.825 | 0.843 | 0.834 |
| 2440 | GG**Trp**GG**Asp**GG | 0.773 | 0.786 | 0.756 | 0.786 | 0.820 | 0.833 | 0.786 |
| 2441 | GG**Trp**GG**Cys**GG | 0.726 | 0.800 | 0.755 | 0.816 | 0.786 | 0.843 | 0.793 |
| 2442 | GG**Trp**GG**Glh**GG | 0.714 | 0.715 | 0.660 | 0.797 | 0.749 | 0.819 | 0.732 |
| 2443 | GG**Trp**GG**Gln**GG | 0.765 | 0.712 | 0.779 | 0.787 | 0.835 | 0.707 | 0.772 |
| 2444 | GG**Trp**GG**Glu**GG | 0.773 | 0.731 | 0.763 | 0.752 | 0.812 | 0.652 | 0.758 |
| 2445 | GG**Trp**GG**Gly**GG | 0.771 | 0.806 | 0.813 | 0.838 | 0.875 | 0.841 | 0.826 |
| 2446 | GG**Trp**GG**Hip**GG | 0.815 | 0.717 | 0.736 | 0.686 | 0.784 | 0.675 | 0.726 |
| 2447 | GG**Trp**GG**His**GG | 0.768 | 0.786 | 0.696 | 0.829 | 0.665 | 0.602 | 0.732 |
| 2448 | GG**Trp**GG**Ile**GG | 0.708 | 0.711 | 0.786 | 0.814 | 0.684 | 0.751 | 0.731 |
| 2449 | GG**Trp**GG**Leu**GG | 0.694 | 0.684 | 0.673 | 0.575 | 0.654 | 0.692 | 0.678 |

This work is shared under a CC BY 4.0 license.

| | Peptide | $r_{replica}(1,2)$ | $r_{replica}(1,3)$ | $r_{replica}(1,4)$ | $r_{replica}(2,3)$ | $r_{replica}(2,4)$ | $r_{replica}(3,4)$ | Median |
|---|---|---|---|---|---|---|---|---|
| 2450 | GG**Trp**GG**Lys**GG | 0.743 | 0.680 | 0.619 | 0.841 | 0.666 | 0.661 | 0.673 |
| 2451 | GG**Trp**GG**Met**GG | 0.689 | 0.766 | 0.820 | 0.769 | 0.718 | 0.850 | 0.768 |
| 2452 | GG**Trp**GG**Phe**GG | 0.803 | 0.723 | 0.755 | 0.811 | 0.783 | 0.753 | 0.769 |
| 2453 | GG**Trp**GG**Pro**GG | 0.766 | 0.680 | 0.619 | 0.857 | 0.815 | 0.865 | 0.790 |
| 2454 | GG**Trp**GG**Ptr**GG | 0.887 | 0.641 | 0.825 | 0.643 | 0.835 | 0.716 | 0.770 |
| 2455 | GG**Trp**GG**S1p**GG | 0.879 | 0.792 | 0.872 | 0.823 | 0.860 | 0.825 | 0.842 |
| 2456 | GG**Trp**GG**Sep**GG | 0.818 | 0.820 | 0.678 | 0.789 | 0.679 | 0.623 | 0.734 |
| 2457 | GG**Trp**GG**Ser**GG | 0.878 | 0.844 | 0.748 | 0.869 | 0.766 | 0.818 | 0.831 |
| 2458 | GG**Trp**GG**T1p**GG | 0.911 | 0.581 | 0.863 | 0.437 | 0.764 | 0.733 | 0.749 |
| 2459 | GG**Trp**GG**Thr**GG | 0.837 | 0.737 | 0.784 | 0.771 | 0.816 | 0.737 | 0.778 |
| 2460 | GG**Trp**GG**Tpo**GG | 0.915 | 0.829 | 0.762 | 0.725 | 0.847 | 0.486 | 0.796 |
| 2461 | GG**Trp**GG**Trp**GG | 0.743 | 0.540 | 0.639 | 0.691 | 0.658 | 0.658 | 0.658 |
| 2462 | GG**Trp**GG**Tyr**GG | 0.687 | 0.695 | 0.770 | 0.741 | 0.678 | 0.706 | 0.701 |
| 2463 | GG**Trp**GG**Val**GG | 0.823 | 0.884 | 0.809 | 0.812 | 0.738 | 0.838 | 0.818 |
| 2464 | GG**Trp**GG**Y1p**GG | 0.894 | 0.911 | 0.781 | 0.903 | 0.824 | 0.746 | 0.859 |
| 2465 | GG**Tyr**GG**Ala**GG | 0.860 | 0.867 | 0.898 | 0.882 | 0.900 | 0.895 | 0.889 |
| 2466 | GG**Tyr**GG**Arg**GG | 0.811 | 0.828 | 0.760 | 0.812 | 0.711 | 0.758 | 0.786 |
| 2467 | GG**Tyr**GG**Ash**GG | 0.832 | 0.611 | 0.782 | 0.691 | 0.853 | 0.707 | 0.745 |
| 2468 | GG**Tyr**GG**Asn**GG | 0.886 | 0.883 | 0.805 | 0.880 | 0.855 | 0.788 | 0.868 |
| 2469 | GG**Tyr**GG**Asp**GG | 0.842 | 0.864 | 0.849 | 0.821 | 0.781 | 0.822 | 0.832 |
| 2470 | GG**Tyr**GG**Cys**GG | 0.795 | 0.723 | 0.835 | 0.767 | 0.870 | 0.738 | 0.781 |
| 2471 | GG**Tyr**GG**Glh**GG | 0.796 | 0.800 | 0.699 | 0.818 | 0.650 | 0.629 | 0.748 |
| 2472 | GG**Tyr**GG**Gln**GG | 0.802 | 0.727 | 0.824 | 0.654 | 0.781 | 0.764 | 0.773 |
| 2473 | GG**Tyr**GG**Glu**GG | 0.796 | 0.773 | 0.699 | 0.831 | 0.731 | 0.691 | 0.752 |
| 2474 | GG**Tyr**GG**Gly**GG | 0.884 | 0.864 | 0.875 | 0.870 | 0.858 | 0.848 | 0.867 |
| 2475 | GG**Tyr**GG**Hip**GG | 0.762 | 0.722 | 0.724 | 0.847 | 0.671 | 0.685 | 0.723 |
| 2476 | GG**Tyr**GG**His**GG | 0.802 | 0.862 | 0.813 | 0.861 | 0.821 | 0.815 | 0.818 |
| 2477 | GG**Tyr**GG**Ile**GG | 0.667 | 0.804 | 0.831 | 0.521 | 0.567 | 0.760 | 0.714 |
| 2478 | GG**Tyr**GG**Leu**GG | 0.707 | 0.694 | 0.672 | 0.793 | 0.826 | 0.861 | 0.750 |
| 2479 | GG**Tyr**GG**Lys**GG | 0.753 | 0.673 | 0.764 | 0.713 | 0.791 | 0.725 | 0.739 |
| 2480 | GG**Tyr**GG**Met**GG | 0.909 | 0.823 | 0.839 | 0.884 | 0.837 | 0.788 | 0.838 |
| 2481 | GG**Tyr**GG**Phe**GG | 0.624 | 0.601 | 0.569 | 0.824 | 0.809 | 0.860 | 0.716 |
| 2482 | GG**Tyr**GG**Pro**GG | 0.794 | 0.853 | 0.889 | 0.809 | 0.801 | 0.856 | 0.831 |
| 2483 | GG**Tyr**GG**Ptr**GG | 0.716 | 0.805 | 0.681 | 0.807 | 0.701 | 0.714 | 0.715 |
| 2484 | GG**Tyr**GG**S1p**GG | 0.751 | 0.679 | 0.588 | 0.737 | 0.679 | 0.684 | 0.682 |
| 2485 | GG**Tyr**GG**Sep**GG | 0.784 | 0.795 | 0.800 | 0.814 | 0.787 | 0.811 | 0.798 |
| 2486 | GG**Tyr**GG**Ser**GG | 0.812 | 0.831 | 0.771 | 0.776 | 0.706 | 0.832 | 0.794 |
| 2487 | GG**Tyr**GG**T1p**GG | 0.539 | 0.769 | 0.745 | 0.382 | 0.378 | 0.785 | 0.642 |
| 2488 | GG**Tyr**GG**Thr**GG | 0.734 | 0.749 | 0.781 | 0.827 | 0.776 | 0.804 | 0.779 |
| 2489 | GG**Tyr**GG**Tpo**GG | 0.627 | 0.746 | 0.458 | 0.712 | 0.777 | 0.611 | 0.670 |
| 2490 | GG**Tyr**GG**Trp**GG | 0.753 | 0.757 | 0.660 | 0.759 | 0.614 | 0.612 | 0.707 |
| 2491 | GG**Tyr**GG**Tyr**GG | 0.721 | 0.724 | 0.691 | 0.802 | 0.812 | 0.780 | 0.752 |
| 2492 | GG**Tyr**GG**Val**GG | 0.605 | 0.730 | 0.758 | 0.661 | 0.601 | 0.740 | 0.696 |
| 2493 | GG**Tyr**GG**Y1p**GG | 0.781 | 0.844 | 0.758 | 0.793 | 0.752 | 0.759 | 0.770 |
| 2494 | GG**Val**GG**Ala**GG | 0.836 | 0.819 | 0.810 | 0.857 | 0.885 | 0.875 | 0.846 |
| 2495 | GG**Val**GG**Arg**GG | 0.689 | 0.672 | 0.716 | 0.734 | 0.777 | 0.802 | 0.725 |
| 2496 | GG**Val**GG**Ash**GG | 0.810 | 0.847 | 0.859 | 0.922 | 0.869 | 0.911 | 0.864 |
| 2497 | GG**Val**GG**Asn**GG | 0.843 | 0.842 | 0.867 | 0.849 | 0.839 | 0.868 | 0.846 |
| 2498 | GG**Val**GG**Asp**GG | 0.558 | 0.713 | 0.571 | 0.568 | 0.789 | 0.553 | 0.570 |
| 2499 | GG**Val**GG**Cys**GG | 0.885 | 0.874 | 0.840 | 0.849 | 0.821 | 0.882 | 0.861 |

This work is shared under a CC BY 4.0 license.

| | Peptide | $r_{replica}(1,2)$ | $r_{replica}(1,3)$ | $r_{replica}(1,4)$ | $r_{replica}(2,3)$ | $r_{replica}(2,4)$ | $r_{replica}(3,4)$ | Median |
|---|---|---|---|---|---|---|---|---|
| 2500 | GG**Val**GG**Glh**GG | 0.884 | 0.823 | 0.747 | 0.842 | 0.744 | 0.807 | 0.815 |
| 2501 | GG**Val**GG**Gln**GG | 0.775 | 0.783 | 0.748 | 0.743 | 0.716 | 0.773 | 0.760 |
| 2502 | GG**Val**GG**Glu**GG | 0.712 | 0.723 | 0.646 | 0.773 | 0.816 | 0.785 | 0.748 |
| 2503 | GG**Val**GG**Gly**GG | 0.867 | 0.842 | 0.873 | 0.823 | 0.877 | 0.834 | 0.854 |
| 2504 | GG**Val**GG**Hip**GG | 0.651 | 0.806 | 0.740 | 0.756 | 0.761 | 0.801 | 0.758 |
| 2505 | GG**Val**GG**His**GG | 0.796 | 0.802 | 0.775 | 0.764 | 0.725 | 0.723 | 0.770 |
| 2506 | GG**Val**GG**Ile**GG | 0.726 | 0.749 | 0.790 | 0.652 | 0.833 | 0.700 | 0.737 |
| 2507 | GG**Val**GG**Leu**GG | 0.840 | 0.745 | 0.771 | 0.796 | 0.785 | 0.820 | 0.790 |
| 2508 | GG**Val**GG**Lys**GG | 0.872 | 0.802 | 0.796 | 0.806 | 0.802 | 0.794 | 0.802 |
| 2509 | GG**Val**GG**Met**GG | 0.780 | 0.700 | 0.820 | 0.688 | 0.800 | 0.699 | 0.740 |
| 2510 | GG**Val**GG**Phe**GG | 0.809 | 0.718 | 0.832 | 0.776 | 0.833 | 0.764 | 0.792 |
| 2511 | GG**Val**GG**Pro**GG | 0.833 | 0.838 | 0.852 | 0.830 | 0.813 | 0.881 | 0.836 |
| 2512 | GG**Val**GG**Ptr**GG | 0.679 | 0.774 | 0.692 | 0.796 | 0.694 | 0.752 | 0.723 |
| 2513 | GG**Val**GG**S1p**GG | 0.746 | 0.796 | 0.810 | 0.801 | 0.774 | 0.826 | 0.799 |
| 2514 | GG**Val**GG**Sep**GG | 0.741 | 0.742 | 0.806 | 0.795 | 0.837 | 0.768 | 0.781 |
| 2515 | GG**Val**GG**Ser**GG | 0.813 | 0.767 | 0.795 | 0.756 | 0.838 | 0.808 | 0.802 |
| 2516 | GG**Val**GG**T1p**GG | 0.409 | 0.781 | 0.380 | 0.360 | 0.834 | 0.421 | 0.415 |
| 2517 | GG**Val**GG**Thr**GG | 0.891 | 0.856 | 0.900 | 0.870 | 0.892 | 0.876 | 0.884 |
| 2518 | GG**Val**GG**Tpo**GG | 0.701 | 0.716 | 0.877 | 0.150 | 0.615 | 0.750 | 0.708 |
| 2519 | GG**Val**GG**Trp**GG | 0.866 | 0.826 | 0.815 | 0.826 | 0.817 | 0.808 | 0.822 |
| 2520 | GG**Val**GG**Tyr**GG | 0.745 | 0.826 | 0.829 | 0.797 | 0.746 | 0.806 | 0.801 |
| 2521 | GG**Val**GG**Val**GG | 0.749 | 0.671 | 0.702 | 0.787 | 0.781 | 0.849 | 0.765 |
| 2522 | GG**Val**GG**Y1p**GG | 0.837 | 0.791 | 0.789 | 0.846 | 0.824 | 0.777 | 0.807 |
| 2523 | GG**Y1p**GG**Ala**GG | 0.897 | 0.867 | 0.882 | 0.899 | 0.905 | 0.910 | 0.898 |
| 2524 | GG**Y1p**GG**Arg**GG | 0.597 | 0.552 | 0.657 | 0.708 | 0.750 | 0.602 | 0.629 |
| 2525 | GG**Y1p**GG**Ash**GG | 0.809 | 0.818 | 0.878 | 0.880 | 0.797 | 0.864 | 0.841 |
| 2526 | GG**Y1p**GG**Asn**GG | 0.833 | 0.835 | 0.603 | 0.848 | 0.759 | 0.653 | 0.796 |
| 2527 | GG**Y1p**GG**Asp**GG | 0.859 | 0.864 | 0.853 | 0.826 | 0.832 | 0.820 | 0.843 |
| 2528 | GG**Y1p**GG**Cys**GG | 0.784 | 0.825 | 0.806 | 0.718 | 0.798 | 0.747 | 0.791 |
| 2529 | GG**Y1p**GG**Glh**GG | 0.674 | 0.722 | 0.717 | 0.839 | 0.844 | 0.814 | 0.768 |
| 2530 | GG**Y1p**GG**Gln**GG | 0.556 | 0.728 | 0.750 | 0.459 | 0.534 | 0.792 | 0.642 |
| 2531 | GG**Y1p**GG**Glu**GG | 0.803 | 0.791 | 0.750 | 0.835 | 0.704 | 0.822 | 0.797 |
| 2532 | GG**Y1p**GG**Gly**GG | 0.900 | 0.839 | 0.851 | 0.797 | 0.846 | 0.823 | 0.842 |
| 2533 | GG**Y1p**GG**Hip**GG | 0.794 | 0.599 | 0.804 | 0.663 | 0.761 | 0.575 | 0.712 |
| 2534 | GG**Y1p**GG**His**GG | 0.574 | 0.554 | 0.482 | 0.865 | 0.736 | 0.731 | 0.652 |
| 2535 | GG**Y1p**GG**Ile**GG | 0.713 | 0.763 | 0.717 | 0.781 | 0.681 | 0.752 | 0.735 |
| 2536 | GG**Y1p**GG**Leu**GG | 0.815 | 0.801 | 0.668 | 0.827 | 0.690 | 0.670 | 0.746 |
| 2537 | GG**Y1p**GG**Lys**GG | 0.654 | 0.602 | 0.571 | 0.633 | 0.632 | 0.788 | 0.633 |
| 2538 | GG**Y1p**GG**Met**GG | 0.836 | 0.791 | 0.814 | 0.715 | 0.806 | 0.743 | 0.799 |
| 2539 | GG**Y1p**GG**Phe**GG | 0.867 | 0.792 | 0.734 | 0.841 | 0.772 | 0.741 | 0.782 |
| 2540 | GG**Y1p**GG**Pro**GG | 0.769 | 0.850 | 0.659 | 0.816 | 0.689 | 0.607 | 0.729 |
| 2541 | GG**Y1p**GG**Ptr**GG | 0.751 | 0.719 | 0.692 | 0.833 | 0.774 | 0.747 | 0.749 |
| 2542 | GG**Y1p**GG**S1p**GG | 0.757 | 0.810 | 0.754 | 0.705 | 0.765 | 0.699 | 0.756 |
| 2543 | GG**Y1p**GG**Sep**GG | 0.502 | 0.693 | 0.695 | 0.648 | 0.573 | 0.811 | 0.670 |
| 2544 | GG**Y1p**GG**Ser**GG | 0.792 | 0.834 | 0.788 | 0.766 | 0.866 | 0.743 | 0.790 |
| 2545 | GG**Y1p**GG**T1p**GG | 0.771 | 0.858 | 0.701 | 0.784 | 0.710 | 0.661 | 0.740 |
| 2546 | GG**Y1p**GG**Thr**GG | 0.817 | 0.829 | 0.894 | 0.793 | 0.826 | 0.814 | 0.821 |
| 2547 | GG**Y1p**GG**Tpo**GG | 0.299 | 0.908 | 0.800 | 0.287 | 0.535 | 0.834 | 0.668 |
| 2548 | GG**Y1p**GG**Trp**GG | 0.685 | 0.627 | 0.676 | 0.631 | 0.666 | 0.731 | 0.671 |
| 2549 | GG**Y1p**GG**Tyr**GG | 0.704 | 0.762 | 0.777 | 0.855 | 0.826 | 0.841 | 0.802 |

This work is shared under a CC BY 4.0 license.

| | Peptide | $r_{replica}(1,2)$ | $r_{replica}(1,3)$ | $r_{replica}(1,4)$ | $r_{replica}(2,3)$ | $r_{replica}(2,4)$ | $r_{replica}(3,4)$ | Median |
|---|---|---|---|---|---|---|---|---|
| 2550 | GG**Y1p**GG**Val**GG | 0.702 | 0.675 | 0.686 | 0.810 | 0.803 | 0.756 | 0.729 |
| 2551 | GG**Y1p**GG**Y1p**GG | 0.531 | 0.536 | 0.514 | 0.472 | 0.583 | 0.468 | 0.522 |
| 2552 | GG**Ala**GGG**Ala**GG | 0.909 | 0.903 | 0.908 | 0.908 | 0.883 | 0.884 | 0.905 |
| 2553 | GG**Ala**GGG**Arg**GG | 0.795 | 0.807 | 0.768 | 0.809 | 0.804 | 0.814 | 0.806 |
| 2554 | GG**Ala**GGG**Ash**GG | 0.818 | 0.873 | 0.866 | 0.875 | 0.852 | 0.881 | 0.869 |
| 2555 | GG**Ala**GGG**Asn**GG | 0.835 | 0.875 | 0.832 | 0.893 | 0.857 | 0.869 | 0.863 |
| 2556 | GG**Ala**GGG**Asp**GG | 0.881 | 0.905 | 0.878 | 0.882 | 0.902 | 0.905 | 0.892 |
| 2557 | GG**Ala**GGG**Cys**GG | 0.864 | 0.880 | 0.883 | 0.858 | 0.866 | 0.865 | 0.866 |
| 2558 | GG**Ala**GGG**Glh**GG | 0.895 | 0.855 | 0.892 | 0.859 | 0.917 | 0.867 | 0.879 |
| 2559 | GG**Ala**GGG**Gln**GG | 0.854 | 0.921 | 0.881 | 0.861 | 0.833 | 0.864 | 0.863 |
| 2560 | GG**Ala**GGG**Glu**GG | 0.851 | 0.887 | 0.878 | 0.884 | 0.905 | 0.907 | 0.885 |
| 2561 | GG**Ala**GGG**Gly**GG | 0.922 | 0.925 | 0.922 | 0.935 | 0.921 | 0.912 | 0.922 |
| 2562 | GG**Ala**GGG**Hip**GG | 0.820 | 0.862 | 0.671 | 0.881 | 0.726 | 0.744 | 0.782 |
| 2563 | GG**Ala**GGG**His**GG | 0.947 | 0.930 | 0.851 | 0.934 | 0.872 | 0.859 | 0.901 |
| 2564 | GG**Ala**GGG**Ile**GG | 0.866 | 0.858 | 0.844 | 0.853 | 0.853 | 0.816 | 0.853 |
| 2565 | GG**Ala**GGG**Leu**GG | 0.874 | 0.869 | 0.815 | 0.878 | 0.882 | 0.902 | 0.876 |
| 2566 | GG**Ala**GGG**Lys**GG | 0.883 | 0.867 | 0.878 | 0.887 | 0.884 | 0.900 | 0.883 |
| 2567 | GG**Ala**GGG**Met**GG | 0.910 | 0.899 | 0.900 | 0.925 | 0.894 | 0.886 | 0.900 |
| 2568 | GG**Ala**GGG**Phe**GG | 0.859 | 0.788 | 0.833 | 0.861 | 0.837 | 0.843 | 0.840 |
| 2569 | GG**Ala**GGG**Pro**GG | 0.878 | 0.916 | 0.892 | 0.844 | 0.871 | 0.845 | 0.874 |
| 2570 | GG**Ala**GGG**Ptr**GG | 0.834 | 0.859 | 0.896 | 0.800 | 0.857 | 0.835 | 0.846 |
| 2571 | GG**Ala**GGG**S1p**GG | 0.851 | 0.849 | 0.883 | 0.855 | 0.840 | 0.812 | 0.850 |
| 2572 | GG**Ala**GGG**Sep**GG | 0.859 | 0.809 | 0.858 | 0.852 | 0.878 | 0.859 | 0.859 |
| 2573 | GG**Ala**GGG**Ser**GG | 0.887 | 0.860 | 0.873 | 0.887 | 0.926 | 0.868 | 0.880 |
| 2574 | GG**Ala**GGG**T1p**GG | 0.680 | 0.401 | 0.442 | 0.877 | 0.884 | 0.943 | 0.778 |
| 2575 | GG**Ala**GGG**Thr**GG | 0.856 | 0.847 | 0.810 | 0.863 | 0.832 | 0.821 | 0.840 |
| 2576 | GG**Ala**GGG**Tpo**GG | 0.354 | 0.801 | 0.935 | 0.769 | 0.164 | 0.659 | 0.714 |
| 2577 | GG**Ala**GGG**Trp**GG | 0.887 | 0.830 | 0.819 | 0.826 | 0.818 | 0.746 | 0.823 |
| 2578 | GG**Ala**GGG**Tyr**GG | 0.863 | 0.876 | 0.880 | 0.860 | 0.893 | 0.875 | 0.876 |
| 2579 | GG**Ala**GGG**Val**GG | 0.759 | 0.727 | 0.763 | 0.838 | 0.812 | 0.854 | 0.788 |
| 2580 | GG**Ala**GGG**Y1p**GG | 0.778 | 0.736 | 0.701 | 0.773 | 0.726 | 0.709 | 0.731 |
| 2581 | GG**Arg**GGG**Ala**GG | 0.797 | 0.772 | 0.839 | 0.805 | 0.818 | 0.765 | 0.801 |
| 2582 | GG**Arg**GGG**Arg**GG | 0.732 | 0.786 | 0.812 | 0.792 | 0.780 | 0.827 | 0.789 |
| 2583 | GG**Arg**GGG**Ash**GG | 0.885 | 0.888 | 0.856 | 0.907 | 0.910 | 0.864 | 0.887 |
| 2584 | GG**Arg**GGG**Asn**GG | 0.849 | 0.873 | 0.856 | 0.839 | 0.847 | 0.807 | 0.848 |
| 2585 | GG**Arg**GGG**Asp**GG | 0.674 | 0.618 | 0.686 | 0.778 | 0.846 | 0.787 | 0.732 |
| 2586 | GG**Arg**GGG**Cys**GG | 0.829 | 0.853 | 0.875 | 0.878 | 0.838 | 0.795 | 0.846 |
| 2587 | GG**Arg**GGG**Glh**GG | 0.787 | 0.856 | 0.800 | 0.802 | 0.857 | 0.797 | 0.801 |
| 2588 | GG**Arg**GGG**Gln**GG | 0.745 | 0.782 | 0.716 | 0.714 | 0.688 | 0.637 | 0.715 |
| 2589 | GG**Arg**GGG**Glu**GG | 0.766 | 0.779 | 0.805 | 0.858 | 0.763 | 0.740 | 0.773 |
| 2590 | GG**Arg**GGG**Gly**GG | 0.894 | 0.909 | 0.899 | 0.917 | 0.902 | 0.909 | 0.905 |
| 2591 | GG**Arg**GGG**Hip**GG | 0.793 | 0.788 | 0.586 | 0.815 | 0.691 | 0.741 | 0.765 |
| 2592 | GG**Arg**GGG**His**GG | 0.815 | 0.843 | 0.834 | 0.849 | 0.808 | 0.845 | 0.839 |
| 2593 | GG**Arg**GGG**Ile**GG | 0.791 | 0.709 | 0.855 | 0.829 | 0.850 | 0.766 | 0.810 |
| 2594 | GG**Arg**GGG**Leu**GG | 0.865 | 0.858 | 0.813 | 0.882 | 0.783 | 0.808 | 0.836 |
| 2595 | GG**Arg**GGG**Lys**GG | 0.839 | 0.787 | 0.786 | 0.776 | 0.735 | 0.863 | 0.786 |
| 2596 | GG**Arg**GGG**Met**GG | 0.893 | 0.880 | 0.818 | 0.904 | 0.897 | 0.839 | 0.886 |
| 2597 | GG**Arg**GGG**Phe**GG | 0.784 | 0.782 | 0.623 | 0.728 | 0.504 | 0.623 | 0.676 |
| 2598 | GG**Arg**GGG**Pro**GG | 0.733 | 0.740 | 0.768 | 0.694 | 0.743 | 0.726 | 0.737 |
| 2599 | GG**Arg**GGG**Ptr**GG | 0.635 | 0.746 | 0.825 | 0.659 | 0.664 | 0.758 | 0.705 |

This work is shared under a CC BY 4.0 license.

| | Peptide | $r_{replica}(1,2)$ | $r_{replica}(1,3)$ | $r_{replica}(1,4)$ | $r_{replica}(2,3)$ | $r_{replica}(2,4)$ | $r_{replica}(3,4)$ | Median |
|---|---|---|---|---|---|---|---|---|
| 2600 | GG**Arg**GGG**S1p**GG | 0.662 | 0.811 | 0.672 | 0.726 | 0.690 | 0.705 | 0.697 |
| 2601 | GG**Arg**GGG**Sep**GG | 0.794 | 0.803 | 0.746 | 0.805 | 0.748 | 0.667 | 0.771 |
| 2602 | GG**Arg**GGG**Ser**GG | 0.784 | 0.809 | 0.800 | 0.820 | 0.867 | 0.815 | 0.812 |
| 2603 | GG**Arg**GGG**T1p**GG | 0.812 | 0.660 | 0.787 | 0.596 | 0.765 | 0.666 | 0.715 |
| 2604 | GG**Arg**GGG**Thr**GG | 0.867 | 0.853 | 0.839 | 0.832 | 0.824 | 0.824 | 0.835 |
| 2605 | GG**Arg**GGG**Tpo**GG | 0.754 | 0.191 | 0.421 | 0.346 | 0.473 | 0.844 | 0.447 |
| 2606 | GG**Arg**GGG**Trp**GG | 0.646 | 0.614 | 0.615 | 0.733 | 0.739 | 0.729 | 0.688 |
| 2607 | GG**Arg**GGG**Tyr**GG | 0.765 | 0.669 | 0.618 | 0.734 | 0.572 | 0.697 | 0.683 |
| 2608 | GG**Arg**GGG**Val**GG | 0.819 | 0.660 | 0.757 | 0.683 | 0.804 | 0.544 | 0.720 |
| 2609 | GG**Arg**GGG**Y1p**GG | 0.672 | 0.582 | 0.718 | 0.637 | 0.682 | 0.585 | 0.655 |
| 2610 | GG**Ash**GGG**Ala**GG | 0.897 | 0.901 | 0.902 | 0.866 | 0.902 | 0.897 | 0.899 |
| 2611 | GG**Ash**GGG**Arg**GG | 0.857 | 0.802 | 0.859 | 0.770 | 0.851 | 0.852 | 0.852 |
| 2612 | GG**Ash**GGG**Ash**GG | 0.843 | 0.852 | 0.811 | 0.888 | 0.864 | 0.856 | 0.854 |
| 2613 | GG**Ash**GGG**Asn**GG | 0.855 | 0.852 | 0.813 | 0.875 | 0.798 | 0.828 | 0.840 |
| 2614 | GG**Ash**GGG**Asp**GG | 0.804 | 0.755 | 0.849 | 0.650 | 0.820 | 0.659 | 0.780 |
| 2615 | GG**Ash**GGG**Cys**GG | 0.777 | 0.784 | 0.760 | 0.842 | 0.830 | 0.817 | 0.800 |
| 2616 | GG**Ash**GGG**Glh**GG | 0.831 | 0.871 | 0.818 | 0.865 | 0.886 | 0.846 | 0.855 |
| 2617 | GG**Ash**GGG**Gln**GG | 0.835 | 0.836 | 0.795 | 0.887 | 0.858 | 0.829 | 0.835 |
| 2618 | GG**Ash**GGG**Glu**GG | 0.802 | 0.824 | 0.853 | 0.751 | 0.869 | 0.830 | 0.827 |
| 2619 | GG**Ash**GGG**Gly**GG | 0.916 | 0.919 | 0.905 | 0.947 | 0.923 | 0.920 | 0.919 |
| 2620 | GG**Ash**GGG**Hip**GG | 0.706 | 0.822 | 0.705 | 0.733 | 0.687 | 0.639 | 0.706 |
| 2621 | GG**Ash**GGG**His**GG | 0.770 | 0.792 | 0.840 | 0.896 | 0.876 | 0.895 | 0.858 |
| 2622 | GG**Ash**GGG**Ile**GG | 0.688 | 0.845 | 0.857 | 0.703 | 0.704 | 0.873 | 0.774 |
| 2623 | GG**Ash**GGG**Leu**GG | 0.782 | 0.788 | 0.886 | 0.814 | 0.829 | 0.826 | 0.820 |
| 2624 | GG**Ash**GGG**Lys**GG | 0.811 | 0.817 | 0.803 | 0.862 | 0.863 | 0.870 | 0.839 |
| 2625 | GG**Ash**GGG**Met**GG | 0.854 | 0.831 | 0.878 | 0.826 | 0.872 | 0.881 | 0.863 |
| 2626 | GG**Ash**GGG**Phe**GG | 0.850 | 0.859 | 0.859 | 0.907 | 0.908 | 0.909 | 0.883 |
| 2627 | GG**Ash**GGG**Pro**GG | 0.897 | 0.911 | 0.885 | 0.912 | 0.852 | 0.899 | 0.898 |
| 2628 | GG**Ash**GGG**Ptr**GG | 0.690 | 0.836 | 0.864 | 0.768 | 0.825 | 0.847 | 0.831 |
| 2629 | GG**Ash**GGG**S1p**GG | 0.880 | 0.816 | 0.839 | 0.896 | 0.908 | 0.875 | 0.877 |
| 2630 | GG**Ash**GGG**Sep**GG | 0.911 | 0.942 | 0.795 | 0.923 | 0.832 | 0.815 | 0.871 |
| 2631 | GG**Ash**GGG**Ser**GG | 0.729 | 0.759 | 0.791 | 0.854 | 0.828 | 0.805 | 0.798 |
| 2632 | GG**Ash**GGG**T1p**GG | 0.630 | 0.817 | 0.859 | 0.305 | 0.494 | 0.853 | 0.724 |
| 2633 | GG**Ash**GGG**Thr**GG | 0.859 | 0.893 | 0.839 | 0.893 | 0.832 | 0.852 | 0.855 |
| 2634 | GG**Ash**GGG**Tpo**GG | 0.912 | 0.388 | 0.949 | 0.416 | 0.938 | 0.456 | 0.684 |
| 2635 | GG**Ash**GGG**Trp**GG | 0.683 | 0.778 | 0.830 | 0.629 | 0.706 | 0.729 | 0.718 |
| 2636 | GG**Ash**GGG**Tyr**GG | 0.802 | 0.810 | 0.780 | 0.846 | 0.835 | 0.814 | 0.812 |
| 2637 | GG**Ash**GGG**Val**GG | 0.855 | 0.886 | 0.876 | 0.815 | 0.835 | 0.830 | 0.845 |
| 2638 | GG**Ash**GGG**Y1p**GG | 0.761 | 0.855 | 0.893 | 0.862 | 0.794 | 0.887 | 0.859 |
| 2639 | GG**Asn**GGG**Ala**GG | 0.883 | 0.810 | 0.866 | 0.846 | 0.870 | 0.840 | 0.856 |
| 2640 | GG**Asn**GGG**Arg**GG | 0.847 | 0.793 | 0.786 | 0.765 | 0.727 | 0.757 | 0.776 |
| 2641 | GG**Asn**GGG**Ash**GG | 0.870 | 0.874 | 0.870 | 0.817 | 0.834 | 0.886 | 0.870 |
| 2642 | GG**Asn**GGG**Asn**GG | 0.906 | 0.878 | 0.938 | 0.862 | 0.921 | 0.853 | 0.892 |
| 2643 | GG**Asn**GGG**Asp**GG | 0.802 | 0.834 | 0.876 | 0.769 | 0.825 | 0.797 | 0.813 |
| 2644 | GG**Asn**GGG**Cys**GG | 0.885 | 0.883 | 0.862 | 0.895 | 0.884 | 0.873 | 0.884 |
| 2645 | GG**Asn**GGG**Glh**GG | 0.838 | 0.832 | 0.884 | 0.821 | 0.905 | 0.829 | 0.835 |
| 2646 | GG**Asn**GGG**Gln**GG | 0.889 | 0.879 | 0.873 | 0.872 | 0.866 | 0.840 | 0.872 |
| 2647 | GG**Asn**GGG**Glu**GG | 0.849 | 0.850 | 0.835 | 0.803 | 0.794 | 0.839 | 0.837 |
| 2648 | GG**Asn**GGG**Gly**GG | 0.933 | 0.916 | 0.944 | 0.926 | 0.931 | 0.901 | 0.929 |
| 2649 | GG**Asn**GGG**Hip**GG | 0.793 | 0.869 | 0.757 | 0.838 | 0.724 | 0.746 | 0.775 |

This work is shared under a CC BY 4.0 license.

| | Peptide | $r_{replica}(1,2)$ | $r_{replica}(1,3)$ | $r_{replica}(1,4)$ | $r_{replica}(2,3)$ | $r_{replica}(2,4)$ | $r_{replica}(3,4)$ | Median |
|---|---|---|---|---|---|---|---|---|
| 2650 | GG**Asn**GGG**His**GG | 0.819 | 0.815 | 0.823 | 0.837 | 0.839 | 0.850 | 0.830 |
| 2651 | GG**Asn**GGG**Ile**GG | 0.845 | 0.838 | 0.816 | 0.883 | 0.812 | 0.858 | 0.842 |
| 2652 | GG**Asn**GGG**Leu**GG | 0.823 | 0.823 | 0.819 | 0.810 | 0.814 | 0.898 | 0.821 |
| 2653 | GG**Asn**GGG**Lys**GG | 0.824 | 0.814 | 0.815 | 0.869 | 0.839 | 0.866 | 0.831 |
| 2654 | GG**Asn**GGG**Met**GG | 0.835 | 0.896 | 0.845 | 0.838 | 0.851 | 0.882 | 0.848 |
| 2655 | GG**Asn**GGG**Phe**GG | 0.892 | 0.895 | 0.881 | 0.873 | 0.880 | 0.883 | 0.882 |
| 2656 | GG**Asn**GGG**Pro**GG | 0.857 | 0.909 | 0.922 | 0.839 | 0.877 | 0.905 | 0.891 |
| 2657 | GG**Asn**GGG**Ptr**GG | 0.821 | 0.766 | 0.826 | 0.900 | 0.849 | 0.858 | 0.837 |
| 2658 | GG**Asn**GGG**S1p**GG | 0.868 | 0.852 | 0.848 | 0.888 | 0.880 | 0.858 | 0.863 |
| 2659 | GG**Asn**GGG**Sep**GG | 0.893 | 0.822 | 0.866 | 0.841 | 0.832 | 0.793 | 0.837 |
| 2660 | GG**Asn**GGG**Ser**GG | 0.825 | 0.878 | 0.877 | 0.835 | 0.865 | 0.879 | 0.871 |
| 2661 | GG**Asn**GGG**T1p**GG | 0.901 | 0.798 | 0.875 | 0.758 | 0.876 | 0.832 | 0.853 |
| 2662 | GG**Asn**GGG**Thr**GG | 0.897 | 0.872 | 0.885 | 0.856 | 0.862 | 0.857 | 0.867 |
| 2663 | GG**Asn**GGG**Tpo**GG | 0.562 | 0.890 | 0.742 | 0.521 | 0.796 | 0.693 | 0.718 |
| 2664 | GG**Asn**GGG**Trp**GG | 0.721 | 0.700 | 0.755 | 0.839 | 0.884 | 0.858 | 0.797 |
| 2665 | GG**Asn**GGG**Tyr**GG | 0.707 | 0.747 | 0.778 | 0.709 | 0.773 | 0.819 | 0.760 |
| 2666 | GG**Asn**GGG**Val**GG | 0.832 | 0.820 | 0.828 | 0.804 | 0.846 | 0.805 | 0.824 |
| 2667 | GG**Asn**GGG**Y1p**GG | 0.891 | 0.866 | 0.886 | 0.882 | 0.912 | 0.894 | 0.889 |
| 2668 | GG**Asp**GGG**Ala**GG | 0.846 | 0.822 | 0.802 | 0.873 | 0.842 | 0.879 | 0.844 |
| 2669 | GG**Asp**GGG**Arg**GG | 0.720 | 0.706 | 0.627 | 0.709 | 0.650 | 0.653 | 0.679 |
| 2670 | GG**Asp**GGG**Ash**GG | 0.769 | 0.830 | 0.813 | 0.747 | 0.749 | 0.836 | 0.791 |
| 2671 | GG**Asp**GGG**Asn**GG | 0.746 | 0.803 | 0.864 | 0.703 | 0.736 | 0.804 | 0.774 |
| 2672 | GG**Asp**GGG**Asp**GG | 0.715 | 0.678 | 0.761 | 0.663 | 0.879 | 0.691 | 0.703 |
| 2673 | GG**Asp**GGG**Cys**GG | 0.822 | 0.810 | 0.821 | 0.828 | 0.769 | 0.746 | 0.816 |
| 2674 | GG**Asp**GGG**Glh**GG | 0.809 | 0.778 | 0.710 | 0.740 | 0.728 | 0.777 | 0.758 |
| 2675 | GG**Asp**GGG**Gln**GG | 0.740 | 0.767 | 0.759 | 0.823 | 0.721 | 0.793 | 0.763 |
| 2676 | GG**Asp**GGG**Glu**GG | 0.785 | 0.775 | 0.798 | 0.843 | 0.803 | 0.822 | 0.800 |
| 2677 | GG**Asp**GGG**Gly**GG | 0.839 | 0.833 | 0.885 | 0.849 | 0.855 | 0.859 | 0.852 |
| 2678 | GG**Asp**GGG**Hip**GG | 0.812 | 0.828 | 0.838 | 0.782 | 0.739 | 0.831 | 0.820 |
| 2679 | GG**Asp**GGG**His**GG | 0.834 | 0.805 | 0.826 | 0.806 | 0.801 | 0.816 | 0.811 |
| 2680 | GG**Asp**GGG**Ile**GG | 0.853 | 0.825 | 0.789 | 0.804 | 0.768 | 0.823 | 0.814 |
| 2681 | GG**Asp**GGG**Leu**GG | 0.749 | 0.775 | 0.840 | 0.764 | 0.788 | 0.850 | 0.782 |
| 2682 | GG**Asp**GGG**Lys**GG | 0.767 | 0.757 | 0.755 | 0.747 | 0.698 | 0.802 | 0.756 |
| 2683 | GG**Asp**GGG**Met**GG | 0.783 | 0.762 | 0.837 | 0.753 | 0.779 | 0.796 | 0.781 |
| 2684 | GG**Asp**GGG**Phe**GG | 0.820 | 0.760 | 0.823 | 0.880 | 0.827 | 0.798 | 0.821 |
| 2685 | GG**Asp**GGG**Pro**GG | 0.830 | 0.861 | 0.784 | 0.832 | 0.833 | 0.779 | 0.831 |
| 2686 | GG**Asp**GGG**Ptr**GG | 0.837 | 0.823 | 0.740 | 0.816 | 0.751 | 0.718 | 0.784 |
| 2687 | GG**Asp**GGG**S1p**GG | 0.816 | 0.770 | 0.757 | 0.822 | 0.826 | 0.848 | 0.819 |
| 2688 | GG**Asp**GGG**Sep**GG | 0.800 | 0.834 | 0.774 | 0.846 | 0.787 | 0.831 | 0.815 |
| 2689 | GG**Asp**GGG**Ser**GG | 0.810 | 0.833 | 0.756 | 0.838 | 0.823 | 0.841 | 0.828 |
| 2690 | GG**Asp**GGG**T1p**GG | 0.830 | 0.365 | 0.330 | 0.515 | 0.484 | 0.804 | 0.499 |
| 2691 | GG**Asp**GGG**Thr**GG | 0.762 | 0.794 | 0.836 | 0.768 | 0.766 | 0.831 | 0.781 |
| 2692 | GG**Asp**GGG**Tpo**GG | 0.509 | 0.875 | 0.152 | 0.626 | 0.816 | 0.263 | 0.567 |
| 2693 | GG**Asp**GGG**Trp**GG | 0.582 | 0.834 | 0.853 | 0.638 | 0.532 | 0.812 | 0.725 |
| 2694 | GG**Asp**GGG**Tyr**GG | 0.767 | 0.738 | 0.792 | 0.700 | 0.719 | 0.770 | 0.753 |
| 2695 | GG**Asp**GGG**Val**GG | 0.774 | 0.786 | 0.850 | 0.837 | 0.861 | 0.813 | 0.825 |
| 2696 | GG**Asp**GGG**Y1p**GG | 0.780 | 0.714 | 0.597 | 0.663 | 0.548 | 0.452 | 0.630 |
| 2697 | GG**Cys**GGG**Ala**GG | 0.809 | 0.858 | 0.811 | 0.892 | 0.848 | 0.896 | 0.853 |
| 2698 | GG**Cys**GGG**Arg**GG | 0.808 | 0.795 | 0.810 | 0.811 | 0.842 | 0.851 | 0.811 |
| 2699 | GG**Cys**GGG**Ash**GG | 0.833 | 0.794 | 0.861 | 0.799 | 0.825 | 0.818 | 0.822 |

This work is shared under a CC BY 4.0 license.

| | Peptide | $r_{replica}(1,2)$ | $r_{replica}(1,3)$ | $r_{replica}(1,4)$ | $r_{replica}(2,3)$ | $r_{replica}(2,4)$ | $r_{replica}(3,4)$ | Median |
|---|---|---|---|---|---|---|---|---|
| 2700 | GG**Cys**GGG**Asn**GG | 0.838 | 0.875 | 0.844 | 0.887 | 0.850 | 0.863 | 0.856 |
| 2701 | GG**Cys**GGG**Asp**GG | 0.881 | 0.891 | 0.865 | 0.875 | 0.869 | 0.884 | 0.878 |
| 2702 | GG**Cys**GGG**Cys**GG | 0.892 | 0.889 | 0.851 | 0.886 | 0.859 | 0.845 | 0.873 |
| 2703 | GG**Cys**GGG**Glh**GG | 0.751 | 0.748 | 0.771 | 0.803 | 0.834 | 0.800 | 0.786 |
| 2704 | GG**Cys**GGG**Gln**GG | 0.857 | 0.839 | 0.846 | 0.829 | 0.795 | 0.816 | 0.834 |
| 2705 | GG**Cys**GGG**Glu**GG | 0.863 | 0.841 | 0.846 | 0.776 | 0.861 | 0.812 | 0.843 |
| 2706 | GG**Cys**GGG**Gly**GG | 0.888 | 0.898 | 0.877 | 0.899 | 0.913 | 0.902 | 0.898 |
| 2707 | GG**Cys**GGG**Hip**GG | 0.641 | 0.678 | 0.704 | 0.790 | 0.826 | 0.828 | 0.747 |
| 2708 | GG**Cys**GGG**His**GG | 0.870 | 0.823 | 0.770 | 0.831 | 0.780 | 0.730 | 0.802 |
| 2709 | GG**Cys**GGG**Ile**GG | 0.761 | 0.798 | 0.777 | 0.794 | 0.768 | 0.825 | 0.785 |
| 2710 | GG**Cys**GGG**Leu**GG | 0.839 | 0.784 | 0.843 | 0.819 | 0.863 | 0.788 | 0.829 |
| 2711 | GG**Cys**GGG**Lys**GG | 0.804 | 0.880 | 0.861 | 0.789 | 0.837 | 0.876 | 0.849 |
| 2712 | GG**Cys**GGG**Met**GG | 0.813 | 0.829 | 0.857 | 0.860 | 0.856 | 0.864 | 0.856 |
| 2713 | GG**Cys**GGG**Phe**GG | 0.784 | 0.784 | 0.843 | 0.835 | 0.816 | 0.849 | 0.825 |
| 2714 | GG**Cys**GGG**Pro**GG | 0.915 | 0.889 | 0.901 | 0.862 | 0.906 | 0.885 | 0.895 |
| 2715 | GG**Cys**GGG**Ptr**GG | 0.743 | 0.692 | 0.806 | 0.703 | 0.722 | 0.633 | 0.713 |
| 2716 | GG**Cys**GGG**S1p**GG | 0.890 | 0.870 | 0.879 | 0.890 | 0.879 | 0.840 | 0.879 |
| 2717 | GG**Cys**GGG**Sep**GG | 0.813 | 0.761 | 0.808 | 0.789 | 0.848 | 0.746 | 0.799 |
| 2718 | GG**Cys**GGG**Ser**GG | 0.903 | 0.867 | 0.871 | 0.881 | 0.893 | 0.910 | 0.887 |
| 2719 | GG**Cys**GGG**T1p**GG | 0.795 | 0.841 | 0.849 | 0.807 | 0.804 | 0.840 | 0.824 |
| 2720 | GG**Cys**GGG**Thr**GG | 0.811 | 0.843 | 0.862 | 0.804 | 0.865 | 0.820 | 0.831 |
| 2721 | GG**Cys**GGG**Tpo**GG | 0.504 | 0.676 | 0.626 | 0.754 | 0.788 | 0.689 | 0.682 |
| 2722 | GG**Cys**GGG**Trp**GG | 0.802 | 0.887 | 0.825 | 0.785 | 0.859 | 0.817 | 0.821 |
| 2723 | GG**Cys**GGG**Tyr**GG | 0.800 | 0.807 | 0.816 | 0.831 | 0.877 | 0.832 | 0.824 |
| 2724 | GG**Cys**GGG**Val**GG | 0.839 | 0.796 | 0.735 | 0.800 | 0.775 | 0.742 | 0.786 |
| 2725 | GG**Cys**GGG**Y1p**GG | 0.847 | 0.839 | 0.788 | 0.788 | 0.834 | 0.740 | 0.811 |
| 2726 | GG**Glh**GGG**Ala**GG | 0.918 | 0.857 | 0.877 | 0.880 | 0.909 | 0.898 | 0.889 |
| 2727 | GG**Glh**GGG**Arg**GG | 0.769 | 0.832 | 0.794 | 0.792 | 0.704 | 0.781 | 0.786 |
| 2728 | GG**Glh**GGG**Ash**GG | 0.837 | 0.829 | 0.824 | 0.878 | 0.873 | 0.889 | 0.855 |
| 2729 | GG**Glh**GGG**Asn**GG | 0.760 | 0.812 | 0.852 | 0.830 | 0.789 | 0.810 | 0.811 |
| 2730 | GG**Glh**GGG**Asp**GG | 0.707 | 0.750 | 0.775 | 0.837 | 0.714 | 0.792 | 0.763 |
| 2731 | GG**Glh**GGG**Cys**GG | 0.897 | 0.867 | 0.873 | 0.883 | 0.889 | 0.906 | 0.886 |
| 2732 | GG**Glh**GGG**Glh**GG | 0.914 | 0.894 | 0.859 | 0.888 | 0.833 | 0.871 | 0.879 |
| 2733 | GG**Glh**GGG**Gln**GG | 0.591 | 0.720 | 0.726 | 0.839 | 0.817 | 0.909 | 0.771 |
| 2734 | GG**Glh**GGG**Glu**GG | 0.843 | 0.815 | 0.849 | 0.856 | 0.766 | 0.780 | 0.829 |
| 2735 | GG**Glh**GGG**Gly**GG | 0.869 | 0.900 | 0.904 | 0.861 | 0.887 | 0.878 | 0.883 |
| 2736 | GG**Glh**GGG**Hip**GG | 0.792 | 0.705 | 0.789 | 0.694 | 0.777 | 0.685 | 0.741 |
| 2737 | GG**Glh**GGG**His**GG | 0.873 | 0.920 | 0.895 | 0.865 | 0.817 | 0.862 | 0.869 |
| 2738 | GG**Glh**GGG**Ile**GG | 0.812 | 0.860 | 0.840 | 0.862 | 0.787 | 0.827 | 0.834 |
| 2739 | GG**Glh**GGG**Leu**GG | 0.867 | 0.873 | 0.459 | 0.869 | 0.463 | 0.483 | 0.675 |
| 2740 | GG**Glh**GGG**Lys**GG | 0.816 | 0.751 | 0.839 | 0.856 | 0.737 | 0.678 | 0.783 |
| 2741 | GG**Glh**GGG**Met**GG | 0.843 | 0.875 | 0.884 | 0.875 | 0.895 | 0.876 | 0.876 |
| 2742 | GG**Glh**GGG**Phe**GG | 0.711 | 0.892 | 0.817 | 0.758 | 0.808 | 0.836 | 0.812 |
| 2743 | GG**Glh**GGG**Pro**GG | 0.878 | 0.802 | 0.835 | 0.836 | 0.871 | 0.858 | 0.847 |
| 2744 | GG**Glh**GGG**Ptr**GG | 0.827 | 0.783 | 0.736 | 0.698 | 0.646 | 0.586 | 0.717 |
| 2745 | GG**Glh**GGG**S1p**GG | 0.823 | 0.856 | 0.874 | 0.870 | 0.875 | 0.891 | 0.872 |
| 2746 | GG**Glh**GGG**Sep**GG | 0.796 | 0.774 | 0.773 | 0.817 | 0.807 | 0.854 | 0.801 |
| 2747 | GG**Glh**GGG**Ser**GG | 0.790 | 0.810 | 0.772 | 0.854 | 0.837 | 0.852 | 0.823 |
| 2748 | GG**Glh**GGG**T1p**GG | 0.767 | 0.754 | 0.688 | 0.866 | 0.785 | 0.850 | 0.776 |
| 2749 | GG**Glh**GGG**Thr**GG | 0.830 | 0.786 | 0.765 | 0.799 | 0.804 | 0.823 | 0.802 |

This work is shared under a CC BY 4.0 license.

| | Peptide | $r_{replica}(1,2)$ | $r_{replica}(1,3)$ | $r_{replica}(1,4)$ | $r_{replica}(2,3)$ | $r_{replica}(2,4)$ | $r_{replica}(3,4)$ | Median |
|---|---|---|---|---|---|---|---|---|
| 2750 | GG**Glh**GGG**Tpo**GG | 0.873 | 0.810 | 0.763 | 0.790 | 0.842 | 0.767 | 0.800 |
| 2751 | GG**Glh**GGG**Trp**GG | 0.787 | 0.797 | 0.707 | 0.797 | 0.754 | 0.792 | 0.790 |
| 2752 | GG**Glh**GGG**Tyr**GG | 0.804 | 0.819 | 0.751 | 0.804 | 0.873 | 0.821 | 0.812 |
| 2753 | GG**Glh**GGG**Val**GG | 0.844 | 0.871 | 0.861 | 0.834 | 0.858 | 0.882 | 0.860 |
| 2754 | GG**Glh**GGG**Y1p**GG | 0.805 | 0.762 | 0.900 | 0.831 | 0.842 | 0.770 | 0.818 |
| 2755 | GG**Gln**GGG**Ala**GG | 0.803 | 0.869 | 0.863 | 0.856 | 0.805 | 0.828 | 0.842 |
| 2756 | GG**Gln**GGG**Arg**GG | 0.797 | 0.764 | 0.811 | 0.764 | 0.736 | 0.773 | 0.768 |
| 2757 | GG**Gln**GGG**Ash**GG | 0.854 | 0.829 | 0.834 | 0.873 | 0.871 | 0.863 | 0.858 |
| 2758 | GG**Gln**GGG**Asn**GG | 0.895 | 0.858 | 0.885 | 0.851 | 0.906 | 0.854 | 0.872 |
| 2759 | GG**Gln**GGG**Asp**GG | 0.847 | 0.866 | 0.835 | 0.876 | 0.889 | 0.870 | 0.868 |
| 2760 | GG**Gln**GGG**Cys**GG | 0.813 | 0.859 | 0.861 | 0.840 | 0.826 | 0.849 | 0.844 |
| 2761 | GG**Gln**GGG**Glh**GG | 0.893 | 0.933 | 0.852 | 0.900 | 0.893 | 0.862 | 0.893 |
| 2762 | GG**Gln**GGG**Gln**GG | 0.761 | 0.801 | 0.782 | 0.843 | 0.852 | 0.873 | 0.822 |
| 2763 | GG**Gln**GGG**Glu**GG | 0.850 | 0.882 | 0.847 | 0.838 | 0.860 | 0.830 | 0.848 |
| 2764 | GG**Gln**GGG**Gly**GG | 0.930 | 0.929 | 0.928 | 0.921 | 0.938 | 0.920 | 0.929 |
| 2765 | GG**Gln**GGG**Hip**GG | 0.802 | 0.753 | 0.734 | 0.829 | 0.777 | 0.808 | 0.790 |
| 2766 | GG**Gln**GGG**His**GG | 0.831 | 0.845 | 0.798 | 0.889 | 0.850 | 0.854 | 0.847 |
| 2767 | GG**Gln**GGG**Ile**GG | 0.807 | 0.826 | 0.855 | 0.733 | 0.801 | 0.838 | 0.816 |
| 2768 | GG**Gln**GGG**Leu**GG | 0.796 | 0.841 | 0.814 | 0.834 | 0.834 | 0.799 | 0.824 |
| 2769 | GG**Gln**GGG**Lys**GG | 0.873 | 0.834 | 0.879 | 0.852 | 0.891 | 0.860 | 0.866 |
| 2770 | GG**Gln**GGG**Met**GG | 0.828 | 0.918 | 0.830 | 0.843 | 0.834 | 0.866 | 0.838 |
| 2771 | GG**Gln**GGG**Phe**GG | 0.907 | 0.853 | 0.890 | 0.857 | 0.900 | 0.849 | 0.874 |
| 2772 | GG**Gln**GGG**Pro**GG | 0.855 | 0.871 | 0.877 | 0.880 | 0.868 | 0.846 | 0.869 |
| 2773 | GG**Gln**GGG**Ptr**GG | 0.815 | 0.848 | 0.826 | 0.840 | 0.816 | 0.874 | 0.833 |
| 2774 | GG**Gln**GGG**S1p**GG | 0.722 | 0.726 | 0.772 | 0.876 | 0.771 | 0.792 | 0.772 |
| 2775 | GG**Gln**GGG**Sep**GG | 0.679 | 0.845 | 0.837 | 0.643 | 0.661 | 0.859 | 0.758 |
| 2776 | GG**Gln**GGG**Ser**GG | 0.826 | 0.834 | 0.816 | 0.776 | 0.790 | 0.808 | 0.812 |
| 2777 | GG**Gln**GGG**T1p**GG | 0.867 | 0.837 | 0.819 | 0.881 | 0.791 | 0.772 | 0.828 |
| 2778 | GG**Gln**GGG**Thr**GG | 0.910 | 0.903 | 0.880 | 0.898 | 0.894 | 0.865 | 0.896 |
| 2779 | GG**Gln**GGG**Tpo**GG | 0.806 | 0.329 | 0.779 | 0.608 | 0.748 | 0.531 | 0.678 |
| 2780 | GG**Gln**GGG**Trp**GG | 0.716 | 0.554 | 0.716 | 0.525 | 0.788 | 0.511 | 0.635 |
| 2781 | GG**Gln**GGG**Tyr**GG | 0.734 | 0.804 | 0.755 | 0.728 | 0.778 | 0.791 | 0.767 |
| 2782 | GG**Gln**GGG**Val**GG | 0.787 | 0.816 | 0.798 | 0.769 | 0.785 | 0.824 | 0.792 |
| 2783 | GG**Gln**GGG**Y1p**GG | 0.705 | 0.785 | 0.850 | 0.755 | 0.749 | 0.858 | 0.770 |
| 2784 | GG**Glu**GGG**Ala**GG | 0.856 | 0.878 | 0.875 | 0.911 | 0.884 | 0.879 | 0.878 |
| 2785 | GG**Glu**GGG**Arg**GG | 0.777 | 0.719 | 0.708 | 0.761 | 0.684 | 0.758 | 0.738 |
| 2786 | GG**Glu**GGG**Ash**GG | 0.871 | 0.872 | 0.836 | 0.864 | 0.835 | 0.880 | 0.867 |
| 2787 | GG**Glu**GGG**Asn**GG | 0.806 | 0.823 | 0.799 | 0.779 | 0.830 | 0.854 | 0.814 |
| 2788 | GG**Glu**GGG**Asp**GG | 0.850 | 0.839 | 0.864 | 0.853 | 0.878 | 0.856 | 0.854 |
| 2789 | GG**Glu**GGG**Cys**GG | 0.835 | 0.840 | 0.803 | 0.829 | 0.815 | 0.813 | 0.822 |
| 2790 | GG**Glu**GGG**Glh**GG | 0.881 | 0.551 | 0.902 | 0.500 | 0.872 | 0.569 | 0.720 |
| 2791 | GG**Glu**GGG**Gln**GG | 0.793 | 0.714 | 0.780 | 0.801 | 0.830 | 0.804 | 0.797 |
| 2792 | GG**Glu**GGG**Glu**GG | 0.844 | 0.848 | 0.799 | 0.808 | 0.789 | 0.823 | 0.815 |
| 2793 | GG**Glu**GGG**Gly**GG | 0.922 | 0.887 | 0.867 | 0.931 | 0.883 | 0.854 | 0.885 |
| 2794 | GG**Glu**GGG**Hip**GG | 0.542 | 0.634 | 0.616 | 0.738 | 0.749 | 0.786 | 0.686 |
| 2795 | GG**Glu**GGG**His**GG | 0.854 | 0.821 | 0.851 | 0.759 | 0.806 | 0.761 | 0.813 |
| 2796 | GG**Glu**GGG**Ile**GG | 0.785 | 0.719 | 0.741 | 0.638 | 0.630 | 0.608 | 0.679 |
| 2797 | GG**Glu**GGG**Leu**GG | 0.809 | 0.809 | 0.809 | 0.819 | 0.771 | 0.774 | 0.809 |
| 2798 | GG**Glu**GGG**Lys**GG | 0.856 | 0.827 | 0.827 | 0.824 | 0.819 | 0.855 | 0.827 |
| 2799 | GG**Glu**GGG**Met**GG | 0.697 | 0.684 | 0.724 | 0.772 | 0.813 | 0.776 | 0.748 |

This work is shared under a CC BY 4.0 license.

| | Peptide | $r_{replica}$(1,2) | $r_{replica}$(1,3) | $r_{replica}$(1,4) | $r_{replica}$(2,3) | $r_{replica}$(2,4) | $r_{replica}$(3,4) | Median |
|---|---|---|---|---|---|---|---|---|
| 2800 | GG**Glu**GGG**Phe**GG | 0.822 | 0.781 | 0.815 | 0.767 | 0.808 | 0.842 | 0.812 |
| 2801 | GG**Glu**GGG**Pro**GG | 0.888 | 0.857 | 0.862 | 0.888 | 0.879 | 0.873 | 0.876 |
| 2802 | GG**Glu**GGG**Ptr**GG | 0.600 | 0.766 | 0.820 | 0.776 | 0.671 | 0.821 | 0.771 |
| 2803 | GG**Glu**GGG**S1p**GG | 0.807 | 0.859 | 0.852 | 0.838 | 0.805 | 0.854 | 0.845 |
| 2804 | GG**Glu**GGG**Sep**GG | 0.837 | 0.824 | 0.742 | 0.805 | 0.699 | 0.774 | 0.790 |
| 2805 | GG**Glu**GGG**Ser**GG | 0.846 | 0.823 | 0.790 | 0.813 | 0.800 | 0.784 | 0.807 |
| 2806 | GG**Glu**GGG**T1p**GG | 0.819 | 0.772 | 0.794 | 0.695 | 0.787 | 0.810 | 0.791 |
| 2807 | GG**Glu**GGG**Thr**GG | 0.790 | 0.821 | 0.802 | 0.875 | 0.819 | 0.846 | 0.820 |
| 2808 | GG**Glu**GGG**Tpo**GG | 0.795 | 0.782 | 0.808 | 0.823 | 0.775 | 0.804 | 0.799 |
| 2809 | GG**Glu**GGG**Trp**GG | 0.764 | 0.768 | 0.676 | 0.738 | 0.632 | 0.672 | 0.707 |
| 2810 | GG**Glu**GGG**Tyr**GG | 0.831 | 0.833 | 0.800 | 0.846 | 0.778 | 0.848 | 0.832 |
| 2811 | GG**Glu**GGG**Val**GG | 0.783 | 0.796 | 0.809 | 0.789 | 0.814 | 0.823 | 0.803 |
| 2812 | GG**Glu**GGG**Y1p**GG | 0.847 | 0.874 | 0.494 | 0.835 | 0.496 | 0.484 | 0.665 |
| 2813 | GG**Gly**GGG**Ala**GG | 0.918 | 0.932 | 0.932 | 0.907 | 0.924 | 0.927 | 0.926 |
| 2814 | GG**Gly**GGG**Arg**GG | 0.828 | 0.835 | 0.821 | 0.816 | 0.776 | 0.795 | 0.818 |
| 2815 | GG**Gly**GGG**Ash**GG | 0.890 | 0.839 | 0.908 | 0.906 | 0.915 | 0.889 | 0.898 |
| 2816 | GG**Gly**GGG**Asn**GG | 0.897 | 0.884 | 0.924 | 0.872 | 0.899 | 0.902 | 0.898 |
| 2817 | GG**Gly**GGG**Asp**GG | 0.910 | 0.945 | 0.924 | 0.934 | 0.898 | 0.940 | 0.929 |
| 2818 | GG**Gly**GGG**Cys**GG | 0.919 | 0.897 | 0.937 | 0.901 | 0.901 | 0.894 | 0.901 |
| 2819 | GG**Gly**GGG**Glh**GG | 0.929 | 0.911 | 0.921 | 0.928 | 0.891 | 0.883 | 0.916 |
| 2820 | GG**Gly**GGG**Gln**GG | 0.894 | 0.886 | 0.873 | 0.886 | 0.895 | 0.926 | 0.890 |
| 2821 | GG**Gly**GGG**Glu**GG | 0.878 | 0.901 | 0.902 | 0.840 | 0.894 | 0.883 | 0.889 |
| 2822 | GG**Gly**GGG**Gly**GG | 0.943 | 0.942 | 0.945 | 0.927 | 0.952 | 0.928 | 0.943 |
| 2823 | GG**Gly**GGG**Hip**GG | 0.864 | 0.852 | 0.820 | 0.858 | 0.819 | 0.793 | 0.836 |
| 2824 | GG**Gly**GGG**His**GG | 0.883 | 0.855 | 0.859 | 0.909 | 0.855 | 0.864 | 0.861 |
| 2825 | GG**Gly**GGG**Ile**GG | 0.867 | 0.842 | 0.818 | 0.791 | 0.889 | 0.721 | 0.830 |
| 2826 | GG**Gly**GGG**Leu**GG | 0.931 | 0.894 | 0.925 | 0.906 | 0.914 | 0.940 | 0.919 |
| 2827 | GG**Gly**GGG**Lys**GG | 0.854 | 0.840 | 0.843 | 0.868 | 0.847 | 0.794 | 0.845 |
| 2828 | GG**Gly**GGG**Met**GG | 0.943 | 0.926 | 0.938 | 0.939 | 0.954 | 0.925 | 0.939 |
| 2829 | GG**Gly**GGG**Phe**GG | 0.877 | 0.887 | 0.878 | 0.875 | 0.921 | 0.843 | 0.877 |
| 2830 | GG**Gly**GGG**Pro**GG | 0.904 | 0.900 | 0.902 | 0.906 | 0.933 | 0.917 | 0.905 |
| 2831 | GG**Gly**GGG**Ptr**GG | 0.796 | 0.892 | 0.906 | 0.856 | 0.868 | 0.905 | 0.880 |
| 2832 | GG**Gly**GGG**S1p**GG | 0.905 | 0.938 | 0.925 | 0.925 | 0.921 | 0.932 | 0.925 |
| 2833 | GG**Gly**GGG**Sep**GG | 0.909 | 0.907 | 0.910 | 0.914 | 0.891 | 0.885 | 0.908 |
| 2834 | GG**Gly**GGG**Ser**GG | 0.923 | 0.898 | 0.900 | 0.910 | 0.912 | 0.886 | 0.905 |
| 2835 | GG**Gly**GGG**T1p**GG | 0.954 | 0.818 | 0.930 | 0.786 | 0.933 | 0.811 | 0.874 |
| 2836 | GG**Gly**GGG**Thr**GG | 0.913 | 0.885 | 0.917 | 0.909 | 0.885 | 0.872 | 0.897 |
| 2837 | GG**Gly**GGG**Tpo**GG | 0.935 | 0.528 | 0.770 | 0.384 | 0.659 | 0.872 | 0.715 |
| 2838 | GG**Gly**GGG**Trp**GG | 0.859 | 0.875 | 0.855 | 0.825 | 0.838 | 0.822 | 0.847 |
| 2839 | GG**Gly**GGG**Tyr**GG | 0.791 | 0.818 | 0.764 | 0.816 | 0.816 | 0.782 | 0.803 |
| 2840 | GG**Gly**GGG**Val**GG | 0.889 | 0.900 | 0.902 | 0.871 | 0.879 | 0.878 | 0.884 |
| 2841 | GG**Gly**GGG**Y1p**GG | 0.849 | 0.824 | 0.797 | 0.801 | 0.755 | 0.861 | 0.813 |
| 2842 | GG**Hip**GGG**Ala**GG | 0.796 | 0.706 | 0.869 | 0.838 | 0.816 | 0.740 | 0.806 |
| 2843 | GG**Hip**GGG**Arg**GG | 0.778 | 0.809 | 0.813 | 0.748 | 0.824 | 0.813 | 0.811 |
| 2844 | GG**Hip**GGG**Ash**GG | 0.806 | 0.858 | 0.856 | 0.857 | 0.855 | 0.863 | 0.856 |
| 2845 | GG**Hip**GGG**Asn**GG | 0.825 | 0.823 | 0.806 | 0.787 | 0.791 | 0.841 | 0.815 |
| 2846 | GG**Hip**GGG**Asp**GG | 0.840 | 0.428 | 0.802 | 0.440 | 0.847 | 0.552 | 0.677 |
| 2847 | GG**Hip**GGG**Cys**GG | 0.859 | 0.761 | 0.857 | 0.793 | 0.846 | 0.830 | 0.838 |
| 2848 | GG**Hip**GGG**Glh**GG | 0.864 | 0.858 | 0.877 | 0.831 | 0.869 | 0.850 | 0.861 |
| 2849 | GG**Hip**GGG**Gln**GG | 0.809 | 0.816 | 0.818 | 0.863 | 0.860 | 0.815 | 0.817 |

This work is shared under a CC BY 4.0 license.

| | Peptide | $r_{replica}$(1,2) | $r_{replica}$(1,3) | $r_{replica}$(1,4) | $r_{replica}$(2,3) | $r_{replica}$(2,4) | $r_{replica}$(3,4) | Median |
|---|---|---|---|---|---|---|---|---|
| 2850 | GG**Hip**GGG**Glu**GG | 0.867 | 0.789 | 0.862 | 0.781 | 0.866 | 0.825 | 0.844 |
| 2851 | GG**Hip**GGG**Gly**GG | 0.861 | 0.894 | 0.901 | 0.866 | 0.874 | 0.887 | 0.880 |
| 2852 | GG**Hip**GGG**Hip**GG | 0.848 | 0.875 | 0.891 | 0.874 | 0.848 | 0.867 | 0.870 |
| 2853 | GG**Hip**GGG**His**GG | 0.848 | 0.805 | 0.824 | 0.850 | 0.860 | 0.831 | 0.839 |
| 2854 | GG**Hip**GGG**Ile**GG | 0.758 | 0.754 | 0.731 | 0.764 | 0.717 | 0.787 | 0.756 |
| 2855 | GG**Hip**GGG**Leu**GG | 0.848 | 0.782 | 0.781 | 0.823 | 0.758 | 0.622 | 0.782 |
| 2856 | GG**Hip**GGG**Lys**GG | 0.748 | 0.769 | 0.867 | 0.852 | 0.812 | 0.803 | 0.808 |
| 2857 | GG**Hip**GGG**Met**GG | 0.778 | 0.796 | 0.727 | 0.821 | 0.794 | 0.847 | 0.795 |
| 2858 | GG**Hip**GGG**Phe**GG | 0.741 | 0.841 | 0.816 | 0.760 | 0.707 | 0.884 | 0.788 |
| 2859 | GG**Hip**GGG**Pro**GG | 0.845 | 0.824 | 0.830 | 0.795 | 0.819 | 0.784 | 0.821 |
| 2860 | GG**Hip**GGG**Ptr**GG | 0.396 | 0.813 | 0.628 | 0.523 | 0.497 | 0.729 | 0.576 |
| 2861 | GG**Hip**GGG**S1p**GG | 0.733 | 0.772 | 0.785 | 0.825 | 0.829 | 0.842 | 0.805 |
| 2862 | GG**Hip**GGG**Sep**GG | 0.652 | 0.715 | 0.780 | 0.767 | 0.619 | 0.716 | 0.715 |
| 2863 | GG**Hip**GGG**Ser**GG | 0.784 | 0.856 | 0.725 | 0.739 | 0.784 | 0.682 | 0.762 |
| 2864 | GG**Hip**GGG**T1p**GG | 0.874 | 0.550 | 0.736 | 0.737 | 0.861 | 0.878 | 0.799 |
| 2865 | GG**Hip**GGG**Thr**GG | 0.738 | 0.791 | 0.792 | 0.785 | 0.758 | 0.837 | 0.788 |
| 2866 | GG**Hip**GGG**Tpo**GG | 0.859 | 0.649 | 0.831 | 0.636 | 0.847 | 0.484 | 0.740 |
| 2867 | GG**Hip**GGG**Trp**GG | 0.819 | 0.783 | 0.573 | 0.788 | 0.607 | 0.766 | 0.775 |
| 2868 | GG**Hip**GGG**Tyr**GG | 0.833 | 0.681 | 0.698 | 0.699 | 0.702 | 0.777 | 0.701 |
| 2869 | GG**Hip**GGG**Val**GG | 0.802 | 0.831 | 0.761 | 0.813 | 0.664 | 0.721 | 0.781 |
| 2870 | GG**Hip**GGG**Y1p**GG | 0.534 | 0.560 | 0.614 | 0.530 | 0.623 | 0.488 | 0.547 |
| 2871 | GG**His**GGG**Ala**GG | 0.767 | 0.792 | 0.823 | 0.814 | 0.810 | 0.813 | 0.811 |
| 2872 | GG**His**GGG**Arg**GG | 0.675 | 0.750 | 0.614 | 0.663 | 0.572 | 0.678 | 0.669 |
| 2873 | GG**His**GGG**Ash**GG | 0.612 | 0.841 | 0.818 | 0.497 | 0.552 | 0.785 | 0.698 |
| 2874 | GG**His**GGG**Asn**GG | 0.793 | 0.715 | 0.793 | 0.836 | 0.794 | 0.752 | 0.793 |
| 2875 | GG**His**GGG**Asp**GG | 0.742 | 0.621 | 0.713 | 0.809 | 0.863 | 0.834 | 0.775 |
| 2876 | GG**His**GGG**Cys**GG | 0.745 | 0.767 | 0.661 | 0.763 | 0.569 | 0.582 | 0.703 |
| 2877 | GG**His**GGG**Glh**GG | 0.841 | 0.834 | 0.784 | 0.799 | 0.782 | 0.799 | 0.799 |
| 2878 | GG**His**GGG**Gln**GG | 0.758 | 0.555 | 0.712 | 0.595 | 0.761 | 0.659 | 0.686 |
| 2879 | GG**His**GGG**Glu**GG | 0.843 | 0.781 | 0.820 | 0.789 | 0.829 | 0.841 | 0.824 |
| 2880 | GG**His**GGG**Gly**GG | 0.866 | 0.836 | 0.855 | 0.814 | 0.852 | 0.869 | 0.854 |
| 2881 | GG**His**GGG**Hip**GG | 0.766 | 0.788 | 0.704 | 0.758 | 0.694 | 0.603 | 0.731 |
| 2882 | GG**His**GGG**His**GG | 0.792 | 0.834 | 0.808 | 0.795 | 0.784 | 0.788 | 0.793 |
| 2883 | GG**His**GGG**Ile**GG | 0.675 | 0.719 | 0.781 | 0.831 | 0.722 | 0.773 | 0.748 |
| 2884 | GG**His**GGG**Leu**GG | 0.752 | 0.724 | 0.658 | 0.775 | 0.739 | 0.809 | 0.746 |
| 2885 | GG**His**GGG**Lys**GG | 0.791 | 0.726 | 0.751 | 0.864 | 0.786 | 0.798 | 0.789 |
| 2886 | GG**His**GGG**Met**GG | 0.861 | 0.847 | 0.790 | 0.829 | 0.832 | 0.738 | 0.831 |
| 2887 | GG**His**GGG**Phe**GG | 0.849 | 0.721 | 0.811 | 0.709 | 0.882 | 0.700 | 0.766 |
| 2888 | GG**His**GGG**Pro**GG | 0.809 | 0.853 | 0.816 | 0.901 | 0.820 | 0.860 | 0.836 |
| 2889 | GG**His**GGG**Ptr**GG | 0.619 | 0.650 | 0.657 | 0.712 | 0.767 | 0.741 | 0.684 |
| 2890 | GG**His**GGG**S1p**GG | 0.668 | 0.642 | 0.723 | 0.509 | 0.757 | 0.526 | 0.655 |
| 2891 | GG**His**GGG**Sep**GG | 0.645 | 0.634 | 0.682 | 0.654 | 0.670 | 0.770 | 0.662 |
| 2892 | GG**His**GGG**Ser**GG | 0.857 | 0.803 | 0.755 | 0.819 | 0.741 | 0.790 | 0.797 |
| 2893 | GG**His**GGG**T1p**GG | 0.739 | 0.619 | 0.578 | 0.471 | 0.521 | 0.662 | 0.599 |
| 2894 | GG**His**GGG**Thr**GG | 0.724 | 0.770 | 0.817 | 0.680 | 0.748 | 0.815 | 0.759 |
| 2895 | GG**His**GGG**Tpo**GG | 0.871 | 0.757 | 0.503 | 0.812 | 0.376 | 0.217 | 0.630 |
| 2896 | GG**His**GGG**Trp**GG | 0.722 | 0.728 | 0.818 | 0.814 | 0.690 | 0.727 | 0.727 |
| 2897 | GG**His**GGG**Tyr**GG | 0.815 | 0.752 | 0.830 | 0.751 | 0.839 | 0.796 | 0.805 |
| 2898 | GG**His**GGG**Val**GG | 0.832 | 0.796 | 0.838 | 0.741 | 0.857 | 0.801 | 0.817 |
| 2899 | GG**His**GGG**Y1p**GG | 0.441 | 0.742 | 0.766 | 0.427 | 0.452 | 0.619 | 0.535 |

This work is shared under a CC BY 4.0 license.

| | Peptide | $r_{replica}(1,2)$ | $r_{replica}(1,3)$ | $r_{replica}(1,4)$ | $r_{replica}(2,3)$ | $r_{replica}(2,4)$ | $r_{replica}(3,4)$ | Median |
|---|---|---|---|---|---|---|---|---|
| 2900 | GG**Ile**GGG**Ala**GG | 0.900 | 0.895 | 0.886 | 0.871 | 0.843 | 0.910 | 0.891 |
| 2901 | GG**Ile**GGG**Arg**GG | 0.774 | 0.785 | 0.746 | 0.815 | 0.732 | 0.756 | 0.765 |
| 2902 | GG**Ile**GGG**Ash**GG | 0.875 | 0.841 | 0.824 | 0.859 | 0.831 | 0.863 | 0.850 |
| 2903 | GG**Ile**GGG**Asn**GG | 0.776 | 0.828 | 0.779 | 0.834 | 0.844 | 0.856 | 0.831 |
| 2904 | GG**Ile**GGG**Asp**GG | 0.751 | 0.801 | 0.826 | 0.760 | 0.737 | 0.812 | 0.780 |
| 2905 | GG**Ile**GGG**Cys**GG | 0.838 | 0.826 | 0.740 | 0.791 | 0.771 | 0.772 | 0.782 |
| 2906 | GG**Ile**GGG**Glh**GG | 0.743 | 0.737 | 0.772 | 0.820 | 0.801 | 0.792 | 0.782 |
| 2907 | GG**Ile**GGG**Gln**GG | 0.743 | 0.749 | 0.762 | 0.816 | 0.852 | 0.801 | 0.782 |
| 2908 | GG**Ile**GGG**Glu**GG | 0.827 | 0.803 | 0.793 | 0.785 | 0.785 | 0.811 | 0.798 |
| 2909 | GG**Ile**GGG**Gly**GG | 0.892 | 0.833 | 0.880 | 0.852 | 0.900 | 0.860 | 0.870 |
| 2910 | GG**Ile**GGG**Hip**GG | 0.788 | 0.768 | 0.748 | 0.768 | 0.761 | 0.736 | 0.764 |
| 2911 | GG**Ile**GGG**His**GG | 0.811 | 0.880 | 0.805 | 0.842 | 0.833 | 0.825 | 0.829 |
| 2912 | GG**Ile**GGG**Ile**GG | 0.801 | 0.739 | 0.849 | 0.783 | 0.757 | 0.744 | 0.770 |
| 2913 | GG**Ile**GGG**Leu**GG | 0.758 | 0.805 | 0.806 | 0.762 | 0.754 | 0.838 | 0.784 |
| 2914 | GG**Ile**GGG**Lys**GG | 0.727 | 0.762 | 0.710 | 0.779 | 0.790 | 0.786 | 0.770 |
| 2915 | GG**Ile**GGG**Met**GG | 0.817 | 0.847 | 0.822 | 0.805 | 0.768 | 0.750 | 0.811 |
| 2916 | GG**Ile**GGG**Phe**GG | 0.814 | 0.763 | 0.830 | 0.735 | 0.787 | 0.784 | 0.786 |
| 2917 | GG**Ile**GGG**Pro**GG | 0.848 | 0.889 | 0.866 | 0.879 | 0.866 | 0.841 | 0.866 |
| 2918 | GG**Ile**GGG**Ptr**GG | 0.858 | 0.821 | 0.812 | 0.813 | 0.838 | 0.772 | 0.817 |
| 2919 | GG**Ile**GGG**S1p**GG | 0.778 | 0.754 | 0.749 | 0.797 | 0.779 | 0.711 | 0.766 |
| 2920 | GG**Ile**GGG**Sep**GG | 0.806 | 0.869 | 0.829 | 0.823 | 0.857 | 0.842 | 0.835 |
| 2921 | GG**Ile**GGG**Ser**GG | 0.802 | 0.811 | 0.839 | 0.803 | 0.855 | 0.837 | 0.824 |
| 2922 | GG**Ile**GGG**T1p**GG | 0.819 | 0.836 | 0.552 | 0.704 | 0.748 | 0.335 | 0.726 |
| 2923 | GG**Ile**GGG**Thr**GG | 0.721 | 0.775 | 0.766 | 0.852 | 0.783 | 0.821 | 0.779 |
| 2924 | GG**Ile**GGG**Tpo**GG | 0.858 | 0.715 | 0.754 | 0.780 | 0.757 | 0.805 | 0.769 |
| 2925 | GG**Ile**GGG**Trp**GG | 0.808 | 0.745 | 0.785 | 0.745 | 0.814 | 0.872 | 0.796 |
| 2926 | GG**Ile**GGG**Tyr**GG | 0.795 | 0.825 | 0.766 | 0.839 | 0.822 | 0.869 | 0.824 |
| 2927 | GG**Ile**GGG**Val**GG | 0.837 | 0.853 | 0.851 | 0.881 | 0.839 | 0.835 | 0.845 |
| 2928 | GG**Ile**GGG**Y1p**GG | 0.813 | 0.819 | 0.789 | 0.809 | 0.837 | 0.792 | 0.811 |
| 2929 | GG**Leu**GGG**Ala**GG | 0.869 | 0.866 | 0.887 | 0.810 | 0.854 | 0.851 | 0.860 |
| 2930 | GG**Leu**GGG**Arg**GG | 0.751 | 0.771 | 0.688 | 0.746 | 0.749 | 0.750 | 0.750 |
| 2931 | GG**Leu**GGG**Ash**GG | 0.907 | 0.896 | 0.860 | 0.893 | 0.927 | 0.865 | 0.894 |
| 2932 | GG**Leu**GGG**Asn**GG | 0.874 | 0.890 | 0.806 | 0.877 | 0.803 | 0.802 | 0.840 |
| 2933 | GG**Leu**GGG**Asp**GG | 0.841 | 0.887 | 0.888 | 0.859 | 0.842 | 0.867 | 0.863 |
| 2934 | GG**Leu**GGG**Cys**GG | 0.877 | 0.778 | 0.866 | 0.831 | 0.863 | 0.837 | 0.850 |
| 2935 | GG**Leu**GGG**Glh**GG | 0.881 | 0.842 | 0.827 | 0.853 | 0.802 | 0.820 | 0.835 |
| 2936 | GG**Leu**GGG**Gln**GG | 0.819 | 0.787 | 0.818 | 0.872 | 0.834 | 0.809 | 0.818 |
| 2937 | GG**Leu**GGG**Glu**GG | 0.896 | 0.869 | 0.854 | 0.882 | 0.845 | 0.874 | 0.871 |
| 2938 | GG**Leu**GGG**Gly**GG | 0.930 | 0.923 | 0.910 | 0.946 | 0.941 | 0.928 | 0.929 |
| 2939 | GG**Leu**GGG**Hip**GG | 0.758 | 0.798 | 0.742 | 0.764 | 0.688 | 0.756 | 0.757 |
| 2940 | GG**Leu**GGG**His**GG | 0.861 | 0.828 | 0.858 | 0.792 | 0.866 | 0.807 | 0.843 |
| 2941 | GG**Leu**GGG**Ile**GG | 0.736 | 0.784 | 0.648 | 0.779 | 0.690 | 0.724 | 0.730 |
| 2942 | GG**Leu**GGG**Leu**GG | 0.783 | 0.768 | 0.864 | 0.802 | 0.760 | 0.795 | 0.789 |
| 2943 | GG**Leu**GGG**Lys**GG | 0.765 | 0.806 | 0.764 | 0.741 | 0.759 | 0.767 | 0.764 |
| 2944 | GG**Leu**GGG**Met**GG | 0.844 | 0.840 | 0.850 | 0.808 | 0.830 | 0.854 | 0.842 |
| 2945 | GG**Leu**GGG**Phe**GG | 0.898 | 0.864 | 0.895 | 0.859 | 0.901 | 0.858 | 0.879 |
| 2946 | GG**Leu**GGG**Pro**GG | 0.882 | 0.922 | 0.899 | 0.891 | 0.888 | 0.917 | 0.895 |
| 2947 | GG**Leu**GGG**Ptr**GG | 0.827 | 0.866 | 0.808 | 0.798 | 0.801 | 0.808 | 0.808 |
| 2948 | GG**Leu**GGG**S1p**GG | 0.769 | 0.818 | 0.823 | 0.765 | 0.777 | 0.802 | 0.790 |
| 2949 | GG**Leu**GGG**Sep**GG | 0.834 | 0.835 | 0.895 | 0.777 | 0.811 | 0.883 | 0.834 |

This work is shared under a CC BY 4.0 license.

| | Peptide | $r_{replica}(1,2)$ | $r_{replica}(1,3)$ | $r_{replica}(1,4)$ | $r_{replica}(2,3)$ | $r_{replica}(2,4)$ | $r_{replica}(3,4)$ | Median |
|---|---|---|---|---|---|---|---|---|
| 2950 | GG**Leu**GGG**Ser**GG | 0.865 | 0.873 | 0.892 | 0.869 | 0.871 | 0.883 | 0.872 |
| 2951 | GG**Leu**GGG**T1p**GG | 0.703 | 0.822 | 0.689 | 0.658 | 0.851 | 0.629 | 0.696 |
| 2952 | GG**Leu**GGG**Thr**GG | 0.771 | 0.828 | 0.834 | 0.816 | 0.780 | 0.859 | 0.822 |
| 2953 | GG**Leu**GGG**Tpo**GG | 0.803 | 0.834 | 0.236 | 0.877 | 0.578 | 0.587 | 0.695 |
| 2954 | GG**Leu**GGG**Trp**GG | 0.558 | 0.572 | 0.576 | 0.811 | 0.680 | 0.765 | 0.628 |
| 2955 | GG**Leu**GGG**Tyr**GG | 0.698 | 0.775 | 0.664 | 0.793 | 0.802 | 0.791 | 0.783 |
| 2956 | GG**Leu**GGG**Val**GG | 0.816 | 0.841 | 0.845 | 0.844 | 0.829 | 0.905 | 0.842 |
| 2957 | GG**Leu**GGG**Y1p**GG | 0.850 | 0.796 | 0.802 | 0.833 | 0.816 | 0.785 | 0.809 |
| 2958 | GG**Lys**GGG**Ala**GG | 0.838 | 0.852 | 0.874 | 0.851 | 0.880 | 0.880 | 0.863 |
| 2959 | GG**Lys**GGG**Arg**GG | 0.788 | 0.791 | 0.806 | 0.787 | 0.749 | 0.773 | 0.787 |
| 2960 | GG**Lys**GGG**Ash**GG | 0.835 | 0.903 | 0.829 | 0.865 | 0.851 | 0.850 | 0.850 |
| 2961 | GG**Lys**GGG**Asn**GG | 0.925 | 0.804 | 0.909 | 0.786 | 0.900 | 0.837 | 0.869 |
| 2962 | GG**Lys**GGG**Asp**GG | 0.889 | 0.884 | 0.855 | 0.899 | 0.877 | 0.885 | 0.885 |
| 2963 | GG**Lys**GGG**Cys**GG | 0.727 | 0.847 | 0.823 | 0.800 | 0.758 | 0.856 | 0.812 |
| 2964 | GG**Lys**GGG**Glh**GG | 0.826 | 0.744 | 0.828 | 0.744 | 0.812 | 0.706 | 0.778 |
| 2965 | GG**Lys**GGG**Gln**GG | 0.770 | 0.800 | 0.798 | 0.867 | 0.813 | 0.854 | 0.807 |
| 2966 | GG**Lys**GGG**Glu**GG | 0.852 | 0.778 | 0.818 | 0.783 | 0.816 | 0.751 | 0.799 |
| 2967 | GG**Lys**GGG**Gly**GG | 0.891 | 0.929 | 0.877 | 0.892 | 0.905 | 0.889 | 0.891 |
| 2968 | GG**Lys**GGG**Hip**GG | 0.789 | 0.816 | 0.827 | 0.855 | 0.807 | 0.822 | 0.819 |
| 2969 | GG**Lys**GGG**His**GG | 0.826 | 0.831 | 0.772 | 0.796 | 0.823 | 0.754 | 0.809 |
| 2970 | GG**Lys**GGG**Ile**GG | 0.798 | 0.790 | 0.766 | 0.823 | 0.845 | 0.796 | 0.797 |
| 2971 | GG**Lys**GGG**Leu**GG | 0.786 | 0.824 | 0.831 | 0.789 | 0.830 | 0.819 | 0.821 |
| 2972 | GG**Lys**GGG**Lys**GG | 0.784 | 0.792 | 0.815 | 0.787 | 0.824 | 0.778 | 0.790 |
| 2973 | GG**Lys**GGG**Met**GG | 0.811 | 0.836 | 0.850 | 0.814 | 0.781 | 0.840 | 0.825 |
| 2974 | GG**Lys**GGG**Phe**GG | 0.801 | 0.803 | 0.850 | 0.843 | 0.854 | 0.841 | 0.842 |
| 2975 | GG**Lys**GGG**Pro**GG | 0.890 | 0.892 | 0.825 | 0.890 | 0.787 | 0.825 | 0.858 |
| 2976 | GG**Lys**GGG**Ptr**GG | 0.818 | 0.748 | 0.698 | 0.767 | 0.822 | 0.650 | 0.758 |
| 2977 | GG**Lys**GGG**S1p**GG | 0.756 | 0.834 | 0.836 | 0.728 | 0.716 | 0.873 | 0.795 |
| 2978 | GG**Lys**GGG**Sep**GG | 0.839 | 0.811 | 0.776 | 0.803 | 0.758 | 0.740 | 0.790 |
| 2979 | GG**Lys**GGG**Ser**GG | 0.877 | 0.858 | 0.885 | 0.837 | 0.853 | 0.850 | 0.855 |
| 2980 | GG**Lys**GGG**T1p**GG | 0.828 | 0.806 | 0.634 | 0.692 | 0.757 | 0.313 | 0.725 |
| 2981 | GG**Lys**GGG**Thr**GG | 0.859 | 0.828 | 0.862 | 0.878 | 0.866 | 0.872 | 0.864 |
| 2982 | GG**Lys**GGG**Tpo**GG | 0.884 | 0.737 | 0.243 | 0.728 | 0.121 | 0.575 | 0.651 |
| 2983 | GG**Lys**GGG**Trp**GG | 0.840 | 0.773 | 0.799 | 0.813 | 0.684 | 0.665 | 0.786 |
| 2984 | GG**Lys**GGG**Tyr**GG | 0.919 | 0.733 | 0.810 | 0.694 | 0.808 | 0.775 | 0.792 |
| 2985 | GG**Lys**GGG**Val**GG | 0.858 | 0.848 | 0.813 | 0.872 | 0.832 | 0.862 | 0.853 |
| 2986 | GG**Lys**GGG**Y1p**GG | 0.689 | 0.789 | 0.814 | 0.739 | 0.741 | 0.764 | 0.753 |
| 2987 | GG**Met**GGG**Ala**GG | 0.835 | 0.855 | 0.851 | 0.846 | 0.839 | 0.883 | 0.849 |
| 2988 | GG**Met**GGG**Arg**GG | 0.748 | 0.806 | 0.807 | 0.779 | 0.756 | 0.830 | 0.792 |
| 2989 | GG**Met**GGG**Ash**GG | 0.917 | 0.868 | 0.875 | 0.867 | 0.881 | 0.853 | 0.872 |
| 2990 | GG**Met**GGG**Asn**GG | 0.814 | 0.863 | 0.861 | 0.782 | 0.793 | 0.841 | 0.828 |
| 2991 | GG**Met**GGG**Asp**GG | 0.863 | 0.862 | 0.873 | 0.914 | 0.914 | 0.891 | 0.882 |
| 2992 | GG**Met**GGG**Cys**GG | 0.888 | 0.847 | 0.864 | 0.826 | 0.829 | 0.857 | 0.852 |
| 2993 | GG**Met**GGG**Glh**GG | 0.796 | 0.820 | 0.811 | 0.808 | 0.783 | 0.801 | 0.804 |
| 2994 | GG**Met**GGG**Gln**GG | 0.799 | 0.829 | 0.883 | 0.849 | 0.828 | 0.852 | 0.839 |
| 2995 | GG**Met**GGG**Glu**GG | 0.772 | 0.758 | 0.732 | 0.815 | 0.694 | 0.820 | 0.765 |
| 2996 | GG**Met**GGG**Gly**GG | 0.926 | 0.934 | 0.916 | 0.904 | 0.929 | 0.905 | 0.921 |
| 2997 | GG**Met**GGG**Hip**GG | 0.835 | 0.815 | 0.852 | 0.769 | 0.825 | 0.794 | 0.820 |
| 2998 | GG**Met**GGG**His**GG | 0.830 | 0.823 | 0.779 | 0.862 | 0.769 | 0.838 | 0.827 |
| 2999 | GG**Met**GGG**Ile**GG | 0.735 | 0.905 | 0.840 | 0.725 | 0.762 | 0.832 | 0.797 |

This work is shared under a CC BY 4.0 license.

| Peptide | $r_{replica}(1,2)$ | $r_{replica}(1,3)$ | $r_{replica}(1,4)$ | $r_{replica}(2,3)$ | $r_{replica}(2,4)$ | $r_{replica}(3,4)$ | Median | |
|---|---|---|---|---|---|---|---|---|
| 3000 | GG**Met**GGG**Leu**GG | 0.885 | 0.820 | 0.729 | 0.871 | 0.785 | 0.713 | 0.802 |
| 3001 | GG**Met**GGG**Lys**GG | 0.813 | 0.800 | 0.840 | 0.826 | 0.870 | 0.847 | 0.833 |
| 3002 | GG**Met**GGG**Met**GG | 0.898 | 0.875 | 0.876 | 0.853 | 0.876 | 0.871 | 0.875 |
| 3003 | GG**Met**GGG**Phe**GG | 0.885 | 0.837 | 0.842 | 0.812 | 0.828 | 0.827 | 0.832 |
| 3004 | GG**Met**GGG**Pro**GG | 0.858 | 0.860 | 0.880 | 0.802 | 0.831 | 0.880 | 0.859 |
| 3005 | GG**Met**GGG**Ptr**GG | 0.869 | 0.805 | 0.782 | 0.851 | 0.757 | 0.609 | 0.793 |
| 3006 | GG**Met**GGG**S1p**GG | 0.897 | 0.881 | 0.851 | 0.874 | 0.874 | 0.859 | 0.874 |
| 3007 | GG**Met**GGG**Sep**GG | 0.818 | 0.799 | 0.875 | 0.742 | 0.836 | 0.800 | 0.809 |
| 3008 | GG**Met**GGG**Ser**GG | 0.883 | 0.843 | 0.909 | 0.871 | 0.864 | 0.802 | 0.867 |
| 3009 | GG**Met**GGG**T1p**GG | 0.849 | 0.634 | 0.795 | 0.645 | 0.799 | 0.351 | 0.720 |
| 3010 | GG**Met**GGG**Thr**GG | 0.859 | 0.867 | 0.833 | 0.894 | 0.839 | 0.870 | 0.863 |
| 3011 | GG**Met**GGG**Tpo**GG | 0.417 | 0.351 | 0.859 | 0.914 | 0.580 | 0.467 | 0.523 |
| 3012 | GG**Met**GGG**Trp**GG | 0.738 | 0.668 | 0.555 | 0.837 | 0.695 | 0.750 | 0.716 |
| 3013 | GG**Met**GGG**Tyr**GG | 0.781 | 0.724 | 0.725 | 0.882 | 0.799 | 0.854 | 0.790 |
| 3014 | GG**Met**GGG**Val**GG | 0.796 | 0.798 | 0.718 | 0.794 | 0.771 | 0.809 | 0.795 |
| 3015 | GG**Met**GGG**Y1p**GG | 0.740 | 0.736 | 0.737 | 0.731 | 0.826 | 0.770 | 0.738 |
| 3016 | GG**Phe**GGG**Ala**GG | 0.870 | 0.879 | 0.834 | 0.858 | 0.823 | 0.859 | 0.858 |
| 3017 | GG**Phe**GGG**Arg**GG | 0.819 | 0.828 | 0.712 | 0.784 | 0.789 | 0.704 | 0.787 |
| 3018 | GG**Phe**GGG**Ash**GG | 0.828 | 0.843 | 0.824 | 0.768 | 0.833 | 0.803 | 0.826 |
| 3019 | GG**Phe**GGG**Asn**GG | 0.864 | 0.848 | 0.798 | 0.833 | 0.803 | 0.811 | 0.822 |
| 3020 | GG**Phe**GGG**Asp**GG | 0.756 | 0.727 | 0.829 | 0.720 | 0.788 | 0.751 | 0.754 |
| 3021 | GG**Phe**GGG**Cys**GG | 0.841 | 0.842 | 0.879 | 0.840 | 0.841 | 0.827 | 0.841 |
| 3022 | GG**Phe**GGG**Glh**GG | 0.794 | 0.846 | 0.817 | 0.915 | 0.830 | 0.871 | 0.838 |
| 3023 | GG**Phe**GGG**Gln**GG | 0.820 | 0.840 | 0.777 | 0.835 | 0.794 | 0.823 | 0.821 |
| 3024 | GG**Phe**GGG**Glu**GG | 0.783 | 0.906 | 0.794 | 0.813 | 0.795 | 0.813 | 0.804 |
| 3025 | GG**Phe**GGG**Gly**GG | 0.917 | 0.894 | 0.903 | 0.878 | 0.910 | 0.889 | 0.898 |
| 3026 | GG**Phe**GGG**Hip**GG | 0.675 | 0.745 | 0.670 | 0.788 | 0.802 | 0.767 | 0.756 |
| 3027 | GG**Phe**GGG**His**GG | 0.858 | 0.886 | 0.899 | 0.836 | 0.828 | 0.845 | 0.851 |
| 3028 | GG**Phe**GGG**Ile**GG | 0.760 | 0.791 | 0.783 | 0.806 | 0.816 | 0.769 | 0.787 |
| 3029 | GG**Phe**GGG**Leu**GG | 0.855 | 0.776 | 0.877 | 0.822 | 0.871 | 0.805 | 0.838 |
| 3030 | GG**Phe**GGG**Lys**GG | 0.790 | 0.804 | 0.809 | 0.845 | 0.855 | 0.854 | 0.827 |
| 3031 | GG**Phe**GGG**Met**GG | 0.844 | 0.801 | 0.859 | 0.816 | 0.874 | 0.821 | 0.832 |
| 3032 | GG**Phe**GGG**Phe**GG | 0.847 | 0.879 | 0.877 | 0.864 | 0.867 | 0.864 | 0.866 |
| 3033 | GG**Phe**GGG**Pro**GG | 0.887 | 0.847 | 0.859 | 0.857 | 0.903 | 0.866 | 0.863 |
| 3034 | GG**Phe**GGG**Ptr**GG | 0.823 | 0.772 | 0.880 | 0.720 | 0.827 | 0.751 | 0.797 |
| 3035 | GG**Phe**GGG**S1p**GG | 0.790 | 0.804 | 0.795 | 0.829 | 0.747 | 0.842 | 0.800 |
| 3036 | GG**Phe**GGG**Sep**GG | 0.743 | 0.776 | 0.748 | 0.867 | 0.866 | 0.874 | 0.821 |
| 3037 | GG**Phe**GGG**Ser**GG | 0.848 | 0.835 | 0.816 | 0.862 | 0.827 | 0.841 | 0.838 |
| 3038 | GG**Phe**GGG**T1p**GG | 0.736 | 0.406 | 0.587 | 0.746 | 0.845 | 0.815 | 0.741 |
| 3039 | GG**Phe**GGG**Thr**GG | 0.864 | 0.849 | 0.893 | 0.863 | 0.873 | 0.852 | 0.864 |
| 3040 | GG**Phe**GGG**Tpo**GG | 0.845 | 0.778 | 0.877 | 0.480 | 0.923 | 0.584 | 0.812 |
| 3041 | GG**Phe**GGG**Trp**GG | 0.853 | 0.778 | 0.750 | 0.805 | 0.830 | 0.787 | 0.796 |
| 3042 | GG**Phe**GGG**Tyr**GG | 0.867 | 0.790 | 0.838 | 0.823 | 0.828 | 0.756 | 0.826 |
| 3043 | GG**Phe**GGG**Val**GG | 0.771 | 0.877 | 0.862 | 0.778 | 0.753 | 0.842 | 0.810 |
| 3044 | GG**Phe**GGG**Y1p**GG | 0.821 | 0.755 | 0.788 | 0.802 | 0.810 | 0.781 | 0.795 |
| 3045 | GG**Pro**GGG**Ala**GG | 0.843 | 0.867 | 0.888 | 0.820 | 0.843 | 0.886 | 0.855 |
| 3046 | GG**Pro**GGG**Arg**GG | 0.841 | 0.843 | 0.834 | 0.844 | 0.858 | 0.802 | 0.842 |
| 3047 | GG**Pro**GGG**Ash**GG | 0.905 | 0.819 | 0.772 | 0.832 | 0.788 | 0.811 | 0.815 |
| 3048 | GG**Pro**GGG**Asn**GG | 0.845 | 0.845 | 0.904 | 0.838 | 0.855 | 0.857 | 0.850 |
| 3049 | GG**Pro**GGG**Asp**GG | 0.908 | 0.886 | 0.854 | 0.915 | 0.886 | 0.863 | 0.886 |

This work is shared under a CC BY 4.0 license.

| | **Peptide** | $r_{replica}(1,2)$ | $r_{replica}(1,3)$ | $r_{replica}(1,4)$ | $r_{replica}(2,3)$ | $r_{replica}(2,4)$ | $r_{replica}(3,4)$ | **Median** |
|---|---|---|---|---|---|---|---|---|
| 3050 | GG**Pro**GGG**Cys**GG | 0.820 | 0.841 | 0.852 | 0.864 | 0.838 | 0.904 | 0.846 |
| 3051 | GG**Pro**GGG**Glh**GG | 0.493 | 0.509 | 0.472 | 0.871 | 0.662 | 0.693 | 0.585 |
| 3052 | GG**Pro**GGG**Gln**GG | 0.809 | 0.850 | 0.839 | 0.840 | 0.792 | 0.838 | 0.838 |
| 3053 | GG**Pro**GGG**Glu**GG | 0.759 | 0.806 | 0.882 | 0.843 | 0.747 | 0.823 | 0.815 |
| 3054 | GG**Pro**GGG**Gly**GG | 0.919 | 0.935 | 0.923 | 0.917 | 0.901 | 0.925 | 0.921 |
| 3055 | GG**Pro**GGG**Hip**GG | 0.851 | 0.850 | 0.811 | 0.835 | 0.826 | 0.828 | 0.831 |
| 3056 | GG**Pro**GGG**His**GG | 0.847 | 0.874 | 0.817 | 0.856 | 0.823 | 0.865 | 0.851 |
| 3057 | GG**Pro**GGG**Ile**GG | 0.875 | 0.790 | 0.781 | 0.864 | 0.838 | 0.787 | 0.814 |
| 3058 | GG**Pro**GGG**Leu**GG | 0.824 | 0.809 | 0.712 | 0.872 | 0.717 | 0.726 | 0.768 |
| 3059 | GG**Pro**GGG**Lys**GG | 0.815 | 0.861 | 0.880 | 0.771 | 0.753 | 0.834 | 0.825 |
| 3060 | GG**Pro**GGG**Met**GG | 0.816 | 0.858 | 0.817 | 0.829 | 0.749 | 0.788 | 0.816 |
| 3061 | GG**Pro**GGG**Phe**GG | 0.827 | 0.826 | 0.764 | 0.834 | 0.858 | 0.795 | 0.827 |
| 3062 | GG**Pro**GGG**Pro**GG | 0.909 | 0.928 | 0.906 | 0.906 | 0.878 | 0.906 | 0.906 |
| 3063 | GG**Pro**GGG**Ptr**GG | 0.746 | 0.819 | 0.761 | 0.758 | 0.753 | 0.783 | 0.759 |
| 3064 | GG**Pro**GGG**S1p**GG | 0.820 | 0.819 | 0.875 | 0.838 | 0.846 | 0.806 | 0.829 |
| 3065 | GG**Pro**GGG**Sep**GG | 0.873 | 0.802 | 0.827 | 0.838 | 0.875 | 0.853 | 0.846 |
| 3066 | GG**Pro**GGG**Ser**GG | 0.759 | 0.885 | 0.876 | 0.763 | 0.729 | 0.850 | 0.807 |
| 3067 | GG**Pro**GGG**T1p**GG | 0.807 | 0.550 | 0.719 | 0.620 | 0.700 | 0.353 | 0.660 |
| 3068 | GG**Pro**GGG**Thr**GG | 0.820 | 0.810 | 0.815 | 0.873 | 0.854 | 0.876 | 0.837 |
| 3069 | GG**Pro**GGG**Tpo**GG | 0.851 | 0.823 | 0.431 | 0.926 | 0.694 | 0.748 | 0.785 |
| 3070 | GG**Pro**GGG**Trp**GG | 0.781 | 0.811 | 0.870 | 0.833 | 0.717 | 0.781 | 0.796 |
| 3071 | GG**Pro**GGG**Tyr**GG | 0.867 | 0.864 | 0.761 | 0.882 | 0.840 | 0.830 | 0.852 |
| 3072 | GG**Pro**GGG**Val**GG | 0.797 | 0.806 | 0.756 | 0.777 | 0.824 | 0.816 | 0.801 |
| 3073 | GG**Pro**GGG**Y1p**GG | 0.850 | 0.831 | 0.841 | 0.797 | 0.816 | 0.742 | 0.824 |
| 3074 | GG**Ptr**GGG**Ala**GG | 0.653 | 0.750 | 0.555 | 0.789 | 0.888 | 0.729 | 0.740 |
| 3075 | GG**Ptr**GGG**Arg**GG | 0.598 | 0.437 | 0.566 | 0.666 | 0.506 | 0.381 | 0.536 |
| 3076 | GG**Ptr**GGG**Ash**GG | 0.809 | 0.733 | 0.726 | 0.703 | 0.683 | 0.692 | 0.715 |
| 3077 | GG**Ptr**GGG**Asn**GG | 0.809 | 0.742 | 0.809 | 0.811 | 0.842 | 0.747 | 0.809 |
| 3078 | GG**Ptr**GGG**Asp**GG | 0.729 | 0.719 | 0.812 | 0.815 | 0.799 | 0.788 | 0.793 |
| 3079 | GG**Ptr**GGG**Cys**GG | 0.768 | 0.769 | 0.831 | 0.756 | 0.808 | 0.789 | 0.779 |
| 3080 | GG**Ptr**GGG**Glh**GG | 0.701 | 0.693 | 0.723 | 0.672 | 0.653 | 0.825 | 0.697 |
| 3081 | GG**Ptr**GGG**Gln**GG | 0.838 | 0.748 | 0.807 | 0.686 | 0.746 | 0.718 | 0.747 |
| 3082 | GG**Ptr**GGG**Glu**GG | 0.791 | 0.431 | 0.783 | 0.534 | 0.857 | 0.519 | 0.658 |
| 3083 | GG**Ptr**GGG**Gly**GG | 0.870 | 0.796 | 0.882 | 0.791 | 0.859 | 0.786 | 0.828 |
| 3084 | GG**Ptr**GGG**Hip**GG | 0.612 | 0.395 | 0.342 | 0.512 | 0.477 | 0.602 | 0.495 |
| 3085 | GG**Ptr**GGG**His**GG | 0.714 | 0.497 | 0.716 | 0.604 | 0.739 | 0.681 | 0.698 |
| 3086 | GG**Ptr**GGG**Ile**GG | 0.771 | 0.742 | 0.708 | 0.772 | 0.727 | 0.731 | 0.737 |
| 3087 | GG**Ptr**GGG**Leu**GG | 0.426 | 0.457 | 0.455 | 0.729 | 0.799 | 0.826 | 0.593 |
| 3088 | GG**Ptr**GGG**Lys**GG | 0.664 | 0.709 | 0.756 | 0.572 | 0.642 | 0.635 | 0.653 |
| 3089 | GG**Ptr**GGG**Met**GG | 0.749 | 0.732 | 0.757 | 0.842 | 0.770 | 0.759 | 0.758 |
| 3090 | GG**Ptr**GGG**Phe**GG | 0.784 | 0.739 | 0.678 | 0.847 | 0.741 | 0.759 | 0.750 |
| 3091 | GG**Ptr**GGG**Pro**GG | 0.862 | 0.830 | 0.786 | 0.881 | 0.815 | 0.864 | 0.846 |
| 3092 | GG**Ptr**GGG**Ptr**GG | 0.626 | 0.696 | 0.754 | 0.762 | 0.668 | 0.714 | 0.705 |
| 3093 | GG**Ptr**GGG**S1p**GG | 0.710 | 0.796 | 0.799 | 0.784 | 0.804 | 0.822 | 0.797 |
| 3094 | GG**Ptr**GGG**Sep**GG | 0.797 | 0.772 | 0.821 | 0.806 | 0.795 | 0.798 | 0.798 |
| 3095 | GG**Ptr**GGG**Ser**GG | 0.763 | 0.771 | 0.827 | 0.851 | 0.805 | 0.789 | 0.797 |
| 3096 | GG**Ptr**GGG**T1p**GG | 0.755 | 0.427 | 0.829 | 0.624 | 0.801 | 0.447 | 0.689 |
| 3097 | GG**Ptr**GGG**Thr**GG | 0.856 | 0.776 | 0.802 | 0.751 | 0.792 | 0.760 | 0.784 |
| 3098 | GG**Ptr**GGG**Tpo**GG | 0.714 | 0.377 | 0.439 | 0.799 | 0.847 | 0.880 | 0.757 |
| 3099 | GG**Ptr**GGG**Trp**GG | 0.710 | 0.688 | 0.672 | 0.710 | 0.724 | 0.715 | 0.710 |

This work is shared under a CC BY 4.0 license.

| | Peptide | $r_{replica}$(1,2) | $r_{replica}$(1,3) | $r_{replica}$(1,4) | $r_{replica}$(2,3) | $r_{replica}$(2,4) | $r_{replica}$(3,4) | Median |
|---|---|---|---|---|---|---|---|---|
| 3100 | GG**Ptr**GGG**Tyr**GG | 0.742 | 0.768 | 0.738 | 0.739 | 0.714 | 0.760 | 0.740 |
| 3101 | GG**Ptr**GGG**Val**GG | 0.670 | 0.724 | 0.736 | 0.689 | 0.712 | 0.783 | 0.718 |
| 3102 | GG**Ptr**GGG**Y1p**GG | 0.720 | 0.730 | 0.601 | 0.744 | 0.734 | 0.580 | 0.725 |
| 3103 | GG**S1p**GGG**Ala**GG | 0.791 | 0.786 | 0.770 | 0.870 | 0.844 | 0.876 | 0.818 |
| 3104 | GG**S1p**GGG**Arg**GG | 0.648 | 0.639 | 0.680 | 0.585 | 0.558 | 0.559 | 0.612 |
| 3105 | GG**S1p**GGG**Ash**GG | 0.698 | 0.760 | 0.681 | 0.866 | 0.814 | 0.821 | 0.787 |
| 3106 | GG**S1p**GGG**Asn**GG | 0.832 | 0.836 | 0.842 | 0.832 | 0.810 | 0.885 | 0.834 |
| 3107 | GG**S1p**GGG**Asp**GG | 0.538 | 0.815 | 0.754 | 0.475 | 0.467 | 0.805 | 0.646 |
| 3108 | GG**S1p**GGG**Cys**GG | 0.729 | 0.773 | 0.716 | 0.855 | 0.766 | 0.778 | 0.770 |
| 3109 | GG**S1p**GGG**Glh**GG | 0.501 | 0.722 | 0.739 | 0.594 | 0.558 | 0.719 | 0.656 |
| 3110 | GG**S1p**GGG**Gln**GG | 0.804 | 0.848 | 0.788 | 0.824 | 0.816 | 0.795 | 0.810 |
| 3111 | GG**S1p**GGG**Glu**GG | 0.720 | 0.716 | 0.757 | 0.667 | 0.820 | 0.677 | 0.718 |
| 3112 | GG**S1p**GGG**Gly**GG | 0.887 | 0.870 | 0.890 | 0.887 | 0.909 | 0.850 | 0.887 |
| 3113 | GG**S1p**GGG**Hip**GG | 0.704 | 0.738 | 0.696 | 0.757 | 0.683 | 0.712 | 0.708 |
| 3114 | GG**S1p**GGG**His**GG | 0.727 | 0.738 | 0.715 | 0.720 | 0.720 | 0.805 | 0.723 |
| 3115 | GG**S1p**GGG**Ile**GG | 0.644 | 0.678 | 0.684 | 0.815 | 0.777 | 0.820 | 0.730 |
| 3116 | GG**S1p**GGG**Leu**GG | 0.716 | 0.788 | 0.786 | 0.867 | 0.796 | 0.865 | 0.792 |
| 3117 | GG**S1p**GGG**Lys**GG | 0.766 | 0.777 | 0.795 | 0.805 | 0.848 | 0.831 | 0.800 |
| 3118 | GG**S1p**GGG**Met**GG | 0.860 | 0.876 | 0.827 | 0.830 | 0.809 | 0.825 | 0.828 |
| 3119 | GG**S1p**GGG**Phe**GG | 0.818 | 0.820 | 0.787 | 0.841 | 0.790 | 0.866 | 0.819 |
| 3120 | GG**S1p**GGG**Pro**GG | 0.825 | 0.831 | 0.858 | 0.801 | 0.756 | 0.824 | 0.825 |
| 3121 | GG**S1p**GGG**Ptr**GG | 0.806 | 0.868 | 0.766 | 0.797 | 0.822 | 0.771 | 0.801 |
| 3122 | GG**S1p**GGG**S1p**GG | 0.786 | 0.764 | 0.768 | 0.838 | 0.802 | 0.847 | 0.794 |
| 3123 | GG**S1p**GGG**Sep**GG | 0.851 | 0.835 | 0.833 | 0.778 | 0.829 | 0.814 | 0.831 |
| 3124 | GG**S1p**GGG**Ser**GG | 0.778 | 0.763 | 0.838 | 0.797 | 0.824 | 0.853 | 0.810 |
| 3125 | GG**S1p**GGG**T1p**GG | 0.502 | 0.393 | 0.673 | 0.758 | 0.777 | 0.677 | 0.675 |
| 3126 | GG**S1p**GGG**Thr**GG | 0.838 | 0.835 | 0.847 | 0.795 | 0.843 | 0.828 | 0.836 |
| 3127 | GG**S1p**GGG**Tpo**GG | 0.670 | 0.820 | 0.793 | 0.541 | 0.767 | 0.783 | 0.775 |
| 3128 | GG**S1p**GGG**Trp**GG | 0.759 | 0.801 | 0.786 | 0.766 | 0.844 | 0.826 | 0.794 |
| 3129 | GG**S1p**GGG**Tyr**GG | 0.776 | 0.780 | 0.826 | 0.794 | 0.821 | 0.790 | 0.792 |
| 3130 | GG**S1p**GGG**Val**GG | 0.745 | 0.777 | 0.806 | 0.679 | 0.764 | 0.775 | 0.770 |
| 3131 | GG**S1p**GGG**Y1p**GG | 0.823 | 0.730 | 0.847 | 0.643 | 0.795 | 0.744 | 0.770 |
| 3132 | GG**Sep**GGG**Ala**GG | 0.829 | 0.833 | 0.840 | 0.788 | 0.852 | 0.806 | 0.831 |
| 3133 | GG**Sep**GGG**Arg**GG | 0.589 | 0.662 | 0.626 | 0.750 | 0.566 | 0.650 | 0.638 |
| 3134 | GG**Sep**GGG**Ash**GG | 0.828 | 0.863 | 0.784 | 0.803 | 0.789 | 0.839 | 0.816 |
| 3135 | GG**Sep**GGG**Asn**GG | 0.770 | 0.822 | 0.867 | 0.817 | 0.817 | 0.827 | 0.819 |
| 3136 | GG**Sep**GGG**Asp**GG | 0.768 | 0.766 | 0.814 | 0.751 | 0.787 | 0.768 | 0.768 |
| 3137 | GG**Sep**GGG**Cys**GG | 0.816 | 0.785 | 0.786 | 0.807 | 0.863 | 0.797 | 0.802 |
| 3138 | GG**Sep**GGG**Glh**GG | 0.754 | 0.720 | 0.702 | 0.787 | 0.726 | 0.721 | 0.723 |
| 3139 | GG**Sep**GGG**Gln**GG | 0.757 | 0.808 | 0.778 | 0.795 | 0.784 | 0.778 | 0.781 |
| 3140 | GG**Sep**GGG**Glu**GG | 0.839 | 0.817 | 0.803 | 0.797 | 0.853 | 0.806 | 0.811 |
| 3141 | GG**Sep**GGG**Gly**GG | 0.639 | 0.769 | 0.727 | 0.804 | 0.804 | 0.893 | 0.786 |
| 3142 | GG**Sep**GGG**Hip**GG | 0.590 | 0.651 | 0.585 | 0.631 | 0.648 | 0.575 | 0.611 |
| 3143 | GG**Sep**GGG**His**GG | 0.709 | 0.793 | 0.880 | 0.628 | 0.676 | 0.861 | 0.751 |
| 3144 | GG**Sep**GGG**Ile**GG | 0.808 | 0.854 | 0.800 | 0.822 | 0.840 | 0.784 | 0.815 |
| 3145 | GG**Sep**GGG**Leu**GG | 0.840 | 0.777 | 0.838 | 0.784 | 0.818 | 0.762 | 0.801 |
| 3146 | GG**Sep**GGG**Lys**GG | 0.750 | 0.703 | 0.652 | 0.780 | 0.768 | 0.750 | 0.750 |
| 3147 | GG**Sep**GGG**Met**GG | 0.780 | 0.770 | 0.770 | 0.853 | 0.823 | 0.843 | 0.802 |
| 3148 | GG**Sep**GGG**Phe**GG | 0.794 | 0.711 | 0.761 | 0.765 | 0.760 | 0.783 | 0.763 |
| 3149 | GG**Sep**GGG**Pro**GG | 0.875 | 0.834 | 0.869 | 0.851 | 0.843 | 0.809 | 0.847 |

This work is shared under a CC BY 4.0 license.

| | Peptide | $r_{replica}$(1,2) | $r_{replica}$(1,3) | $r_{replica}$(1,4) | $r_{replica}$(2,3) | $r_{replica}$(2,4) | $r_{replica}$(3,4) | Median |
|---|---|---|---|---|---|---|---|---|
| 3150 | GG**Sep**GGG**Ptr**GG | 0.638 | 0.659 | 0.642 | 0.690 | 0.817 | 0.695 | 0.674 |
| 3151 | GG**Sep**GGG**S1p**GG | 0.859 | 0.880 | 0.646 | 0.899 | 0.563 | 0.562 | 0.752 |
| 3152 | GG**Sep**GGG**Sep**GG | 0.749 | 0.741 | 0.759 | 0.722 | 0.796 | 0.796 | 0.754 |
| 3153 | GG**Sep**GGG**Ser**GG | 0.834 | 0.831 | 0.792 | 0.860 | 0.840 | 0.847 | 0.837 |
| 3154 | GG**Sep**GGG**T1p**GG | 0.587 | 0.841 | 0.409 | 0.626 | 0.697 | 0.459 | 0.606 |
| 3155 | GG**Sep**GGG**Thr**GG | 0.858 | 0.833 | 0.794 | 0.809 | 0.758 | 0.807 | 0.808 |
| 3156 | GG**Sep**GGG**Tpo**GG | 0.779 | 0.787 | 0.699 | 0.859 | 0.814 | 0.845 | 0.800 |
| 3157 | GG**Sep**GGG**Trp**GG | 0.652 | 0.751 | 0.735 | 0.602 | 0.637 | 0.739 | 0.694 |
| 3158 | GG**Sep**GGG**Tyr**GG | 0.792 | 0.795 | 0.622 | 0.850 | 0.781 | 0.776 | 0.787 |
| 3159 | GG**Sep**GGG**Val**GG | 0.796 | 0.767 | 0.793 | 0.785 | 0.873 | 0.762 | 0.789 |
| 3160 | GG**Sep**GGG**Y1p**GG | 0.815 | 0.843 | 0.843 | 0.789 | 0.803 | 0.860 | 0.829 |
| 3161 | GG**Ser**GGG**Ala**GG | 0.862 | 0.849 | 0.836 | 0.848 | 0.858 | 0.843 | 0.849 |
| 3162 | GG**Ser**GGG**Arg**GG | 0.816 | 0.773 | 0.829 | 0.811 | 0.829 | 0.796 | 0.813 |
| 3163 | GG**Ser**GGG**Ash**GG | 0.877 | 0.838 | 0.872 | 0.850 | 0.888 | 0.887 | 0.874 |
| 3164 | GG**Ser**GGG**Asn**GG | 0.850 | 0.867 | 0.870 | 0.880 | 0.892 | 0.894 | 0.875 |
| 3165 | GG**Ser**GGG**Asp**GG | 0.848 | 0.799 | 0.795 | 0.785 | 0.800 | 0.716 | 0.797 |
| 3166 | GG**Ser**GGG**Cys**GG | 0.859 | 0.867 | 0.838 | 0.852 | 0.853 | 0.859 | 0.856 |
| 3167 | GG**Ser**GGG**Glh**GG | 0.800 | 0.799 | 0.788 | 0.811 | 0.797 | 0.789 | 0.798 |
| 3168 | GG**Ser**GGG**Gln**GG | 0.832 | 0.830 | 0.867 | 0.785 | 0.847 | 0.809 | 0.831 |
| 3169 | GG**Ser**GGG**Glu**GG | 0.722 | 0.869 | 0.821 | 0.733 | 0.703 | 0.837 | 0.777 |
| 3170 | GG**Ser**GGG**Gly**GG | 0.924 | 0.901 | 0.930 | 0.911 | 0.904 | 0.915 | 0.913 |
| 3171 | GG**Ser**GGG**Hip**GG | 0.774 | 0.765 | 0.811 | 0.812 | 0.817 | 0.746 | 0.793 |
| 3172 | GG**Ser**GGG**His**GG | 0.726 | 0.793 | 0.858 | 0.832 | 0.803 | 0.879 | 0.818 |
| 3173 | GG**Ser**GGG**Ile**GG | 0.830 | 0.721 | 0.792 | 0.676 | 0.763 | 0.702 | 0.742 |
| 3174 | GG**Ser**GGG**Leu**GG | 0.821 | 0.827 | 0.823 | 0.908 | 0.856 | 0.893 | 0.842 |
| 3175 | GG**Ser**GGG**Lys**GG | 0.804 | 0.757 | 0.838 | 0.739 | 0.820 | 0.780 | 0.792 |
| 3176 | GG**Ser**GGG**Met**GG | 0.841 | 0.855 | 0.805 | 0.862 | 0.830 | 0.829 | 0.836 |
| 3177 | GG**Ser**GGG**Phe**GG | 0.868 | 0.889 | 0.860 | 0.867 | 0.858 | 0.926 | 0.868 |
| 3178 | GG**Ser**GGG**Pro**GG | 0.880 | 0.895 | 0.868 | 0.886 | 0.877 | 0.885 | 0.883 |
| 3179 | GG**Ser**GGG**Ptr**GG | 0.783 | 0.732 | 0.777 | 0.739 | 0.859 | 0.760 | 0.768 |
| 3180 | GG**Ser**GGG**S1p**GG | 0.897 | 0.889 | 0.852 | 0.859 | 0.860 | 0.845 | 0.860 |
| 3181 | GG**Ser**GGG**Sep**GG | 0.834 | 0.787 | 0.795 | 0.733 | 0.764 | 0.780 | 0.784 |
| 3182 | GG**Ser**GGG**Ser**GG | 0.849 | 0.880 | 0.863 | 0.859 | 0.856 | 0.862 | 0.861 |
| 3183 | GG**Ser**GGG**T1p**GG | 0.934 | 0.735 | 0.929 | 0.792 | 0.938 | 0.794 | 0.861 |
| 3184 | GG**Ser**GGG**Thr**GG | 0.786 | 0.865 | 0.824 | 0.766 | 0.807 | 0.855 | 0.816 |
| 3185 | GG**Ser**GGG**Tpo**GG | 0.580 | 0.869 | 0.817 | 0.825 | 0.167 | 0.634 | 0.725 |
| 3186 | GG**Ser**GGG**Trp**GG | 0.711 | 0.818 | 0.877 | 0.674 | 0.703 | 0.825 | 0.764 |
| 3187 | GG**Ser**GGG**Tyr**GG | 0.867 | 0.875 | 0.753 | 0.849 | 0.835 | 0.803 | 0.842 |
| 3188 | GG**Ser**GGG**Val**GG | 0.826 | 0.802 | 0.840 | 0.804 | 0.829 | 0.786 | 0.815 |
| 3189 | GG**Ser**GGG**Y1p**GG | 0.794 | 0.862 | 0.828 | 0.790 | 0.734 | 0.895 | 0.811 |
| 3190 | GG**T1p**GGG**Ala**GG | 0.776 | 0.715 | 0.728 | 0.612 | 0.824 | 0.498 | 0.722 |
| 3191 | GG**T1p**GGG**Arg**GG | 0.687 | 0.816 | 0.805 | 0.774 | 0.672 | 0.784 | 0.779 |
| 3192 | GG**T1p**GGG**Ash**GG | 0.660 | 0.320 | 0.825 | 0.679 | 0.693 | 0.315 | 0.670 |
| 3193 | GG**T1p**GGG**Asn**GG | 0.615 | 0.826 | 0.464 | 0.718 | 0.798 | 0.637 | 0.677 |
| 3194 | GG**T1p**GGG**Asp**GG | 0.533 | 0.759 | 0.487 | 0.732 | 0.766 | 0.664 | 0.698 |
| 3195 | GG**T1p**GGG**Cys**GG | 0.791 | 0.773 | 0.813 | 0.749 | 0.787 | 0.742 | 0.780 |
| 3196 | GG**T1p**GGG**Glh**GG | 0.387 | 0.775 | 0.879 | 0.749 | 0.560 | 0.832 | 0.762 |
| 3197 | GG**T1p**GGG**Gln**GG | 0.695 | 0.711 | 0.718 | 0.632 | 0.544 | 0.888 | 0.703 |
| 3198 | GG**T1p**GGG**Glu**GG | 0.512 | 0.705 | 0.397 | 0.768 | 0.836 | 0.730 | 0.717 |
| 3199 | GG**T1p**GGG**Gly**GG | 0.720 | 0.914 | 0.578 | 0.674 | 0.882 | 0.532 | 0.697 |

This work is shared under a CC BY 4.0 license.

| | Peptide | $r_{replica}$(1,2) | $r_{replica}$(1,3) | $r_{replica}$(1,4) | $r_{replica}$(2,3) | $r_{replica}$(2,4) | $r_{replica}$(3,4) | Median |
|---|---|---|---|---|---|---|---|---|
| 3200 | GG**T1p**GGG**Hip**GG | 0.526 | 0.530 | 0.799 | 0.715 | 0.412 | 0.376 | 0.528 |
| 3201 | GG**T1p**GGG**His**GG | 0.650 | 0.752 | 0.772 | 0.549 | 0.555 | 0.805 | 0.701 |
| 3202 | GG**T1p**GGG**Ile**GG | 0.432 | 0.643 | 0.743 | 0.637 | 0.555 | 0.630 | 0.633 |
| 3203 | GG**T1p**GGG**Leu**GG | 0.392 | 0.859 | 0.890 | 0.414 | 0.368 | 0.881 | 0.637 |
| 3204 | GG**T1p**GGG**Lys**GG | 0.715 | 0.824 | 0.763 | 0.768 | 0.726 | 0.819 | 0.766 |
| 3205 | GG**T1p**GGG**Met**GG | 0.765 | 0.746 | 0.580 | 0.593 | 0.379 | 0.815 | 0.669 |
| 3206 | GG**T1p**GGG**Phe**GG | 0.406 | 0.356 | 0.646 | 0.851 | 0.768 | 0.769 | 0.707 |
| 3207 | GG**T1p**GGG**Pro**GG | 0.553 | 0.503 | 0.833 | 0.904 | 0.603 | 0.569 | 0.586 |
| 3208 | GG**T1p**GGG**Ptr**GG | 0.374 | 0.415 | 0.384 | 0.784 | 0.783 | 0.797 | 0.599 |
| 3209 | GG**T1p**GGG**S1p**GG | 0.444 | 0.809 | 0.480 | 0.401 | 0.775 | 0.385 | 0.462 |
| 3210 | GG**T1p**GGG**Sep**GG | 0.709 | 0.530 | 0.873 | 0.820 | 0.782 | 0.606 | 0.746 |
| 3211 | GG**T1p**GGG**Ser**GG | 0.670 | 0.796 | 0.660 | 0.607 | 0.875 | 0.562 | 0.665 |
| 3212 | GG**T1p**GGG**T1p**GG | 0.513 | 0.778 | 0.729 | 0.479 | 0.454 | 0.819 | 0.621 |
| 3213 | GG**T1p**GGG**Thr**GG | 0.795 | 0.795 | 0.568 | 0.752 | 0.504 | 0.741 | 0.747 |
| 3214 | GG**T1p**GGG**Tpo**GG | 0.547 | 0.381 | 0.564 | 0.813 | 0.688 | 0.734 | 0.626 |
| 3215 | GG**T1p**GGG**Trp**GG | 0.721 | 0.622 | 0.646 | 0.498 | 0.759 | 0.444 | 0.634 |
| 3216 | GG**T1p**GGG**Tyr**GG | 0.838 | 0.453 | 0.329 | 0.465 | 0.419 | 0.882 | 0.459 |
| 3217 | GG**T1p**GGG**Val**GG | 0.715 | 0.756 | 0.709 | 0.769 | 0.819 | 0.763 | 0.760 |
| 3218 | GG**T1p**GGG**Y1p**GG | 0.732 | 0.736 | 0.783 | 0.730 | 0.598 | 0.686 | 0.731 |
| 3219 | GG**Thr**GGG**Ala**GG | 0.821 | 0.842 | 0.863 | 0.876 | 0.825 | 0.887 | 0.853 |
| 3220 | GG**Thr**GGG**Arg**GG | 0.877 | 0.837 | 0.719 | 0.809 | 0.762 | 0.724 | 0.785 |
| 3221 | GG**Thr**GGG**Ash**GG | 0.870 | 0.853 | 0.877 | 0.778 | 0.902 | 0.776 | 0.861 |
| 3222 | GG**Thr**GGG**Asn**GG | 0.884 | 0.899 | 0.778 | 0.865 | 0.805 | 0.788 | 0.835 |
| 3223 | GG**Thr**GGG**Asp**GG | 0.795 | 0.850 | 0.757 | 0.806 | 0.791 | 0.760 | 0.793 |
| 3224 | GG**Thr**GGG**Cys**GG | 0.862 | 0.896 | 0.865 | 0.877 | 0.772 | 0.808 | 0.863 |
| 3225 | GG**Thr**GGG**Glh**GG | 0.765 | 0.789 | 0.757 | 0.811 | 0.806 | 0.797 | 0.793 |
| 3226 | GG**Thr**GGG**Gln**GG | 0.807 | 0.838 | 0.899 | 0.855 | 0.876 | 0.878 | 0.866 |
| 3227 | GG**Thr**GGG**Glu**GG | 0.841 | 0.869 | 0.813 | 0.804 | 0.773 | 0.834 | 0.823 |
| 3228 | GG**Thr**GGG**Gly**GG | 0.891 | 0.915 | 0.897 | 0.912 | 0.921 | 0.911 | 0.911 |
| 3229 | GG**Thr**GGG**Hip**GG | 0.867 | 0.816 | 0.858 | 0.848 | 0.872 | 0.811 | 0.853 |
| 3230 | GG**Thr**GGG**His**GG | 0.721 | 0.779 | 0.789 | 0.815 | 0.652 | 0.761 | 0.770 |
| 3231 | GG**Thr**GGG**Ile**GG | 0.864 | 0.833 | 0.805 | 0.797 | 0.782 | 0.802 | 0.804 |
| 3232 | GG**Thr**GGG**Leu**GG | 0.845 | 0.895 | 0.845 | 0.848 | 0.841 | 0.884 | 0.847 |
| 3233 | GG**Thr**GGG**Lys**GG | 0.821 | 0.818 | 0.842 | 0.810 | 0.806 | 0.826 | 0.820 |
| 3234 | GG**Thr**GGG**Met**GG | 0.893 | 0.883 | 0.853 | 0.864 | 0.839 | 0.748 | 0.858 |
| 3235 | GG**Thr**GGG**Phe**GG | 0.824 | 0.809 | 0.843 | 0.796 | 0.796 | 0.853 | 0.817 |
| 3236 | GG**Thr**GGG**Pro**GG | 0.791 | 0.879 | 0.861 | 0.794 | 0.870 | 0.859 | 0.860 |
| 3237 | GG**Thr**GGG**Ptr**GG | 0.819 | 0.818 | 0.860 | 0.737 | 0.808 | 0.836 | 0.818 |
| 3238 | GG**Thr**GGG**S1p**GG | 0.793 | 0.727 | 0.842 | 0.800 | 0.802 | 0.740 | 0.797 |
| 3239 | GG**Thr**GGG**Sep**GG | 0.868 | 0.858 | 0.679 | 0.873 | 0.639 | 0.629 | 0.769 |
| 3240 | GG**Thr**GGG**Ser**GG | 0.863 | 0.815 | 0.867 | 0.830 | 0.844 | 0.864 | 0.853 |
| 3241 | GG**Thr**GGG**T1p**GG | 0.785 | 0.815 | 0.793 | 0.845 | 0.860 | 0.886 | 0.830 |
| 3242 | GG**Thr**GGG**Thr**GG | 0.793 | 0.808 | 0.734 | 0.800 | 0.764 | 0.791 | 0.792 |
| 3243 | GG**Thr**GGG**Tpo**GG | 0.258 | 0.927 | 0.630 | 0.358 | 0.859 | 0.679 | 0.655 |
| 3244 | GG**Thr**GGG**Trp**GG | 0.753 | 0.580 | 0.580 | 0.806 | 0.785 | 0.819 | 0.769 |
| 3245 | GG**Thr**GGG**Tyr**GG | 0.769 | 0.822 | 0.803 | 0.752 | 0.722 | 0.756 | 0.763 |
| 3246 | GG**Thr**GGG**Val**GG | 0.745 | 0.774 | 0.769 | 0.739 | 0.771 | 0.772 | 0.770 |
| 3247 | GG**Thr**GGG**Y1p**GG | 0.806 | 0.789 | 0.783 | 0.820 | 0.863 | 0.850 | 0.813 |
| 3248 | GG**Tpo**GGG**Ala**GG | 0.888 | 0.617 | 0.809 | 0.554 | 0.884 | 0.344 | 0.713 |
| 3249 | GG**Tpo**GGG**Arg**GG | 0.398 | 0.557 | 0.430 | 0.621 | 0.455 | 0.515 | 0.485 |

This work is shared under a CC BY 4.0 license.

| | Peptide | $r_{replica}$(1,2) | $r_{replica}$(1,3) | $r_{replica}$(1,4) | $r_{replica}$(2,3) | $r_{replica}$(2,4) | $r_{replica}$(3,4) | Median |
|---|---|---|---|---|---|---|---|---|
| 3250 | GG**Tpo**GGG**Ash**GG | 0.722 | 0.791 | 0.369 | 0.871 | 0.700 | 0.690 | 0.711 |
| 3251 | GG**Tpo**GGG**Asn**GG | 0.602 | 0.703 | 0.629 | 0.847 | 0.807 | 0.855 | 0.755 |
| 3252 | GG**Tpo**GGG**Asp**GG | 0.949 | 0.164 | 0.789 | 0.227 | 0.808 | 0.632 | 0.711 |
| 3253 | GG**Tpo**GGG**Cys**GG | 0.641 | 0.681 | 0.681 | 0.843 | 0.802 | 0.777 | 0.730 |
| 3254 | GG**Tpo**GGG**Glh**GG | 0.762 | 0.715 | 0.729 | 0.459 | 0.890 | 0.363 | 0.722 |
| 3255 | GG**Tpo**GGG**Gln**GG | 0.677 | 0.854 | 0.842 | 0.628 | 0.772 | 0.809 | 0.790 |
| 3256 | GG**Tpo**GGG**Glu**GG | 0.831 | 0.812 | 0.699 | 0.840 | 0.595 | 0.455 | 0.756 |
| 3257 | GG**Tpo**GGG**Gly**GG | 0.888 | 0.875 | 0.867 | 0.866 | 0.837 | 0.850 | 0.866 |
| 3258 | GG**Tpo**GGG**Hip**GG | 0.580 | 0.681 | 0.661 | 0.624 | 0.565 | 0.583 | 0.603 |
| 3259 | GG**Tpo**GGG**His**GG | 0.328 | 0.832 | 0.743 | 0.188 | 0.643 | 0.716 | 0.680 |
| 3260 | GG**Tpo**GGG**Ile**GG | 0.791 | 0.743 | 0.633 | 0.727 | 0.600 | 0.585 | 0.680 |
| 3261 | GG**Tpo**GGG**Leu**GG | 0.427 | 0.211 | 0.797 | 0.848 | 0.539 | 0.338 | 0.483 |
| 3262 | GG**Tpo**GGG**Lys**GG | 0.711 | 0.777 | 0.288 | 0.835 | 0.642 | 0.584 | 0.677 |
| 3263 | GG**Tpo**GGG**Met**GG | 0.714 | 0.730 | 0.729 | 0.692 | 0.714 | 0.712 | 0.714 |
| 3264 | GG**Tpo**GGG**Phe**GG | 0.861 | 0.933 | 0.558 | 0.866 | 0.795 | 0.576 | 0.828 |
| 3265 | GG**Tpo**GGG**Pro**GG | 0.786 | 0.795 | 0.763 | 0.804 | 0.743 | 0.760 | 0.774 |
| 3266 | GG**Tpo**GGG**Ptr**GG | 0.741 | 0.665 | 0.776 | 0.626 | 0.669 | 0.710 | 0.690 |
| 3267 | GG**Tpo**GGG**S1p**GG | 0.758 | 0.752 | 0.812 | 0.845 | 0.822 | 0.844 | 0.817 |
| 3268 | GG**Tpo**GGG**Sep**GG | 0.911 | 0.461 | 0.374 | 0.402 | 0.300 | 0.929 | 0.432 |
| 3269 | GG**Tpo**GGG**Ser**GG | 0.925 | 0.894 | 0.471 | 0.934 | 0.425 | 0.336 | 0.683 |
| 3270 | GG**Tpo**GGG**T1p**GG | 0.732 | 0.531 | 0.525 | 0.363 | 0.492 | 0.614 | 0.528 |
| 3271 | GG**Tpo**GGG**Thr**GG | 0.796 | 0.773 | 0.767 | 0.728 | 0.702 | 0.827 | 0.770 |
| 3272 | GG**Tpo**GGG**Tpo**GG | 0.260 | -0.017 | 0.902 | 0.585 | 0.138 | -0.007 | 0.199 |
| 3273 | GG**Tpo**GGG**Trp**GG | 0.677 | 0.747 | 0.675 | 0.783 | 0.746 | 0.714 | 0.730 |
| 3274 | GG**Tpo**GGG**Tyr**GG | 0.583 | 0.632 | 0.655 | 0.617 | 0.760 | 0.751 | 0.644 |
| 3275 | GG**Tpo**GGG**Val**GG | 0.749 | 0.554 | 0.738 | 0.465 | 0.718 | 0.468 | 0.636 |
| 3276 | GG**Tpo**GGG**Y1p**GG | 0.799 | 0.818 | 0.507 | 0.783 | 0.564 | 0.425 | 0.674 |
| 3277 | GG**Trp**GGG**Ala**GG | 0.774 | 0.780 | 0.788 | 0.706 | 0.763 | 0.719 | 0.769 |
| 3278 | GG**Trp**GGG**Arg**GG | 0.704 | 0.620 | 0.525 | 0.704 | 0.531 | 0.549 | 0.585 |
| 3279 | GG**Trp**GGG**Ash**GG | 0.711 | 0.734 | 0.705 | 0.862 | 0.723 | 0.725 | 0.724 |
| 3280 | GG**Trp**GGG**Asn**GG | 0.696 | 0.750 | 0.810 | 0.810 | 0.758 | 0.821 | 0.784 |
| 3281 | GG**Trp**GGG**Asp**GG | 0.820 | 0.779 | 0.812 | 0.837 | 0.852 | 0.845 | 0.828 |
| 3282 | GG**Trp**GGG**Cys**GG | 0.641 | 0.743 | 0.651 | 0.731 | 0.495 | 0.677 | 0.664 |
| 3283 | GG**Trp**GGG**Glh**GG | 0.697 | 0.746 | 0.749 | 0.750 | 0.648 | 0.763 | 0.748 |
| 3284 | GG**Trp**GGG**Gln**GG | 0.813 | 0.612 | 0.785 | 0.679 | 0.809 | 0.650 | 0.732 |
| 3285 | GG**Trp**GGG**Glu**GG | 0.752 | 0.840 | 0.846 | 0.778 | 0.737 | 0.807 | 0.792 |
| 3286 | GG**Trp**GGG**Gly**GG | 0.879 | 0.818 | 0.830 | 0.835 | 0.881 | 0.719 | 0.833 |
| 3287 | GG**Trp**GGG**Hip**GG | 0.791 | 0.776 | 0.818 | 0.773 | 0.719 | 0.783 | 0.779 |
| 3288 | GG**Trp**GGG**His**GG | 0.823 | 0.806 | 0.855 | 0.832 | 0.852 | 0.808 | 0.827 |
| 3289 | GG**Trp**GGG**Ile**GG | 0.732 | 0.736 | 0.777 | 0.775 | 0.791 | 0.839 | 0.776 |
| 3290 | GG**Trp**GGG**Leu**GG | 0.713 | 0.699 | 0.811 | 0.726 | 0.663 | 0.744 | 0.720 |
| 3291 | GG**Trp**GGG**Lys**GG | 0.793 | 0.551 | 0.581 | 0.714 | 0.570 | 0.344 | 0.576 |
| 3292 | GG**Trp**GGG**Met**GG | 0.636 | 0.641 | 0.693 | 0.781 | 0.754 | 0.744 | 0.719 |
| 3293 | GG**Trp**GGG**Phe**GG | 0.675 | 0.800 | 0.722 | 0.675 | 0.652 | 0.642 | 0.675 |
| 3294 | GG**Trp**GGG**Pro**GG | 0.723 | 0.656 | 0.643 | 0.765 | 0.598 | 0.747 | 0.690 |
| 3295 | GG**Trp**GGG**Ptr**GG | 0.581 | 0.740 | 0.604 | 0.627 | 0.558 | 0.597 | 0.600 |
| 3296 | GG**Trp**GGG**S1p**GG | 0.693 | 0.695 | 0.781 | 0.677 | 0.773 | 0.718 | 0.706 |
| 3297 | GG**Trp**GGG**Sep**GG | 0.739 | 0.663 | 0.707 | 0.707 | 0.716 | 0.819 | 0.712 |
| 3298 | GG**Trp**GGG**Ser**GG | 0.684 | 0.666 | 0.742 | 0.854 | 0.860 | 0.875 | 0.798 |
| 3299 | GG**Trp**GGG**T1p**GG | 0.657 | 0.890 | 0.881 | 0.497 | 0.410 | 0.899 | 0.769 |

This work is shared under a CC BY 4.0 license.

| | Peptide | $r_{replica}$(1,2) | $r_{replica}$(1,3) | $r_{replica}$(1,4) | $r_{replica}$(2,3) | $r_{replica}$(2,4) | $r_{replica}$(3,4) | Median |
|---|---|---|---|---|---|---|---|---|
| 3300 | GG**Trp**GGG**Thr**GG | 0.596 | 0.590 | 0.562 | 0.749 | 0.686 | 0.802 | 0.641 |
| 3301 | GG**Trp**GGG**Tpo**GG | 0.804 | 0.696 | 0.828 | 0.797 | 0.786 | 0.664 | 0.791 |
| 3302 | GG**Trp**GGG**Trp**GG | 0.808 | 0.822 | 0.662 | 0.850 | 0.630 | 0.689 | 0.749 |
| 3303 | GG**Trp**GGG**Tyr**GG | 0.820 | 0.759 | 0.635 | 0.789 | 0.752 | 0.676 | 0.755 |
| 3304 | GG**Trp**GGG**Val**GG | 0.739 | 0.761 | 0.757 | 0.683 | 0.689 | 0.756 | 0.747 |
| 3305 | GG**Trp**GGG**Y1p**GG | 0.724 | 0.695 | 0.697 | 0.665 | 0.716 | 0.631 | 0.696 |
| 3306 | GG**Tyr**GGG**Ala**GG | 0.826 | 0.873 | 0.844 | 0.847 | 0.875 | 0.893 | 0.860 |
| 3307 | GG**Tyr**GGG**Arg**GG | 0.768 | 0.788 | 0.788 | 0.751 | 0.707 | 0.785 | 0.777 |
| 3308 | GG**Tyr**GGG**Ash**GG | 0.825 | 0.847 | 0.858 | 0.814 | 0.837 | 0.877 | 0.842 |
| 3309 | GG**Tyr**GGG**Asn**GG | 0.718 | 0.753 | 0.733 | 0.874 | 0.835 | 0.829 | 0.791 |
| 3310 | GG**Tyr**GGG**Asp**GG | 0.847 | 0.808 | 0.720 | 0.740 | 0.823 | 0.681 | 0.774 |
| 3311 | GG**Tyr**GGG**Cys**GG | 0.849 | 0.822 | 0.870 | 0.841 | 0.860 | 0.895 | 0.855 |
| 3312 | GG**Tyr**GGG**Glh**GG | 0.734 | 0.826 | 0.846 | 0.671 | 0.696 | 0.857 | 0.780 |
| 3313 | GG**Tyr**GGG**Gln**GG | 0.853 | 0.851 | 0.814 | 0.804 | 0.850 | 0.784 | 0.832 |
| 3314 | GG**Tyr**GGG**Glu**GG | 0.820 | 0.777 | 0.790 | 0.738 | 0.835 | 0.695 | 0.783 |
| 3315 | GG**Tyr**GGG**Gly**GG | 0.832 | 0.882 | 0.867 | 0.813 | 0.834 | 0.860 | 0.847 |
| 3316 | GG**Tyr**GGG**Hip**GG | 0.727 | 0.793 | 0.707 | 0.777 | 0.737 | 0.720 | 0.732 |
| 3317 | GG**Tyr**GGG**His**GG | 0.837 | 0.842 | 0.810 | 0.879 | 0.846 | 0.820 | 0.840 |
| 3318 | GG**Tyr**GGG**Ile**GG | 0.869 | 0.801 | 0.832 | 0.802 | 0.795 | 0.799 | 0.802 |
| 3319 | GG**Tyr**GGG**Leu**GG | 0.805 | 0.847 | 0.796 | 0.851 | 0.851 | 0.861 | 0.849 |
| 3320 | GG**Tyr**GGG**Lys**GG | 0.745 | 0.764 | 0.697 | 0.716 | 0.788 | 0.743 | 0.744 |
| 3321 | GG**Tyr**GGG**Met**GG | 0.808 | 0.593 | 0.788 | 0.551 | 0.817 | 0.607 | 0.698 |
| 3322 | GG**Tyr**GGG**Phe**GG | 0.810 | 0.851 | 0.797 | 0.826 | 0.774 | 0.793 | 0.804 |
| 3323 | GG**Tyr**GGG**Pro**GG | 0.864 | 0.862 | 0.834 | 0.861 | 0.879 | 0.862 | 0.862 |
| 3324 | GG**Tyr**GGG**Ptr**GG | 0.742 | 0.701 | 0.747 | 0.854 | 0.728 | 0.715 | 0.735 |
| 3325 | GG**Tyr**GGG**S1p**GG | 0.802 | 0.825 | 0.755 | 0.839 | 0.750 | 0.753 | 0.778 |
| 3326 | GG**Tyr**GGG**Sep**GG | 0.836 | 0.833 | 0.834 | 0.831 | 0.840 | 0.839 | 0.835 |
| 3327 | GG**Tyr**GGG**Ser**GG | 0.807 | 0.837 | 0.799 | 0.862 | 0.871 | 0.859 | 0.848 |
| 3328 | GG**Tyr**GGG**T1p**GG | 0.864 | 0.806 | 0.831 | 0.788 | 0.877 | 0.711 | 0.819 |
| 3329 | GG**Tyr**GGG**Thr**GG | 0.792 | 0.851 | 0.747 | 0.852 | 0.752 | 0.783 | 0.788 |
| 3330 | GG**Tyr**GGG**Tpo**GG | 0.855 | 0.302 | 0.329 | 0.421 | 0.418 | 0.914 | 0.419 |
| 3331 | GG**Tyr**GGG**Trp**GG | 0.786 | 0.785 | 0.688 | 0.878 | 0.734 | 0.767 | 0.776 |
| 3332 | GG**Tyr**GGG**Tyr**GG | 0.760 | 0.806 | 0.798 | 0.821 | 0.843 | 0.854 | 0.813 |
| 3333 | GG**Tyr**GGG**Val**GG | 0.773 | 0.759 | 0.716 | 0.799 | 0.790 | 0.793 | 0.781 |
| 3334 | GG**Tyr**GGG**Y1p**GG | 0.673 | 0.724 | 0.728 | 0.756 | 0.833 | 0.855 | 0.742 |
| 3335 | GG**Val**GGG**Ala**GG | 0.856 | 0.856 | 0.875 | 0.834 | 0.877 | 0.887 | 0.866 |
| 3336 | GG**Val**GGG**Arg**GG | 0.747 | 0.704 | 0.746 | 0.720 | 0.759 | 0.694 | 0.733 |
| 3337 | GG**Val**GGG**Ash**GG | 0.820 | 0.860 | 0.817 | 0.716 | 0.725 | 0.858 | 0.818 |
| 3338 | GG**Val**GGG**Asn**GG | 0.772 | 0.782 | 0.777 | 0.849 | 0.810 | 0.802 | 0.792 |
| 3339 | GG**Val**GGG**Asp**GG | 0.852 | 0.842 | 0.790 | 0.836 | 0.810 | 0.809 | 0.823 |
| 3340 | GG**Val**GGG**Cys**GG | 0.851 | 0.818 | 0.795 | 0.839 | 0.794 | 0.833 | 0.825 |
| 3341 | GG**Val**GGG**Glh**GG | 0.739 | 0.699 | 0.752 | 0.614 | 0.638 | 0.792 | 0.719 |
| 3342 | GG**Val**GGG**Gln**GG | 0.836 | 0.845 | 0.877 | 0.843 | 0.821 | 0.851 | 0.844 |
| 3343 | GG**Val**GGG**Glu**GG | 0.782 | 0.834 | 0.815 | 0.719 | 0.715 | 0.889 | 0.798 |
| 3344 | GG**Val**GGG**Gly**GG | 0.923 | 0.828 | 0.816 | 0.839 | 0.816 | 0.909 | 0.834 |
| 3345 | GG**Val**GGG**Hip**GG | 0.806 | 0.775 | 0.781 | 0.807 | 0.795 | 0.835 | 0.801 |
| 3346 | GG**Val**GGG**His**GG | 0.872 | 0.821 | 0.851 | 0.862 | 0.887 | 0.832 | 0.856 |
| 3347 | GG**Val**GGG**Ile**GG | 0.817 | 0.831 | 0.776 | 0.815 | 0.838 | 0.796 | 0.816 |
| 3348 | GG**Val**GGG**Leu**GG | 0.867 | 0.839 | 0.806 | 0.889 | 0.840 | 0.827 | 0.839 |
| 3349 | GG**Val**GGG**Lys**GG | 0.817 | 0.734 | 0.661 | 0.734 | 0.651 | 0.619 | 0.697 |

This work is shared under a CC BY 4.0 license.

| | Peptide | $r_{replica}(1,2)$ | $r_{replica}(1,3)$ | $r_{replica}(1,4)$ | $r_{replica}(2,3)$ | $r_{replica}(2,4)$ | $r_{replica}(3,4)$ | Median |
|---|---|---|---|---|---|---|---|---|
| 3350 | GG**Val**GGG**Met**GG | 0.820 | 0.843 | 0.846 | 0.806 | 0.840 | 0.858 | 0.841 |
| 3351 | GG**Val**GGG**Phe**GG | 0.837 | 0.868 | 0.901 | 0.874 | 0.856 | 0.862 | 0.865 |
| 3352 | GG**Val**GGG**Pro**GG | 0.806 | 0.795 | 0.808 | 0.821 | 0.902 | 0.835 | 0.814 |
| 3353 | GG**Val**GGG**Ptr**GG | 0.820 | 0.748 | 0.683 | 0.746 | 0.695 | 0.796 | 0.747 |
| 3354 | GG**Val**GGG**S1p**GG | 0.863 | 0.817 | 0.806 | 0.857 | 0.727 | 0.686 | 0.811 |
| 3355 | GG**Val**GGG**Sep**GG | 0.879 | 0.836 | 0.749 | 0.824 | 0.793 | 0.763 | 0.808 |
| 3356 | GG**Val**GGG**Ser**GG | 0.745 | 0.834 | 0.801 | 0.850 | 0.789 | 0.859 | 0.817 |
| 3357 | GG**Val**GGG**T1p**GG | 0.614 | 0.607 | 0.337 | 0.781 | 0.673 | 0.741 | 0.644 |
| 3358 | GG**Val**GGG**Thr**GG | 0.745 | 0.780 | 0.819 | 0.777 | 0.755 | 0.852 | 0.779 |
| 3359 | GG**Val**GGG**Tpo**GG | 0.644 | 0.660 | 0.763 | 0.347 | 0.784 | 0.440 | 0.652 |
| 3360 | GG**Val**GGG**Trp**GG | 0.623 | 0.612 | 0.780 | 0.685 | 0.695 | 0.752 | 0.690 |
| 3361 | GG**Val**GGG**Tyr**GG | 0.824 | 0.844 | 0.752 | 0.832 | 0.805 | 0.813 | 0.818 |
| 3362 | GG**Val**GGG**Val**GG | 0.808 | 0.827 | 0.843 | 0.856 | 0.818 | 0.866 | 0.835 |
| 3363 | GG**Val**GGG**Y1p**GG | 0.748 | 0.737 | 0.825 | 0.751 | 0.711 | 0.739 | 0.743 |
| 3364 | GG**Y1p**GGG**Ala**GG | 0.816 | 0.711 | 0.732 | 0.796 | 0.859 | 0.828 | 0.806 |
| 3365 | GG**Y1p**GGG**Arg**GG | 0.747 | 0.613 | 0.707 | 0.586 | 0.649 | 0.488 | 0.631 |
| 3366 | GG**Y1p**GGG**Ash**GG | 0.807 | 0.817 | 0.840 | 0.784 | 0.846 | 0.842 | 0.828 |
| 3367 | GG**Y1p**GGG**Asn**GG | 0.692 | 0.742 | 0.689 | 0.817 | 0.708 | 0.751 | 0.725 |
| 3368 | GG**Y1p**GGG**Asp**GG | 0.849 | 0.775 | 0.791 | 0.829 | 0.738 | 0.697 | 0.783 |
| 3369 | GG**Y1p**GGG**Cys**GG | 0.837 | 0.795 | 0.664 | 0.832 | 0.757 | 0.636 | 0.776 |
| 3370 | GG**Y1p**GGG**Glh**GG | 0.741 | 0.729 | 0.692 | 0.816 | 0.792 | 0.779 | 0.760 |
| 3371 | GG**Y1p**GGG**Gln**GG | 0.750 | 0.663 | 0.760 | 0.722 | 0.799 | 0.710 | 0.736 |
| 3372 | GG**Y1p**GGG**Glu**GG | 0.807 | 0.831 | 0.820 | 0.851 | 0.825 | 0.812 | 0.822 |
| 3373 | GG**Y1p**GGG**Gly**GG | 0.922 | 0.890 | 0.930 | 0.887 | 0.920 | 0.905 | 0.913 |
| 3374 | GG**Y1p**GGG**Hip**GG | 0.666 | 0.701 | 0.710 | 0.632 | 0.776 | 0.632 | 0.683 |
| 3375 | GG**Y1p**GGG**His**GG | 0.695 | 0.765 | 0.821 | 0.816 | 0.747 | 0.742 | 0.756 |
| 3376 | GG**Y1p**GGG**Ile**GG | 0.755 | 0.832 | 0.824 | 0.698 | 0.729 | 0.812 | 0.783 |
| 3377 | GG**Y1p**GGG**Leu**GG | 0.651 | 0.691 | 0.731 | 0.657 | 0.803 | 0.778 | 0.711 |
| 3378 | GG**Y1p**GGG**Lys**GG | 0.792 | 0.766 | 0.707 | 0.739 | 0.653 | 0.709 | 0.724 |
| 3379 | GG**Y1p**GGG**Met**GG | 0.809 | 0.743 | 0.831 | 0.650 | 0.732 | 0.727 | 0.738 |
| 3380 | GG**Y1p**GGG**Phe**GG | 0.875 | 0.793 | 0.617 | 0.837 | 0.611 | 0.591 | 0.705 |
| 3381 | GG**Y1p**GGG**Pro**GG | 0.641 | 0.853 | 0.696 | 0.713 | 0.749 | 0.716 | 0.714 |
| 3382 | GG**Y1p**GGG**Ptr**GG | 0.611 | 0.841 | 0.804 | 0.626 | 0.655 | 0.820 | 0.729 |
| 3383 | GG**Y1p**GGG**S1p**GG | 0.878 | 0.810 | 0.802 | 0.840 | 0.814 | 0.785 | 0.812 |
| 3384 | GG**Y1p**GGG**Sep**GG | 0.499 | 0.489 | 0.509 | 0.775 | 0.771 | 0.852 | 0.640 |
| 3385 | GG**Y1p**GGG**Ser**GG | 0.860 | 0.842 | 0.831 | 0.838 | 0.839 | 0.840 | 0.840 |
| 3386 | GG**Y1p**GGG**T1p**GG | 0.351 | 0.317 | 0.630 | 0.666 | 0.712 | 0.675 | 0.648 |
| 3387 | GG**Y1p**GGG**Thr**GG | 0.836 | 0.812 | 0.821 | 0.777 | 0.737 | 0.841 | 0.817 |
| 3388 | GG**Y1p**GGG**Tpo**GG | 0.308 | 0.870 | 0.793 | 0.329 | 0.392 | 0.837 | 0.593 |
| 3389 | GG**Y1p**GGG**Trp**GG | 0.637 | 0.738 | 0.604 | 0.803 | 0.830 | 0.783 | 0.761 |
| 3390 | GG**Y1p**GGG**Tyr**GG | 0.724 | 0.746 | 0.696 | 0.796 | 0.841 | 0.767 | 0.757 |
| 3391 | GG**Y1p**GGG**Val**GG | 0.706 | 0.834 | 0.635 | 0.710 | 0.595 | 0.627 | 0.671 |
| 3392 | GG**Y1p**GGG**Y1p**GG | 0.607 | 0.800 | 0.838 | 0.643 | 0.719 | 0.822 | 0.760 |

This work is shared under a CC BY 4.0 license.
